# Identification of a short, single site matriglycan that maintains neuromuscular function in the mouse

**DOI:** 10.1101/2023.12.20.572361

**Authors:** Tiandi Yang, Ishita Chandel, Miguel Gonzales, Hidehiko Okuma, Sally J. Prouty, Sanam Zarei, Soumya Joseph, Keith W. Garringer, Saul Ocampo Landa, Takahiro Yonekawa, Ameya S. Walimbe, David P. Venzke, Mary E. Anderson, Jeffery M. Hord, Kevin P. Campbell

## Abstract

Matriglycan (−1,3-β-glucuronic acid-1,3-α-xylose-) is a polysaccharide that is synthesized on α-dystroglycan, where it functions as a high-affinity glycan receptor for extracellular proteins, such as laminin, perlecan and agrin, thus anchoring the plasma membrane to the extracellular matrix. This biological activity is closely associated with the size of matriglycan. Using high-resolution mass spectrometry and site-specific mutant mice, we show for the first time that matriglycan on the T317/T319 and T379 sites of α-dystroglycan are not identical. T379-linked matriglycan is shorter than the previously characterized T317/T319-linked matriglycan, although it maintains its laminin binding capacity. Transgenic mice with only the shorter T379-linked matriglycan exhibited mild embryonic lethality, but those that survived were healthy. The shorter T379-linked matriglycan exists in multiple tissues and maintains neuromuscular function in adult mice. In addition, the genetic transfer of α-dystroglycan carrying just the short matriglycan restored grip strength and protected skeletal muscle from eccentric contraction-induced damage in muscle-specific dystroglycan knock-out mice. Due to the effects that matriglycan imparts on the extracellular proteome and its ability to modulate cell-matrix interactions, our work suggests that differential regulation of matriglycan length in various tissues optimizes the extracellular environment for unique cell types.

## INTRODUCTION

Glycosylation is carbohydrate modification found on biological macromolecules including nucleotides (*1*), lipids (*2*, *3*) and especially proteins (*4*, *5*). Protein glycosylation has been the target of extensive research efforts because glycan modifications not only modulate the functions of the proteins but also act as receptors/ligands for their lectin counterparts. The glycan-lectin interactions are associated with numerous biological processes, such as quality control of protein folding (*6*), cellular growth and migration (*7*), host-pathogen/microbiome interaction (*8*), transmembrane signaling (*9*), and more. One of the best functionally characterized glycan receptors is a unique matriglycan found on dystroglycan (DG). DG is encoded by the DAG1 gene (*10*) which is widely expressed in different tissues and is conserved among various organisms. DAG1 is transcribed into a single mRNA, but the translated DG polypeptide post-translationally cleaves into DG N-terminus (DG-Nt) (*11*), α- and β-DG (*10*). The α subunit has a heavily *O*-glycosylated mucin-like domain that harbors the matriglycan, and a C-terminal domain (DG-Ct) that forms a complex with the transmembrane β-DG. Matriglycan serves as the carbohydrate receptor for extracellular matrix (ECM) proteins containing laminin G-like (LG) domain (*12*, *13*), such as laminin (*10*, *14*), agrin (*15*), and perlecan (*12*); while α- and β-DG are sub-units of a dystrophin-glycoprotein complex (DGC) (*16*) that anchors to cytoskeletal actin fibers (**Fig. 1a**). Therefore, matriglycan plays a central role of bridging the ECM to the plasma membrane, thus mediating cell-matrix interactions in different tissues including skeletal muscle and the central nervous system.

**Fig. 1.**
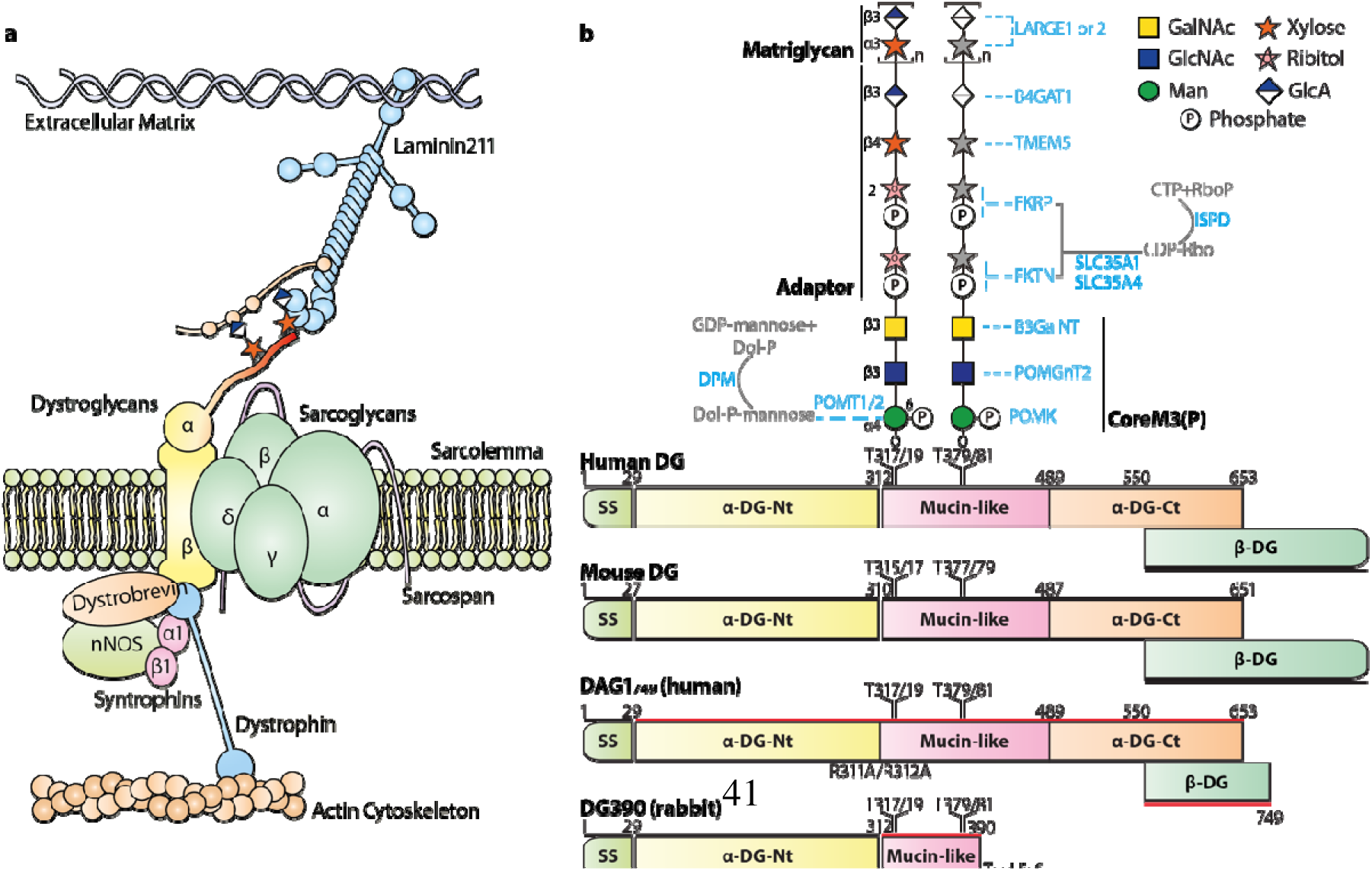
Dystroglycan complex. (**a**) A laminin-matriglycan-dystroglycan complex-dystrophin-actin axis plays a central role in bridging the plasma membrane and extracellular matrix. (**b**) The dystroglycan polypeptide chain auto-cleaves into DG N-terminus (DG-Nt), α- and β-DG. α-DG has two domains, a mucin-like domain presenting the matriglycan and a C-terminal domain (α-DG-Ct) binding to β-DG. α-DG is highly conserved in different organisms. The “most prominent” difference between human and mice is the two amino acid difference between their signal sequences (SS). Matriglycosylation requires a cooperation of multiple enzymes (colored blue). Previous research revealed two TPT motifs on α-DG can carry coreM3 *O*-glycan. The matriglycan links on the T317T319 motif coreM3 via a ribitol-phosphate containing adaptor. It was not known if the T379T381 coreM3 is extended into the matriglycan.

Matriglycan is a polymer of [1,3-β-glucuronic acid (GlcA)-1,3-α-xylose (Xyl)-] repeating disaccharide (*17*) and its biological activities almost solely depend on its length: longer matriglycans exhibit drastically higher affinity to laminin than shorter ones (*18*). Matriglycan is synthesized on a β-GlcA-1,4-β-Xyl-1,2-ribitol-1-phosphate (RboP)-5-RboP adaptor (*19–24*) (**Fig. 1b**) by the bi-functional glycosyltransferases LARGE1 or LARGE2 (*17*, *25–27*). The adaptor is attached to phosphorylated coreM3 (*28–32*) [coreM3(P)] forming the matriglycan precursor. Multiple enzymes are required for proper matriglycosylation, and an interruption of the process leads to a group of disorders in muscle and the central nervous systems, which are called α-dystroglycanopathies (*33*). α-dystroglycanopathy can range in phenotypic expression from Limb-Girdle Muscular Dystrophy (LGMD) to Congenital Muscular Dystrophy to Muscle Eye Brain Disease and Walker-Warburg Syndrome (WWS) in order of increasing disease severity. Disease severity hinges on the ability of matriglycan to bind ECM ligands which in turn depends on the expression and length of matriglycan. A complete absence of matriglycan leads to mammalian embryonic lethality. Normally, dystroglycanopathy patients carry matriglycan, but they are drastically reduced and insufficient to maintain normal cell-matrix interactions (*34*, *35*)

The mucin-like domain of α-DG that harbors the matriglycan contains roughly 50 potential *O*-glycosylation sites, but the coreM3(P) was merely found on three threonine residues (T317, T319, and T379), and matriglycan was only found on a T317/T319 motif before (*23*, *24*, *32*, *36*, *37*). It was reported that T379 can receive LARGE-mediated matriglycosylation under the LARGE over-expression condition (*38*). In the present work, we established an ultra-high resolution and sensitivity mass spectrometric method mapping matriglycosylation sites of α-DG. We confirmed T379 as a secondary matriglycosylation site harboring a short matriglycan without LARGE over-expression and existing in multiple organs. Although the short matriglycan led to embryonic lethality in mice, it appeared to be completely functional in adult skeletal muscle. Simultaneous mutations of the two matriglycosylation sites led to muscular pathology, while neuromuscular junction (NMJ) integrity required modification at both matriglycan sites. In addition, cellular proteomics indicated that matriglycan can modulate the ECM proteome possibly by altering its size or switching matriglycan sites.

## RESULTS

### Matriglycosylation site(s) on α-DG

Our research started with the question of whether there are more than one matriglycan motif on α-DG. We infected HEK293 cells with an adenoviral vector expressing Fc-tagged α-DG (DG-Fc) protein with or without T317A/T319A site-directed mutations. A laminin overlay assay strongly suggests the T317A/T319A mutation fails to eliminate laminin binding (**Fig. s1a**). Because DG-Fc showed significantly lower electrophoretic migration, we performed laminin overlay guided in-gel digestion and mass spectrometry (MS) based proteomics to verify DG-Fc as the main protein in the 250 KDa bands (**Table s1** and **2**).

These data indicated additional matriglycosylation site(s) without LARGE over-expression and drove us to establish a nano-flow liquid chromatography (nanoLC) MS method to map matriglycosylation sites. Matriglycosylated peptides are extraordinarily challenging targets for electrospray ionization (ESI) MS, and to date, there are no published ESI-MS data identifying matriglycosylated peptides. We expressed a DAG1_749_ recombinant protein in HEK293F cells, which mimics the extracellular region of human pro-dystroglycan with R311A/R312A mutations (**Fig. 1b**, amino acid 28-749, DG-Nt attached). *O*-glycomics showed the protein carried multiple mucin-type *O*-glycans and *O*-mannose glycans, indicating proper glycosylation and secretion by our cell model (**Fig. s1b**). We then used a glycosidase mixture (β-hexosaminidase, β-1,4-galactosidase, fucosidase, sialidase, and *O*-glycosidase, see Materials and Methods) to reduce the sample complexity caused by numerous glycoforms. Previous analytical efforts normally introduced a lysine residue for Lys-C digestion so that truncated matriglycopeptides can be enriched by ion exchange chromatography and characterized by matrix-laser desorption/ionization time-of-flight (MALDI-TOF) MS (*23*). A similar strategy was used to collect high-resolution ESI-Orbitrap data on truncated DG glycopeptides carrying the precursor of matriglycan (*24*). Our approach keeps longer peptide backbones to generate comprehensive information on *O*-mannosylation and to facilitate electrospray ionization for ESI-Orbitrap to achieve ultra-high resolution and sensitivity. The addition of RboP and GlcA-Xyl residues was shown to delay glycopeptides LC elution (*24*). Therefore, we eluted the LC column with a more concentrated acetonitrile solution to compensate for the possible ion-paring effect.

Our optimized MS strategy detected the canonical T317/T319 motif containing glycopeptide with an amino acid sequence of VAAQIHA**<u>T</u>**P**<u>T</u>**PVTAIGPPTTAIQEPPSR (note the R to A mutations in **Fig. s1c-f**). This glycopeptide carries the matriglycan precursor, five mannose, and one hexNAc, giving rise to an MS signal at *m/z* 1305.55^4+^. We were able to get a glycopeptide with a nearly native sequence analyzed, and our data are consistent with previous research identifying T317/T319 as a matriglycan motif. We were also able to detect matriglycan on the secondary T379/T381 site in a glycopeptide with a sequence of GAIIQ**<u>T</u>**P**<u>T</u>**LGPIQPTR. This is supported by two MS signals at *m/z* 1124.46^3+^ and 1227.15^3+^ which correspond to the glycopeptides carrying the two mannoses plus the matriglycan precursor and one matriglycan disaccharide elongation, respectively (**Fig. s1g-m**). This is the first time that a matriglycosylated peptide has been successfully analyzed by nanoLC-ESI-MS, and our data indicate all three previously mapped coreM3 can be elongated into matriglycans under normal physiological conditions.

### Confirmation of matriglycosylation on T317/T319 and T379/T381 two motifs

As crystallographic research has revealed that matriglycans with two disaccharide repeats strongly bind to the laminin LG4 domain (*13*), we next sought to better characterize matriglycosylated peptides for both motifs. The excessive peptide and glycopeptide MS signals detected in the DAG1_749_ sample could have suppressed MS signals of matriglycosylated peptides, so we expressed a shorter His-tagged DG390 protein with HEK293F cells. DG390 is designed based on the rabbit DAG1 sequence (see **Fig 1b. native rbt DG**), covering both matriglycan motifs identified on DAG1_749_. DG390 was also subjected to *O*-glycomic analysis, and the detected mucin-type and *O*-mannose *O*-glycans proved this much shorter construct was heavily glycosylated (**Fig. s2a** and **b**). In addition, MALDI-TOF also detected sodiated permethylated (GlcA-Xyl)_n_-Rbo glycan released by mild hydrofluoric acid (HF) hydrolysis from DG390 (**Fig. s2c**). The MS detected matriglycan repeats up to two times, and the MS peak intensities of the matriglycan decreased as the matriglycan got longer. The longer matriglycans are expected to show even lower intensities and might not be detected because of a reduced sensitivity when extensive chemical reactions are implemented before MS analysis. Subsequent MALDI-TOF/TOF analysis of the glycans confirmed their sharing of a linear architecture (**Fig. 2d-f**).

**Fig. 2.**
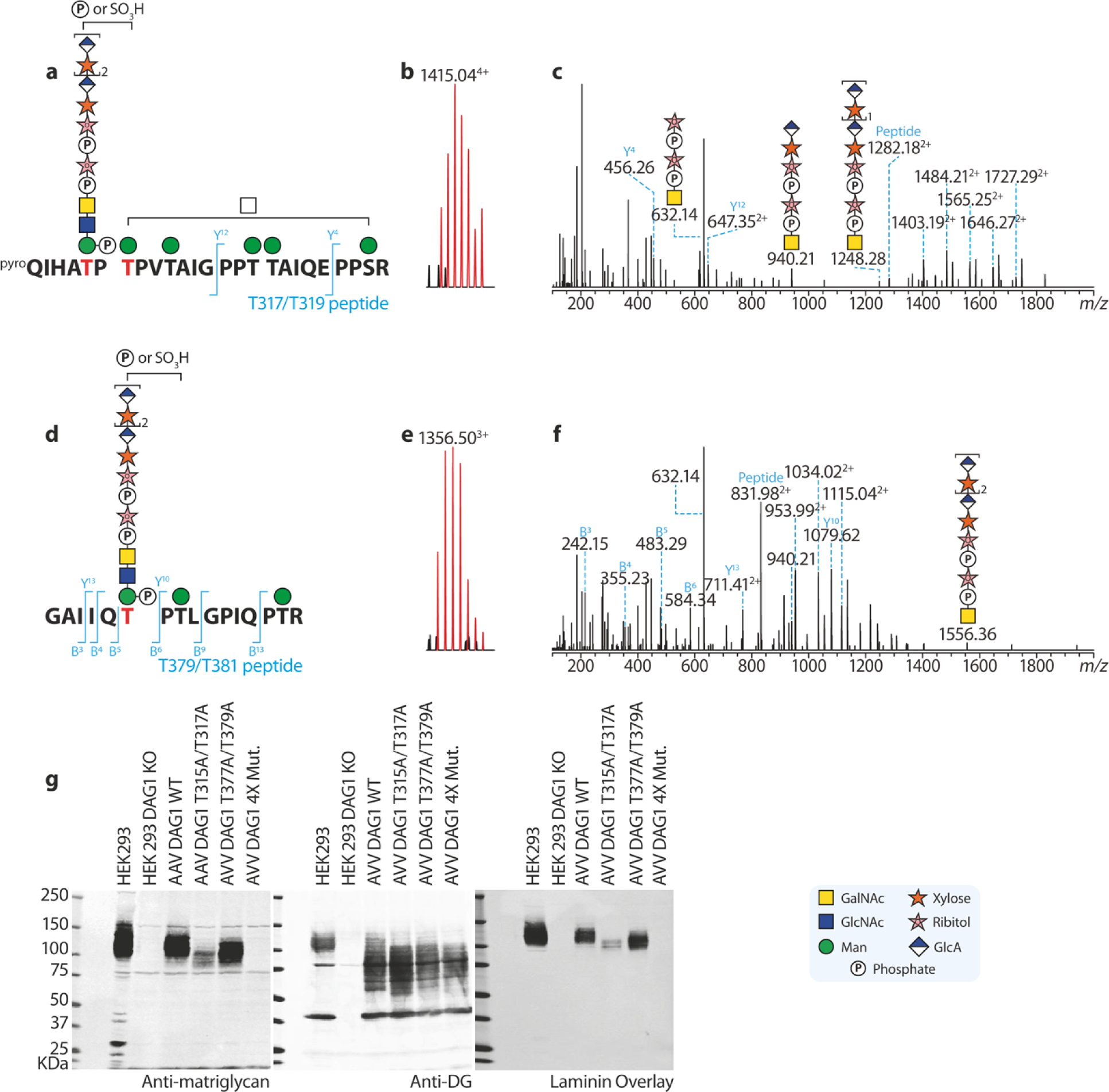
Mapping of T317T319 motif (a-c) and T379T381 (d-f) matriglycan motifs. (**a**) and (**d**) Diagrams showing structures of two matriglycosylated peptides. (**b**) and (**e**) MS signals of the glycopeptides in (**a**) and (**d**). (**c**) and (**f**) MS/MS spectra of the correspondence MS signals. Both peptide and matriglycan fragment ions were detected, confirming the glycopeptides carries matriglycans. There are only two matriglycan motifs detectable by MS. (**g**) Immunoblotting and laminin overlay assays supporting the exitance of only two matriglycan motifs in HEK cells. For immunoblotting, antibodies against both matriglycan and DG backbone were used. The blots indicate the T379T381 motif carries a shorter matriglycan than the T317T319 motif.

The DG390 was enzymatically de-glycosylated (**Fig. s2b**), tryptically digested, and subjected to nanoLC-ESI-MS-based glycoproteomics. A representative MS peak at *m/z* 1415.04^4+^ (**Fig. 2b**) corresponds to a T317/T319 glycopeptide with a sequence of QIHATPTPVTAIGPPTTAIQEPPSR carrying two matriglycan repeats on the precursor, five hexoses, one hexNAc, and an extra phosphate or sulfate (**Fig. 2a** and **Fig. s3**). As DG390 does not bear any mutation, the peptide backbone is slightly shorter than its counterpart found in DAG1_749_. The glycan composition indicates all serine and threonine residues are modifiable by *O*-mannoses, and one mannose in the T317/T319 motif is further synthesized to mature matriglycans (**Fig. 2a**). Further MS/MS analysis confirmed our structural annotation (**Fig. 2c**). First, we found fragment ions at *m/z* 632.14^+^, 940.21^+^ and 1248.28^+^ showing the matriglycan assembled on the RboP-containing adaptor. Secondly, the molecular ion of the peptide backbone was detected at *m/z* 1282.18^2+^, and two peptide fragment ions were observed at *m/z* 456.26^+^ (y4) and 647.35^2+^ (y12). An addition of mannose-phosphate (242.02 Da) shifts the peptide molecular ion from 1282.18^2+^ to 1403.19^2+^, to which more mannose is added, forming glycopeptide ions with *m/z* of 1484.21^2+^, 1565.25^2+^, 1646.27^2+^, and 1727.29^2+^. This proves at least five out of six *O*-glycosylation sites in the peptides are *O*-mannosylated. No fragment ion corresponds to a HexNAc addition to the peptide backbone, indicating all six *O*-glycosylation sites are *O*-mannosylated.

The T379/T381 motif is in the GAIIQ**<u>T</u>**P**<u>T</u>**LGPIQPTR glycopeptide (also see **Fig. s4**), which is the same as in DAG1_749_. In addition to the MS signals at *m/z* 1124.46^3+^ and 1227.15^3+^, we were able to find a representative signal at *m/z* 1356.50^3+^ (**Fig. 2e**) corresponding to the peptide with two matriglycan repeats on the precursor with two hexoses, one HexNAc and an extra phosphate or sulphate (**Fig. 2d**). The glycan composition again suggests that all three *O*-glycosylation sites are modified by *O*-mannoses and one of them was biosynthesized into the matriglycan. MS/MS analysis (**Fig. 2f**) detected matriglycan fragments at *m/z* 632.14^+^, 940.21^+^, and 1556.36^+^, and the molecular ion of the peptide backbone at *m/z* 831.98^2+^. The peptide is partially sequenced by fragment ions at *m/z* 242.15^+^ (b3), 355.23^+^ (b4), 483.29^+^ (b5), and 584.34^+^ (b6), and y ions at m/z 1079.62^+^ (y10) and 711.41^2+^ (y13). A combination of the presence of fragment ions at *m/z* 952.99^2+^ (addition of a ManP), 1034.02^2+^ (addition of man, HexP), and 1115.04^2+^ (addition of 2Man, ManP) and absence of 933.52^2+^ ion further confirms all three potential *O*-glycosylation sites are *O*-mannosylated.

To verify our MS observation that there are two matriglycan motifs among the roughly 50 *O*-glycosylation sites, we infected DG KO HEK293 cells with adenoviral vectors encoding WT DG and DG-bearing T-to-A mutations in the MS-characterized matriglycan motifs (**Fig. 2g**). When both motifs get mutated (4X mut.), no matriglycan was detected by both immunoblots and laminin overlay despite an over-expression of dystroglycan backbone. This confirms there are only two matriglycan motifs on α-DG. α-DG with T317/T319 matriglycan exhibited lower electrophoretic migration in polyacrylamide gels than with T379/T381 matriglycan, indicating the former matriglycan is considerably longer than the latter when the protein is expressed in HEK293F cells.

### T379 carries a matriglycan repeating for at least nine disaccharide units

Previous research has revealed both T317 and T319 can carry matriglycan (*37*). To pinpoint the exact site for matriglycosylation in the T379/T381 motif, we performed electron-transfer dissociation (ETD) on the T379/T381 glycopeptides (**Fig. 3a-c**). We found a series of C7 ions at *m/z* 1346.60^+^, 1560.62^+^, and 1774.63^+^, proving the T379 residue carries a coreM3(P) that can be synthesized towards matriglycan. These MS spectra determine T379 as a major carrier of the matriglycan in the motif.

**Fig. 3.**
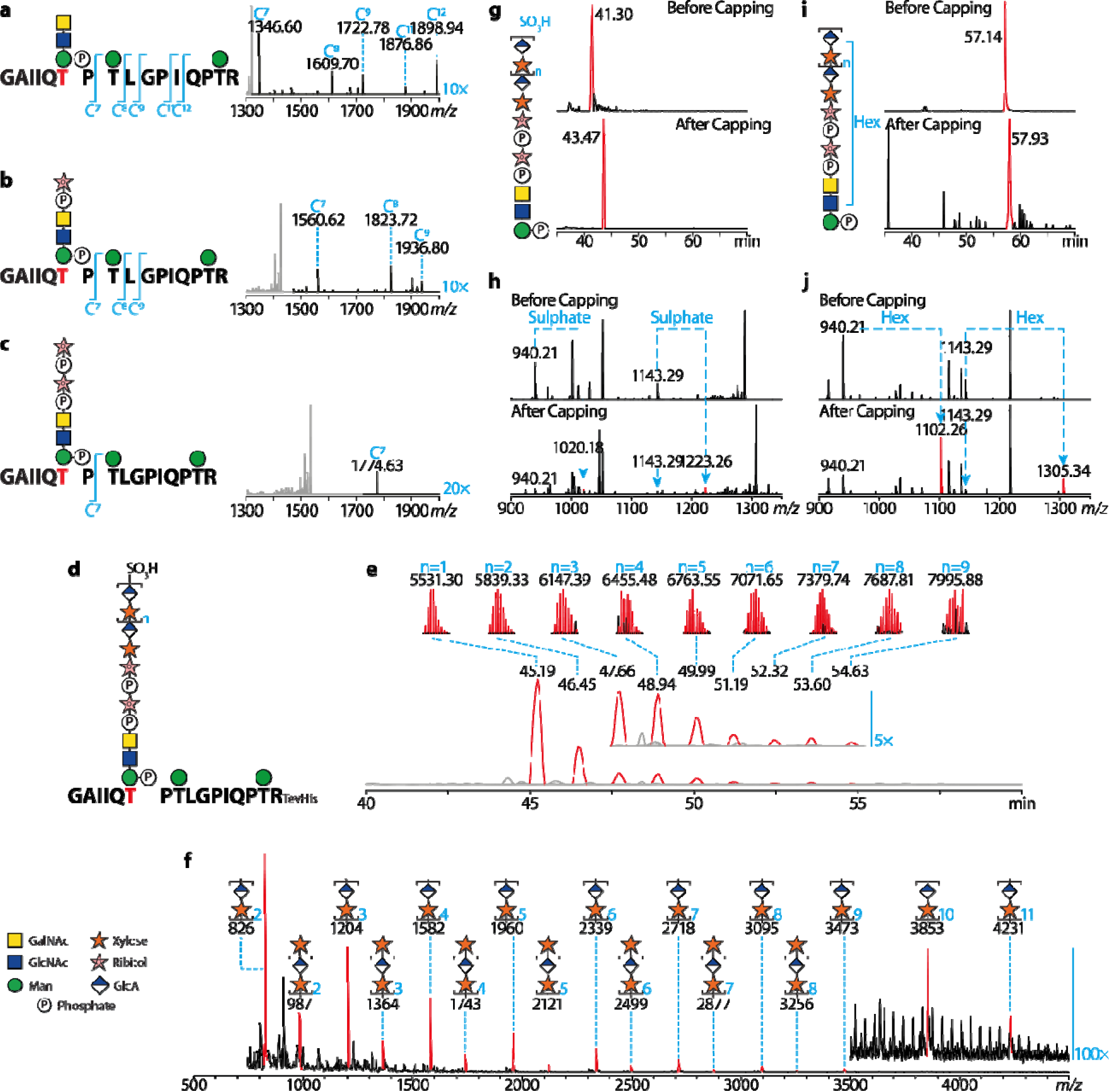
MS investigation of the T379 matriglycan. Electron transfer dissociation (ETD) confirming T379 as the main carrier of the matriglycan in the T377T379 motif (**a-c**). Fragmentation of the peptide backbones are annotated at the left of the correspondence MS/MS spectra. Because of the low ETD efficiency, black regions of the spectra are amplified by 10 or 20-fold. MS detected the T379 matriglycan repeats for up to nine times (**d-f**). (d) A diagram of T379 glycopeptides carrying extended matriglycans. (**e**) Overlayed XICs of MS signals on T379 glycopeptides carrying matriglycans with different disaccharide unit numbers. The XIC peak are annotated with retention times, MS signals of the glycopeptide ions, molecular weights of the ions and disaccharide repeating numbers (colored blue). Note the heights of the XIC peaks are normalized to the same level and they do not represent relative intensities. (**f**) LARGE1 assay *in vitro*. MALDI-TOF profile of permethylated LARGE1 products are shown. MS signal corresponding to sodiated permethylated matriglycans are colored red and annotated with glycan structures. Higher molecular weight region of the spectrum is amplified by 100-fold. The matriglycans can be capped by a sulphate group (**g and h**) or hexose residue (**i and j**). The XIC peaks shown in (**g**) and (**h**) are colored red and annotated with retention times. EThcD spectra detected the sulphate (**i**) and hexose (**j**) caps on the *m/z* 940.21 fragment ions (structure annotated in Fig. 1c).

We next measured the length of matriglycan on the T379/T381 motif. Note that we deliberately attached a His-tag to DG390 when designing it, and this plays a critical role in the nanoLC-MS analysis of longer matriglycans. The poly-histidine tag has two functions: first, it normalizes the nanoLC retention of matriglycosylated peptides (**Fig. s5c** and **d**) and second, it greatly facilitates positive ESI of matriglycosylated peptides (**Fig. s5e** and **f**). We found the T379/T381 motif carrying matriglycans up to nine GlcA-Xyl repeats (**Fig. 3d-e**) and the biggest observed matriglycosylated peptide had a molecular weight of around 8 KDa. Structural annotations of the dominant MS signals were verified by MS/MS analysis (**Fig. s5g-j**), but some MS intensities for longer matriglycan get too low to analyze.

Consistent with matriglycans released by mild HF hydrolysis, deconvoluted MS spectra strongly suggested MS intensities of matriglycosylated peptides drop rapidly as the matriglycan length increases (**Fig. s5k**). This phenomenon is also observed with our matriglycan *in vitro* biosynthesis assay (**Fig. 3f**). The synthesized matriglycan can only get up to 10 repeats after 48 hours of incubation with excessive LARGE1, and such length is comparable with our DG390 results. MALDI-TOF detected matriglycans terminated by both GlcA and xylose, but GlcA-terminated matriglycans are considerably more abundant than xylose-terminated ones. In addition, MS intensities of the matriglycans decrease as they get longer just like the T379 matriglycan synthesized in HEK293 cells.

Collectively, the data suggest that matriglycan synthesized on T379 in HEK293 cells is roughly nine to ten units long; and a longer matriglycan corresponds to the lower amount. Matriglycan synthesis could be rate-determined by the xylosyltransferase activity. Previous MALDI-TOF analysis indicated that T317/T319 matriglycan repeats 17 times (*23*), which is considerably longer than the T379 matriglycan that is characterized in this research. This means the two matriglycan motifs are not necessarily functionally identical, as the matriglycan length is important for its receptor binding affinity.

### Termination of matriglycan elongation can be achieved by sulphate and hexose capping

MS signals for both T317/T319 and T379/T381 glycopeptides imply an extra modification of roughly 79.96-79.97 Da mass (**Fig. 2 a-f**), which matches the mass of a sulphate or phosphate group. This group can be found on all dominant polymeric matriglycans on the T379/T381 motif (**Fig. 3d** and **e**), although uncapped polymeric matriglycans were also observed at considerably lower intensities. We luckily found a single MS/MS spectrum showing the 79.97 Da modification directly linked to the matriglycan fragments (**Fig. 3g** and **h**). The extremely labile modification was kept on glycans most likely due to the long peptide backbone dispersing the fragmentation energy during the MS/MS processes. Our MS analysis cannot directly determine if the group is a sulphate or phosphate, but we attributed the extra mass to a sulphate cap because of two reasons. First, previous research identified HNK-1 as a sulfotransferase that caps the matriglycan (*38*, *39*). Secondly, MS/MS analysis of matriglycosylated peptides with the extra 79.97 Da did not detect fragment ions corresponding to glycopeptides carrying two mannose-phosphate (**Fig. s5i**) that were easily detected in non-matriglycosylated peptides (**Fig. s5n-o**). In addition to the presumable sulphate caps, we found a small fraction of the matriglycan precursors are capped by a hexose residue. We are not able to determine if the hexose modification is a consequence of enzymatic catalysis or chemical ligation. However, we only found the matriglycan precursor capped by the hexose, indicating that the hexose cap can compete with LARGE to prevent the matriglycan biosynthesis from the very beginning if the process is enzymatically catalyzed.

### SLC35A1 serves as a RboP transporter

The characterization of the T379/T381 matriglycosylation provided us with a platform to explore the functions of all genes involved in matriglycan biosynthesis. Since the T379/T381 matriglycopeptides exist in far simpler glycoforms than the T317/T319 glycopeptides, we can use a series of representative MS signals at *m/z* 879.41^3+^, 950.76^3+^, 1022.10^3+^, 1124.46^3+^, and 1227.15^3+^ (their extracted ion chromatograms in **Fig. s6a**) to monitor glycan structures along the matriglycan biosynthesis in cell models. Therefore, we applied the MS strategy to characterize two novel pathological genes, *SLC35A1* encoding a CMP-sialic acid transporter, and *POMGnT1* encoding an *O*-mannose β-1,2-GlcNAc transferase. Both activities are not directly associated with the matriglycosylation pathway, but their mutations have previously been reported to cause human muscular disorders resembling dystroglycanopathies (*40–42*).

We first tried to knockout (KO) *SLC35A1* in our HEK293F bioreactor (**Fig. s6b**) with a CRISPR/Cas9 system. Although sialylation is significantly reduced, we were still able to glycomically observe sialylated N-glycans. Further analysis indicated nearly all sialylation is absent on the secreted DG390 (**Fig. s6c**), whereas a highly intense MS signal is present for the glycopeptide carrying the precursor of matriglycan (**Fig. s6d**). To eliminate the possible effects of residual activity of SLC35A1 and/or other similar-function transporters in the HEK cells, we infected HAP1 *SLC35A1* KO cells (**Fig. s6e**) with an adenoviral vector expressing DG390 and were able to detect the matriglycosylation pathway halted before RboP (FKTN activity) by MS (**Fig. 4a)**, as the relative intensity of one and two RboP containing glycopeptides reduced by about 70% and 97%, respectively. Only a trace amount of matriglycan precursor was observed and matriglycan was nearly undetectable (**Fig. s6f** and **g**). This is consistent with immuno- and laminin blots, which also support a sharp downregulation of matriglycan (**Fig. 4b**). The *SLC35A1* and *ISPD* (encoding a CDP-ribitol synthase) gene transfer can rescue the phenotype independently (**Fig. 4c**), indicating *SLC35A1* could be involved in CDP-Rbo transport. These observations provide MS evidence for recently published research showing *SLC35A1* is involved in the translocation of RboP (*43*).

**Fig. 4.**
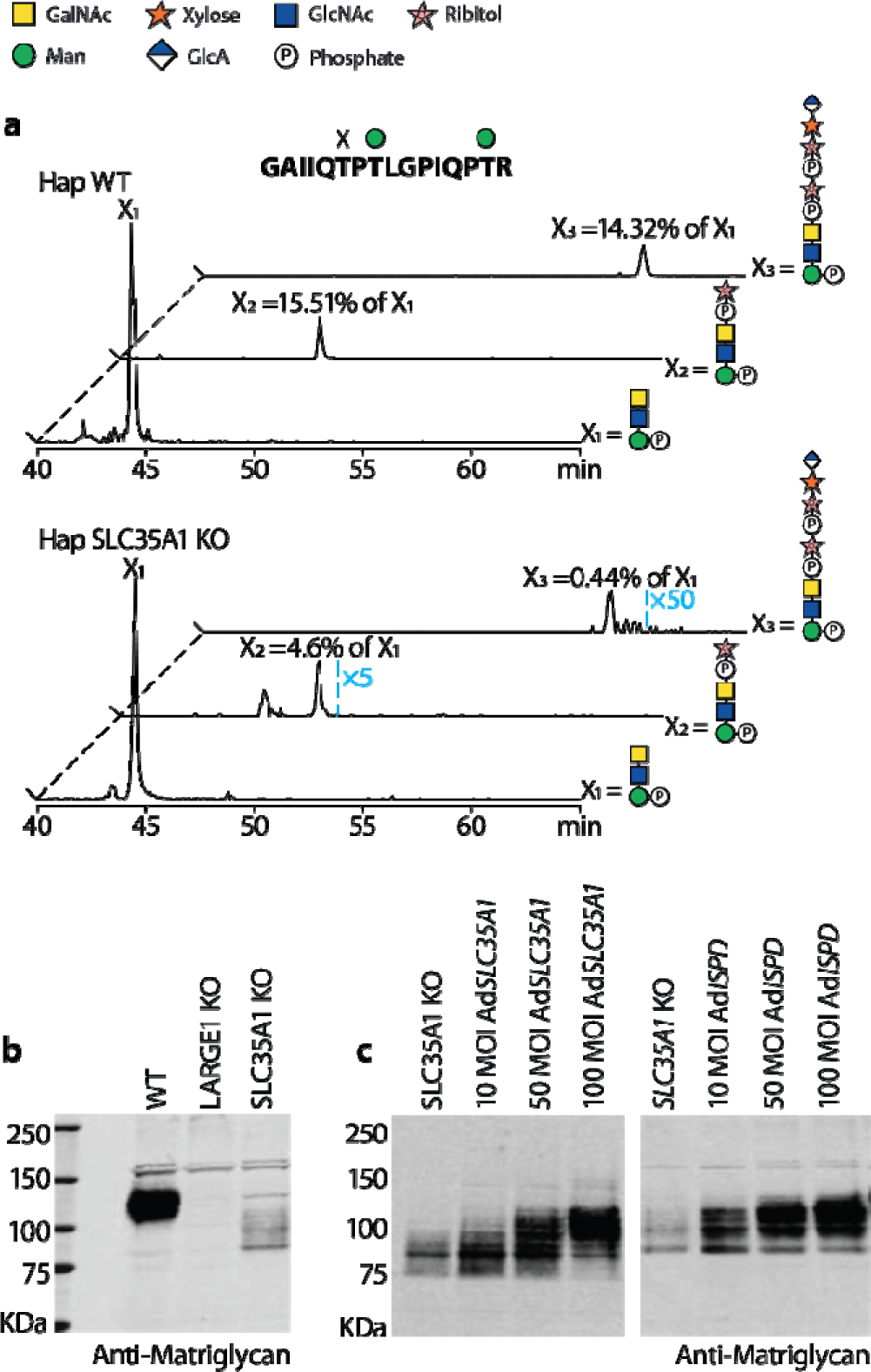
SLC35A1 functions as a ribitol-phosphate transporter. (**a**) Extracted ion chromatograms of the T379T381 glycopeptides containing coreM3 (X_1_), RboP-coreM3 (X_2_) and (RboP)_2_-coreM3 (X_3_) in WT (upper panel) and SLC35A1 KO (lower panel) haploid (Hap) cells. The RboP containing peaks (X_2_ and X_3_) in SLC35A1 KO cells are amplified by 5 and 50 times, and the roughly 70% and 97% reduction in their intensity relative to the coreM3 (X_1_) peak strongly indicates the biosynthesis of the matriglycan halted at the step of RboP addition. The relative intensities are the calculated ratios between the integrated area under the RboP containing peaks and the coreM3 peak. Immunoblotting indicates the SLC35A1 KO leads to a dramatic downregulation of matriglycan (**b**), adenoviral vectors encoding SLC35A1 or ISPD can rescue the matriglycan phenotype (**c**).

We next infected HAP1 *POMGnT1* KO cells with the same viral vector and detected all the representative ions of matriglycosylation. The MS signal at *m/z* 1124.45^3+^ corresponding to the T379 glycopeptides carrying the precursor of matriglycosylation is very strong. The MS signal at *m/z* 1227.15^3+^ characteristic of matriglycosylation is low (**Fig. s6g** and **h**), but it unambiguously proves the existence of matriglycan. This is partially contrary to previously reported immunoblotting results (*44*), where no or very little matriglycan signal was observed.

### Both matriglycans exist and are functional in vivo

After the comprehensive MS analysis of matriglycosylation based on cell models, we set out to create transgenic mice carrying T315A/T317A and T377A/T379A mutations for further interrogating the MS-determined matriglycan motifs *in vivo*. Note that murine DG is two amino acids shorter within its sequence signal than the human and rabbit versions (**Fig. 1b**), so the amino acid number is different. Despite both DG T315A/T317A and T377A/T379A mice are viable, the former mice suffer mild embryonic lethality: of 220 liveborn mice only 22 were T315A/T317A homozygous, giving a birth rate of 10% that fails the χ2 Goodness of Fit Test (**Table s3** and **4**); while the rate was 22% for T377A/T379A homogeneous mice (53 out of 239). Considering *Large^myd^/Large^myd^*(*myd*) mice also show a reduced birth rate, our observation indicates the long T315/T317 matriglycan is important for mice reproduction and/or embryonic development. We were unable to generate live mice carrying both T315A/T317A and T377A/T379A mutations (4X mut.). This is not surprising because the 4X mutation in theory eliminates functions of both LARGE1 and 2, and this observation supports there are only two matriglycan motifs *in vivo*.

We first tested for matriglycans’ presence and function in skeletal muscle, in our various mouse models. Being consistent with the cell models, immunoblotting confirmed matriglycosylation in skeletal muscle of both T315A/T317A and T377A/T379A mice (**Fig. 5c**), and the former motif carries a longer matriglycan (**Fig. 5a**). Immunofluorescence and hematoxylin and eosin (H&E) staining further supported that matriglycans exist in both mice models. Skeletal muscle from both mouse models shows normal morphology (**Fig. 5b**) but losing one matriglycan site seems to slightly reduce fiber size (**Fig. 5c**). When compared to WT controls, extensor digitorum longus (EDL) from both mouse models shows similar absolute and specific tentative forces (**Fig. 5d** and **f**) and similar susceptibility to injuries induced by eccentric contractions (ECCs, **Fig. 5g**), which indicates a single matriglycan, even relatively short, is sufficient to maintain healthy muscle in adult mice. The only observed difference is slightly altered cross-sectional areas (CSA) in the T315A/T317A EDL muscle (**Fig. 5e**).

**Fig. 5.**
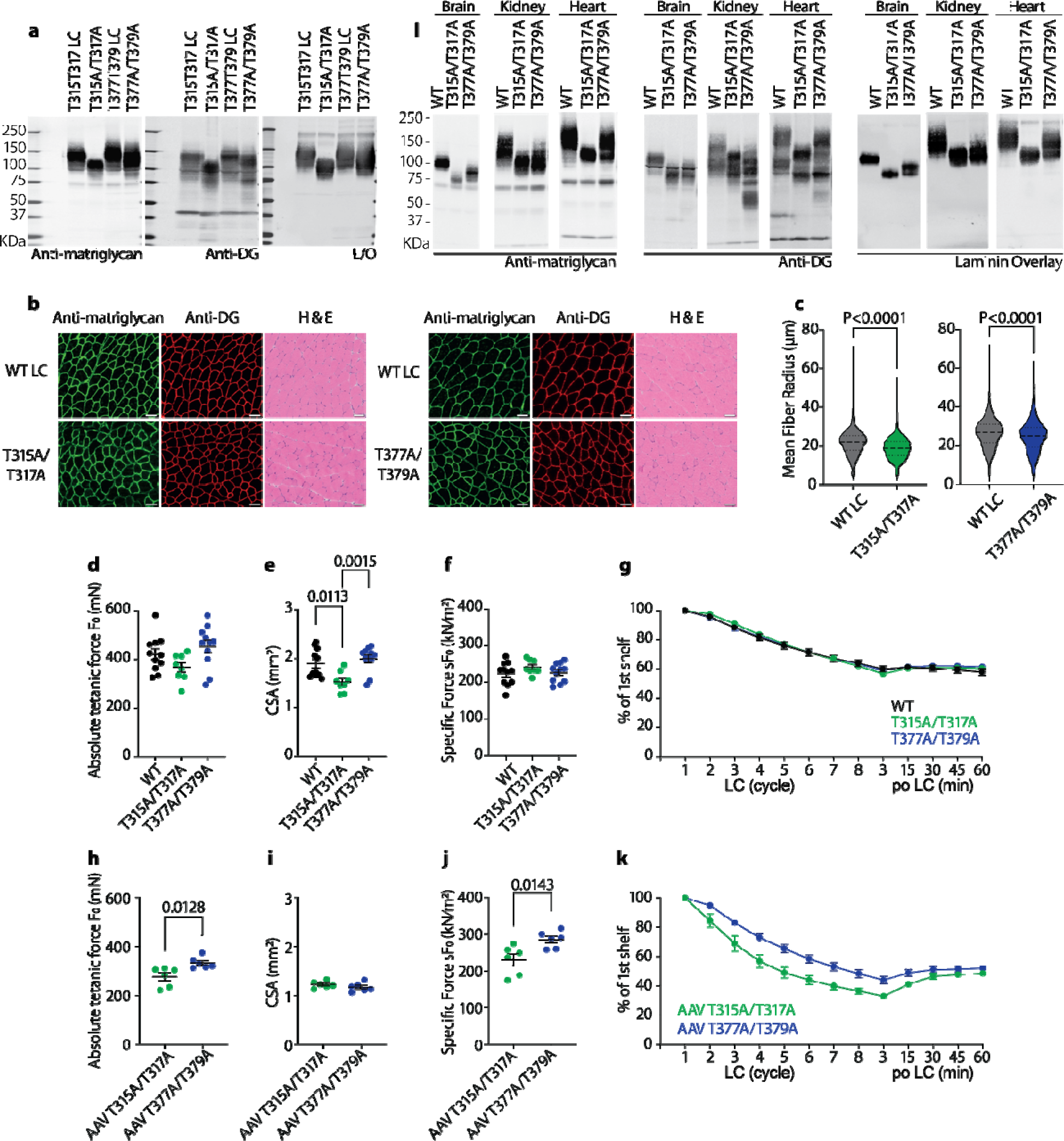
Both MS determined matriglycan motifs exist and functional *in vivo*. (**a**) Immunoblotting and laminin assays of the WGA-enriched skeletal muscle suggest both motifs can be modified by the matriglycan and the T377T379 matriglycan is shorter in skeletal muscle. (**b**) Both of the transgenic mice show no apparent muscle pathology as proved by IF and H&E stain, indicating both are functionally sufficient to maintain a healthy muscle. (**c**) Fiber size i slightly reduced in skeletal muscle from mice with a single matriglycan compared to their litter mates. (**d-g**) Extensor digitorum longus (EDL) from mice with a single matriglycan showed similar absolute tetanic (F_0_) and specific (sF_0_) forces and injuries caused eccentric contraction (ECC) to WT mice. The only observed difference is the slightly altered cross-sectional area (CSA). (**h-k**) DAG1KO muscle rescued by AAV T315A/T317A DAG1 has similar CSA (**i**) but slightly lower F_0_ (**h**) and sF_0_ (**j**) compared to that rescued by AAV T377A/T379A DAG1. The former muscle is also more susceptible to ECC-induced injuries (**k**). (**l**) The two motifs also exist in tissues beyond muscle as showed by the immunoblotting and laminin assays of the WGA-enriched brain, kidney, and heart lysates. The T377T379 matriglycan is consistently shorter. P values for (**k**) are summarized in **sTable5**.

To further compare the two matriglycosylation motifs, we rescued muscle-specific DAG1 KO mice with gene transfer mediated by adeno-associated viruses (AAV) encoding T315A/T317A and T377A/T379A DG. Although the m*yf5-Cre* system cannot eliminate dystroglycan in muscle stem cells, most matriglycan signals detected by immunoblotting are from the AAV-induced DG (**Fig. s7c**). Muscle rescued by AAV encoding T377A/T379A DG showed a mildly higher absolute tetanic force (F_0_) and specific force (sF_0_) than AAV encoding T315A/T317A DG. T315A/T317A DG rescued EDL are also more susceptible to ECC-induced injuries. These data indicate longer matriglycan is superior in terms of rescuing dystroglycanopathy-related muscle pathology (also check **Fig. s7a** and **b**).

We then characterized the two matriglycan motifs in the brain, the kidney, and the heart (**Fig. 5l**). Matriglycan length varies in different tissues. For example, matriglycan in the heart and skeletal muscle is a lot longer than that in the brain. However, T315/T317 matriglycan is consistently longer than T377/T379 matriglycan. In addition, the length of the two matriglycans is individually regulated in different tissues. In skeletal muscle and brain, the T315/T317 matriglycan is far longer than the T377/T379, whereas these two matriglycans are similar in size when synthesized in the kidney, suggesting that there are tissue-specific factors that regulate site specific matriglycan length.

### Skeletal muscle pathophysiology emerges when both matriglycan motifs are mutated

Next, we wanted to know what happens when both matriglycan motifs are mutated. Since direct breeding of heterozygous mice did not generate any 4X mutant mice, we created heterozygous mice whose one allele encodes a floxed WT DAG1 and another allele encodes a DAG1 4X mutant (**Fig. 6a**). Immunoblotting shows that there is no matriglycan present in skeletal muscle when one copy of WT *DAG1* is removed by breeding the heterozygous 4x mutant DG mice with *Pax7-Cre* mice or by injecting AAV-*CMV-Cre* (**Fig. 6b**). This indicates that there are only two matriglycan sites on DG and enables us to pinpoint the neuromuscular functions of matriglycan in the mouse models. The skeletal muscles of 4X mutant mice showed fibers with central nuclei, adipose tissue infiltration, and fibrosis (**Fig. 6c**), which resembles the muscular phenotypes of the *myd* mice (*45*). AAV encoding 4X mutant DG lost its ability to rescue grip strength (**Fig. 6d**). The 4X mutant mice consistently show lower forelimb grip strength and F_0_ than WT mice (**Fig. 6e-h, j-l**). They are also much more susceptible to ECC-induced injuries (**Fig. 6i** and **m**). In addition, NMJs in tibialis anterior (TA), EDL, and soleus (SOL) muscles from *Pax7-Cre* 4X mutant mice are more fragmented than WT mice and mice with a single matriglycan (**Fig. 6n** and **o**). Interestingly, NMJs from mice possessing only one matriglycan show a low degree of irregularity, which indicates the existence of two matriglycans is critical for NMJ integrity. These data once again highlight the function of matriglycan as an ECM receptor and strongly support that there are at least two matriglycan motifs in skeletal muscle *in vivo*.

**Fig. 6.**
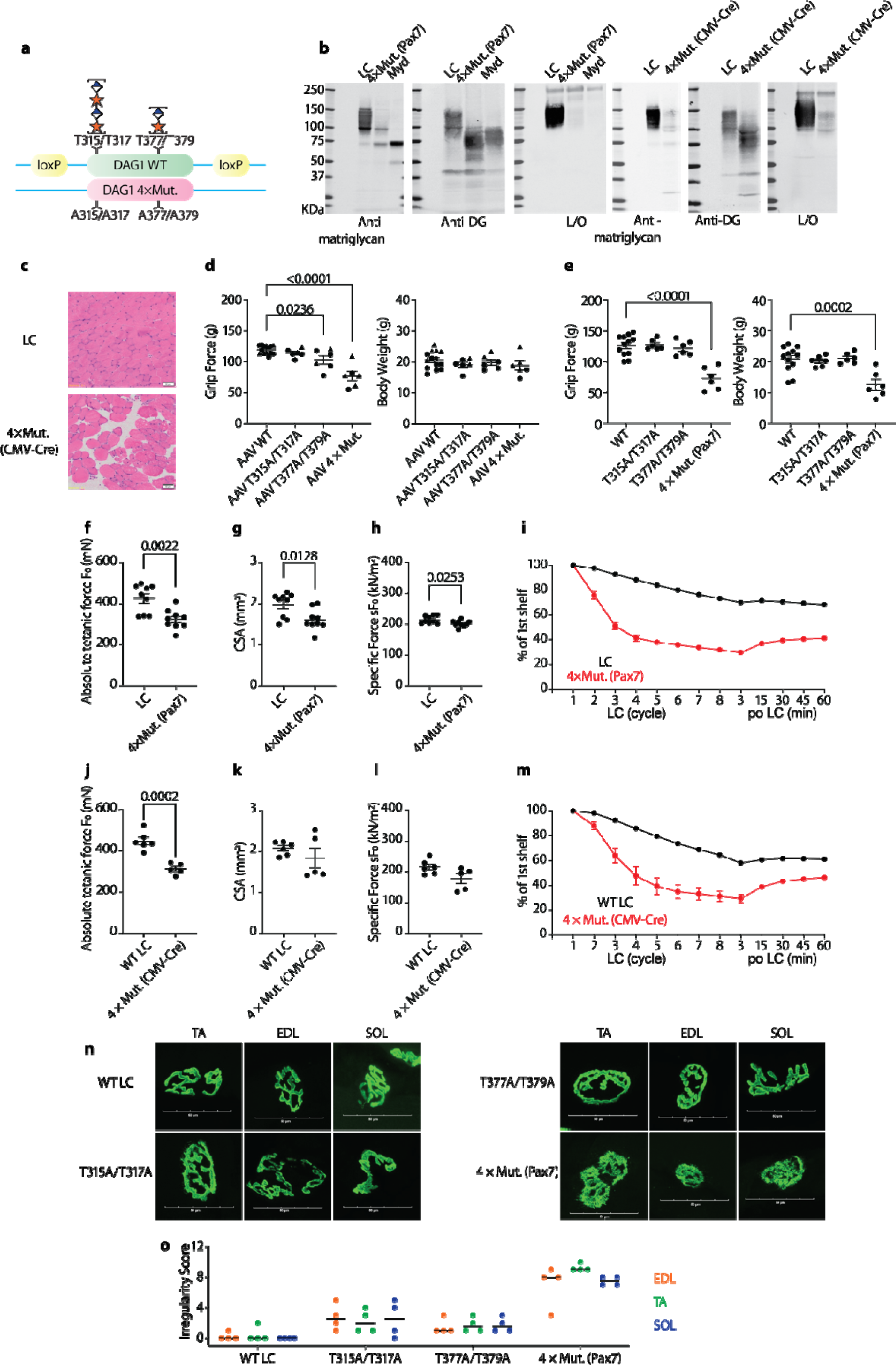
Mutation of both matriglycans in skeletal muscle leads to muscular disorders. (**a**) Diagrams illustrating alleles of heterozygous mice carrying floxed WT DAG1 and DAG1 4X mutant. The Floxed DAG1 can be KO by breeding with pax7 Cre or by injecting AAV Cre (**b**). The KO leads to pathologic muscle as shown by H&E stain (**c**). When both matriglycosylation sites get mutated, forelimb grip force (**d-e**), absolute tetanic (F_0_) and specific (sF_0_) forces (**f-h, j-l**) are all reduced, while muscles are more susceptible to ECC-induced injuries (**i and m**). Mice with T315A/T317A, T377A T379A, and Pax7 Cre DAG1 result in post-synaptic defects. Neuromuscular junctions from TA, EDL, and SOL muscles were obtained from 10-14-week-old mice. (**n)** Representative images of post-synaptic terminals (α-BTX-488) from TA, EDL, and SOL muscles. Scale bars = 50 µm. (**o**) Dot chart of irregularity score for post-synaptic defects (scoring criteria described in Materials and Methods). P values for (**i**) and (**m**) are summarized in **sTable5**.

### ECM proteome is modulated by the length of matriglycan

We initially expected the mice with only the T377/T379 matriglycan to show severe pathologic phenotypes as the matriglycan is short compared to the T315/T317 matriglycan. In the brain, the matriglycosylated T315A/T317A α-DG smear shows less apparent molecular weight than the smear corresponding to mucin-type *O*-glycosylated α-DG (**Fig.5l**). Surprisingly, not only are the transgenic DG T315A/T317A mice generally healthy, but even the AAV encoding DG T315A/T317A can rescue DG KO mice to nearly healthy conditions. We speculated that other ECM proteins might compensate for the effect of the reduced matriglycosylation. We sought to tackle the question with label-free, quantitative shotgun proteomics. We first performed a “quality control” experiment on LARGE1 over-expressed HEK cells to confirm the reproducibility of our proteomic workflow and to make sure we can identify LARGE1 over-expression (**Fig. s8a** and **Table s6**).

Considering one of the major secretors of ECM proteins is the fibroblast cell, and matriglycans are differentially downregulated in human patients suffering different types of dystroglycanopathies, we decided to compare the proteome of cultured primary fibroblast cells from a WWS patient carrying an ISPD mutation and LGMD patients carrying FKRP mutations. The ISPD mutation can severely truncate the matriglycan, so the WWS dystroglycan was selected to resemble the glycan found in the DG T315A/T317A mice. Similarly, the downregulation of matriglycan caused by the FKRP was selected to resemble the DG T377A/T379A mutation. We performed the same workflow on fibroblast cells from three different WWS patients and three different LGMD patients and quantified 3582 (**Fig. s8b** and **c**, and **Fig. 7a**) out of 4235 detected proteins at 1% cut-off false discovery rate (FDR) of both proteins and peptides. Principle component analysis (PCA) accurately separated WWS from LGMD samples (**Fig. 7b**), while unsupervised hierarchical clustering analysis (HCA) robustly segregated their proteomes (**Fig. s8d**). Although certain collagens are mildly upregulated, most detected fibroblast biomarkers including many collagens, integrins, and vimentin were not that differently expressed (**Fig. s8e**), so they should not be used for separating the two groups. On the contrary, WWS cells are characterized by ECM and membrane proteins such as THBS2, COL18A1, FBN2, and HLA-A, while LGMD cells are better characterized by cytosolic and nucleus proteins such as CUL7, RIF1, and KMT2A (**Fig. 7c**).

**Fig. 7.**
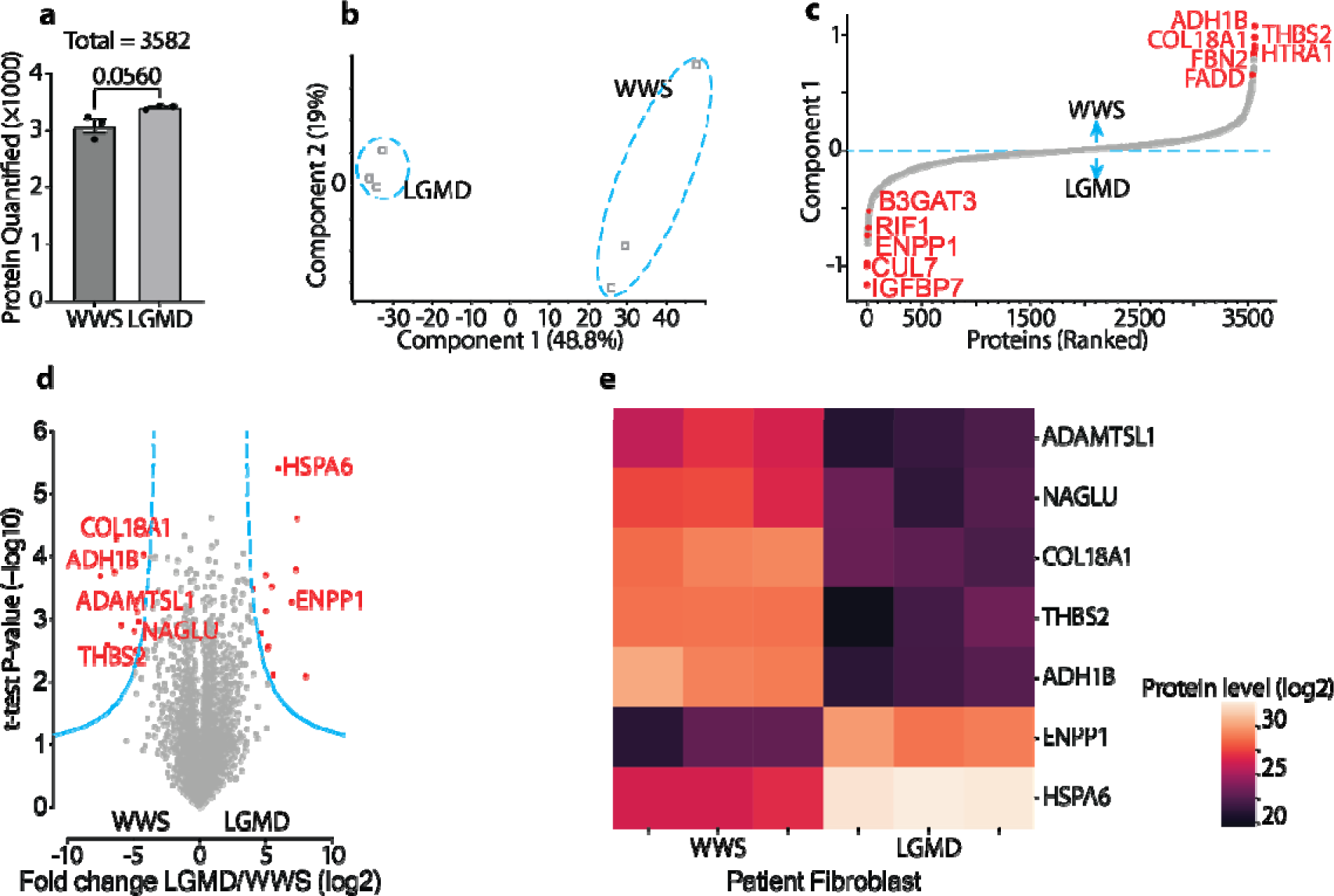
Label free, quantitative proteomics of primary fibroblast indicates ECM modulation by matriglycans. (**a**) Number of unique proteins quantified by MaxLFQ from each group of diseases (n = 3 patients for each disease). (**b**) Principal component analysis (PCA) groups limb-girdle muscular dystrophy (LGMD) and Walker-Warburg Syndrome (WWS) samples. (**c**) Ranking of proteins by expression in WWS cells versus LGMD cells shows WWS proteomes characterized of extracellular matrix proteins. (**d**) Volcano plots comparing LGMD and WWS cells screened out 25 significantly differentially expressed proteins (n red). Two-sided t-test FDR < 0.01. Seven proteins mainly expressed by fibroblast in skeletal muscle are annotated with gene names. (**e**) Heat map of the seven proteins differentially expressed in the patients’ fibroblasts. A dark purple color presents a low expression while bright beige color presents a high expression.

Further analysis screened out 25 differentially expressed protein groups (**Fig. 7d**). While the proteins with higher expression in LGMD cells did not show any apparent features, half of the 12 proteins increased in WWS cells are ECM proteins (THBS2, COL18A1, HTRA1, FBN2, NAGLU, and ADAMTSL1). The 25 altered proteins were further manually scrutinized against the single-cell RNA sequencing database on the Human Protein Atlas website (https://www.proteinatlas.org/) to pinpoint proteins mainly expressed by fibroblast cells in skeletal muscle (**Fig. 7e**). The proteins upregulated in LGMD cells were mostly filtered out and six out of seven remaining targets, which are mostly upregulated in WWS cells, belong to the ECM proteome, and the left one is an enzyme related to ECM degradation (*46*, *47*). Meanwhile, HCA also leads to the clustering of the WWS samples characterized by upregulated ECM region proteins (**Fig. s8f**). Collectively, our data indicate the ECM proteome is directly modulated by the matriglycan and shorter matriglycan tends to upregulate ECM proteins. The modulation can be achieved by directly altering the secretion protein profile of fibroblasts.

## Discussion

### The optimized MS strategy facilitates characterization of “negatively charged” high molecular weight glycoconjugates

The nanoLC-ESI-MS technique has become the most important tool for protein PTM investigations due to its high throughput, ultrahigh sensitivity, and wide compatibility with all modifications. However, glycopeptides carrying long sugars modified by acidic groups, such as phosphate and sulphate, are inherently challenging for existing glycoproteomic methods because both their high molecular weight carbohydrate moieties and the modification groups can hinder ionization. This severely limits the application of MS in characterizing glycosylation processes including glycosaminoglycans, matriglycans, and sometimes sulphated mucin-type *O*-glycans. Here, we provide a nanoLC-ESI-MS-based solution for analyzing matriglycosylated peptides. Instead of inserting additional protease sites to digest big glycopeptides into shorter, more manageable, parts, we selected to maintain the peptide backbone and added a hexa-histidine tag to aid the ionization. The resulting His-tagged peptides get ionized into ions with multiple charges, comfortably fitting the mass range of modern MS instruments. More importantly, their big sizes stabilize fragile groups and keep protein modifications intact when collisional energy is imposed during MS/MS experiments. This is especially beneficial to characterize sulphated and phosphorylated carbohydrates. In addition, even when we used an LC elution program with more concentrated acetonitrile, the sulphate modification still delays matriglycopeptides to very late retention times, which happens to be compensated by adding His-tags to glycopeptides (**Fig. s5c** and **d**). All these experimental considerations and designs facilitated a routine detection of nearly intact matriglycans in a positive ion mode bottom-up shotgun glycoproteomic setup. Our MS strategy might contribute to further research on glycosaminoglycans and sulphated mucin-type *O*-glycans, which are attracting more and more biological research interests.

### Elongation of matriglycans is potentially regulated by two mechanisms

Our MS methods and data provide new insights into matriglycan elongation, although being limited by the DG over-expression model. The biosynthesis of matriglycan might not be the most efficient, considering only a very small portion of matriglycan precursor eventually gets matriglycosylated (**Fig.s6a**). When the matriglycan reaches about 10 disaccharide units during its elongation process, it tends to dissociate from LARGE. The matriglycan might bind to LARGE again for another round of elongation but it can also be capped by sulphate or a hexose group by other enzymes. At least in our cell models, LARGE xylosyltransferase activity tends to be lower than its GlcA transferase activity, it might be outcompeted by capping enzymes. As a result, the matriglycans tend to get capped and their elongation processes are terminated. More interestingly, capping reactions can happen on the adaptor of matriglycan (**Fig. 3g**-**j**), which might stop matriglycosylation at the very beginning. This partially explains why the amount of matriglycosylated peptides is so small. The sulphation process mediated by HNK-1 explained why matriglycans in brains are a lot shorter, but it seems that a similar mechanism exists in other tissues, such as the kidney. As such, our model provides a possible mechanism for the postnatal regulation of matriglycan elongation in different tissues.

LARGE1 activity may be increased by molecular recognition of DG-Nt and the phosphate on coreM3, thereby explaining the existence of very long matriglycans (*11*, *48*, *49*). If the observed 80 Da modification corresponds to a phosphate on an *O*-mannose (see **Fig.2**), it is likely the extra mannose-phosphate can further boost LARGE activity. We speculate the T317/T319 matriglycan represents the boosted LARGE1 activity whereas the T379/T381 matriglycan is the product of LARGE1 without other chaperones. T377A/T379A α-DG exhibited considerably higher electrophoretic migration than the WT. This indicates both matriglycosylation sites can be simultaneously occupied, although the T-to-A mutation also affects mucin-type *O*-glycosylation. We have not elucidated the reason why two matriglycans are required, but our limited evidence indicates this is important for the integrity of NMJ. In addition, we noticed both matriglycan motifs from α-DG can be completely deglycosylated by the glycosidases that remove mucin-type *O*-glycans. This indicates α-DG exists in more than one glycoform carrying only the long, the short, or both matriglycans. Two different matriglycan motifs might provide another straightforward mechanism to combinatorially regulate the length of matriglycan tissue-specifically, during development and muscle regeneration.

### Matriglycans modulates ECM proteome

We found that mice with only the short matriglycan suffer mild embryonic lethality. This reveals the critical roles of the elongation of matriglycan during mammalian embryonic development. However, our data very strongly indicate that if the mice survived the developmental stage, they grew into healthy adults. More surprisingly, it is evident that the T377/T379 motif carrying the shorter matriglycan is more capable of restoring grip strength. This means α-DG with only shorter matriglycan maintains cell-matrix interactions well at least in mice after development. The phenotypical comparison, especial for the fiber size (**Fig.5c**) and NMJ morphology (**Fig.6o**), between the DG T315A/T317A and T377A/T379A mice needs further validation because the current data are limited by a small number of mice available for experiments.

We tried to address these physiological observations by performing label-free proteomic quantification on fibroblast cells that are known to secrete ECM proteins directly related to muscular functions. Comparative proteomics previously revealed muscular dystrophy caused by mutations in dystrophin leads to an upregulation of ECM proteins such as collagen and fibronectin (*43*). We identified a few upregulated proteins in WWS cells with truncated matriglycans including THBS2 which was reported to stabilize muscle membranes (*50*), and ADAMTSL1 was shown to help muscle regeneration (*51*). The tendency of ECM protein upregulation in matriglycan truncated cells is also supported by HCA. We propose the upregulation of these proteins might compensate for the “side effect” of reduced matriglycan, which explains why mice with shorter matriglycans can be postnatally healthy. Notably, this ECM “rescue effort” is likely ineffective during embryonic development, which highlights the essential role of matriglycan in mammalian development. We plan to apply our proteomic approach in analyzing the brain and skeletal muscle, which should be more physiologically relevant. Meanwhile, such experiments will likely provide more biochemical insights on functions of the two matriglycans.

## MATERIALS AND METHODS

### Viral vectors

#### DAG1 WT, T315A/T317A, T377A/T379A, T315A/T317A/T377A/T379A (4X mut.) viral vectors

Sequences encoding mouse WT and site-mutated DAG1 were synthesized (GenScript, Piscataway, NJ) and cloned into adeno-associated virus 2/9 (AAV2/9) backbones under the transcriptional control of the ubiquitous CMV promoter (University of Iowa Viral Vector Core Facility). The advantages of the AAV2/9 vector were previously described (*1*). Briefly, the vectors show improved muscle transduction efficiency and altered tropism. The plasmid was also used for making adenoviral vectors for infecting HEK293F cells.

#### DG390 TevHis6 (DG390) adenoviral vector

The adenoviral vector was previously reported (*2*). Briefly, FseI-x-390 AAs-TEV-6xHis-NotI was obtained using pcDNA3rbtDG as a PCR template and ligated into the pcDNA3DG340TEVHis digested with FseI and NotI to construct DG390TEVHis, which includes 1–390 AAs of rabbit DG, TEV site, and 6x Histidine. The construct was inserted in the pacAd shuttle plasmid from the University of Iowa Viral Vector Core Facility to generate the vector. E1-deficient recombinant adenoviruses were generated by the University of Iowa Viral Vector Core using the RAPAd system. The vectors were used to infect HAP1 cells for the investigation of SLC35A1 and POMGnT1 functions.

### Genetically modified cell lines

#### HEK293F bioreactor constitutively expressing DG390TevHis6

HEK293F cells were stably transfected (XtremeGENETM 9 DNA transfection reagent) according to the manufacturer’s protocol with pcDNA3 encoding DG390TevHis6. The cells were cultured with a serum-free culture medium and were constantly selected by neomycin.

#### DAG1 KO HEK293 cells

The cells were purchased from Abcam (ab266263). Knockout achieved by using CRISPR/Cas9, homozygous: 1 bp insertion in exon 2.

#### SLC35A1 knockout (KO) HAP1 cells

HAP1 cells bearing a 2 bp deletion of exon 2 of SLC35A1, generated using the CRISPR/Cas9 system, were purchased from Horizon Discovery (HZGHC010158c010). The sequence of guide RNA is TTCTGTGATACACAACGGCTG. SLC35A1 KO was confirmed by Horizon Discovery via PCR amplification and Sanger sequencing. PCR primers used for DNA sequencing are SLC35A1 Forward CACGTATTTCCAGACAATGTCACT, and SLC35A1 Reverse ACAAATACCCTCACCAGGTAGAAAA. The KO was further functionally confirmed by mass spectrometry-based glycomics. To generate DG390 proteins for MS analyses, HAP1 cells were grown in IMDM with 10% FBS and 1% penicillin/streptomycin on p150 plates. When plates were 80% confluent, cells were washed twice with DPBS, media changed to serum-free IMDM with 1% penicillin/streptomycin (Invitrogen), and cells infected at high MOI (250–1000) of adenovirus expressing DG390TEVHis. The culture media were harvested after 72 hours and stored at 4 □C.

#### POMGnT1 knockout (KO) HAP1 cells

HAP1 cells bearing an 8 bp deletion of exon 2 of POMGnT1, generated using the CRISPR/Cas9 system, were purchased from Horizon Discovery (HZGHC000747c012). The sequence of guide RNA is AGCGGAGCTGGTACCTTACC. POMGnT1 KO was confirmed by Horizon Discovery via PCR amplification and Sanger sequencing. PCR primers used for DNA sequencing are SLC35A1 Forward TATAGGGACATGACACATAGAGGTC, and SLC35A1 Reverse ATCAGACTTTGGGAGGAGCATCTTA.

### Proteins expression and purification

#### DG390TevHis6 (DG390)

DG390 secreted by HEK293F, WT HAP1, and SLC35A1 KO HAP1 cells was purified by HisPur™ Ni-NTA Resin (Thermo Fisher Scientific, US). The nickel resin was eluted by 300 mM imidazole, which was removed by extensive dialysis against a 50 mM ammonium bicarbonate buffer. The sample is lyophilized before being stored at −20°C.

#### Dystroglycan protein DAG1_28–749_^R311A/R312A^ (DG749)

DG749 purification was previously described (*3*). Briefly, HEK293F cells were stably transfected (XtremeGENE^TM^ 9 DNA transfection reagent; Sigma-Aldrich) according to the manufacturer’s protocol with pcDNA3.1 + encoding human dystroglycan residues 1–749 with a mutation in the furin cleavage site (R311A/R312A). The secreted dystroglycan protein DAG1_28–749_^R311A/R312A^ was purified using His60 Ni IMAC resin (Takara) and subjected to anion exchange using DEAE resin. The sample is lyophilized before being stored at −20°C.

#### DG-Fc

DG-Fc protein was reported (*4*, *5*), and it was purified by protein G beads from cell culture media.

#### LARGE1dTM

Pure recombinant LARGE proteins were obtained as previously described (*1*). Briefly, LARGE1dTM secreted by stably transfected HEK293F cells were purified from the media using TALON resin. The eluate was dialyzed against 50 mM ammonium bicarbonate before lyophilization and stored at −20°C.

### Genetically modified mice

#### myf5Cre/+; Dag1fl/fl (myf5-Cre/DAG1 KO)

Mice homozygous for floxed DG, in which loxP sites flank exon 2 of Dag1, were crossed with transgenic mice expressing a cre-mediated recombination system (007845; The Jackson Laboratory). This recombination system was driven by the Myf5 promoter. Male offspring heterozygous for the floxed Dag1 and Cre-positive were crossed with female mice heterozygous for the floxed Dag1 allele. When bred to mice with a cre recombinase gene under the control of the Myf5 promoter of interest, exon 2 of the Dag1 gene is deleted in skeletal muscle.

#### Matriglycosylation motif mutant transgenic mice

The transgenic mice were generated by using a CRISPR-Cas9 system according to previous publications (*6*).

C57BL/6J mice were purchased from The Jackson Laboratory (000664; Bar Harbor, ME). Male mice older than 8 weeks were used to breed with 3-5 weeks old super-ovulated females to produce zygotes for electroporation. Female ICR (Envigo; Hsc:ICR(CD-1)) mice were used as recipients for embryo transfer. All animals were maintained in a climate-controlled environment at 25°C and a 12/12 light/dark cycle. Animal care and procedures conformed to the standards of the Institutional Animal Care and Use Committee of the Office of Animal Resources at the University of Iowa.

Chemically modified CRISPR-Cas9 crRNAs and CRISPR-Cas9 tracrRNAs were purchased from IDT (Alt-R® CRISPR-Cas9 crRNA; Alt-R® CRISPR-Cas9 tracrRNA (Cat# 1072532)). The crRNAs and tracrRNA were suspended in T10E0.1 and combined to 1 µg/µl (∼29.5 µM) final concentration in a 1:2 (weight-to-weight) ratio. The RNAs were heated at 98°C for 2 minutes and allowed to cool slowly to 20°C in a thermal cycler. The annealed cr:tracrRNAs were aliquoted to single-use tubes and stored at −80°C.

Cas9 nuclease was also purchased from IDT (Alt-R® S.p. HiFi Cas9 Nuclease). Individual Cr:tracr:Cas9 ribonucleoprotein complexes were made by combining Cas9 protein and cr:tracrRNA in Opti-MEM (Gibco; 31985062; final concentrations: 1 µg/µl (∼6.2 uM) Cas9 protein and 0.6 µg/µl (∼17.7 µM) cr:tracrRNA). The Cas9 protein and annealed RNAs were incubated at 37°C for 10 minutes. The RNP complex was mixed with repair oligo (IDT ultramer) resulting in final concentrations of 100 ng/ul (∼0.62 µM) Cas9 protein and 60 ng/ul (∼1.8 µM) cr:tracrRNA and 0.5 µg/µl donor oligo.

Pronuclear-stage embryos were collected using described methods (*6*). Embryos were collected in KSOM media (Millipore; MR101D) and washed 3 times to remove cumulous cells. The embryos were treated for 10 seconds in acid Tyrode’s solution (Sigma; T1788) to weaken the zona. The embryos were then washed 5 times with Opti-MEM and 25-30 were aligned between the electrodes of a slide glass plate electrode with a 1 mm gap and platinum leads. The acid-treated embryos were overlayed with the electroporation mix and electroporated for 7 cycles at 30V (3 msec on:100 msec off). Electroporation was performed using an ECM830 square wave electroporator (BTX). Embryos were immediately implanted into pseudo-pregnant ICR females.

PCR-genotyping and Sanger sequencing was done to identify founders with the intended DAG1 mutations.

Dag1_T377A_T379A_CrA ATTATTCAGACCCCAACTCT

Dag1_T377A_T379A_CrB CAGAGTTGGGGTCTGAATAA

Dag1_T315A_T317A_CrX CCAATGGCAGTAACAGGTGT

Dag1_T315A_T317A_CrY GGCAGTAACAGGTGTAGGTG

Dag1_T377A_T379A_601 TAGTGCCTACGCCTACATCT

Dag1_T377A_T379A_602 GGACACTCGTGGCTTCTTT

Dag1_T315A_T317A_901 GCTGCTCCTTGAACCAGAATA

Dag1_T315A_T317A_902 TGCAATGGCTGGAGATGTAG

Dag1_T377A_T379A_Donor

AGGGATCCTGTTCCAGGGAAGCCCACGGTCACCATTCGGACGCGAGGTGCCATTATTCAGGCTCCTGCACTCGGCCCTATCCAGCCTACTCGGGTGTCAGAAGCTGGTACCACGGTTCCTGGCCAGATTCGC

Dag1_T315A_T317A_Donor

ACATTGCCAATAAGAAGCCCACTCTCCCCAAACGACTCCGGAGGCAGATCCACGCAGCACCTGCACCTGTTACTGCCATTGGACCCCCAACCACGGCCATTCAGGAGCCACCATCGCGGA

### Mass spectrometry-based glycan and glycopeptide analysis

#### Glycosidases Treatment

DG390 was deglycosylated by a mixture of β (1–3,4)-Galactosidase, β-N-Acetylhexosaminidase, O-glycanase, α (1–2,3,4,6) fucosidase and Sialidase A (all enzymes were purchased from Glyko/ProZyme, Inc.) in 100 mM ammonium acetated buffer (pH = 5.5) containing 1X MS-SAFE Protease and Phosphatase Inhibitor. Deglycosylated DG390 was lyophilized before further experiments.

#### Reductive Elimination

*O*-glycans were reductively eliminated from DG390 proteins and purified on a 50 W Dowex column according to published methods (*7*, *8*). Briefly, lyophilized DG390 was resuspended in 55 mg/mL potassium borohydride in 1 mL of a 0.1 M potassium hydroxide solution. The mixture was incubated overnight at 45 □C and quenched by adding 5-6 drops of acetic acid. The sample was loaded on the Dowex column and eluted with 5% acetic acid. The collected solution was concentrated and lyophilized before excessive borates were removed with 10% methanolic acetic acid under a stream of nitrogen.

#### Permethylation and deutero-permethylation

Three to five pellets per sample of sodium hydroxide were crushed in 3 mL dry dimethyl sulfoxide. The resulting slurry (0.75 mL) as well as iodomethane or iodomethane-d3 (500 µL) were added to the sample. The mixture was agitated for 60 minutes before quenched by adding 2 mL ultrapure water with shaking. Derivatized glycans were extracted with chloroform (2 mL) and washed with ultrapure water two more times before it was dried down under a stream of nitrogen.

Before MS analysis, the derivatized glycans were dissolved in 50% methanol and loaded onto a preconditioned C18 Sep-Pak cartridge (Waters). The cartridge was washed with 5 mL ultrapure water and 3 mL of 15% acetonitrile before eluting by 3 mL of 50% acetonitrile. The purified glycans were lyophilized and dissolved in 10 µL of methanol. One µL of the solution was mixed with 1 µL of 20 mg/mL 3,4-diaminobenzophenone and analyzed by a Bruker ultrafleXtreme MALDI-TOF/TOF.

#### Glycoproteomics

Twenty µg of lyophilized deglycosylated DG390 was digested by 1 µg MS grade trypsin (Promega) in a 50 mM ammonium bicarbonate buffer (pH=8.4) for 8-10 hours at 37°C. The buffer was lyophilized before the tryptic peptides were dissolved in 10-20 µL of 0.1% formic acid. 1 µL of the solution was injected into an UltiMate^TM^ NCS-3500 nanoLC (Thermo Fisher Scientific, US) coupled to an orbitrap Fusion Lumos mass spectrometer (Thermo Fisher Scientific, US). A two-solution (solution A: 0.1% formic acid; solution B: 80% acetonitrile, 0.1% formic acid) gradient was used to elute an Acclaim™ PepMap™ 100 C18 LC Columns (25 cm, 2 µm, 100Å, 75 µm, Thermo Fisher Scientific, US): 0-1% B in 8 minutes, 1-5% B in 1 minute, 5-60% B in 90 minutes and kept at 60% for another 6 minutes. The solution B was then elevated to 95% to wash the column for 5 minutes before coming back to 0%. The LC flow rate was 300 nL/minute, and a guard column was used to further desalt the samples. The column temperature was 45 □C.

MS methods were slightly modified for different samples according to instrumental status. Detailed methods are included in each raw data file. General methods used for routinely analysis are below. For MS, the mass spectrometer was operated under the positive mode. The spray voltage is 1900 V. The ion transfer line temperature is 275°C. MS resolution was set to 120K and the mass range was 400-2000 Da. The maximum injection time was 72 ms. The AGC target was 400000. S-lens RF level was 60. MS was active from 8 to 120 minutes.

For MS/MS, higher-energy collision dissociation (HCD) triggered collision-induced dissociation (CID), CID triggered electron transfer dissociation (ETD), EThcD triggered HCD, and HCD triggered MS3 modes were used. High resolution ETD dissociation was also tested but the data are of lower quality than EThcD. The trigger ions include 204.087, 366.140, 274.092, 163.094, 940.213, 632.134, and 418.110. The initial MS/MS were recorded by the orbitrap (resolution set to 30K), while triggered MS/MS were recorded by the ion trap. All MS/MS started recording from 100 Da. Isolation for Orbitrap was all set to quadrupole. Two microscans were accumulated to generate initial MS/MS spectra, while four were accumulated to generate triggered MS/MS spectra. For HCD triggered CID, HCD maximum injection time was 150 ms and the AGC target was 150000. A stepped HCD collision set to 26%, 28%, and 30% was used to dissociate glycopeptides. Triggered CID has an AGC target of 5000 and maximum injection of 75 ms. Activation time was 10 ms and Q value was 0.25. Stepped CID collisional energy was set to 33%, 35% and 48%. For CID triggered ETD, the maximum injection time for CID analysis was 150 ms, and the AGC target was 150000. A stepped CID collision was set to 33%, 35% and 37%. For the ETD, EThcD was active to facilitate fragmentation. AGC target was set to 10000 and maximum injection was 75 ms. For EThcD triggered HCD, EThcD MS/MS resolution was 200 ms and the AGC target was 200000. A hcD collision energy at 15% was used to facilitate ETD fragmentation. For the triggered HCD, a stepped HCD collision was set to 26%, 28% and 33%.

Extracted ion chromatograms were generated by Skyline software. Protein deconvolution was done by Byonic Intact Mass software. All shown MS signals were found by a manual inspection of raw MS data.

### Mass spectrometry-based proteomics

#### Protein in-gel digestion

Proteins were separated by 10-20% tris-tricine gel and in-gel digested according to a previous published protocol with modifications (*9*).

Briefly, gel pieces containing proteins were cut, transferred, and mashed into small pieces. The pieces were incubated in 500 µL of acetonitrile for 10 minutes until they shrunk. The shrunken pieces were incubated in 50 mM dithiothreitol (Roche) in 50 mM ammonium bicarbonate at 56°C for 30 minutes, washed with 500 µL acetonitrile, incubated in 100 mM iodoacetamide in dark at room temperature for 20 minutes, and washed with 500 µL acetonitrile.

The gel pieces were then incubated with 60 µL of a 50 mM ammonium bicarbonate solution of trypsin (MS grade, Promega) at 4°C for 90 minutes and incubated at 37°C overnight. The protease to protein ratio is roughly 1:50. The liquid in the tubes were collected before gel pieces were eluted by 100 µL of 66.6% acetonitrile in 0.1% formic acid. The solutions were combined and concentrated with a SpeedVac (Thermo Fisher Scientific, US).

#### Filter-aided sample preparation for proteomics of human fibroblasts

Human fibroblasts were cultured in Dulbecco’s Modified Eagles Medium (DMEM, Thermo Fisher Scientific, US) supplemented with 20% fetal bovine serum (FBS, Thermo Fisher Scientific, US), 2% L-glutamine (Thermo Fisher Scientific, US), and 1% penicillin/streptomycin (Thermo Fisher Scientific, US). Cells were maintained in a humid 5% CO2 atmosphere at 37°C and were passaged using 1x trypsin-EDTA approximately every 2-3 days (when cells reached 60-80% confluency). Fibroblasts were harvested by gently perturbing plates rinsed with 1x PBS. Fibroblast pellets were homogenized by adding a solution containing 4% SDS, 100 mM Tris (pH=8), 100 mM Dithiothreitol (DTT) at 95°C for 3 minutes. The follow-up filter-based protein digestion method was previous described (*10*). With the aid of the 30 KDa cut-off filter (Microcon), the solution was replaced by 8 M urea and disulfide bonds were blocked by iodoacetamide (IAA). The urea and IAA solutions were eventually exchanged into a 50 mM ammonium bicarbonate buffer, and the proteins were tryptic digested (Promega) at 37°C overnight. The digested samples were de-salted by classical C18 Sep-Pak cartridges (Waters).

#### Proteomics

The concentrated solutions were diluted to 2-3 mg/mL by 0.1% formic acid and 1 µL of the solution was injected into the same nanoLC-MS instrument. A two-solution (solution A: 0.1% formic acid; solution B: 80% acetonitrile, 0.1% formic acid) nanoLC gradient was used to elute an Acclaim™ PepMap™ 100 C18 LC Columns (25 cm, 2 µm, 100Å, 75 µm, Thermo Fisher Scientific, US): 0-1% B in 8 minutes, 1-5% B in 1 minute, 5-40% B in 200 minutes, 40-50% B in 5 minutes, 40-50% B in 5 minutes. The solution B was then elevated to 95% to wash the column for 5 minutes before coming back to 0%. The LC flow rate was 300 nL/minute. Column temperature was kept at 40°C. A guard column was used to further desalt the samples. The mass spectrometer was operated under the positive mode. The spray voltage is 1900 V. The ion transfer line temperature was 275°C. MS resolution was set to 120K and the mass range was 400-2000 Da. The maximum injection time was 72 ms. The AGC target was 4e5. S-lens RF level was 60. HCD MS/MS resolution was 30K. The maximum injection time was 75 ms. The AGC target was 1.2e5. The HCD collision energy was 28%. CID MS/MS was performed by linear ion trap, collisional energy was set to 35%, activation time set to 10 ms, and activation Q value was 0.25. MS/MS isolation offset was 1.6. MS was active from 8 to 240 minutes.

#### Data analysis

MS Raw data were processed by MaxQuant. The peptide and protein identification were determined by the MaxQuant built-in Andromeda engine (*11*) that searches raw data against a Uniprot human proteome. False discovery rate threshold was set to 1%. The MaxLFQ algorithm (*12*) was used for label-free proteome quantification. The minimum ratio count for quantification was set to 1, while the final list of protein quantifications was filtered for proteins with a minimum of two peptides for each protein. The ‘Match Between Runs’ feature was enabled. All statistics and bioinformatics were done by using Perseus (*13*, *14*). For the two-sample test, we performed student t-test with a P value threshold of 5%. Missing values were replaced based on a normal distribution (width, 0.3; downshift, 1.8). Pearson correlation for **fig. s7a-c** was done by using Python.

### Western blot and laminin overlay

Homogenization A TBS buffer containing 1% Triton-X, 10mM EDTA and proteinase inhibitors (leupeptin, pepstatin A, aprotinin and PMSF) was used to lyse mice tissues (brain, heart, kidney, and skeletal muscle) and cells. Skeletal muscle was minced with a scalpel and homogenized with a Kinematica PT2500 E Polytron. Heart and brain tissues were processed by a “beads beat” method using a Next Advance Bullet Blender. The tissue lysates were centrifuged at 20K xg, 4□ for 30 minutes, and the supernatants were collected.

Wheat-germ agglutinin (WGA) affinity chromatography. Tissue and cell lysates supernatants were added to WGA-agarose resin. The mixtures were rotated overnight at 4L. The resin was then washed three times with 0.1% Triton X-100. The slurry was combined with SDS-loading dye and the sample heated for 10 minutes at 99L.

#### SDS-PAGE

Proteins were resolved on homemade 3-15% polyacrylamide gradient gels in SDS-glycine buffer for either 6 hours at 200 V or 16 hours at 60 V at room temperature. The proteins were transferred to Immobilon-FL (Millipore IPFL00010).

#### Western blotting

Membranes were blocked in either 2% skim milk in low-salt TBS-T or fish gelatin and incubated with antibodies against matriglycan (IIH6), dystroglycan (AF6868 or 199768), or laminin overlay. Antibodies against matriglycan (IIH6) and dystroglycan (AF6868) were made in house. Mouse laminin111 and its antibody used for laminin overlay was purchased from Thermo Fisher Scientific (23017015) and Sigma-Aldrich (L9393).

### Rescuing myf5-Cre/DAG1 KO mice with AAV2/9CMV mDag1

Myf5 cre + flox DAG1 mice at 2-7 days of age were injected with one dose of 10 uL of AAV vectors (1.8 – 5.1 × 10e11 vg) via the retro-orbital sinus. Retro-orbital injections are administered in pups with a 31-gauge, 0.3125-inch needle attached to a 0.3 ml insulin syringe (BD Ultra-Fine II, Becton, Dickinson and Co., Franklin Lakes, NJ). To carry out the injections, we use a dissecting microscope with a fiber optic point source light source. Sunflower seeds were added to the cage after injection to reduce cannibalism.

AAV2/9CMV mDag1 n=16

AAV2/9CMV mDag1 T315A,T317A n=6

AAV2/9CMV mDag1 T377A, T379A n=6

AAV2/9CMV mDag1 T315A,T317A T377A, T379A n=6

### Forelimb grip strength test

The methods were previously described (*1*). Mice were first weighed on a Scout scale (OHAUS CORPORATION in Parsippany, NJ). Mice were then brought to rigid wire grid about 4in. x 4in. in size. This rigid wire grid is attached to a Chatillon E-DFE-002 Force Gauge (Columbus Instruments, Columbus, OH) at a 45-degree angle. The mouse is then allowed to grasp the rigid wire grid with just two forelimbs; trials in which the mouse gripped with one limb, or more than 2 limbs were rejected. The mouse was then gently and steadily pulled back until its grip was broken from the grid; the attached force gauge registers the highest force exerted by the mouse on the pull; this value is recorded. The grip of the mouse and the gentle pull are then immediately repeated 2 more times for a total of 3 times per session. After a session, the mouse was allowed to rest for 1 minute, and 4 more sessions were conducted, for a total of 15 pulls, 5 sessions. The three highest force (in grams) exerted by the mouse were averaged to find the final grip strength of the mouse. Ten–fourteen-week-old mice were (n=6 for each group) used. Statistics were calculated using GraphPad Prism software and two-sided Student’s unpaired t-test or one-way ANOVA were used. Differences were considered significant at a p-value less than 0.05.

Statistics were calculated using GraphPad Prism software and one-way ANOVA was used. Differences were considered significant at a p-value less than 0.05.

### Measurement of muscle function *in vitro*

To compare the contractile properties of muscles, extensor digitorum longus (EDL) muscles were surgically removed as described previously (*15–17*). The muscle was immediately placed in a bath containing a buffered physiological salt solution (composition in mM: NaCl, 137; KCl, 5; CaCl_2_, 2; MgSO_4_, 1; NaH_2_PO_4_, 1; NaHCO_3_, 24; glucose, 11). The bath was maintained at 25°C, and the solution was bubbled with 95% O_2_ and 5% CO_2_ to stabilize pH at 7.4. The proximal tendon was clamped to a post and the distal tendon tied to a dual mode servomotor (Model 305C; Aurora Scientific, Aurora, ON, Canada). Optimal current and whole muscle length (L_0_) were determined by monitoring isometric twitch force. Optimal frequency and maximal isometric tetanic force (F_0_) were also determined. The muscle was then subjected to an eccentric contraction (ECC) protocol consisting of eight eccentric contractions (ECCs) at 3-minute intervals. A fiber length L_f_-to-L_0_ ratio of 0.45 was used to calculate L_f_. Each ECC consisted of an initial 100 millisecond isometric contraction at optimal frequency immediately followed by a stretch of Lo to 30% of L_f_ beyond Lo at a velocity of 1 L_f_/s at optimal frequency. The muscle was then passively returned to L_o_ at the same velocity. At 3, 15, 30, 45, and 60 minutes after the ECC protocol, isometric tetanic force was measured. After the analysis of the contractile properties, the muscle was weighed. The CSA of muscle was determined by dividing the muscle mass by the product of L_f_ and the density of mammalian skeletal muscle (1.06 g/cm3). The specific force was determined by dividing F_o_ by the CSA (kN/mm2). For AAV rescued mice, n = 5 to 8 muscles used for each analysis. For transgenic mice, n=5 to 11 muscles used for each analysis. Each data point represents an individual EDL. Statistics were calculated using GraphPad Prism software and two-sided Student’s unpaired t-test or one-way ANOVA were used. Differences were considered significant at a p-value less than 0.05.

### NMJ imaging and quantification

Upon harvest, Extensor digitorum longus (EDL), Tibias anterior (TA), and Soleus (SOL) were washed in Phosphate Buffered Saline (PBS) 3 times for 5 minutes each time. Each muscle group was fixed in 4% paraformaldehyde in PBS for 20 minutes then followed by 3 washes with PBS. Fixed muscle was split apart into muscle fiber bundles then incubated in 3% Triton X-100 in PBS for 3 hours at room temperature. Muscle bundles were washed in PBS for 5 minutes, then incubated in Background Buster (Innovex; NB306) for 4 hours at room temperature. The muscle bundles are washed with PBS for 5 minutes then incubated with Alexa Fluor 488-conjugated α-bungarotoxin (α-BTX) (Invitrogen; B13422) diluted in 5% background buster at 1:500 ratio for 2 hours. Muscle bundles were then washed for 4 hours in PBS exchanging the buffer every 30 minutes at room temperature. Images were acquired using an Olympus FLUOVIEW FV3000 confocal laser scanning microscope. Complete en face NMJs were identified and acquired with Z-stacks using 100x objectives. Maximum intensity Z-stacks were reconstructed with the FV31S (Olympus) software. Observer analyzes a-BTX-488-labeled acetylcholine receptor (AChR) cluster formations to determine irregularities, fragmentation, synaptic size, and dispersion. Irregularities such as AChR plaques, AChR perforated plaques, ring-shaped or c-shaped clusters, and extensive fragmentation were scored. C-shaped, or ring-shaped NMJs receive a score of 1, fragmented NMJs receive a score of 2, perforated NMJs receive a score of 3, and plaque NMJs receive a score of 4. If more than one irregularity is identified in an NMJ, then the score is the sum of the individual scores (e.g. ring shaped (1) + perforated (3) + plaque (4) = 7). Fragmentation was determined by the number of identifiable individual AChR clusters within the footprint of the a-BTX-488-labeled AChR cluster. Fiji ImageJ software was used for semi-automatic analysis of AChR clusters. Synaptic size refers to the total area of the AChR footprint identified by a-BTX-488-labeled AChR. AChR cluster dispersion was determined by the following equation: total stained area/total area*100. GraphPad Prism was used to quantitate irregular %, fragments, synaptic size, and AChR cluster dispersion.

### H&E and immunofluorescence

The H&E and immunofluorescence cryosections were cut at 10 microns on a Leica CM3050S Research Cryostat then put on Superfrost Plus Microscope Slides (Fisher Scientific, Pittsburgh, PA), and airdried at room temperature for 20 minutes before slides were stained. For H&E, slides were fixed in 10% Neutral Buffered Formalin for 5 minutes, washed in dH_2_O 3×5 minutes, 1 minute in Hematoxylin (Leica SelecTech 560, Richmond, IL), washed in dH_2_O x3 on/off, 0.5 % acid alcohol for a quick dip, washed in dH_2_Ox 3 on/off, Scott’s Tap water for 1 minute, washed in dH_2_O, 80% ethyl alcohol for 1 minute, Eosin (Leica SelecTech 515LT, Richmond, IL) for 20 seconds, dehydrated through a series of 95-100% ethyl alcohol, xylene 3 x 1 minute, mounting medium (Permount ThermoFisher Scientific, UK) and cover slipped (Leica Surgipath coverglass, LeicaBiosystems.com). For Immunofluorescence, slides were washed in 1xPBS for 5 minutes, placed in 1:1 methanol/acetone (precooled to −20°C) for 10 minutes in a flammable storage freezer set at −20°C, washed in 1xPBS, 3×10 minutes, blocked in Background Buster (Innovex Biosciences, Richmond, CA) overnight then washed in 1xPBS, 2×2 minutes before incubation in primary antibodies. The first primary antibody that was used was a mouse monoclonal antibody IIH6 to matriglycan on alpha-dystroglycan (αDG; 1:25; Campbell Laboratory, University of Iowa), was incubated overnight at 4□°C in a humid chamber, washed in 1xPBS for 3×10 minutes followed by secondary antibody Alexa Fluor-conjugated 488 goat anti-mouse IgM (1:500; Invitrogen Thermo Fisher Scientific, Waltham, MA) for 40 minutes in the dark, washed in 1xPBS 3×10 minutes then mounting medium, (ProLong Gold Antifade Mountant, Thermo Fisher Scientific, Waltham, MA) was added and slides were cover slipped (Surgipath coverglass, LeicaBiosystems.com) and dried 24 hours before imaging. Another primary antibody used was affinity purified βDG rabbit polyclonal AP83 (1:50; Campbell Laboratory, University of Iowa) followed by Alexa-Fluor-conjugated 555 goat anti-rabbit IgG (1:500; Invitrogen Thermo Fisher Scientific, Waltham, MA). The steps for this primary and secondary antibody were the same as stated above for the other primary and secondary antibody used. Digital images of stained sections were captured with a VS120-S5-FL Olympus slide scanner upright microscope (Olympus Cooperation, Tokyo, Japan). For measuring fiber radius, rabbit polyclonal anti-collagen VI primary antibody (1:100, Fitzgerald Industries International) and goat anti-Rabbit IgG (H+L) secondary antibody (1:500, Alexa Fluor^TM^ Plus 647, Thermo Fisher Scientific) were used to visualize collagen VI. Monoclonal primary antibody (1:200, A7L6, Thermo Fisher Scientific) and the goat anti-rat IgG (H+L) secondary antibody (1:250, Alexa Fluor^TM^ 488, Thermo Fisher Scientific) were used to visualize perlecan. The slides were dried for 10 minutes, washed in PBS for 5 minutes, and blocked by 3%BSA in PBS for 30 minutes at room temperature. The primary antibodies were added in PBS containing 1% BSA, and the slides were incubated in the mixture at 4°C overnight. The slides were then washed 5 minutes in PBS three times, incubated with PBS containing 1% BSA and the secondary antibodies for 30 minutes, and washed with PBS three times at room temperature. The slides were stored in dark, and images were collected by using SlideScanner 24 hours later. Feret’s diameter of muscle fibers was measured with VS-DESKTOP software (Olympus).

## Acknowledgments

We thank the University of Iowa Viral Vector Core for generating the adeno-associated viral vectors (http://www.medicine.uiowa.edu/vectorcore), and the University of Iowa High Resolution Mass Spectrometry Core Facility for the maintenance of mass spectrometers, especially the MALDI-TOF (https://hrmsf.research.uiowa.edu/). We are grateful to Liping Yu, Gang Wu, Nan Jia, Yifan Yu, Dongli Lu, Mohui Wei and Nianyi Zeng for proof reading and useful suggestions. We are grateful to the Scientific Editing and Research Communication Core at the University of Iowa Carver College of Medicine. We are also grateful to Amber Mower and Jaeda Harmon for assistance with administrative support. Sanam Zarei was a Howard Hughes Medical Institute Research Fellow.

## Ethics

Animal experimentation: This study was performed in strict accordance with the recommendations in the Guide for the Care and Use of Laboratory Animals of the National Institutes of Health. All animal experiments were approved by the Institutional Animal Care and Use Committee (IACUC) protocols of the University of Iowa (#0081122).

---

**TABLE 1.**
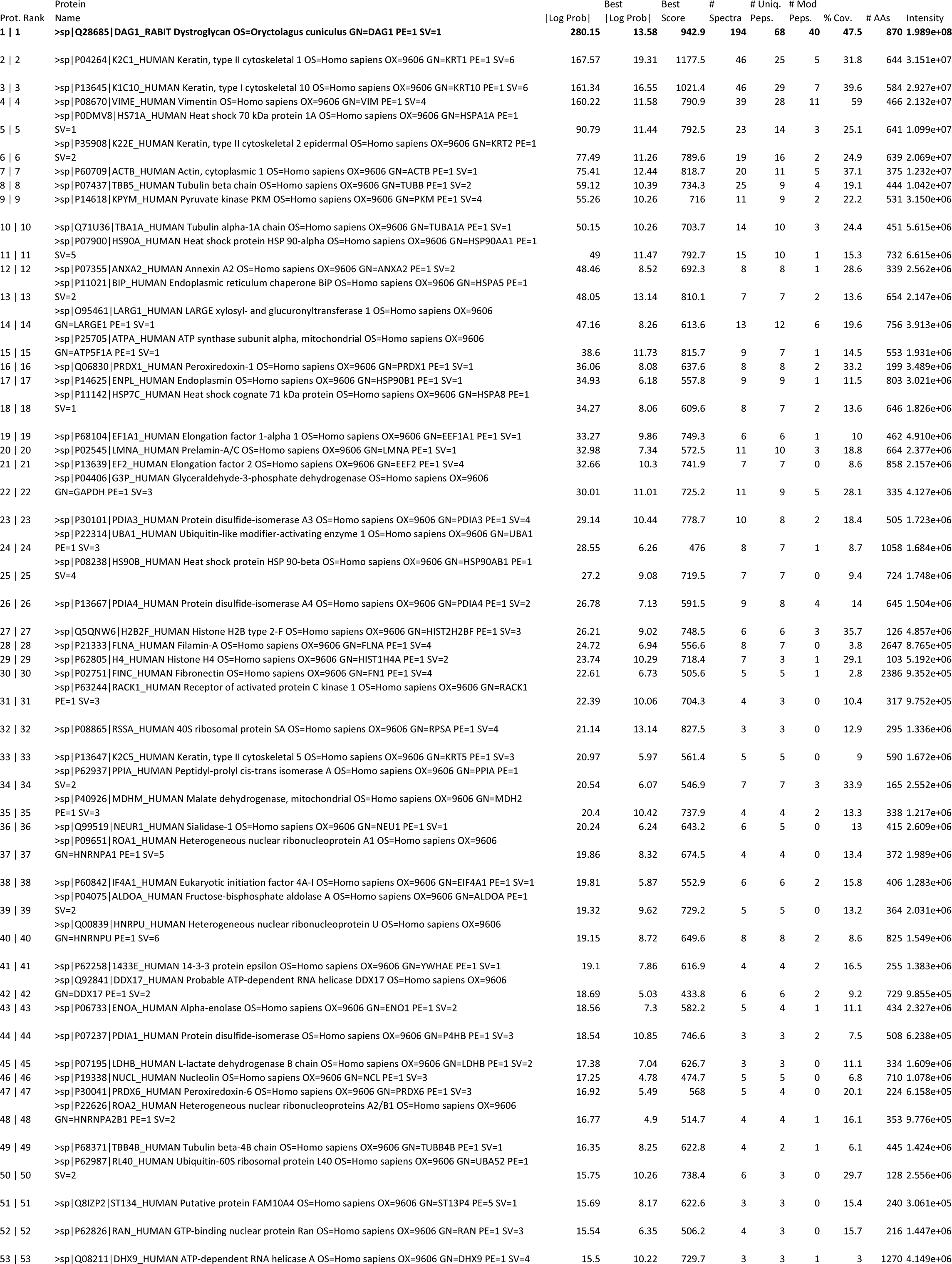
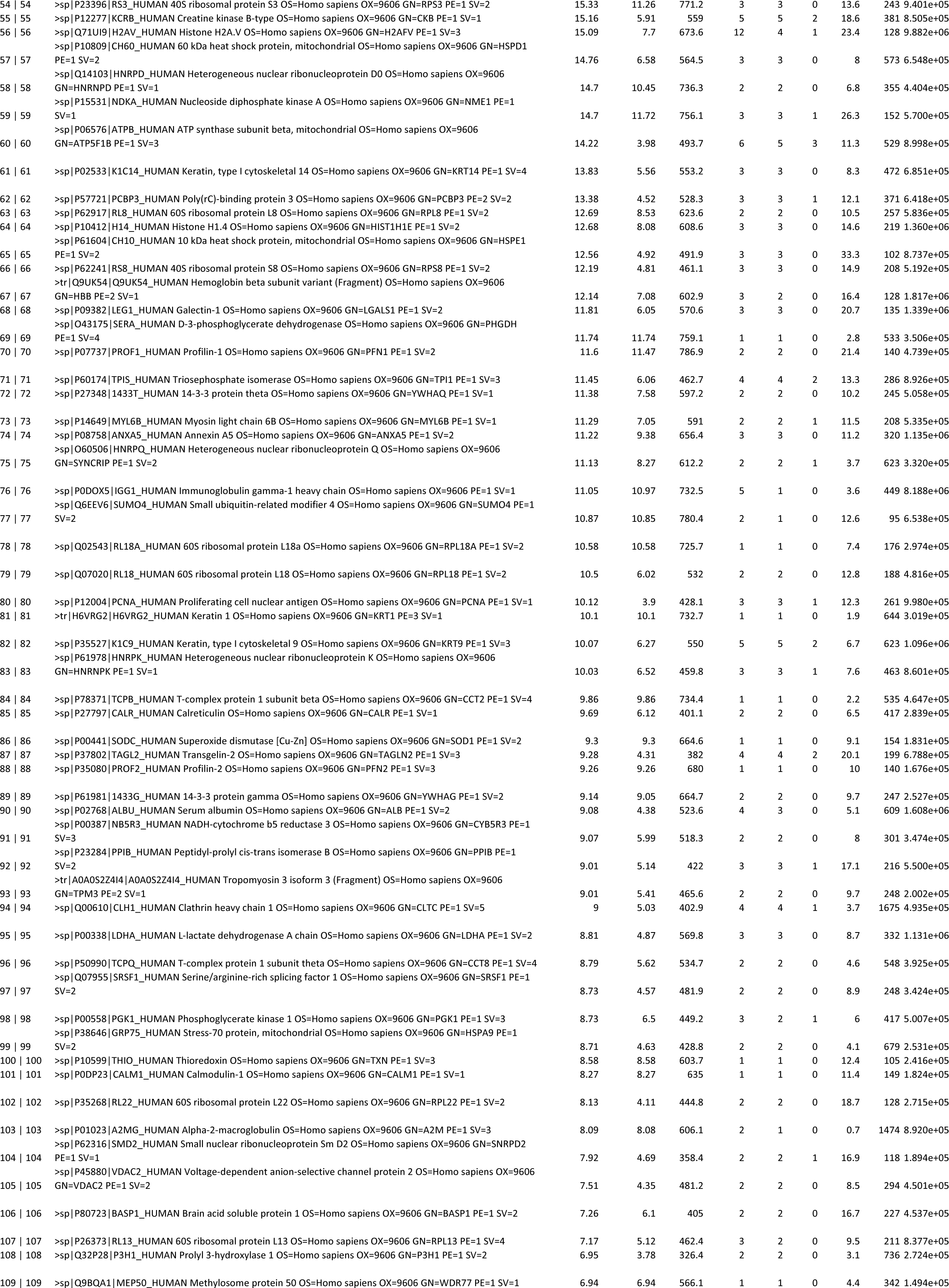
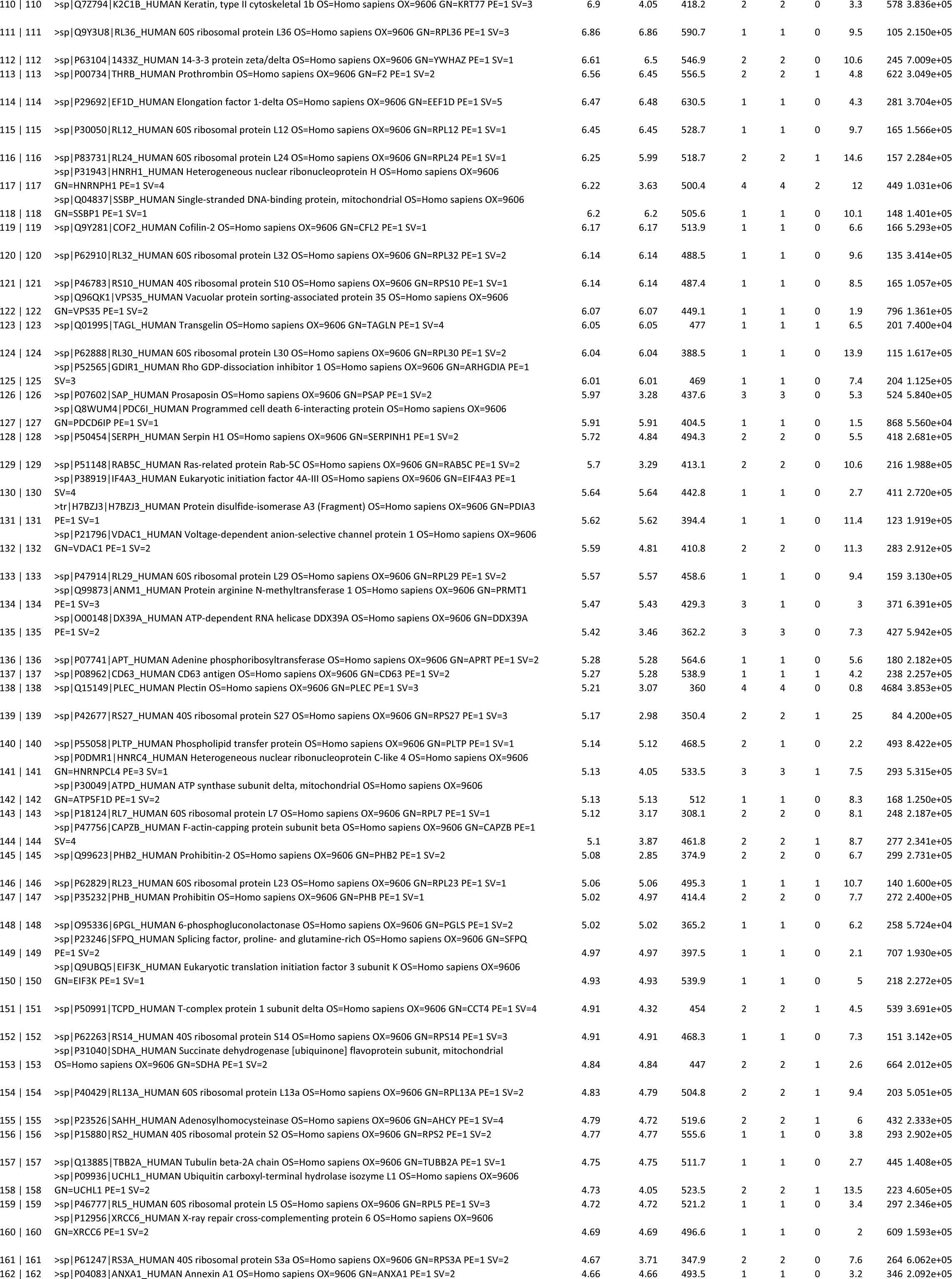
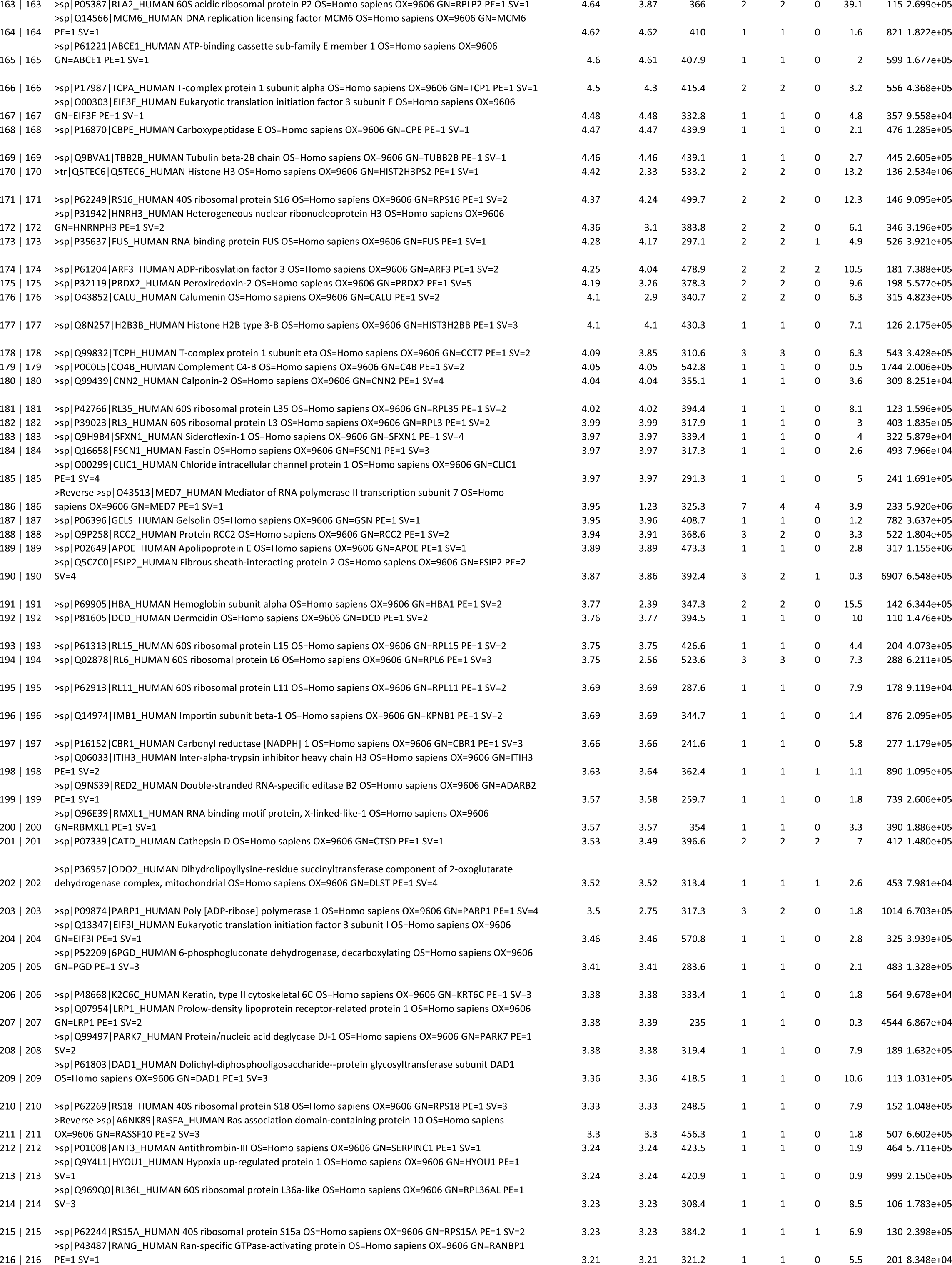
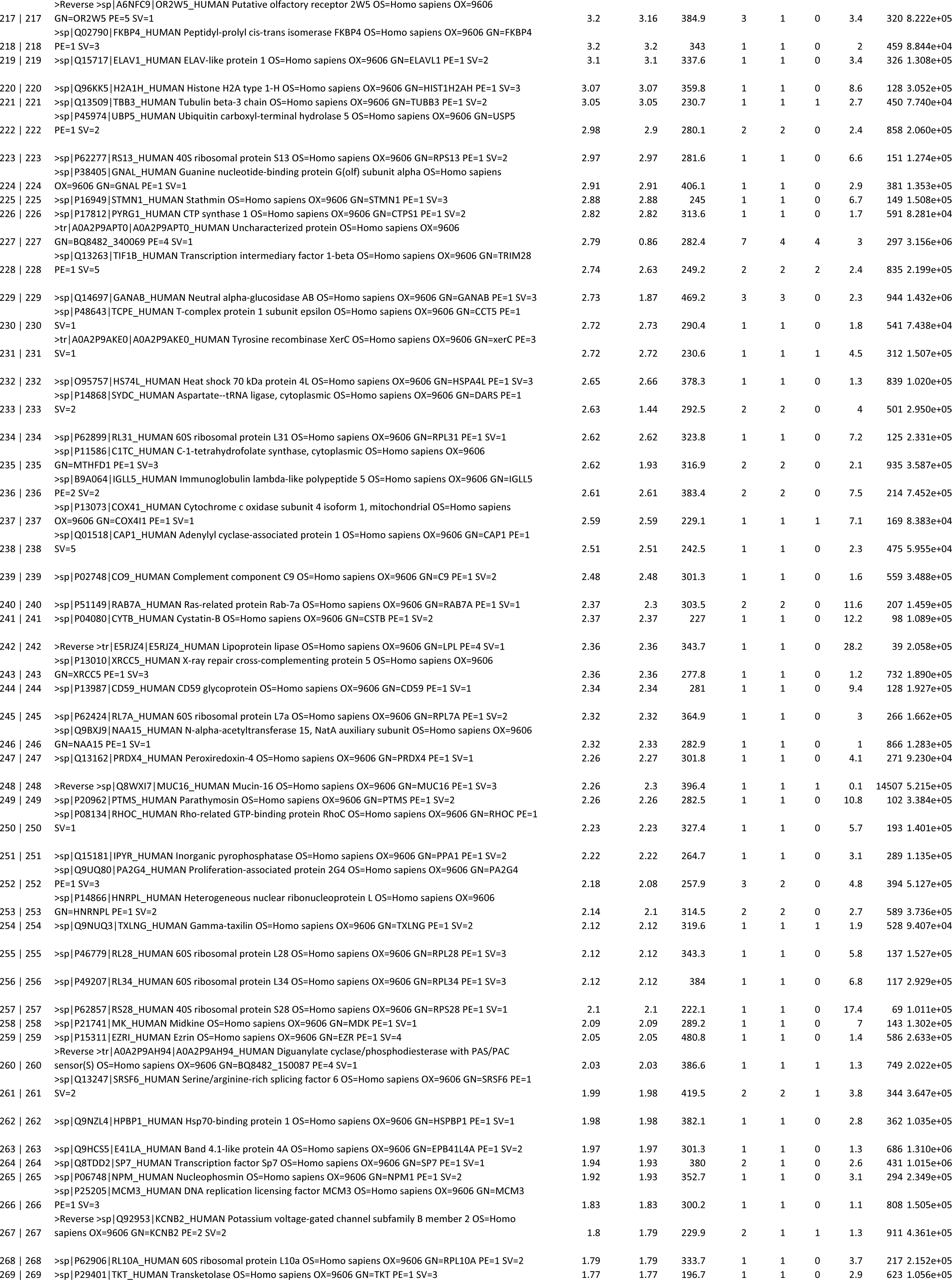
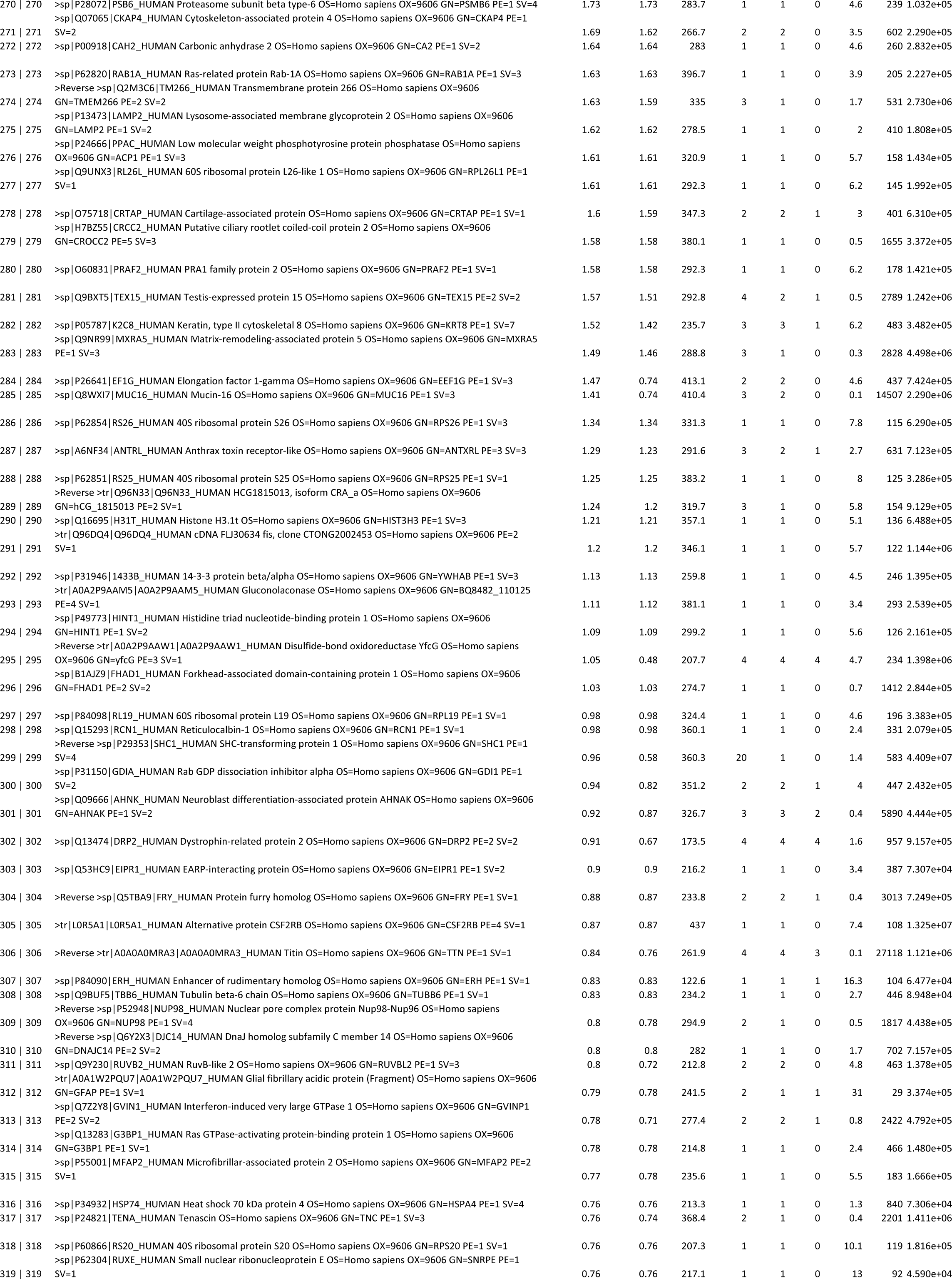
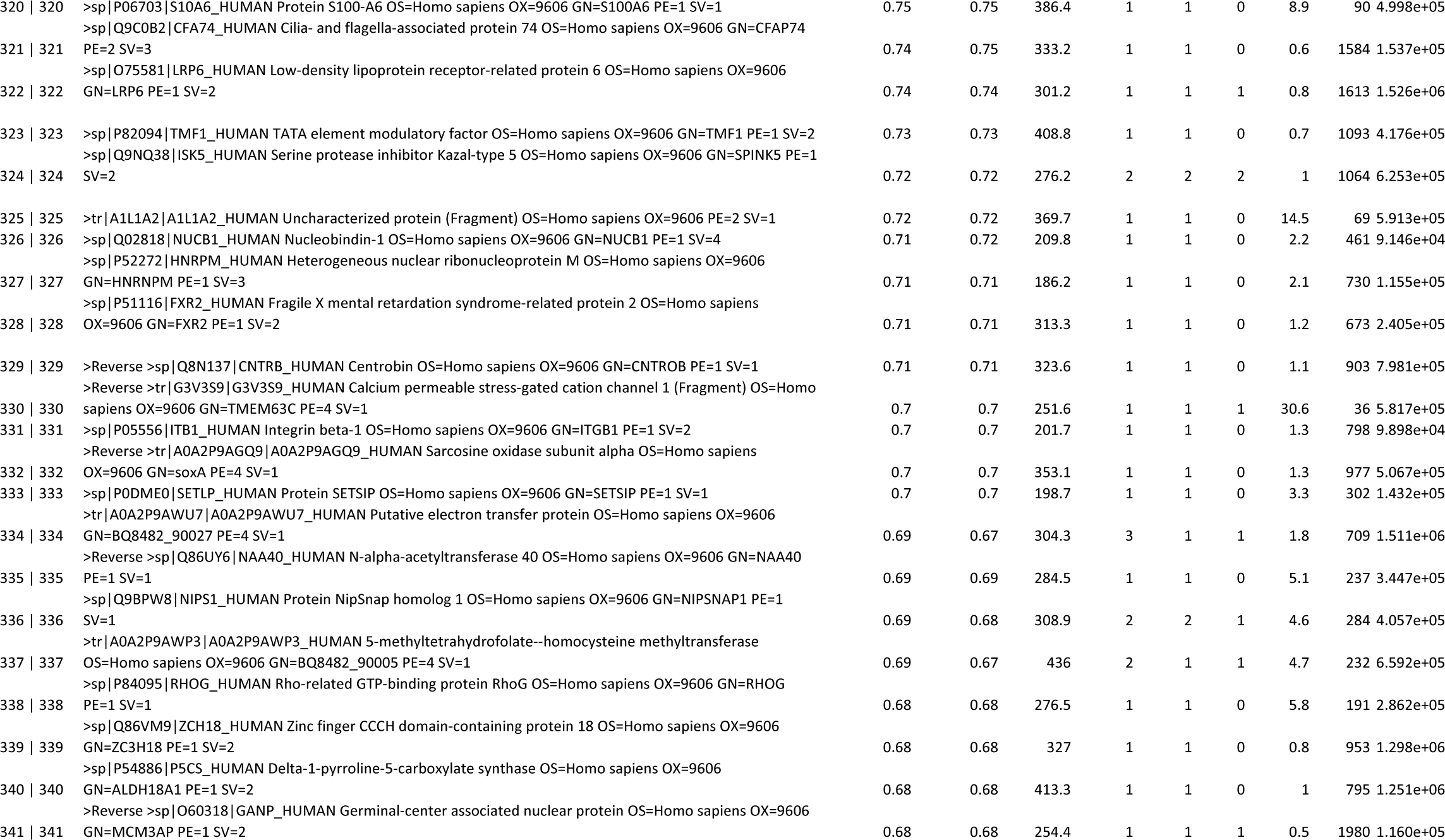
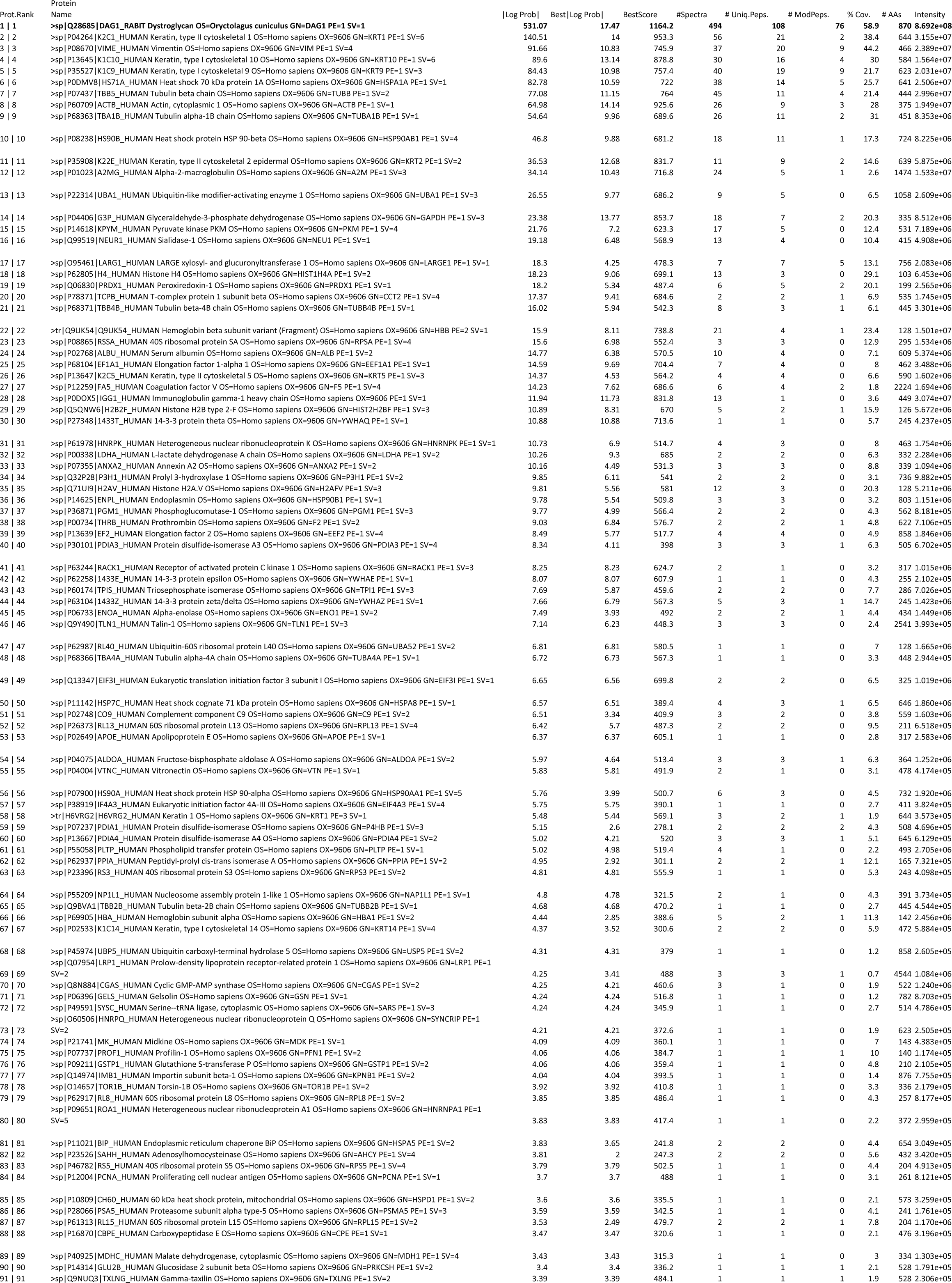
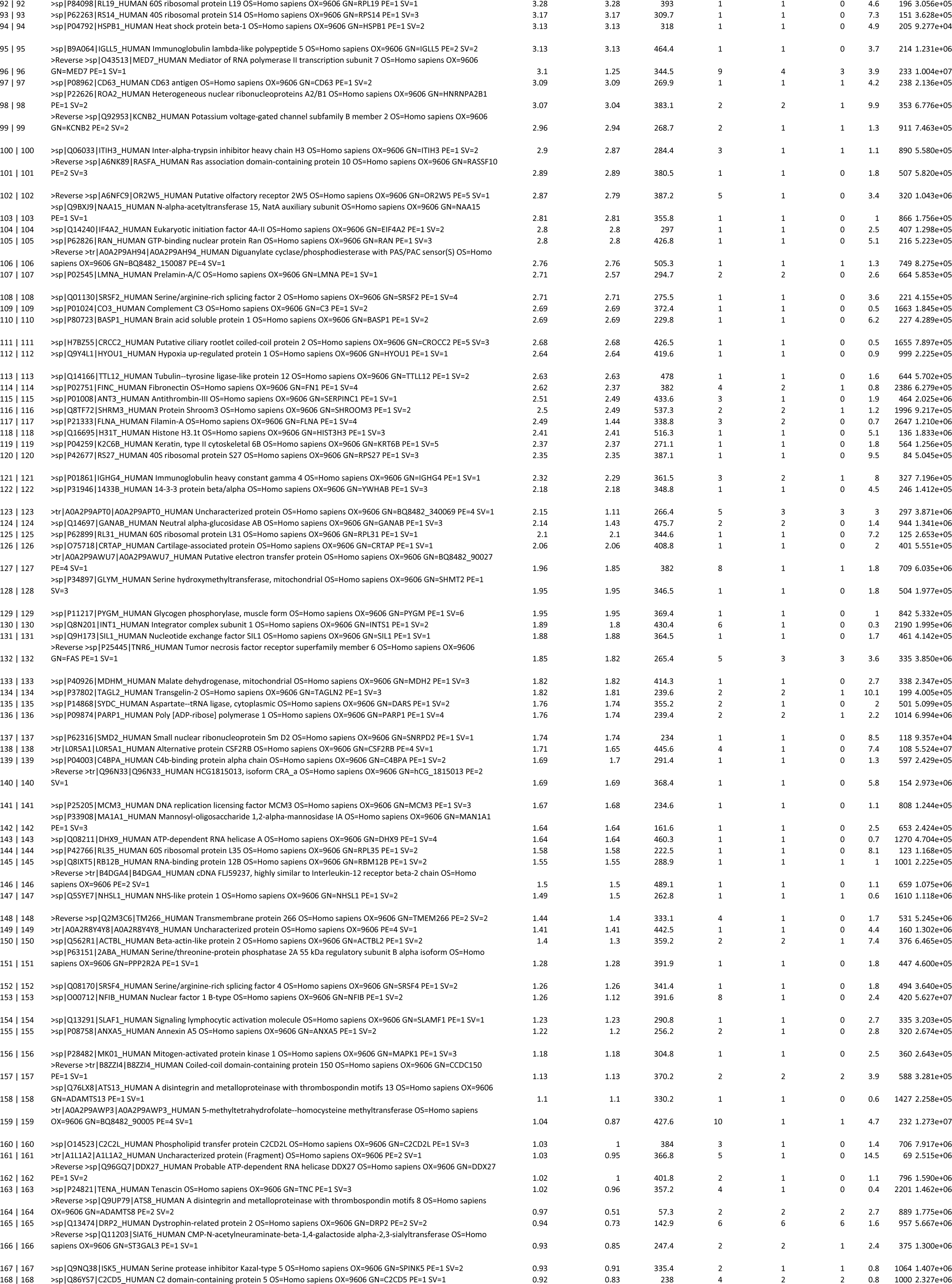
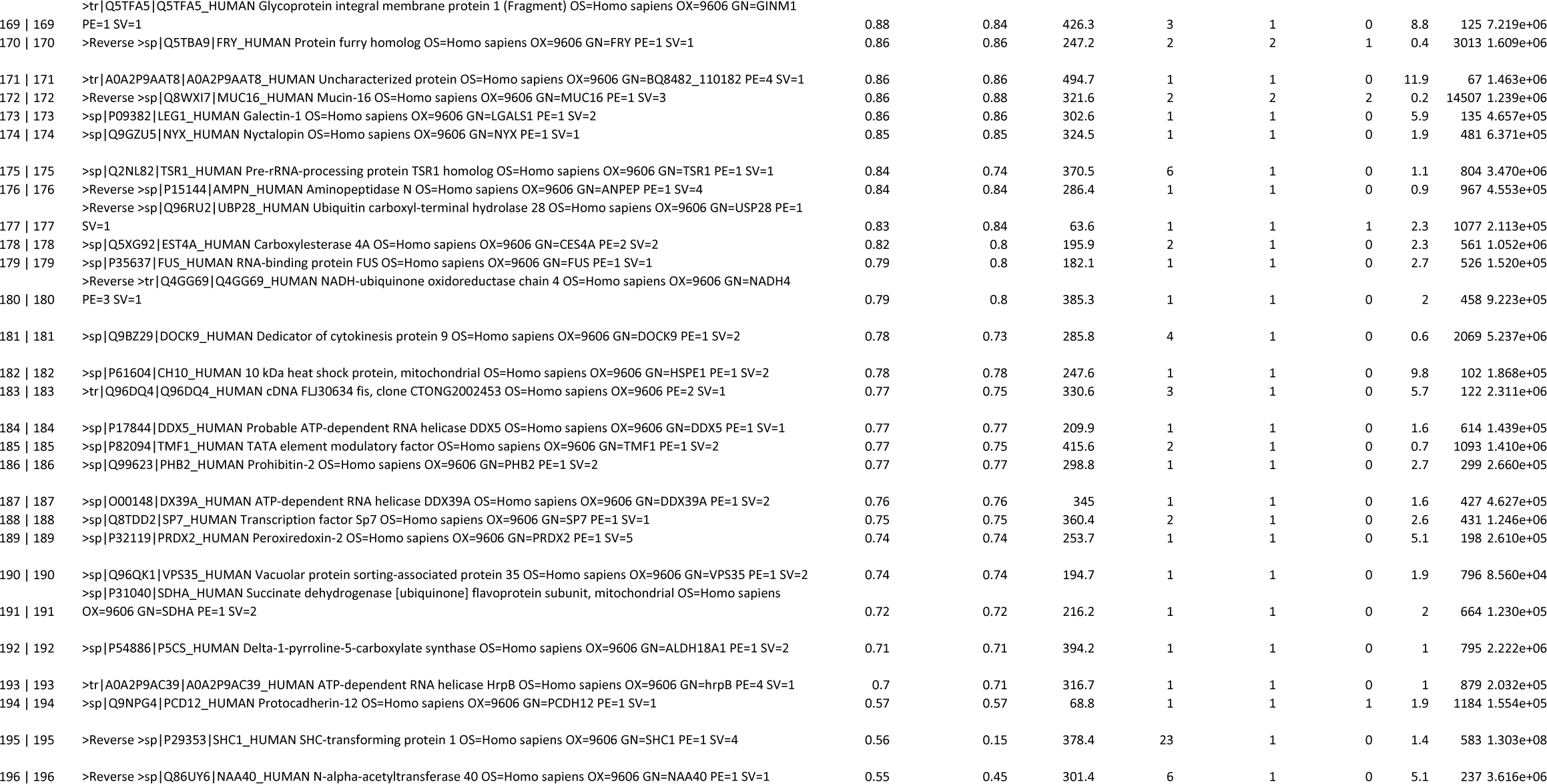

**Table 2.**

|  |  |  |  |  |
| --- | --- | --- | --- | --- |
|  | 315.317 |  |  |  |
|  | Observed # | Expected # | Observed % | Expected % |
| Het | 118 | 110 | 0.536 | 0.5 |
| WT | 80 | 55 | 0.364 | 0.25 |
| **Mut** | **22** | **55** | **0.1** | **0.25** |
| Total: | 220 | 220 | 1 | 1 |
|  | 377.379 |  |  |  |
|  | Observed # | Expected # | Observed % | Expected % |
| Het | 111 | 119.5 | 0.464 | 0.5 |
| WT | 75 | 59.75 | 0.314 | 0.25 |
| Mut | 53 | 59.75 | 0.222 | 0.25 |
| Total: | 239 | 239 | 1 | 1 |

**Table 3.**
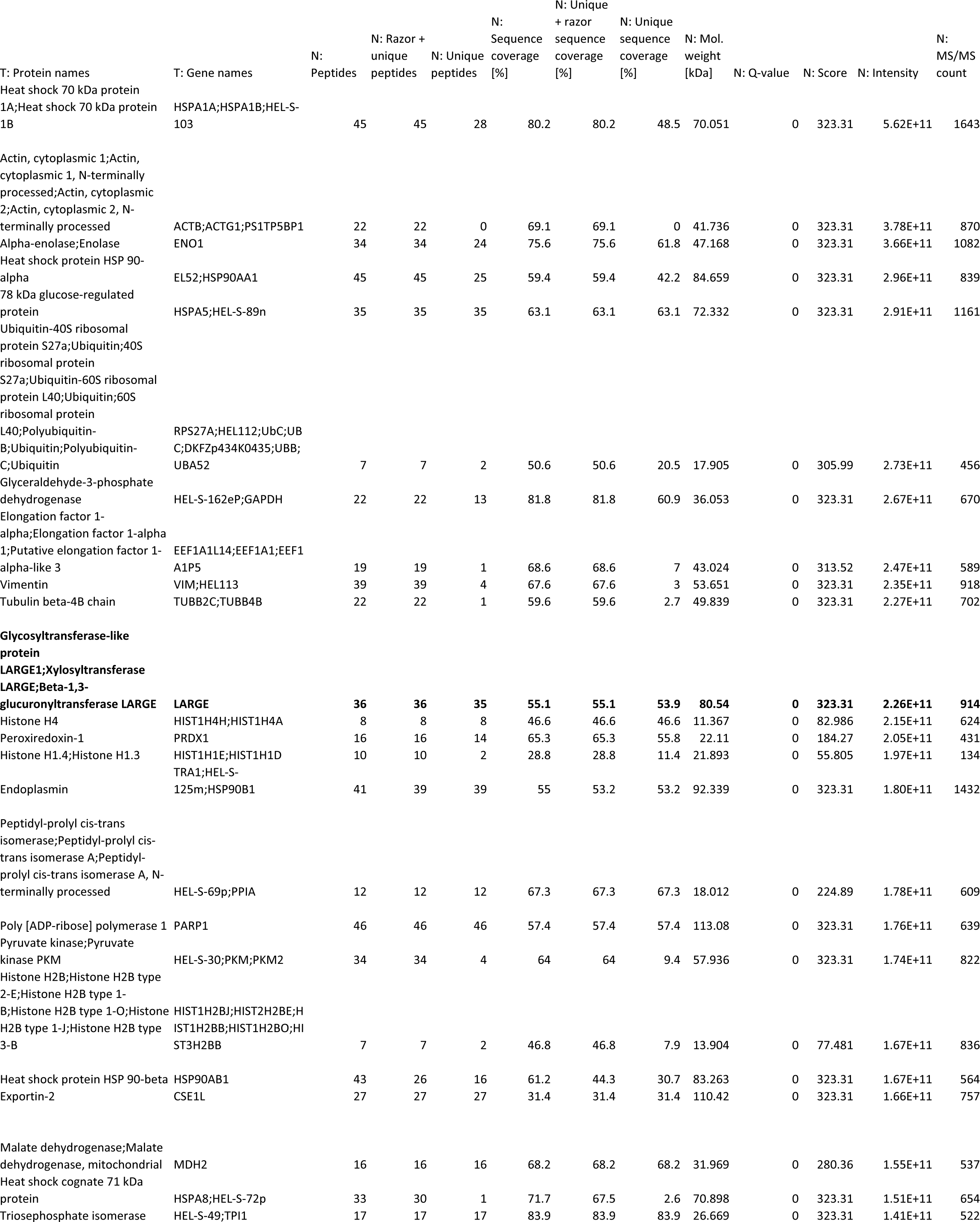
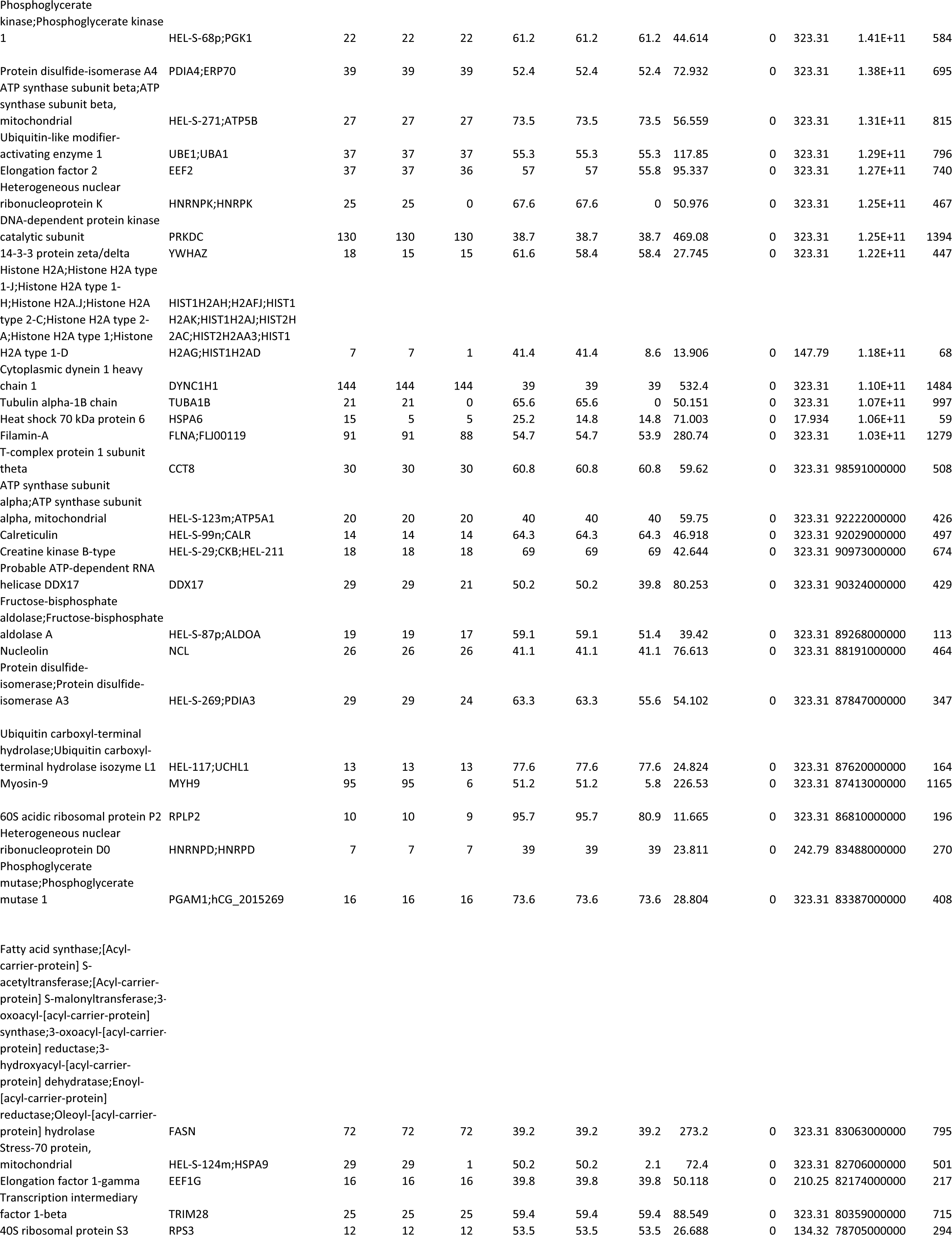
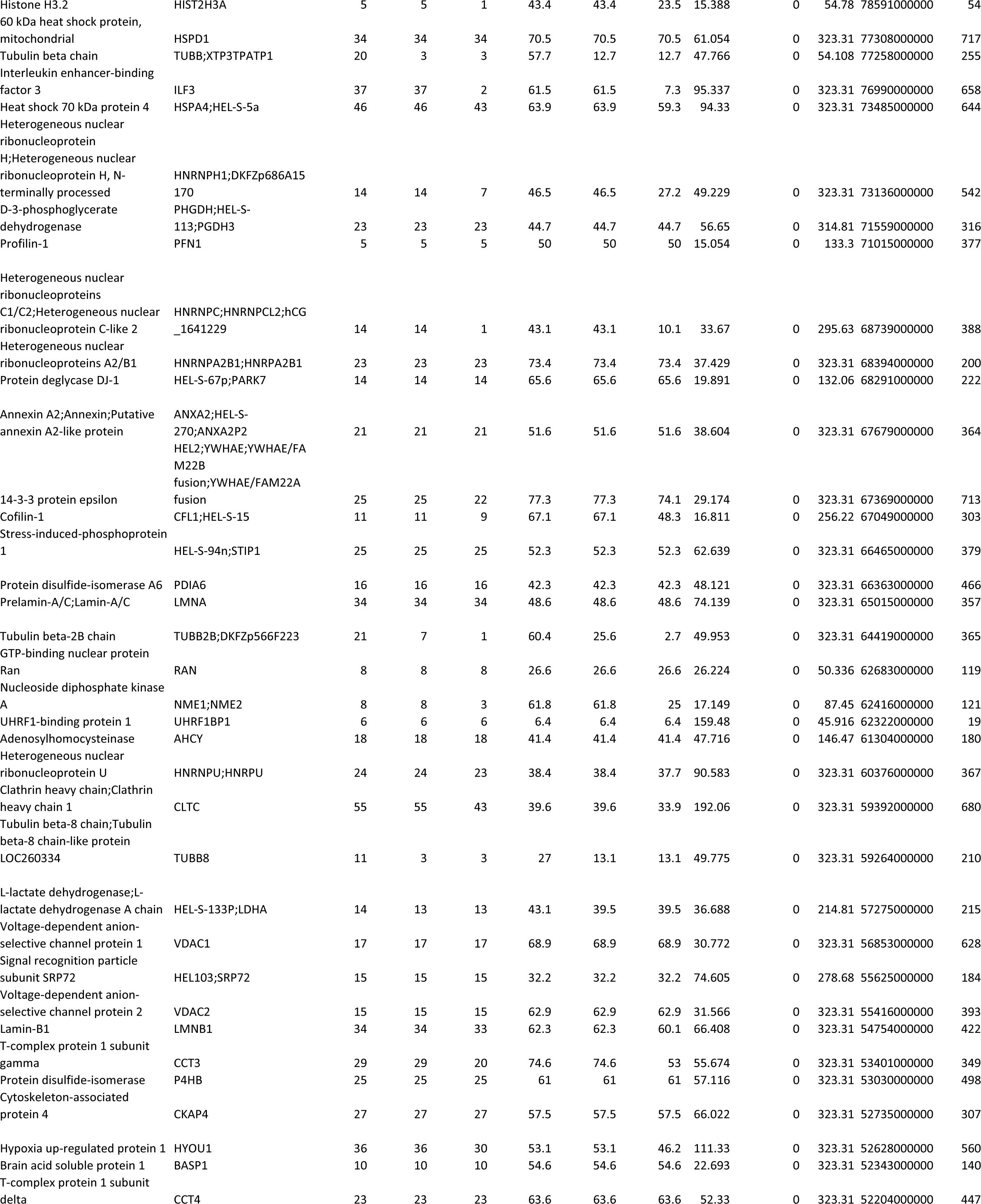
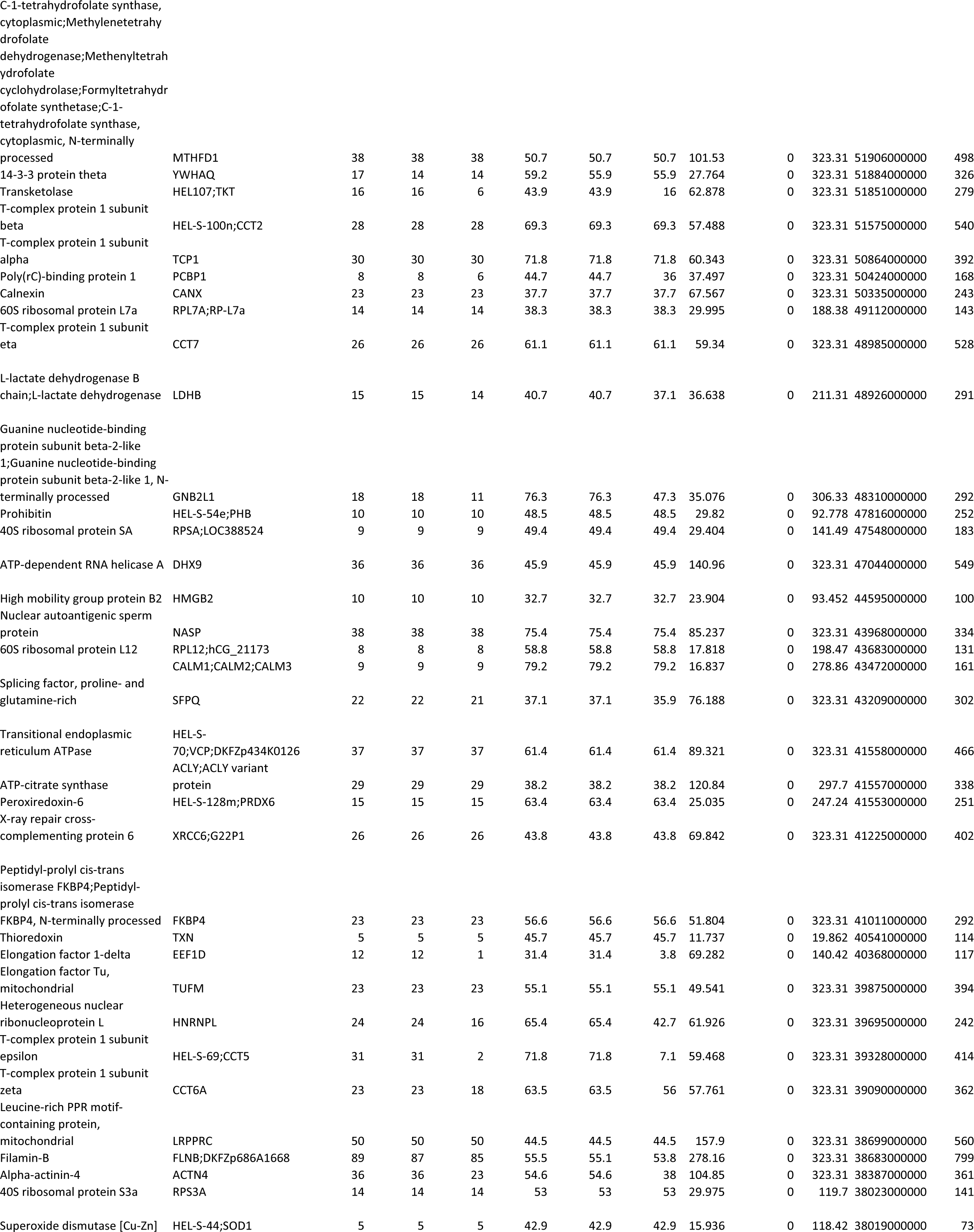
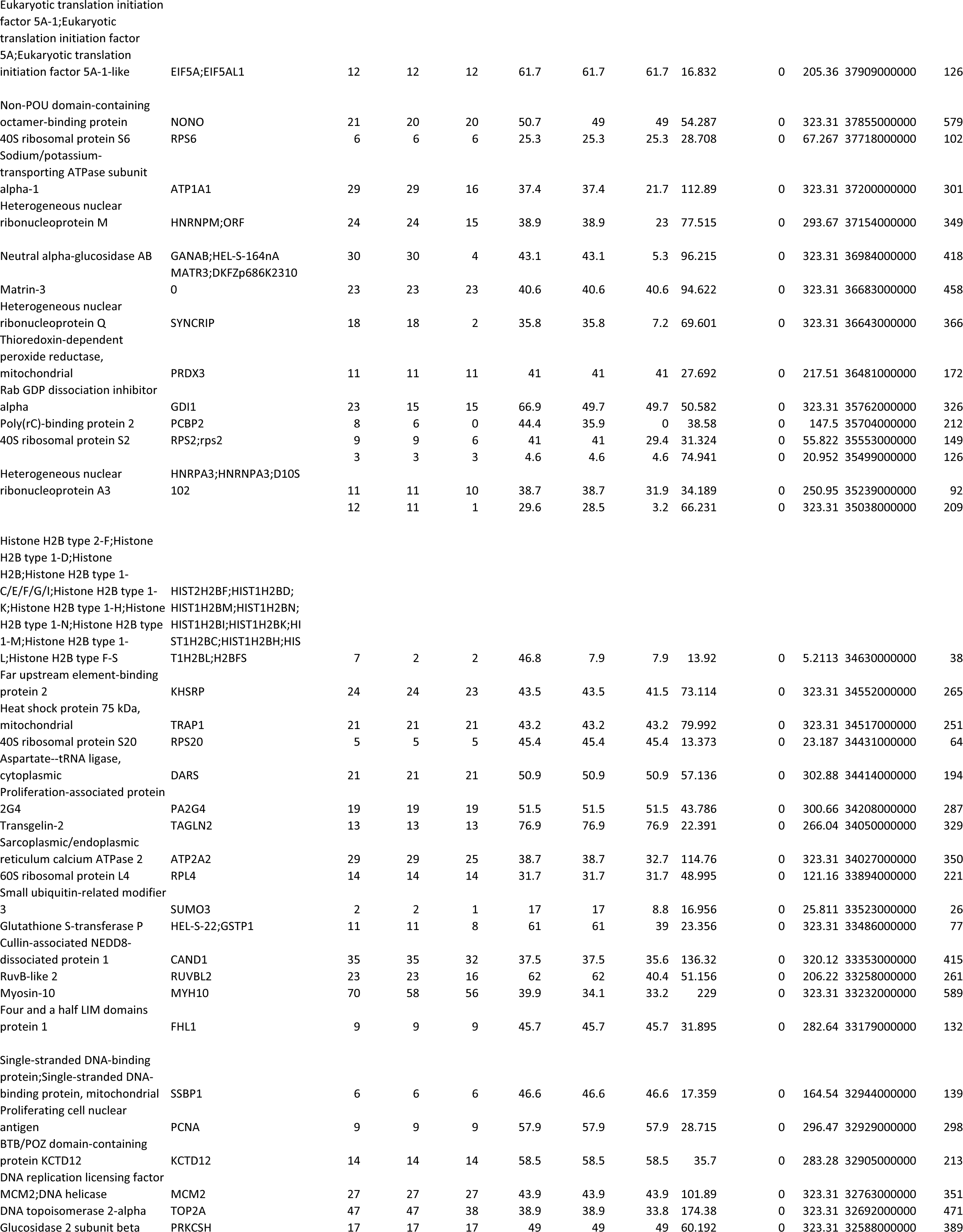
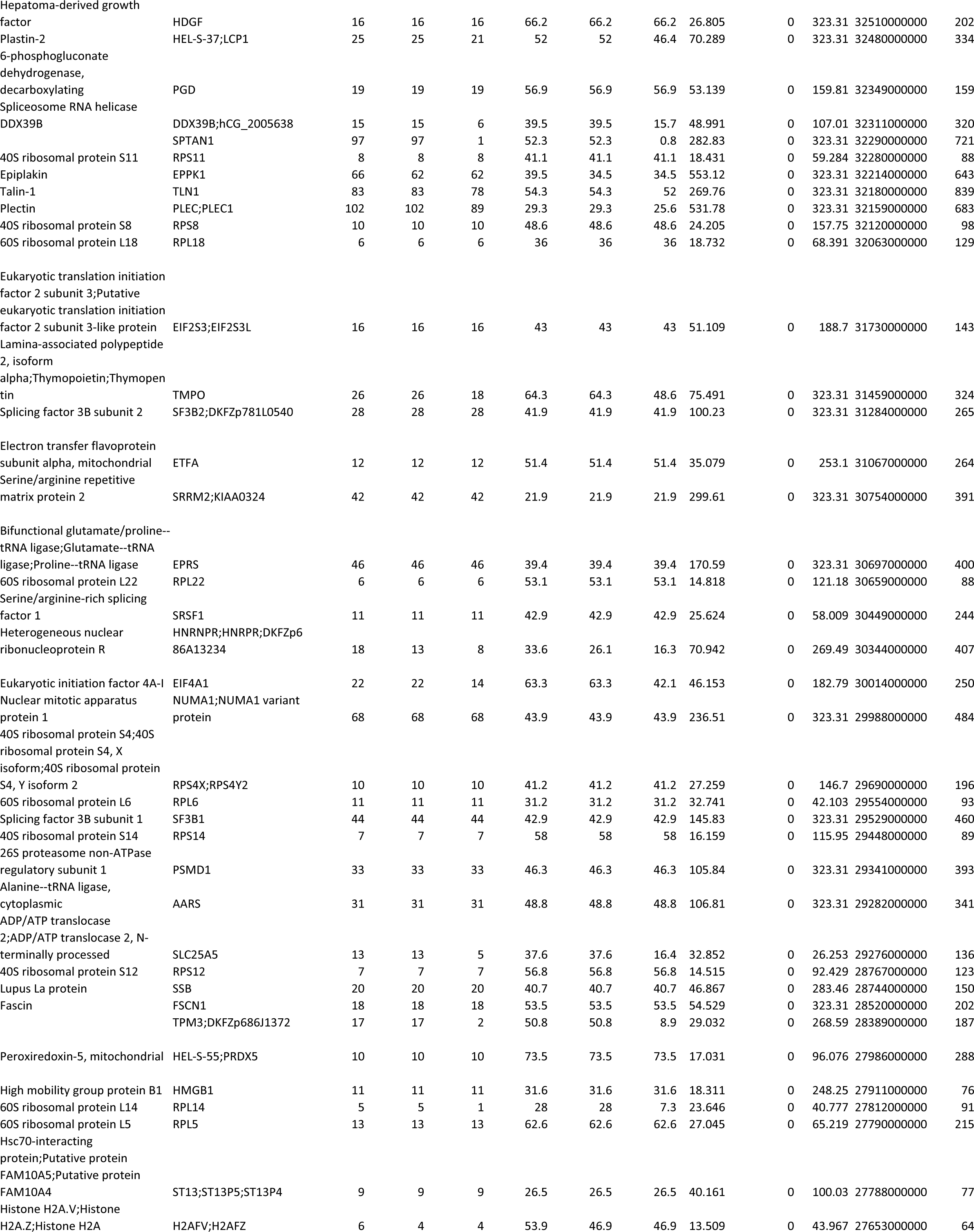
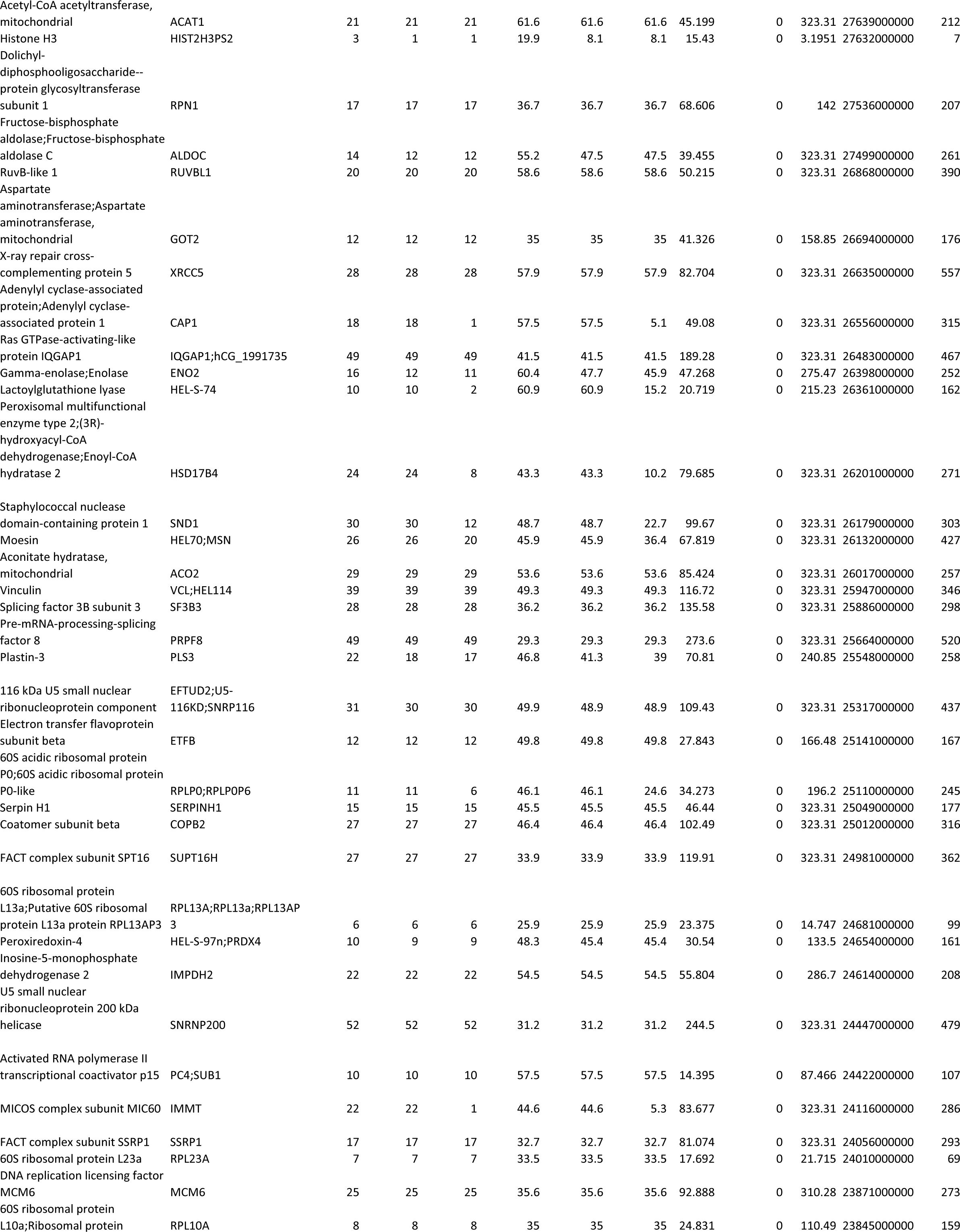
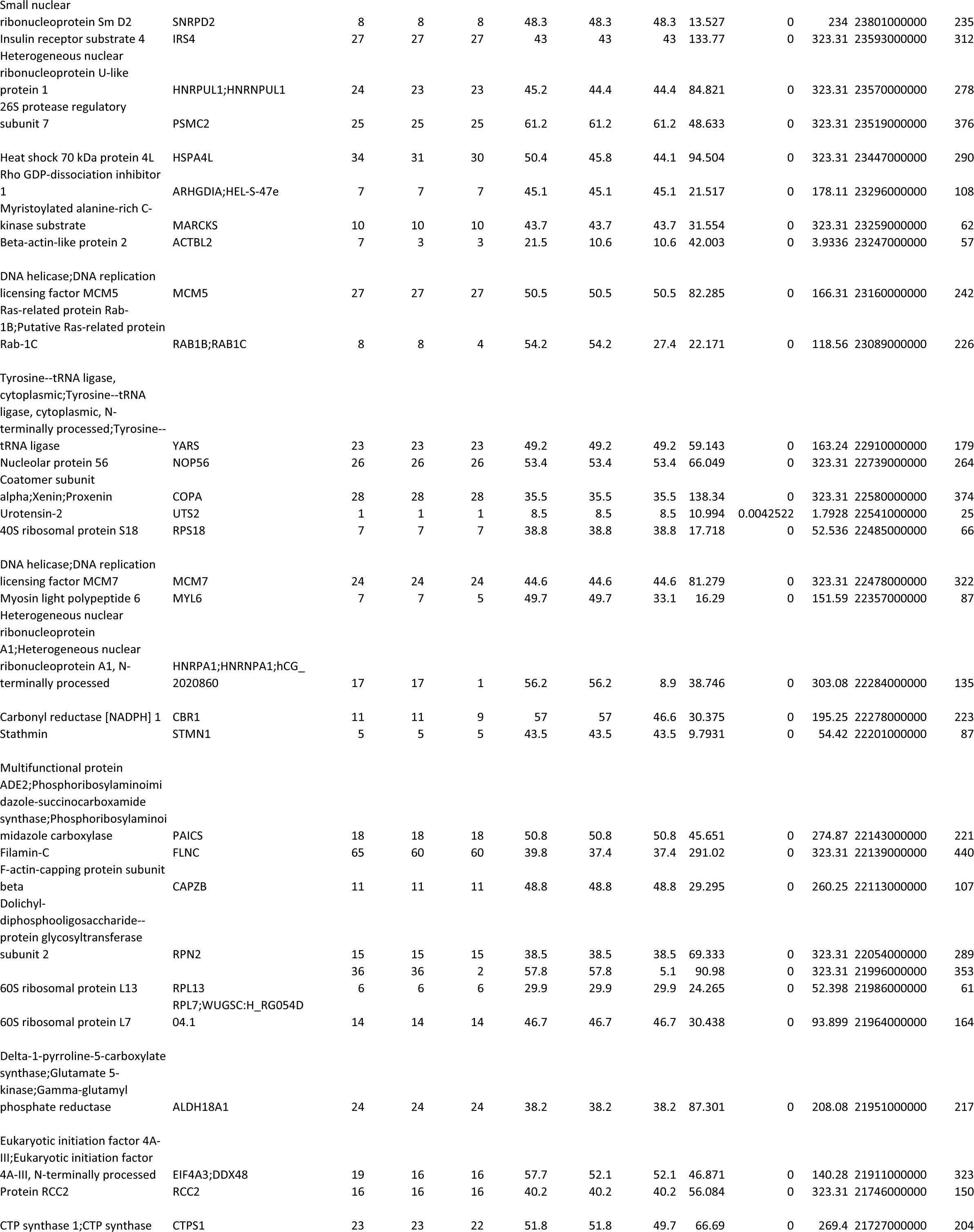
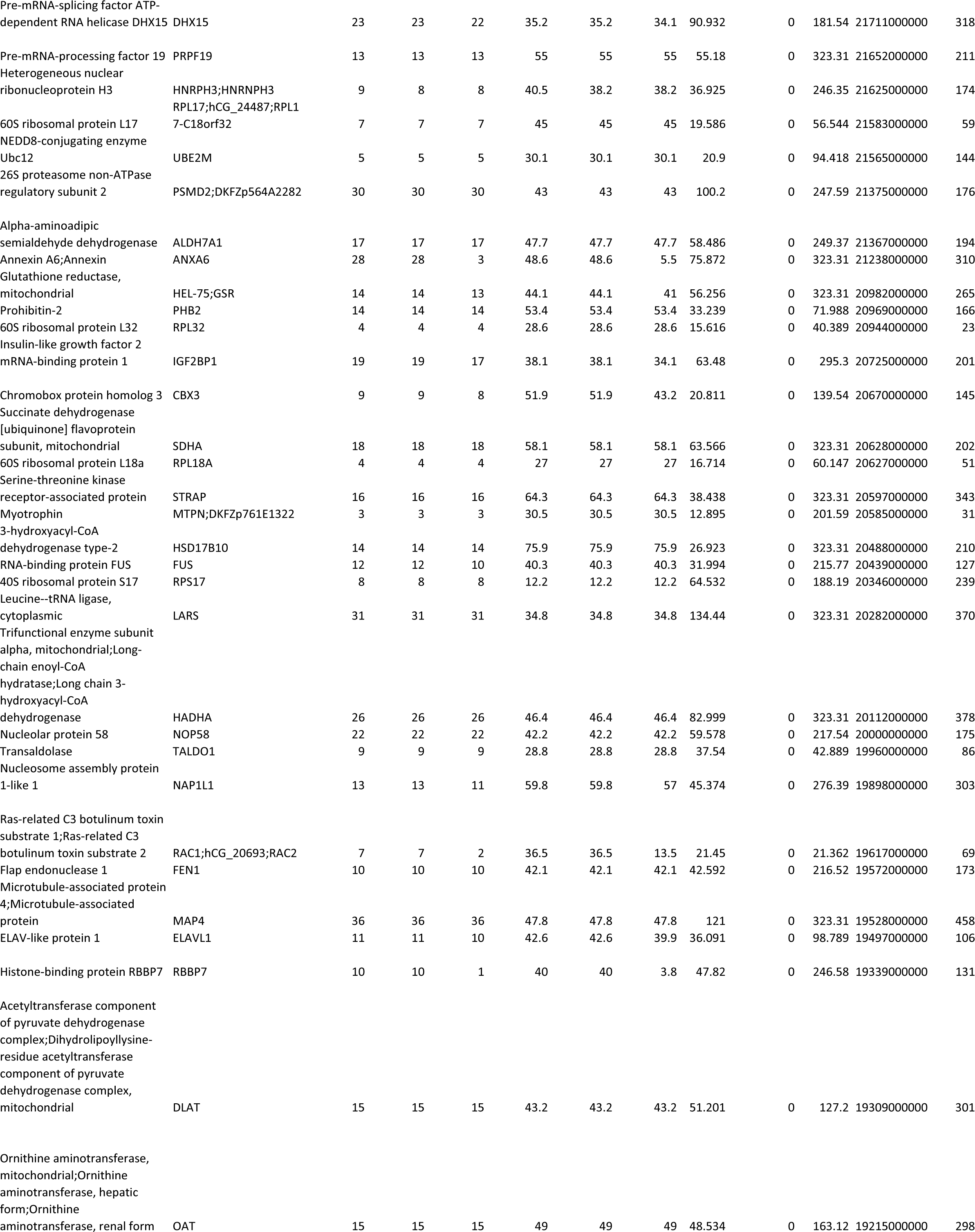
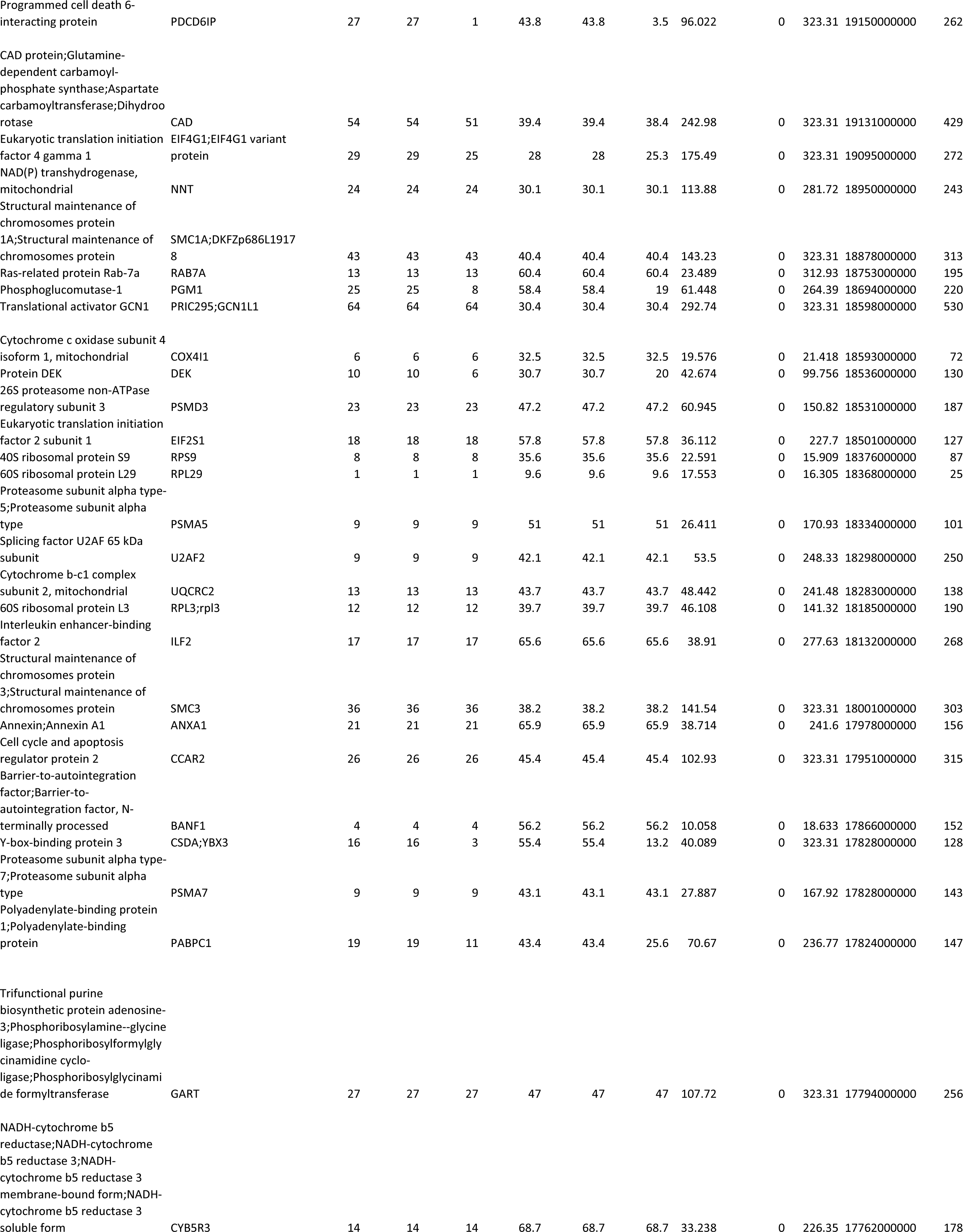
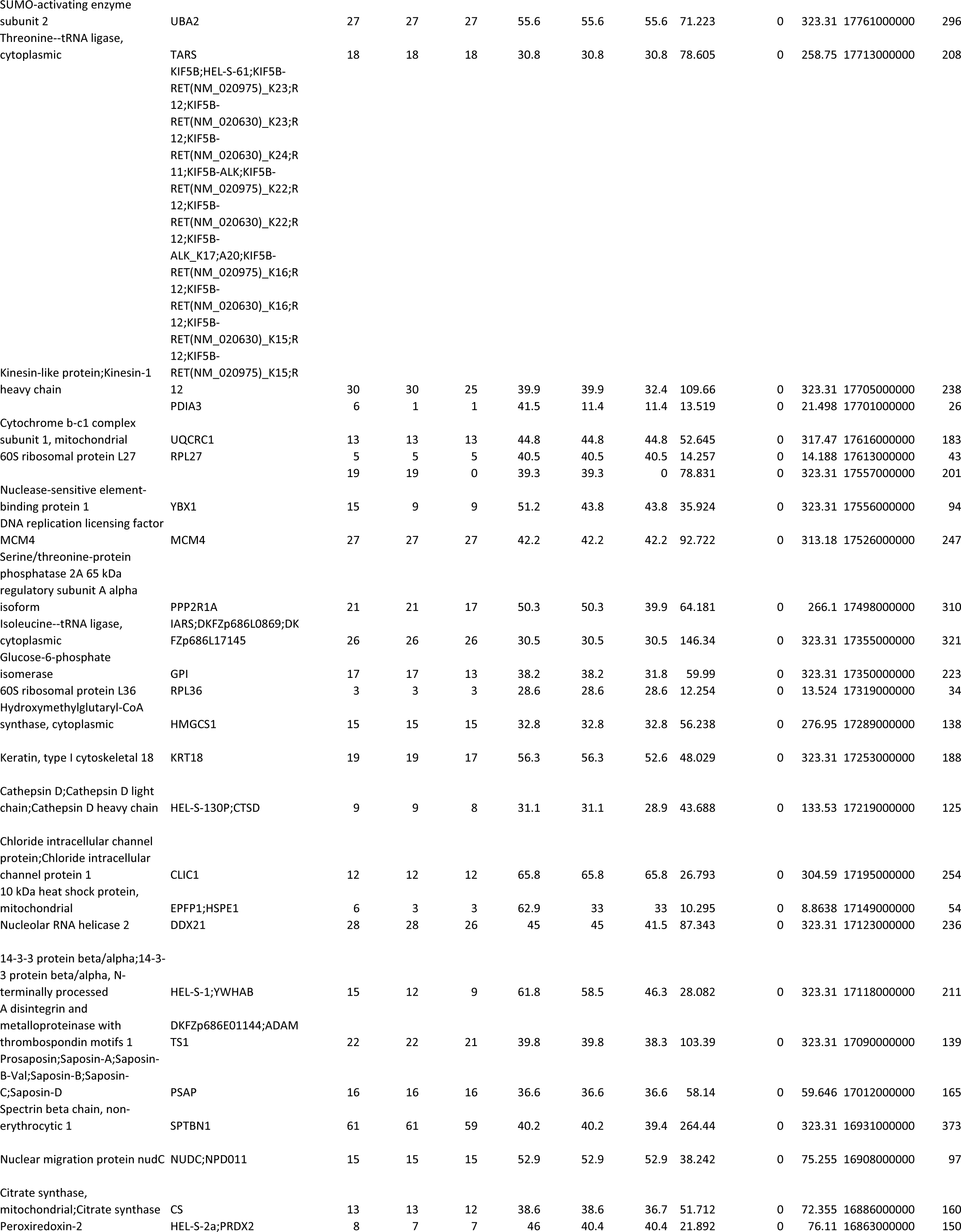
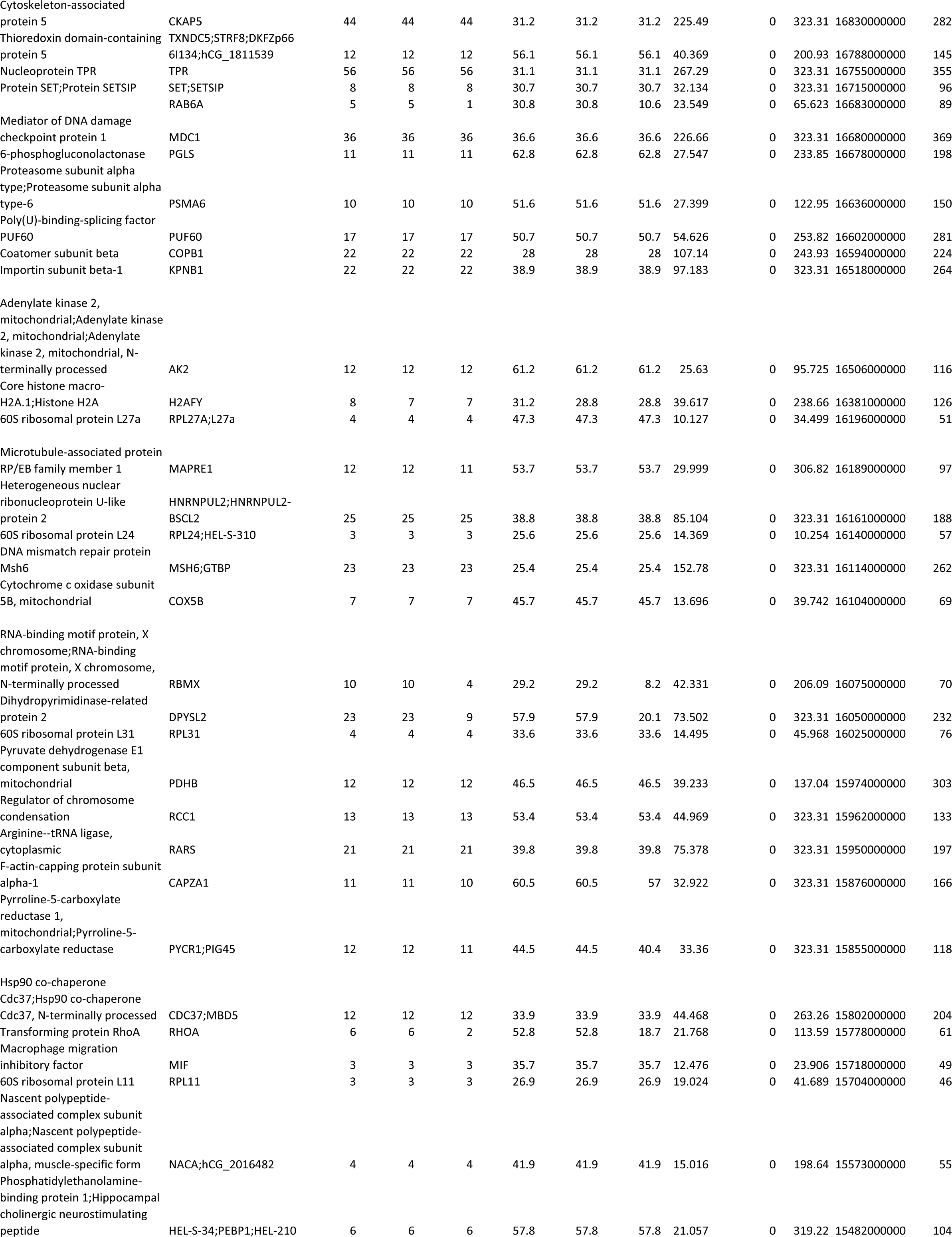
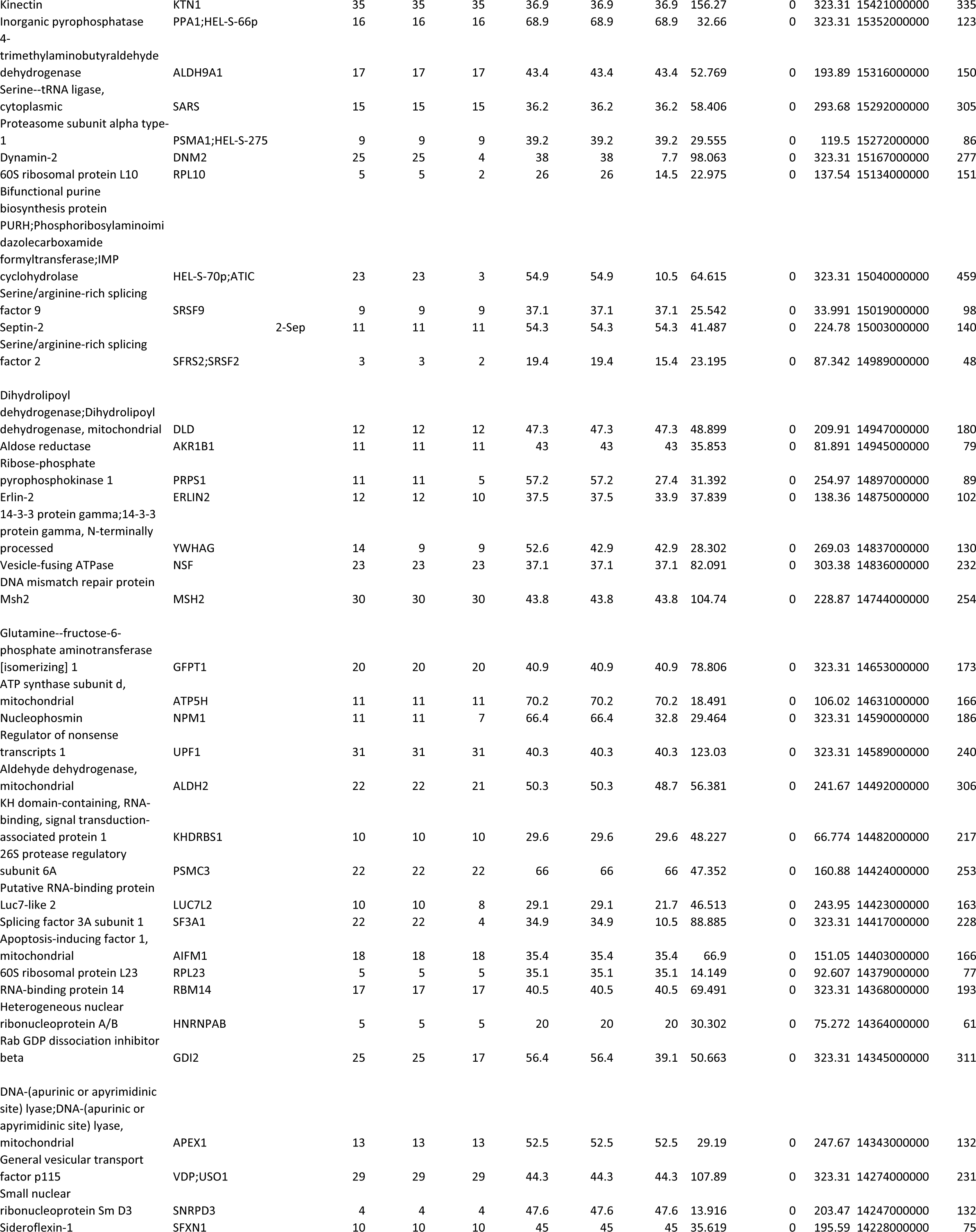
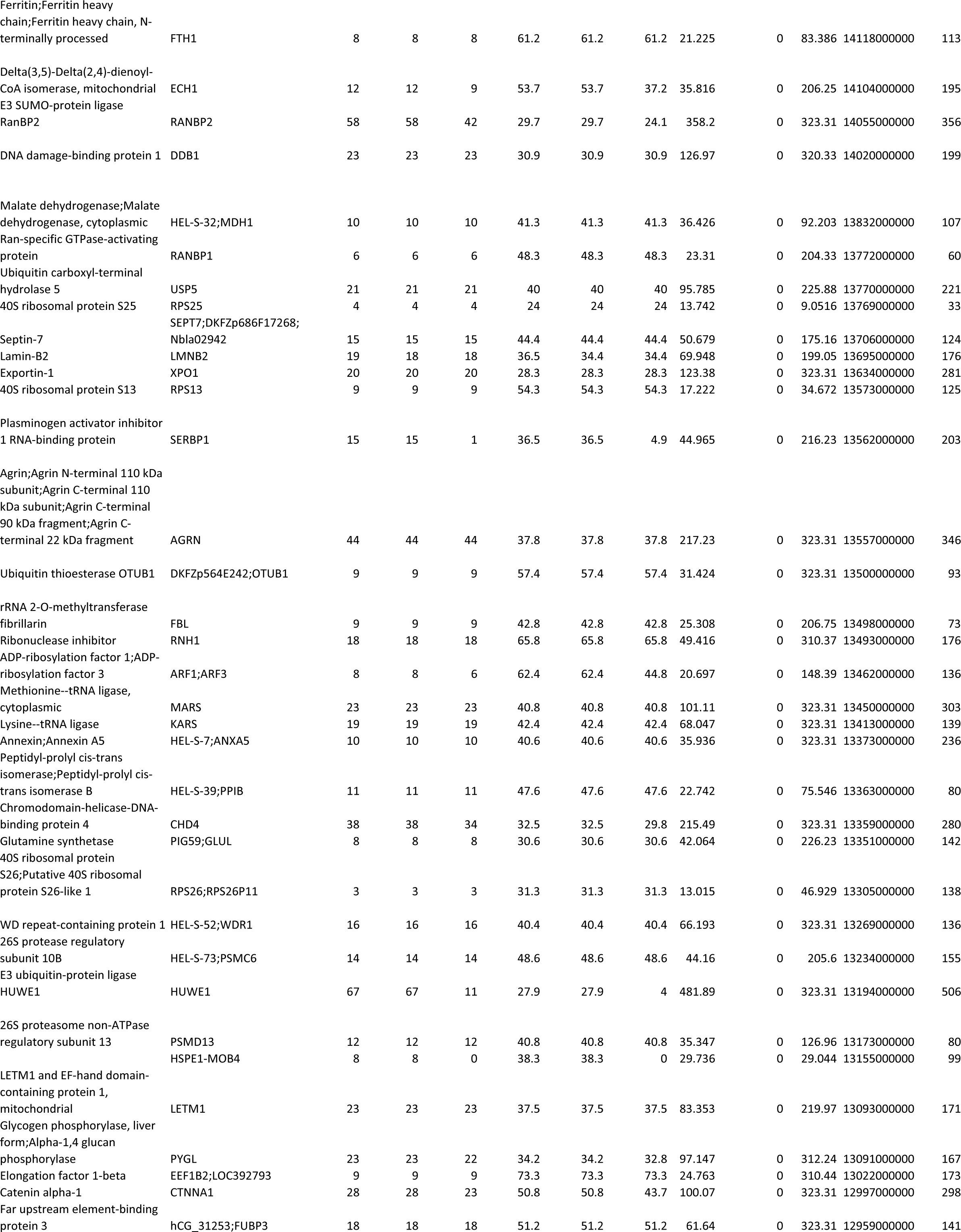
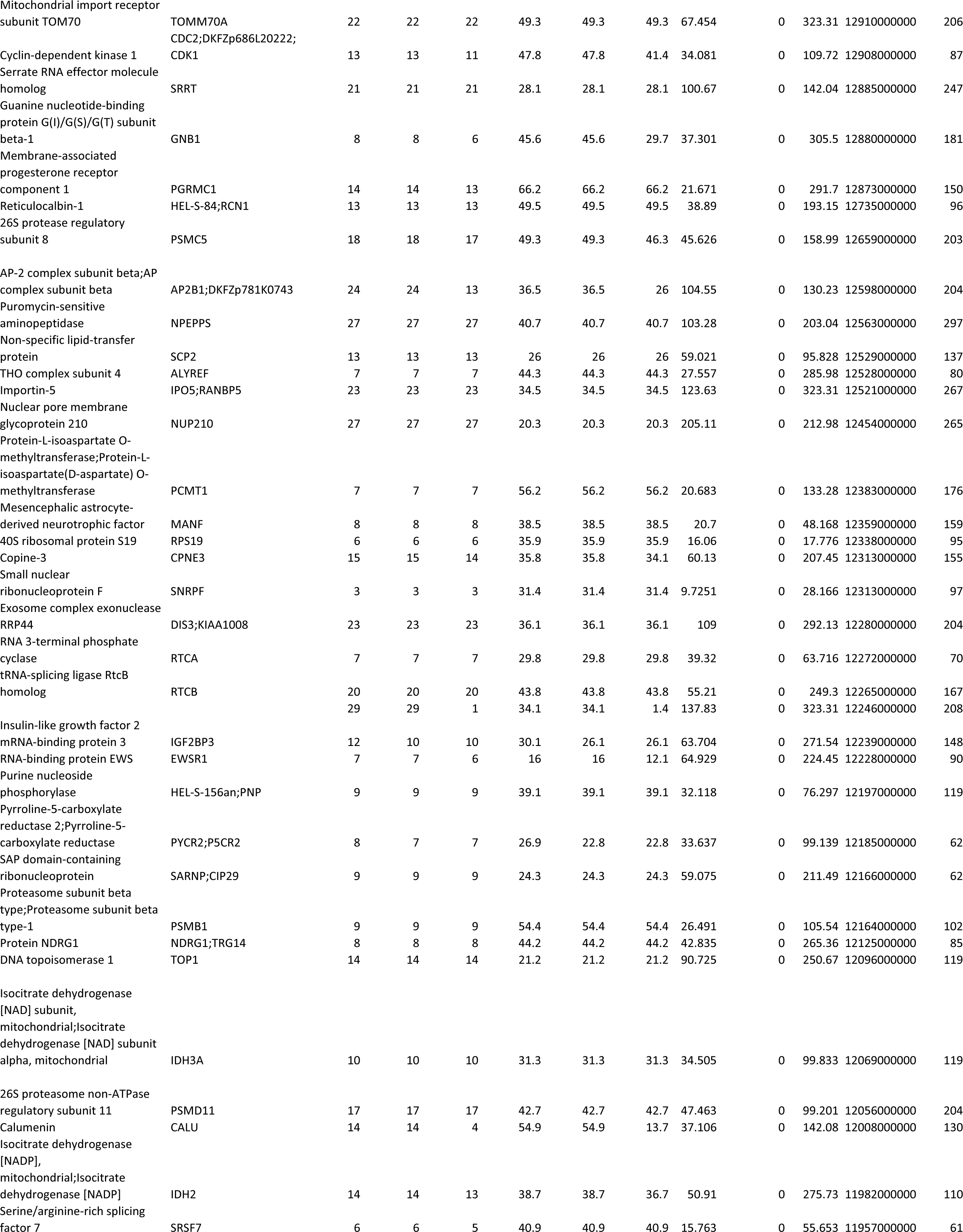
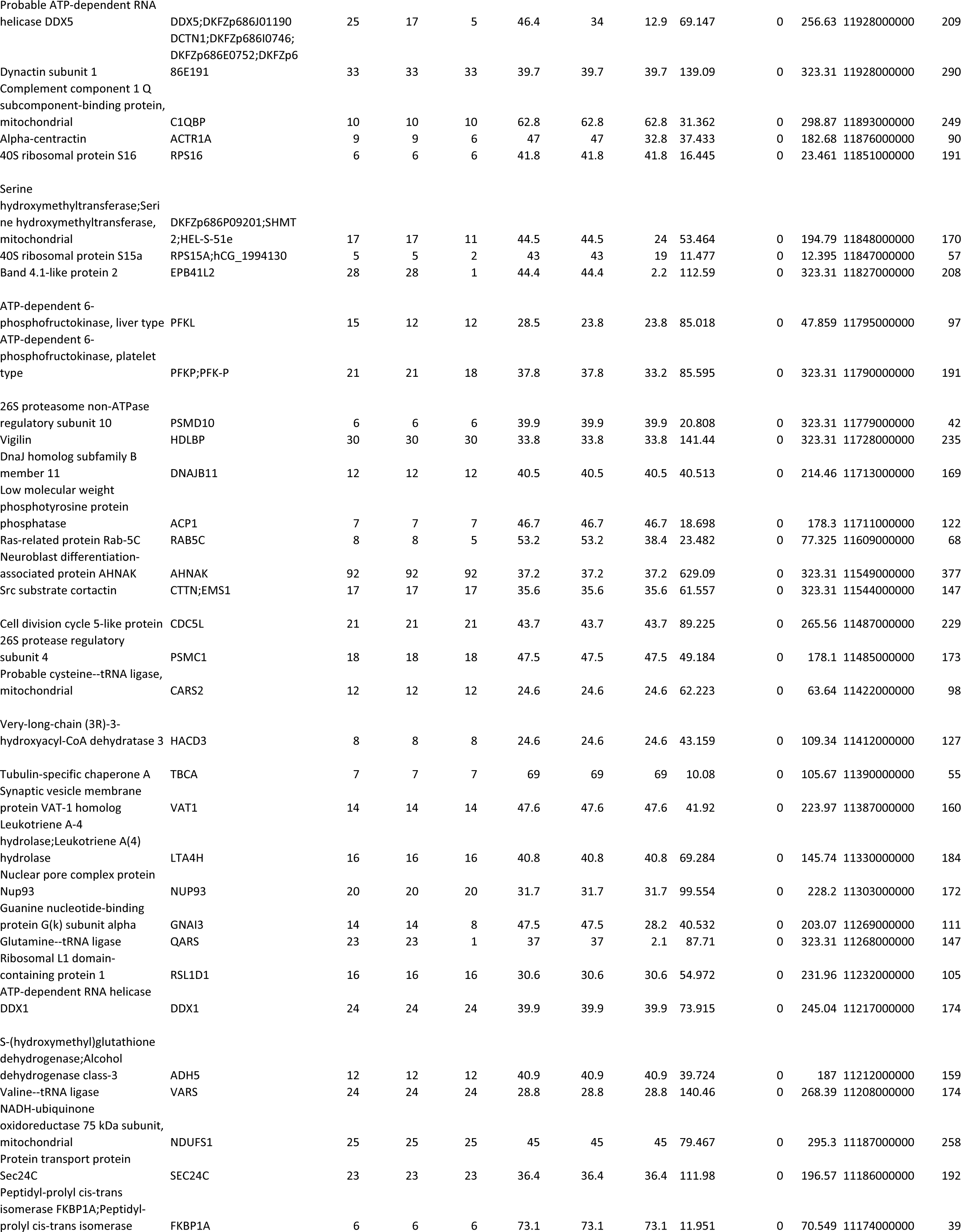
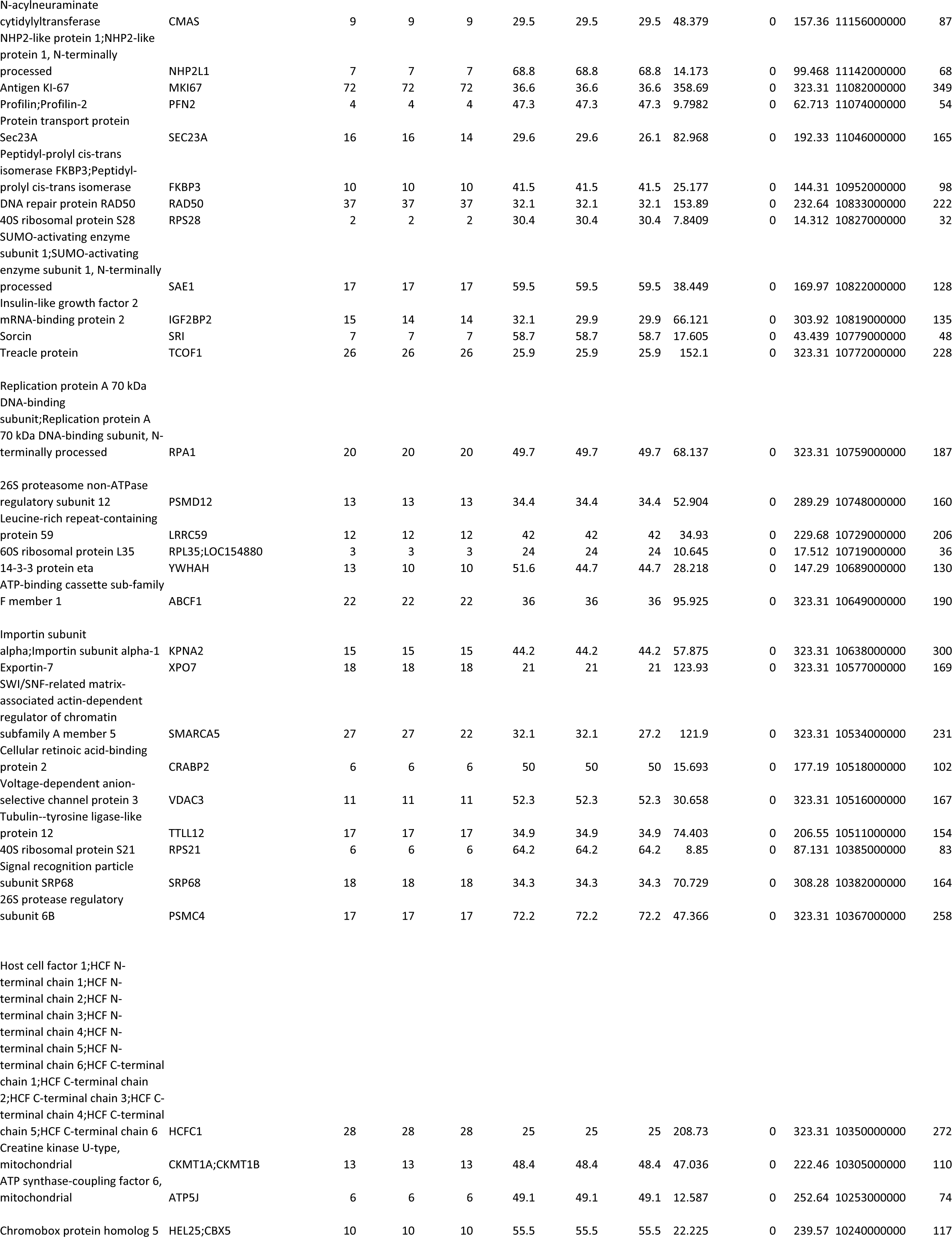
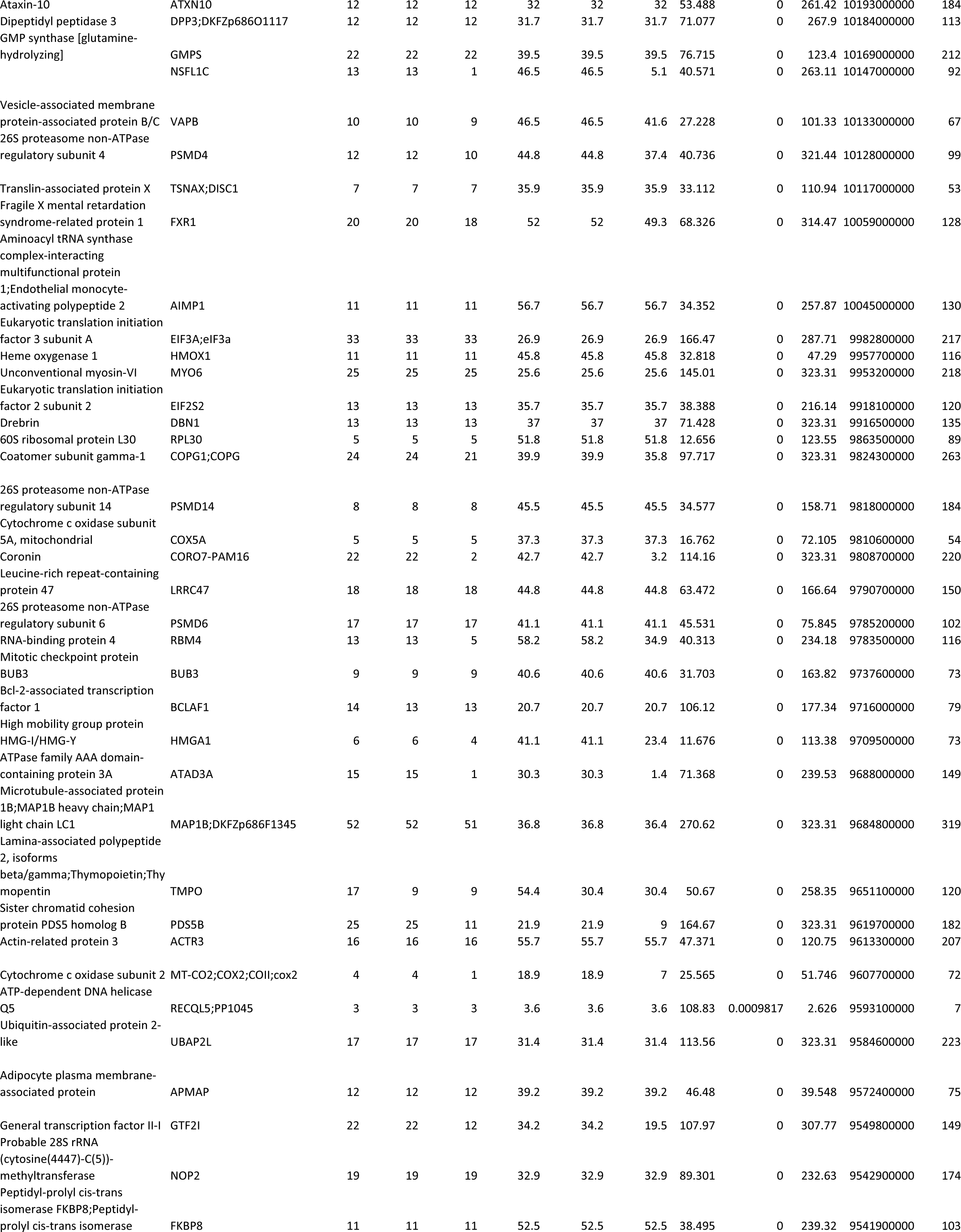
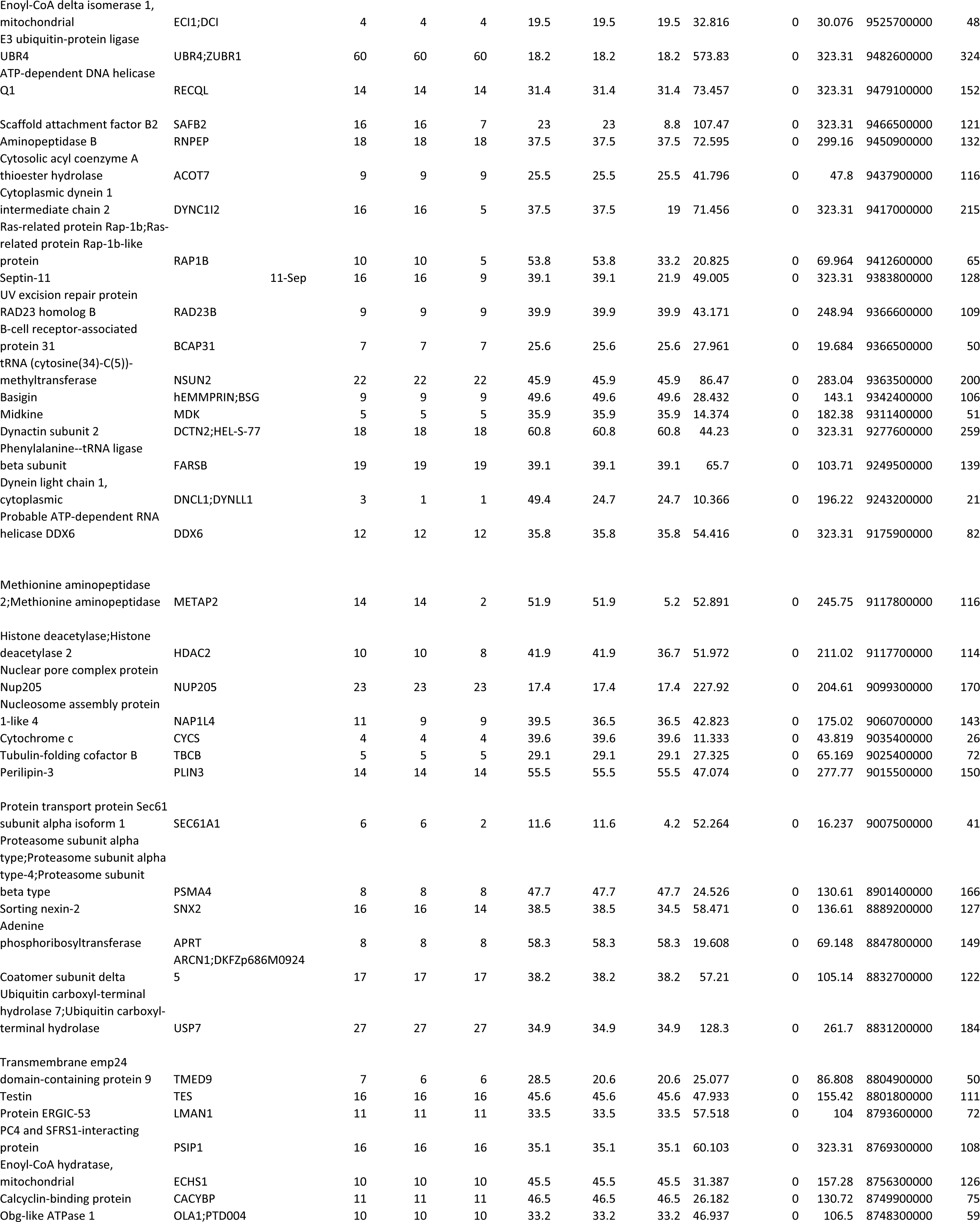
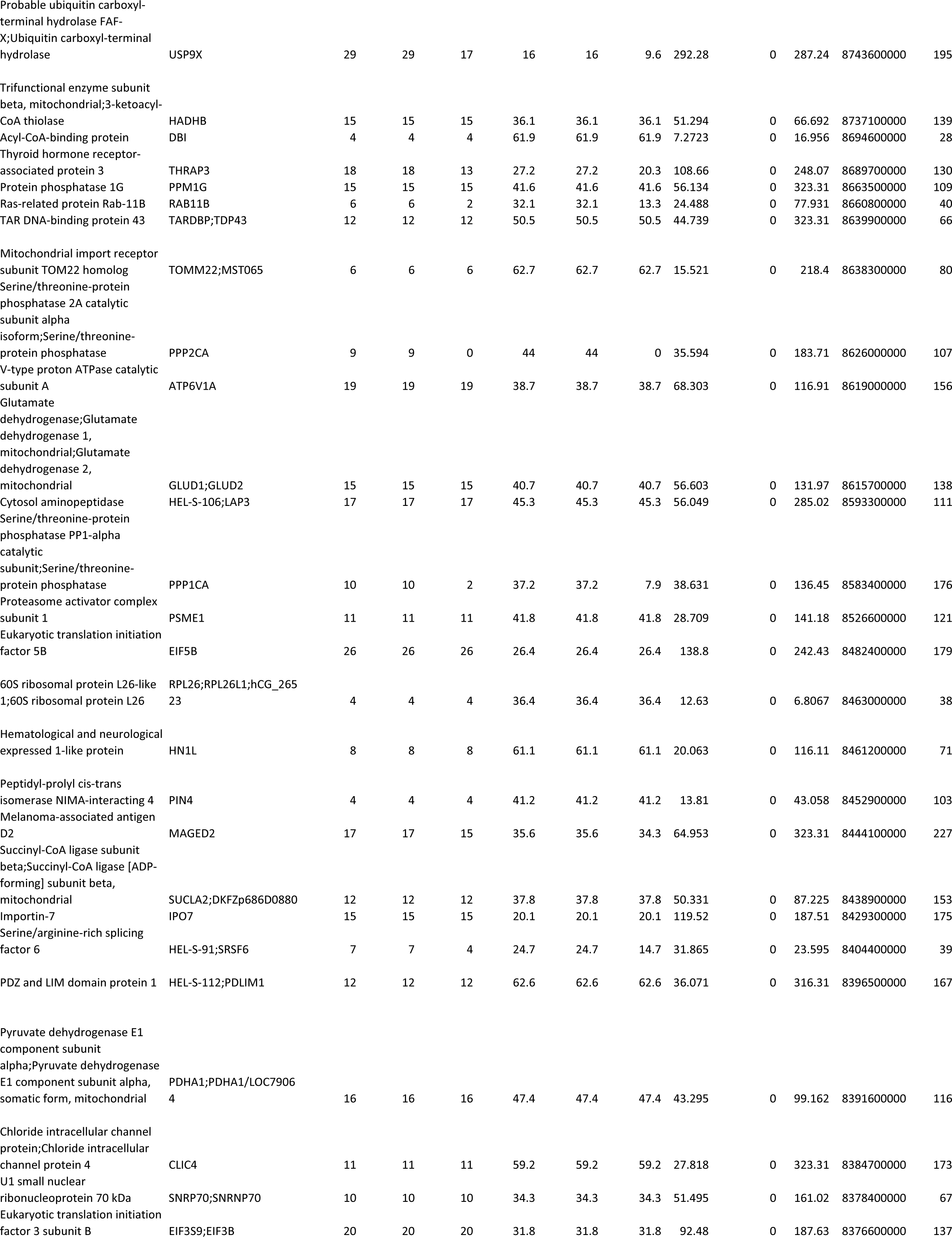
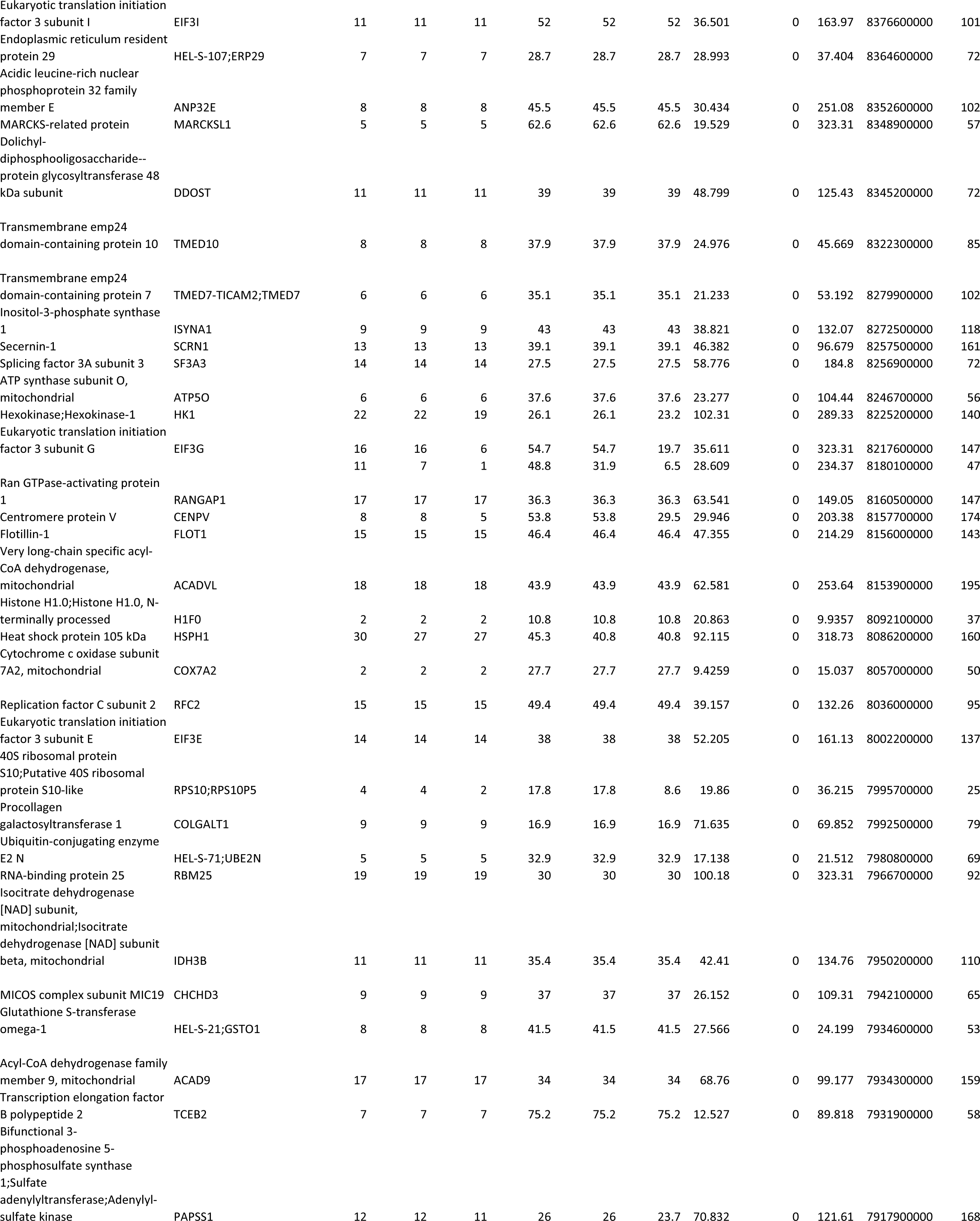
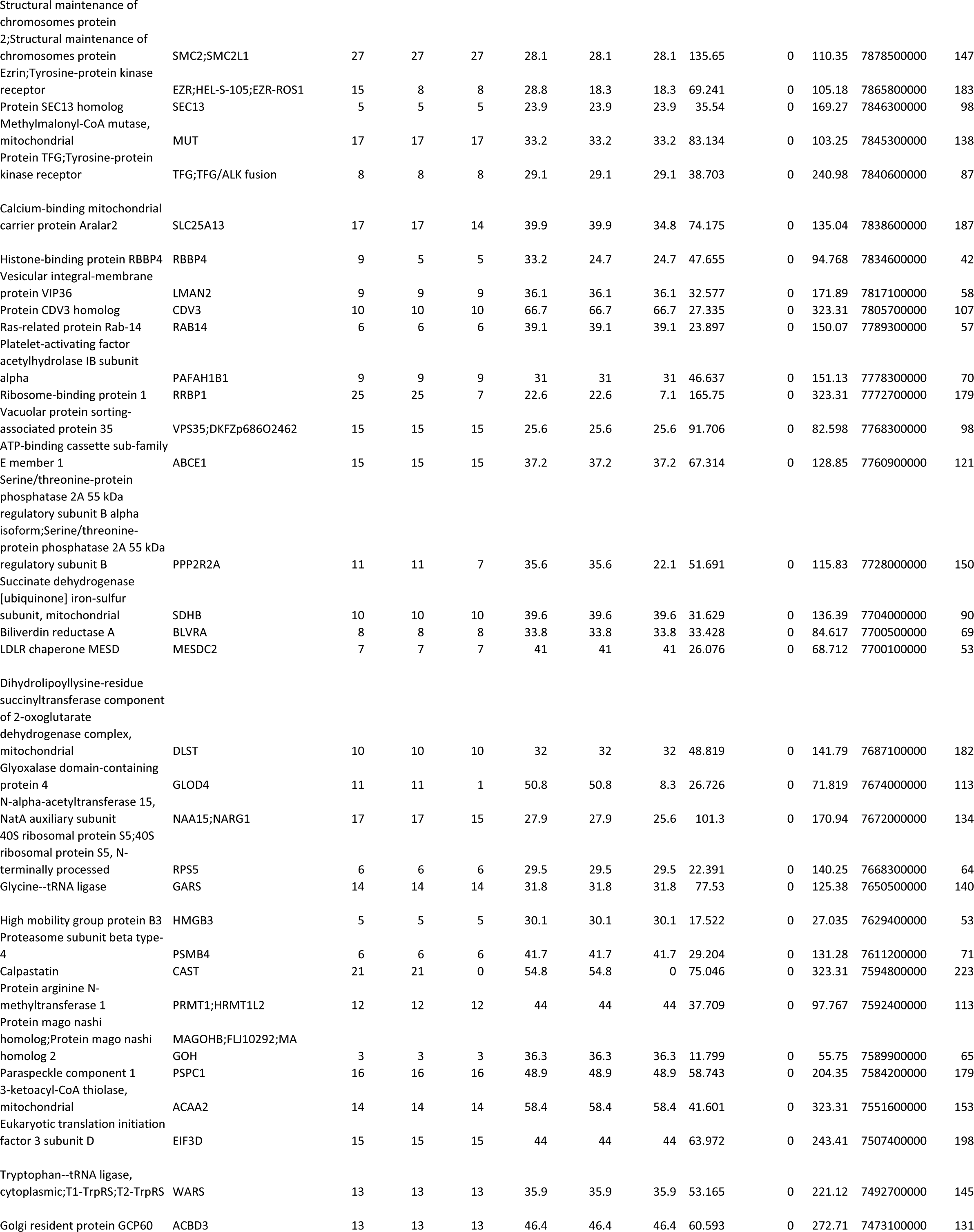
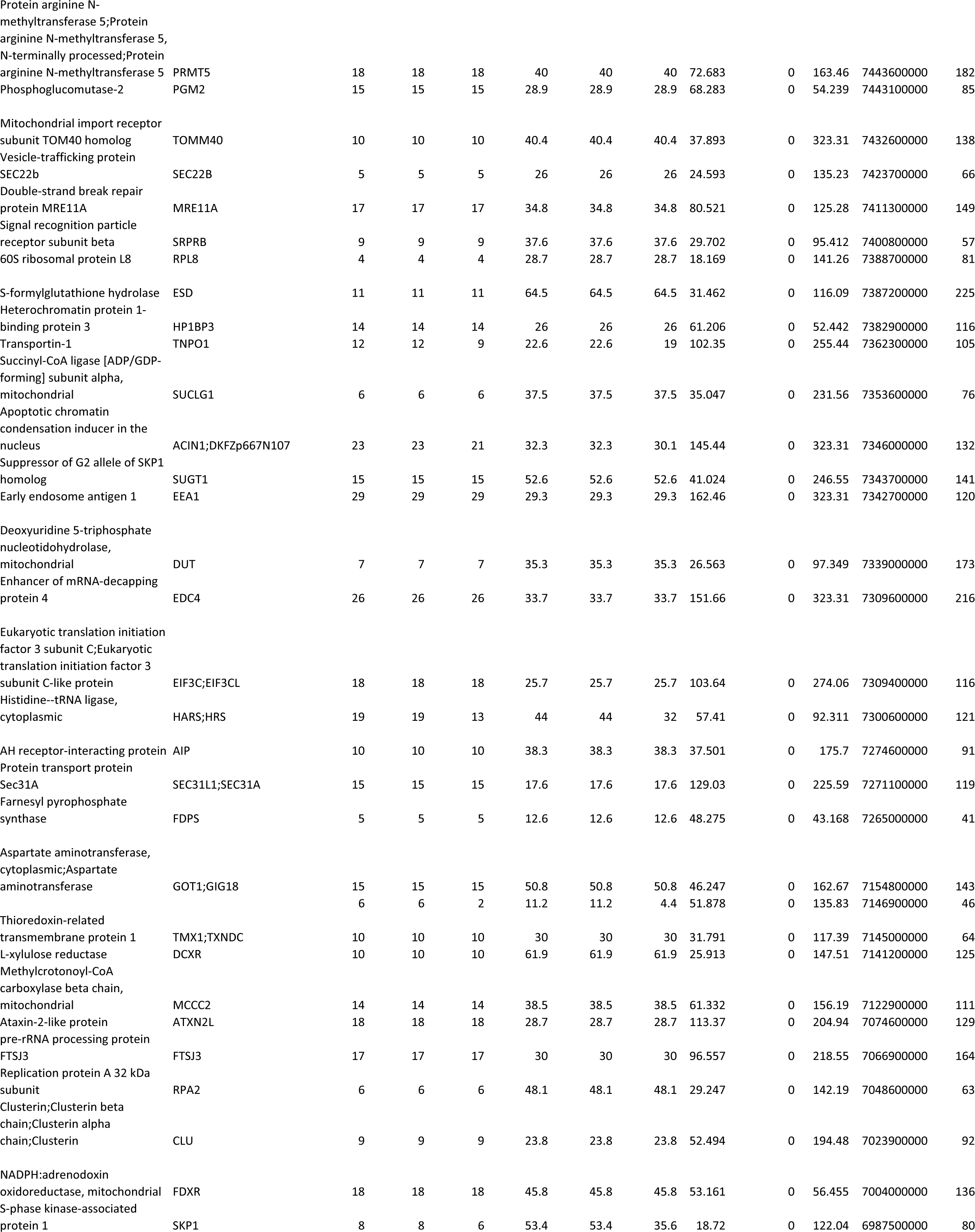
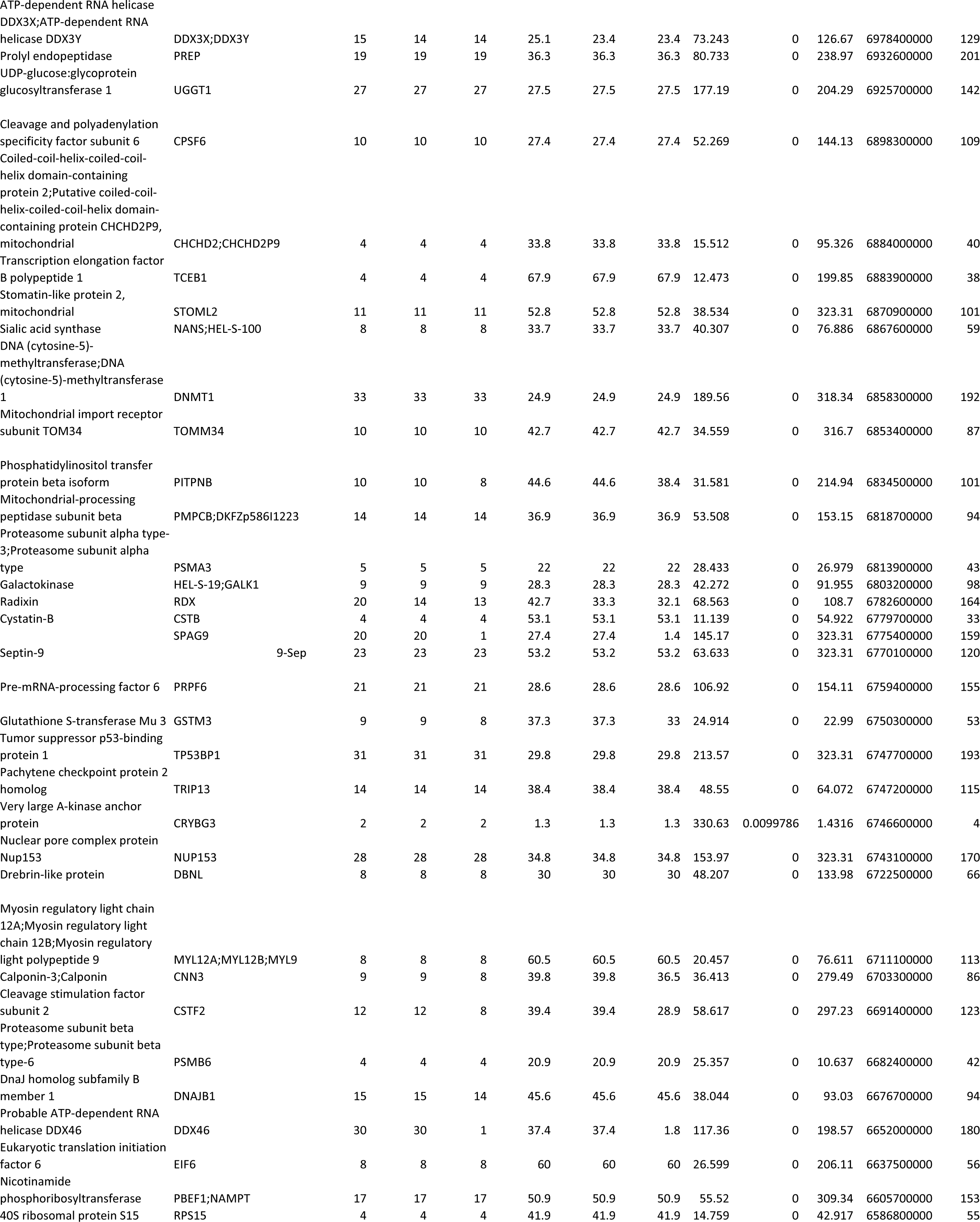
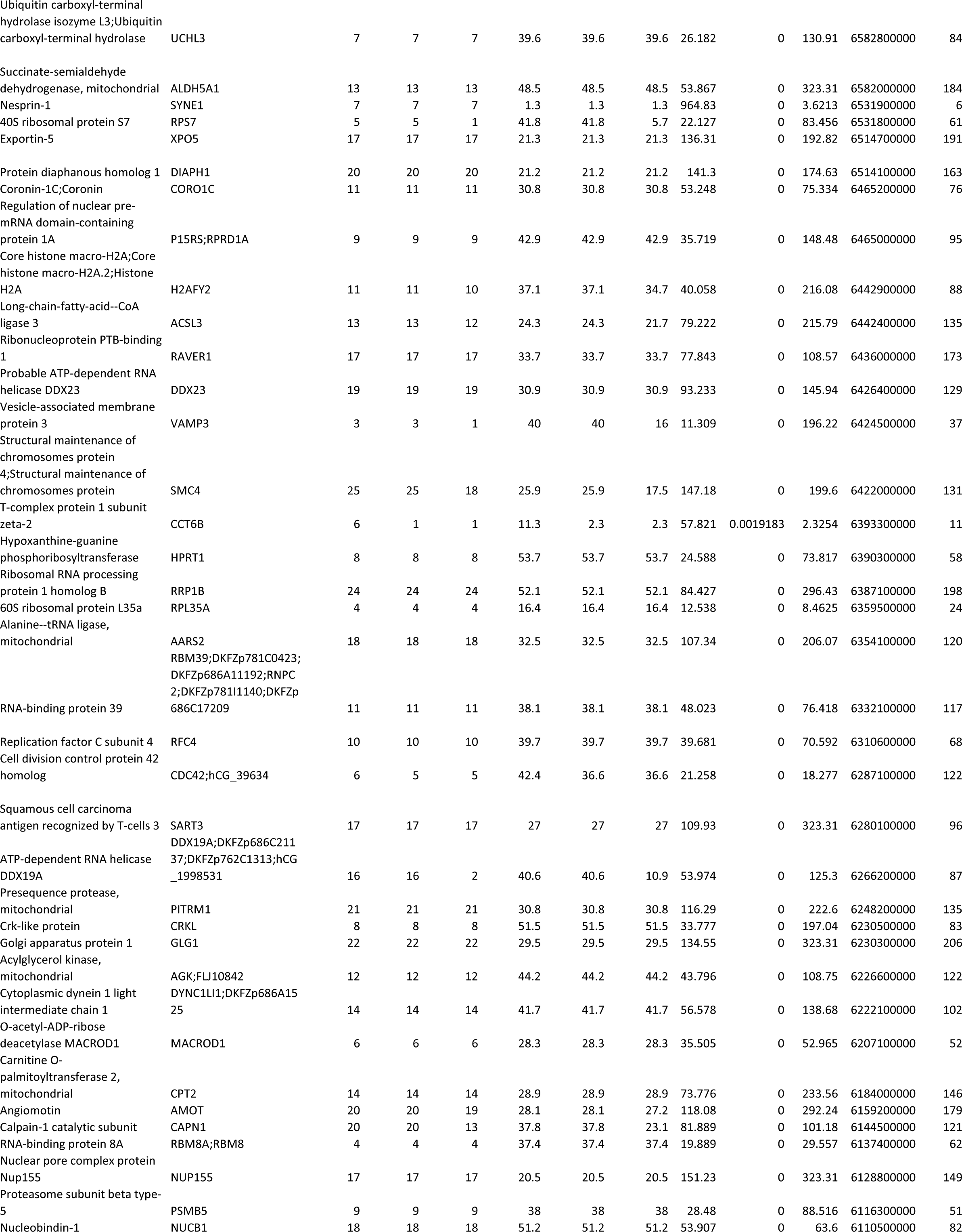
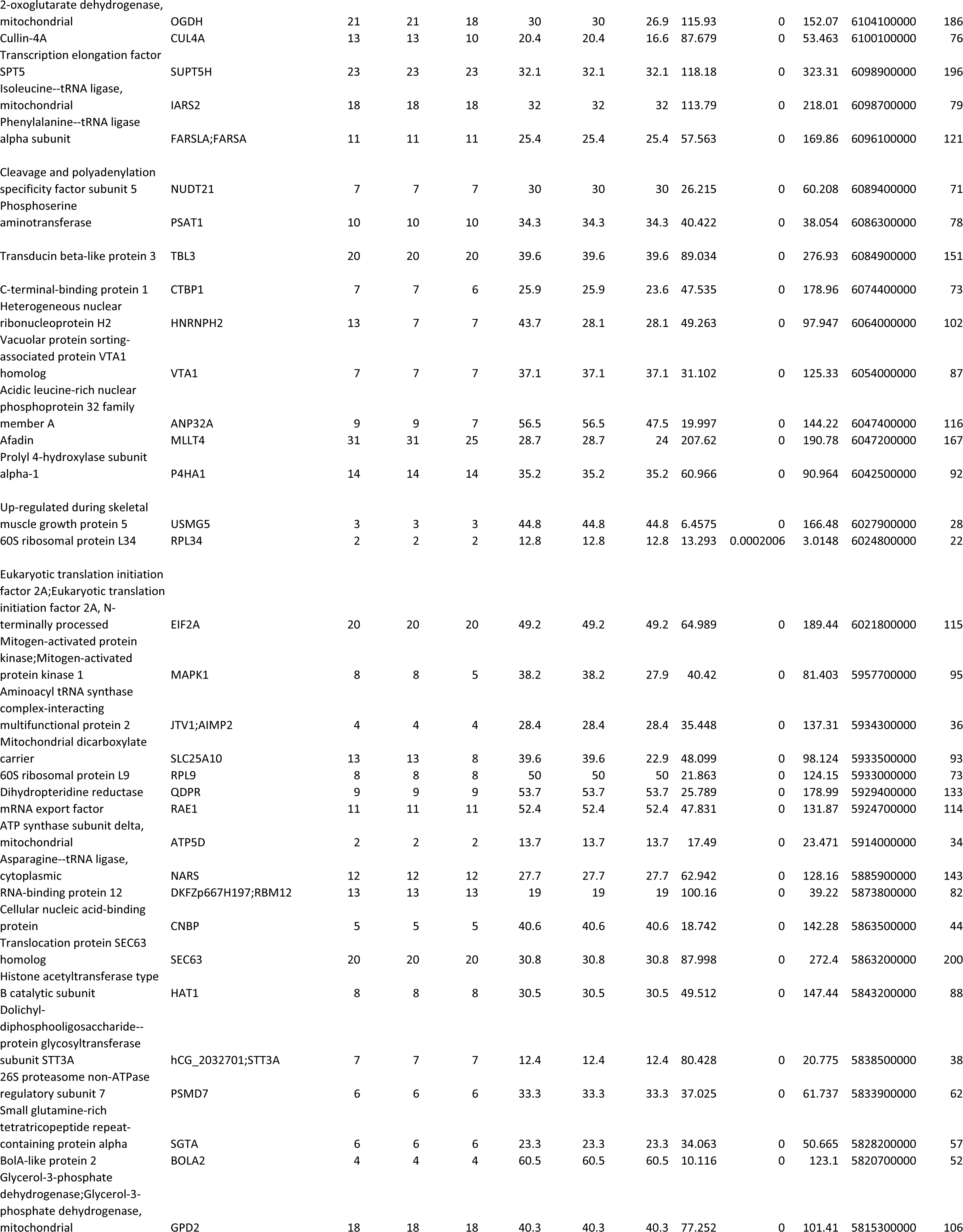
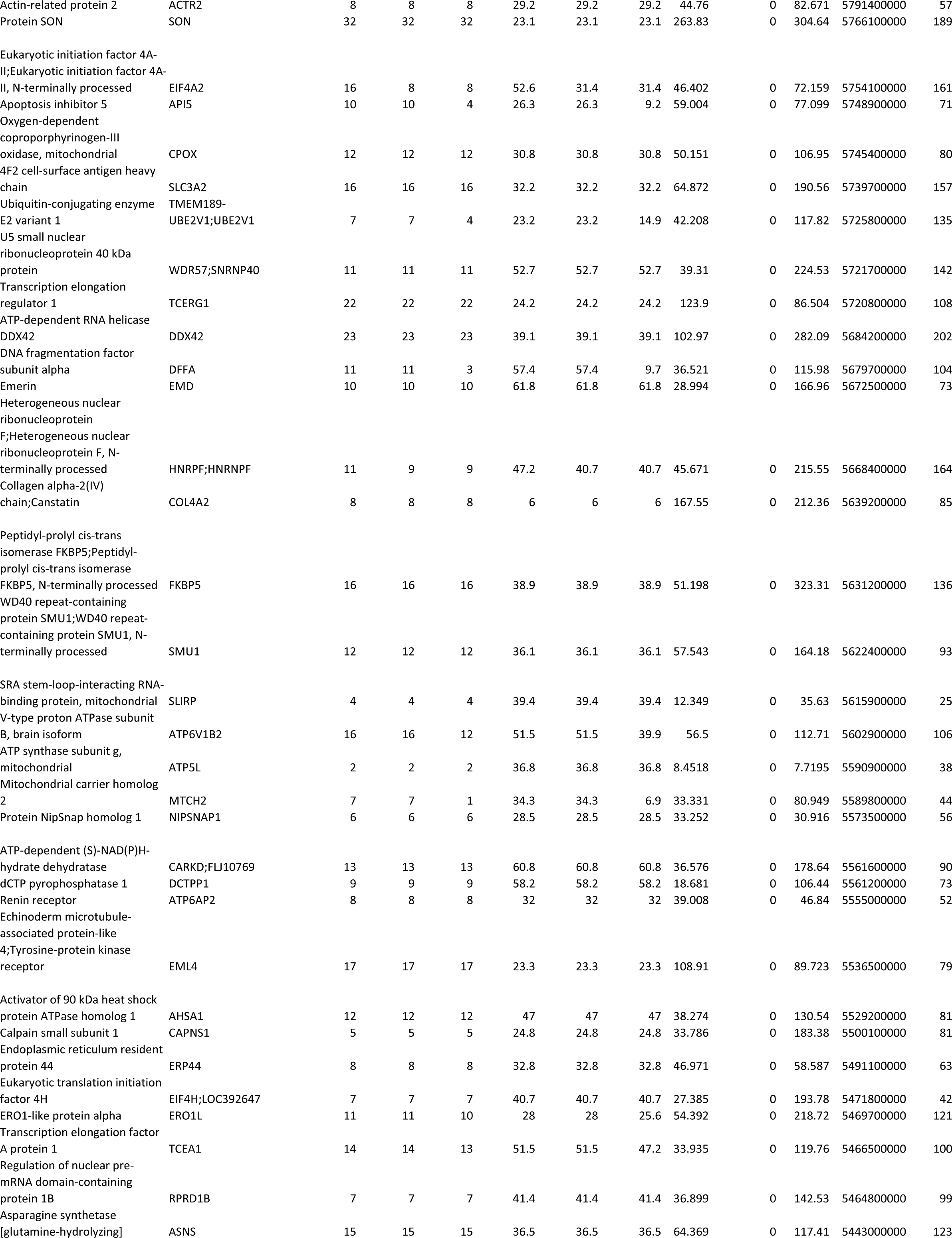
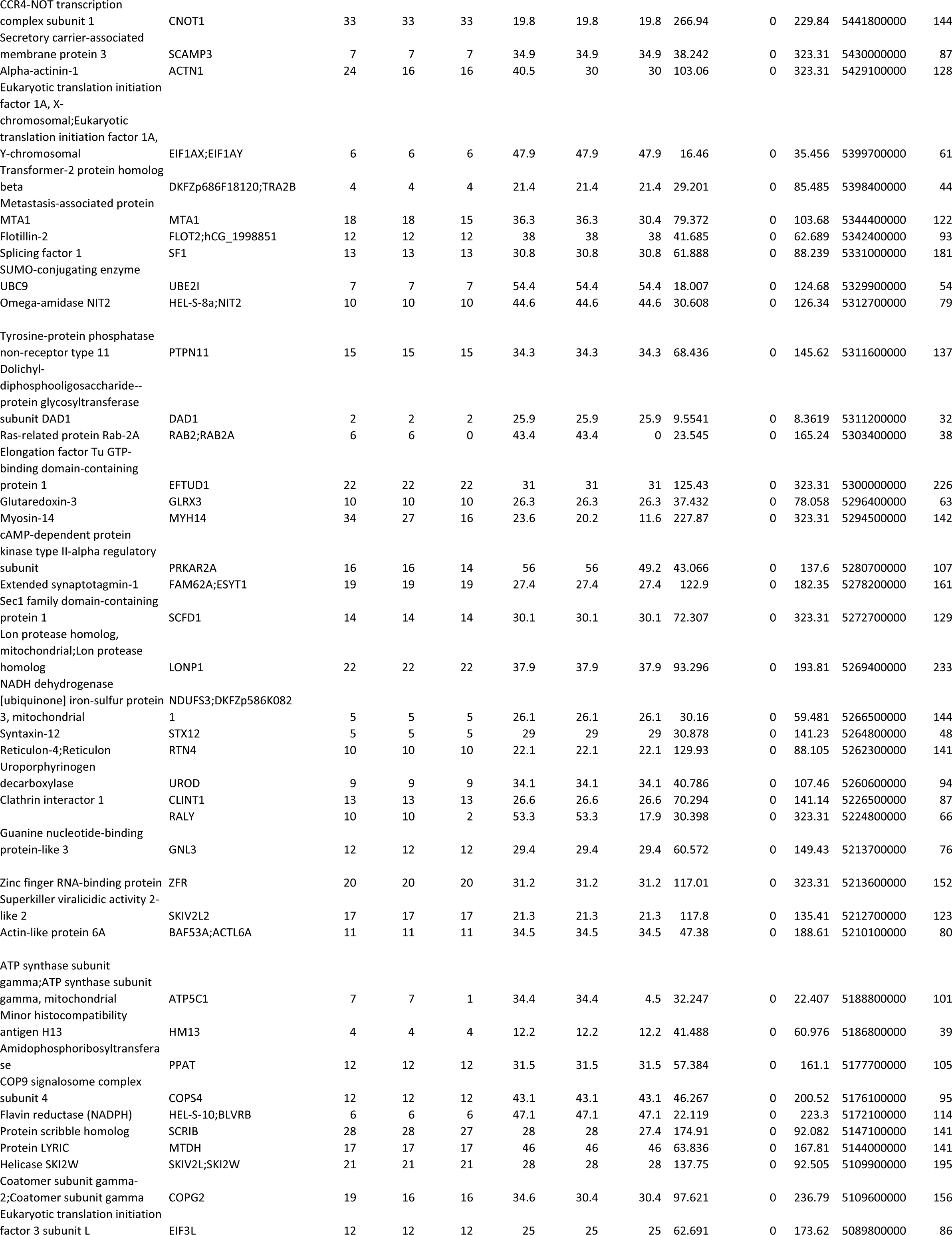
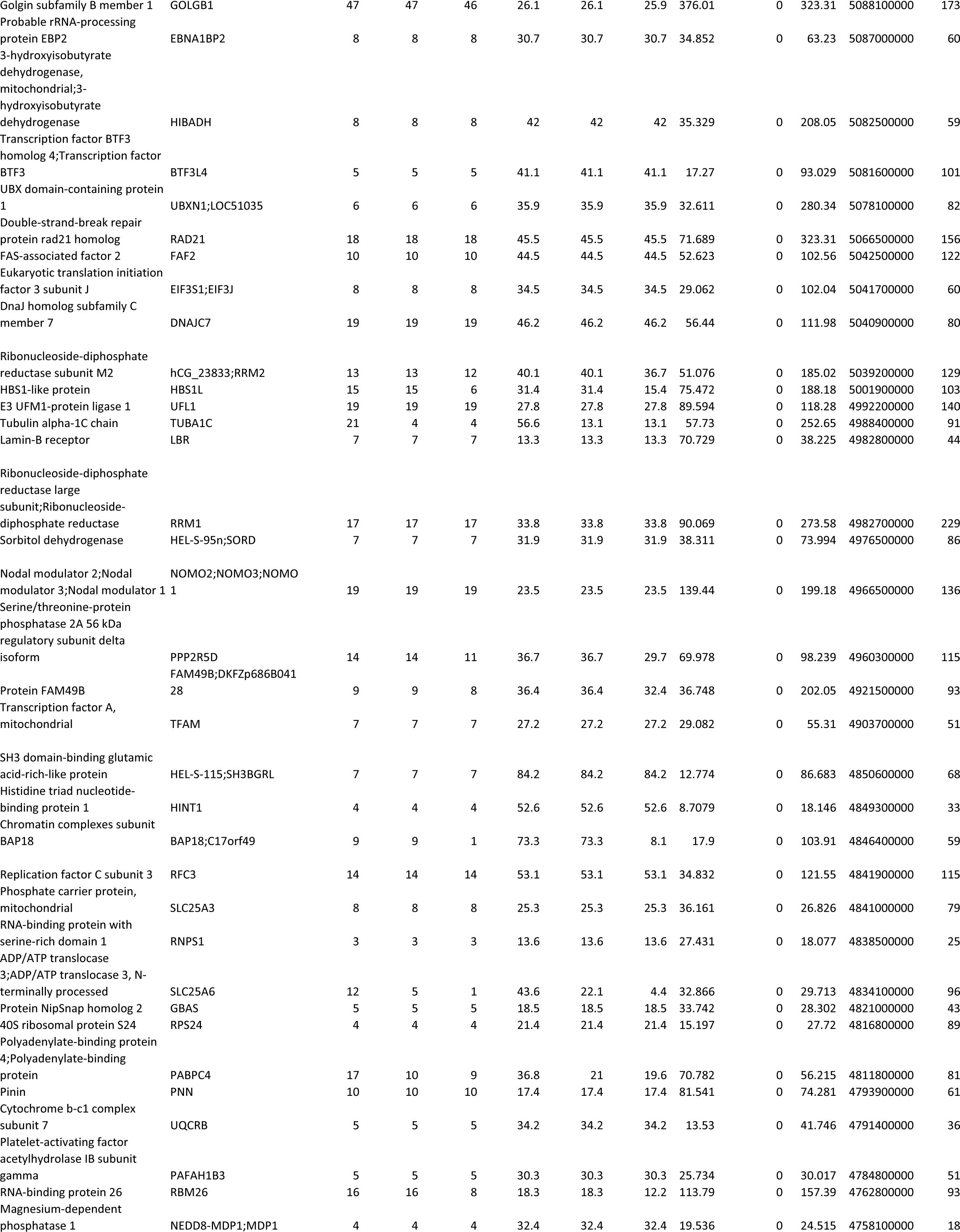
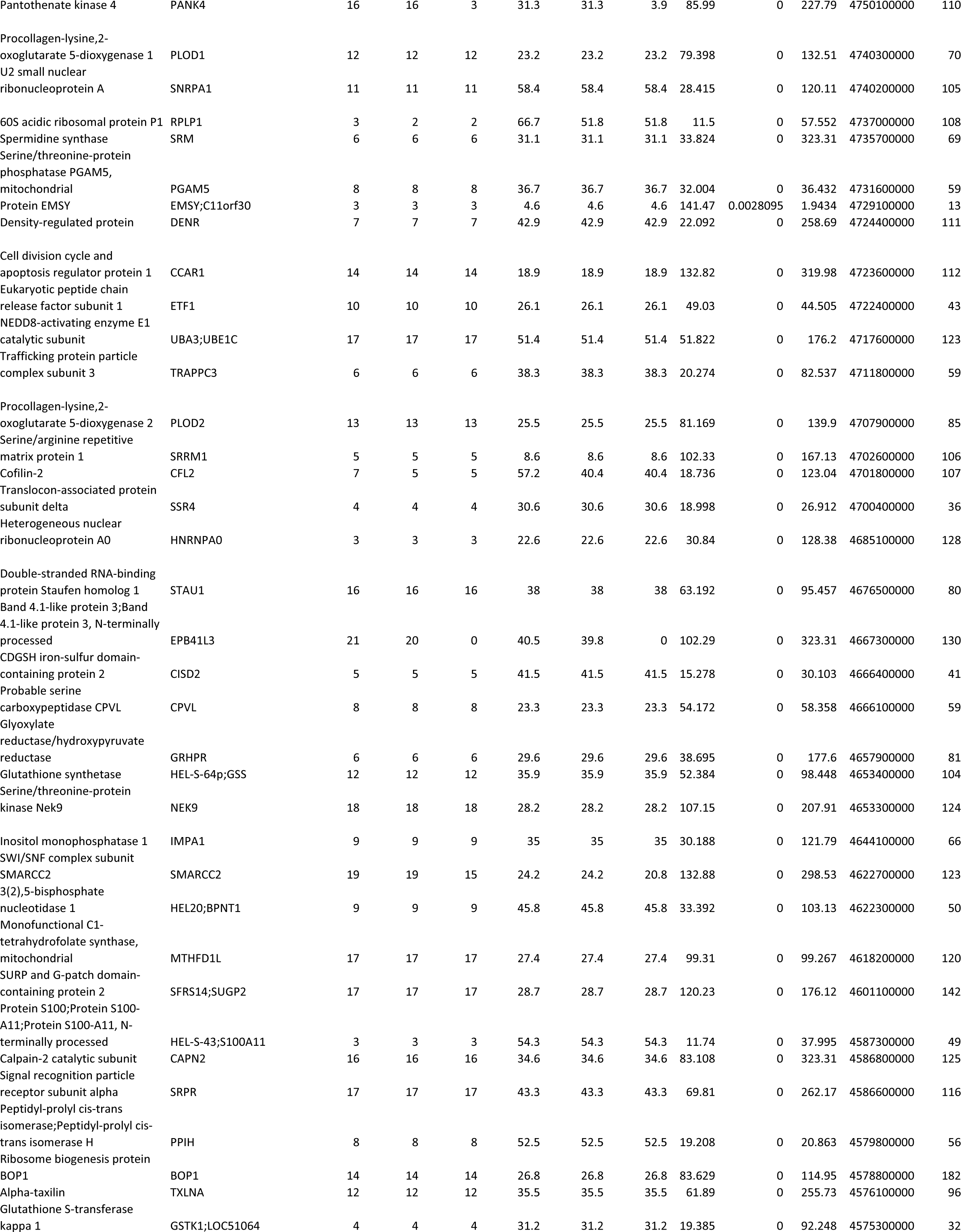
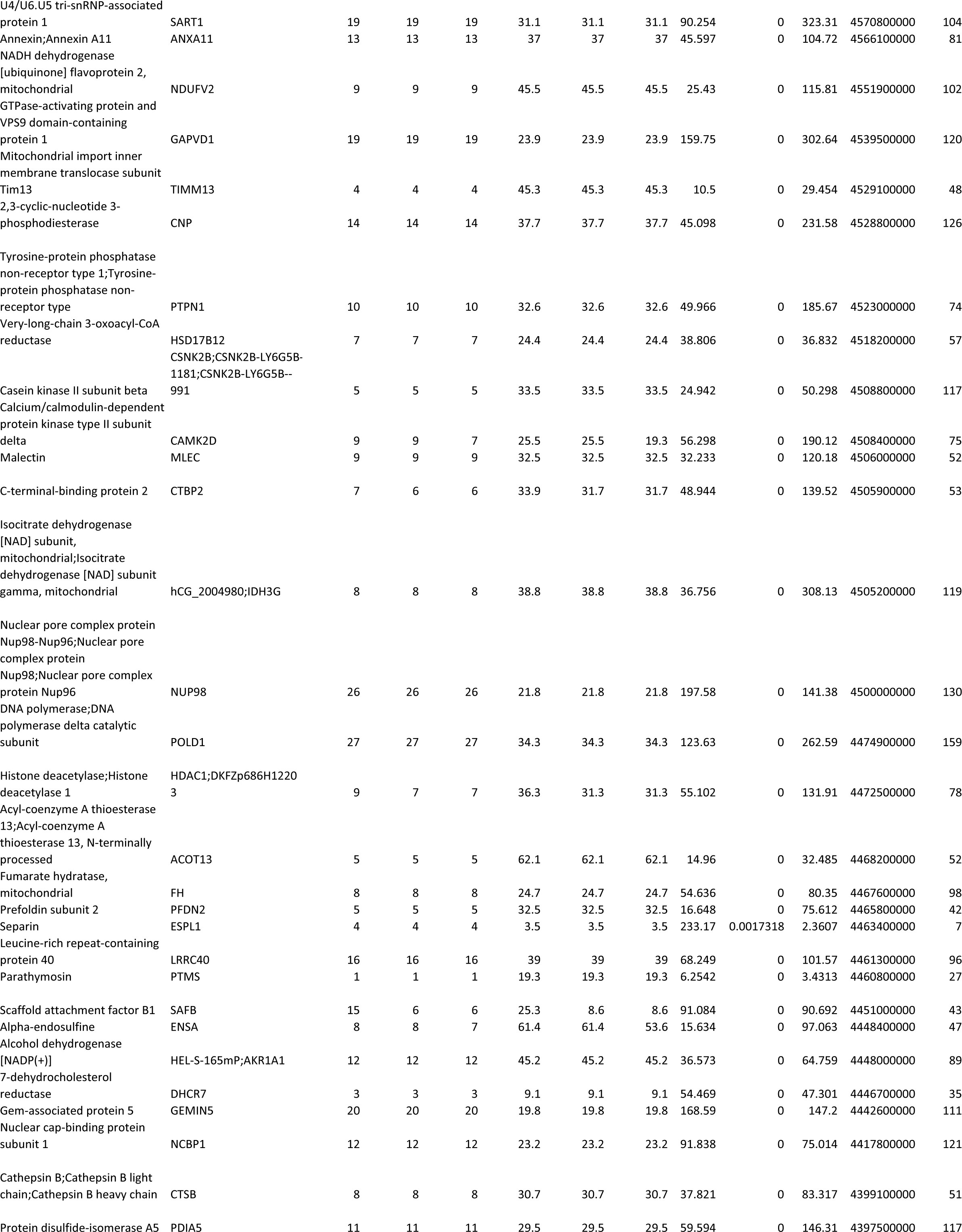
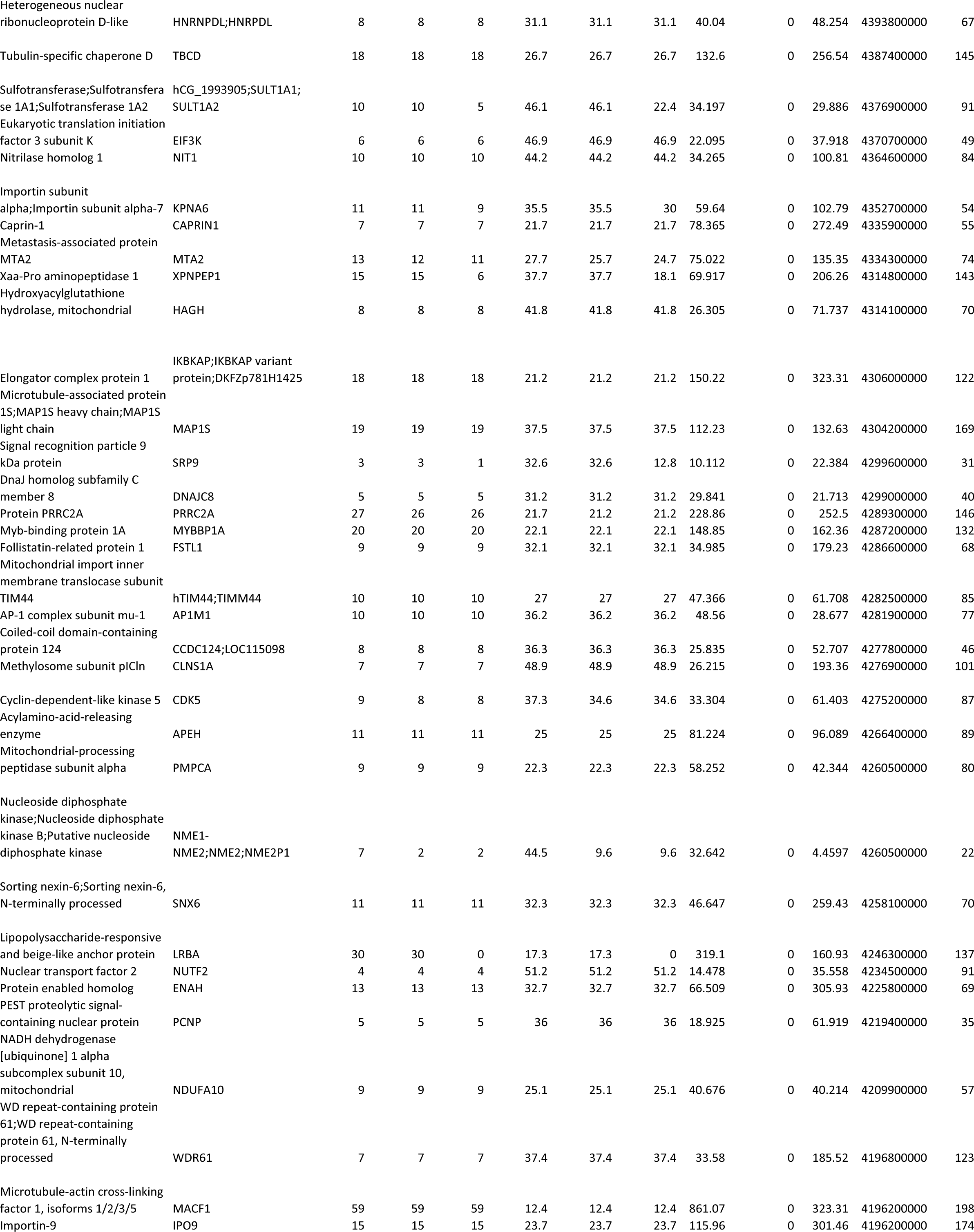
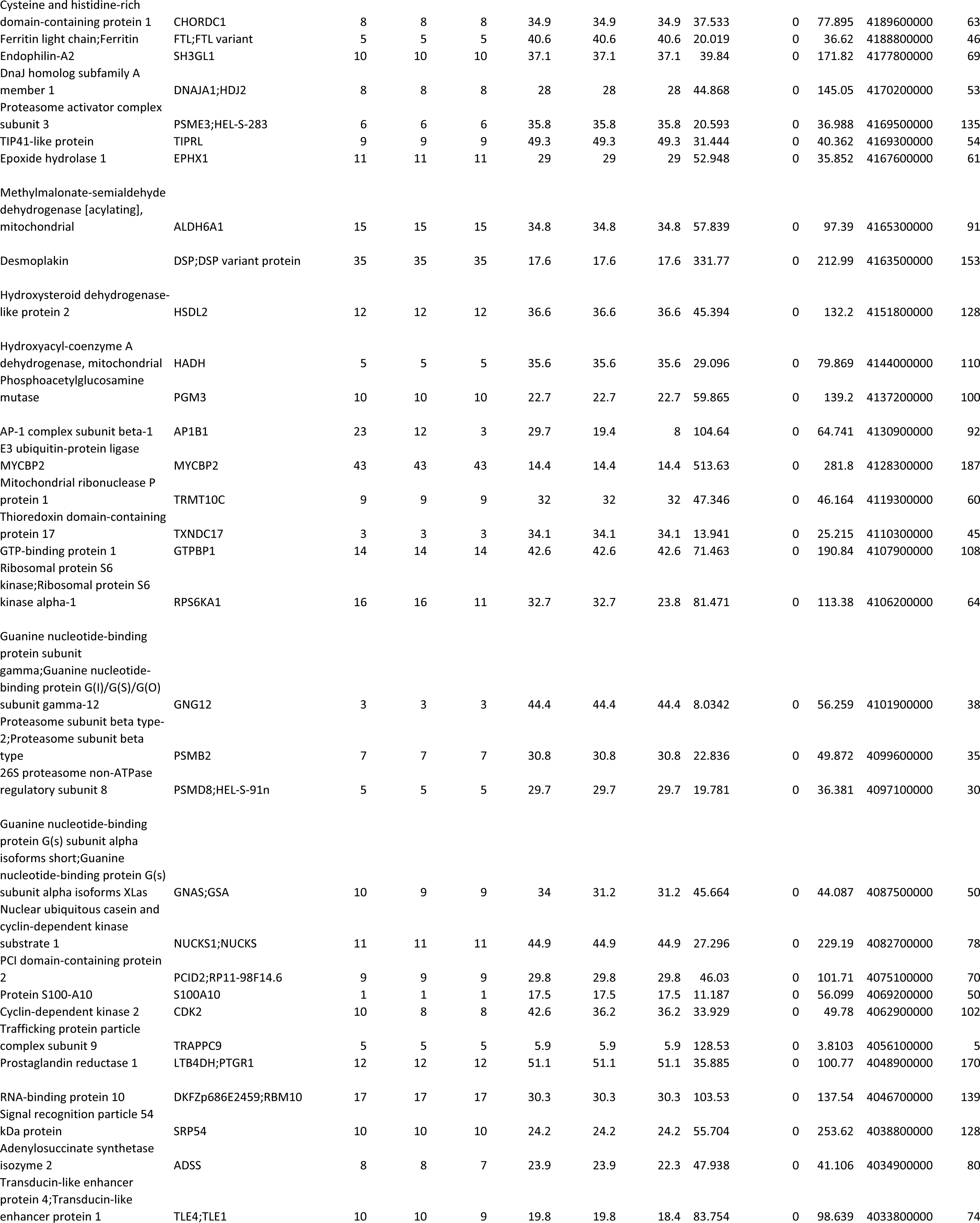
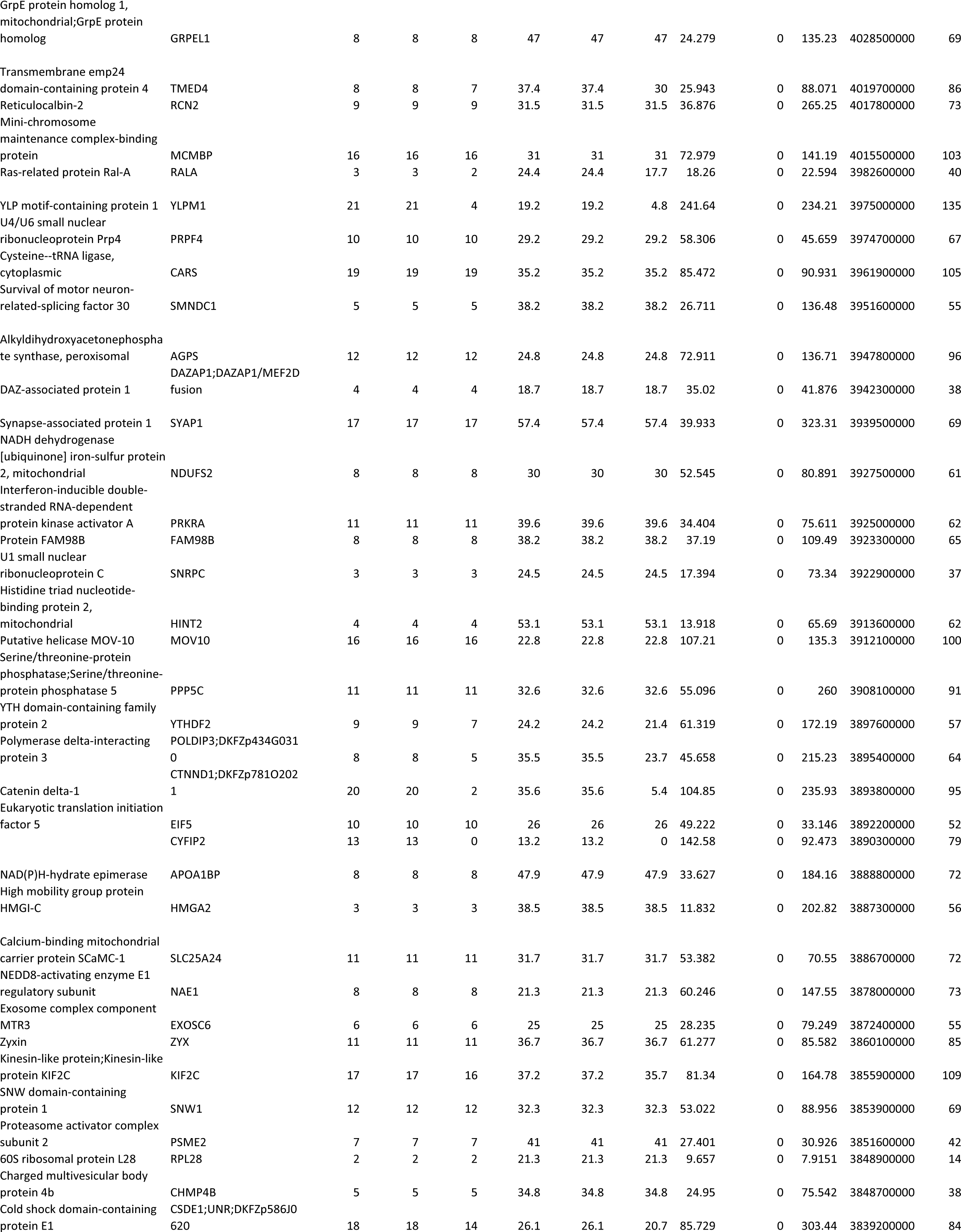
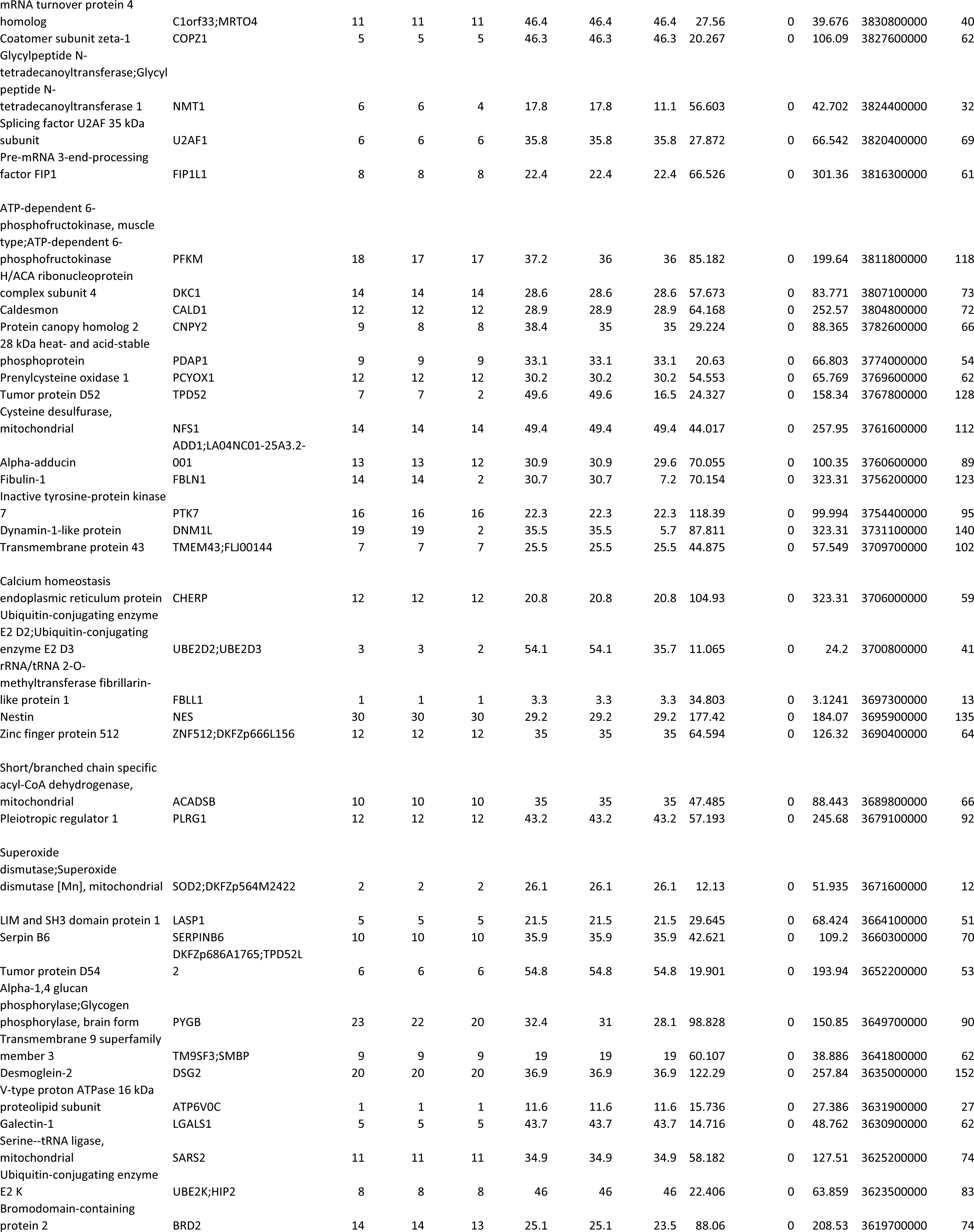
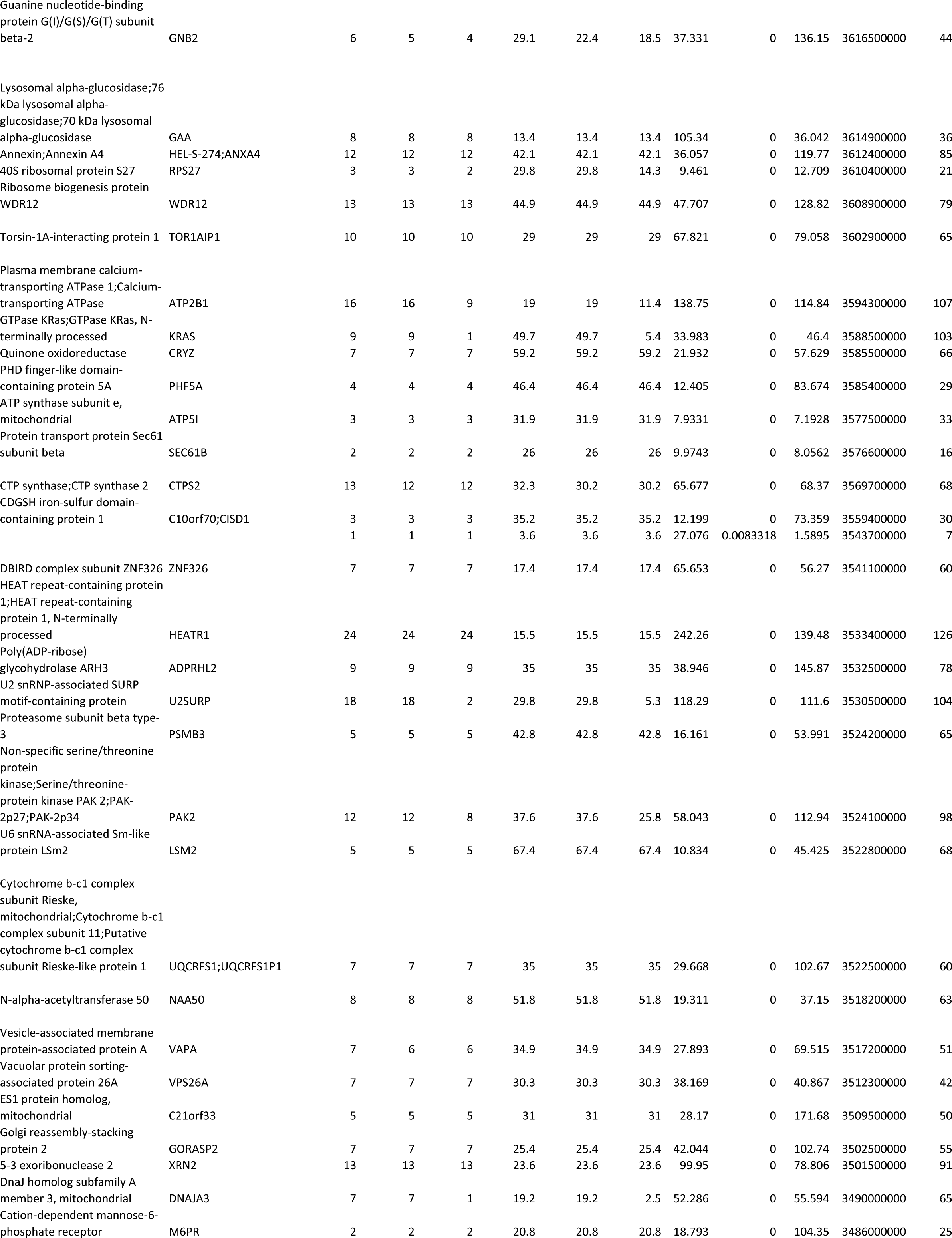
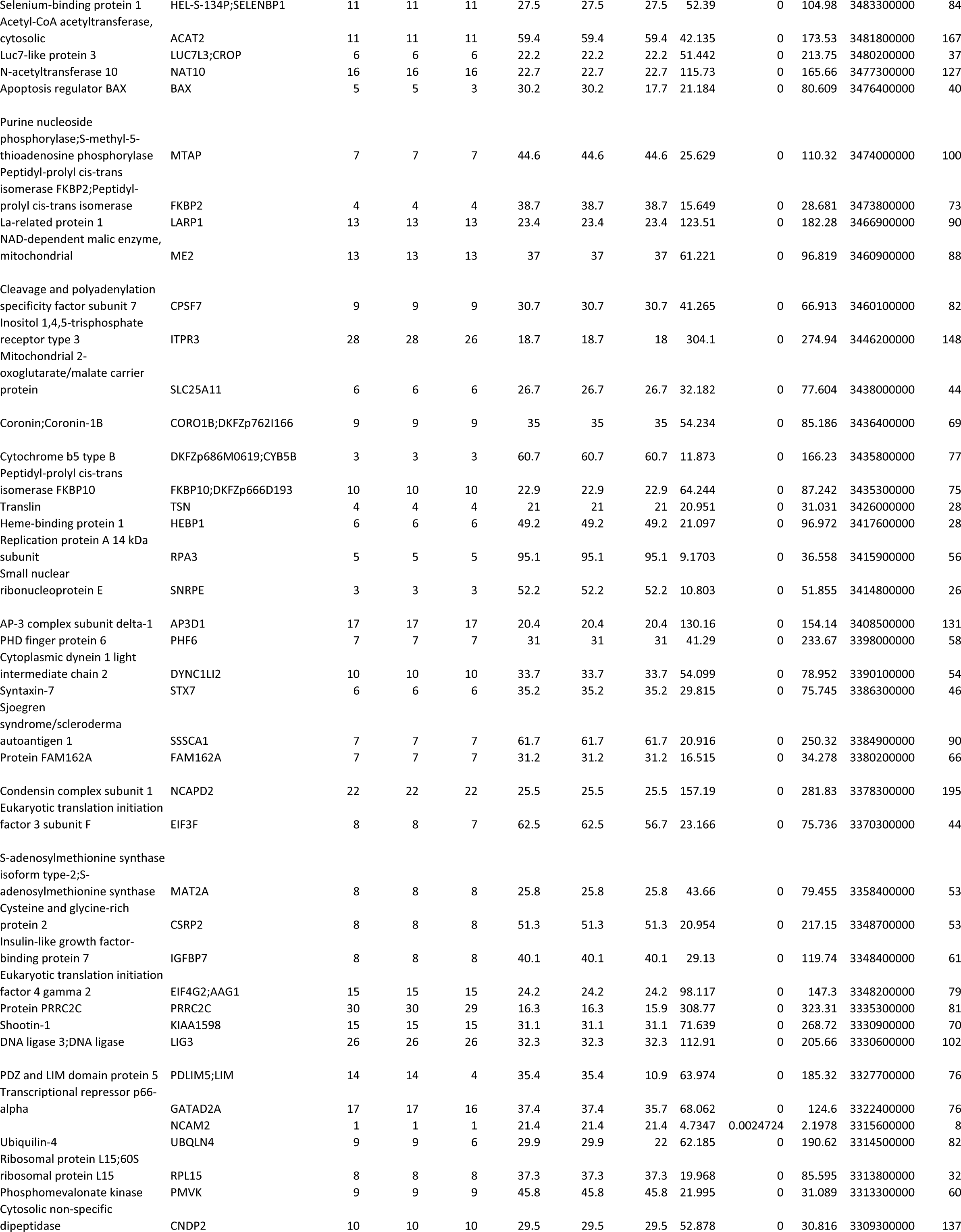
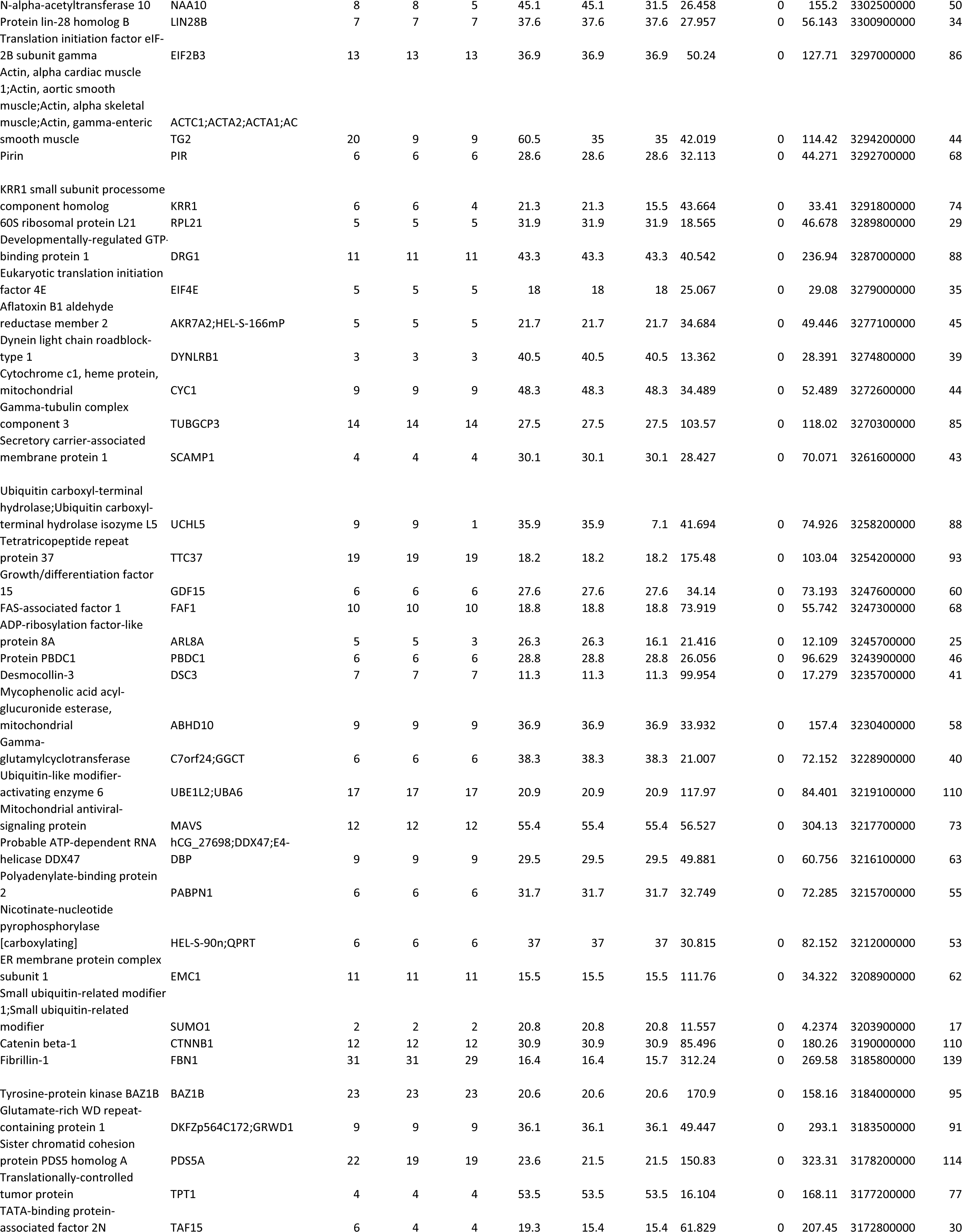
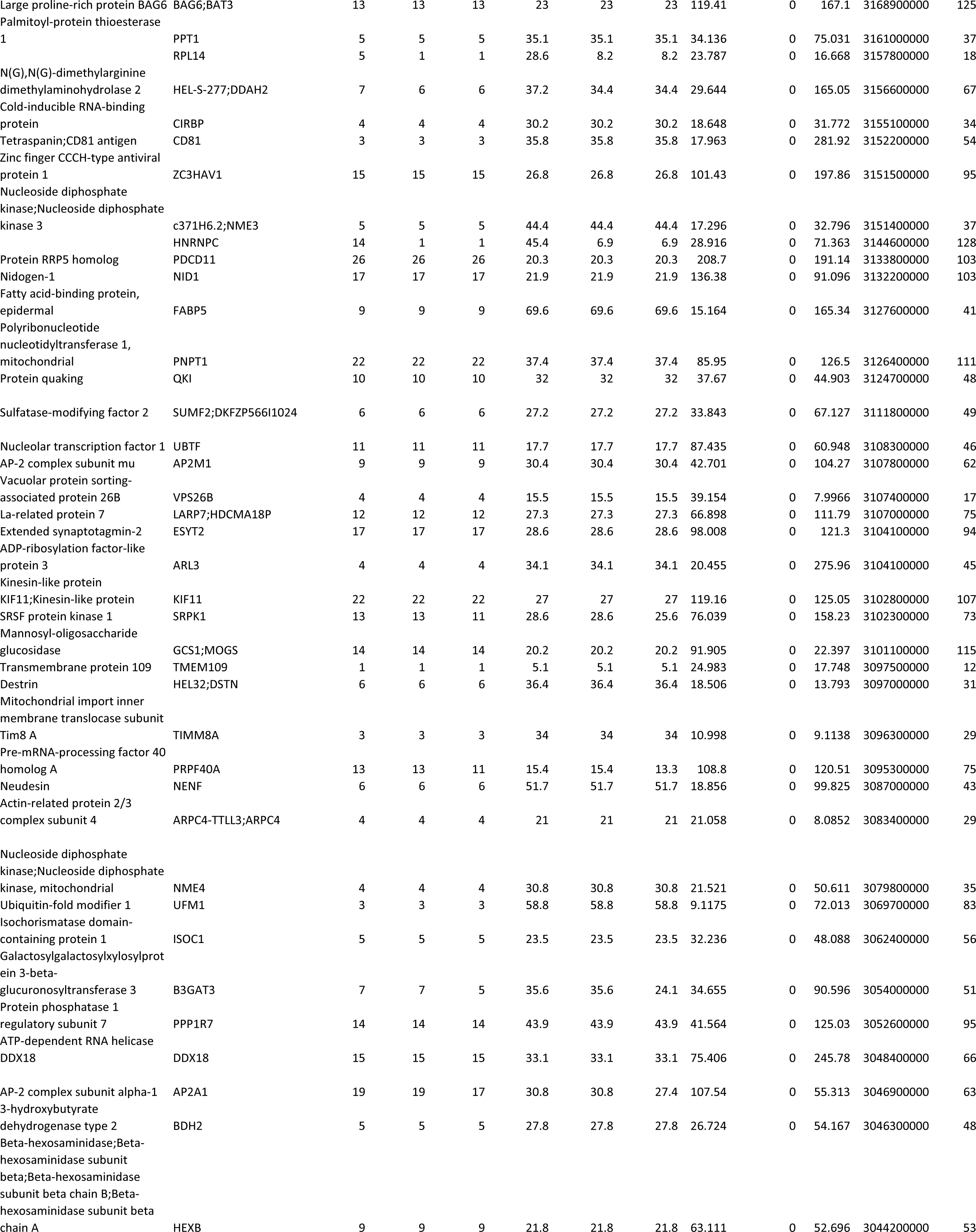
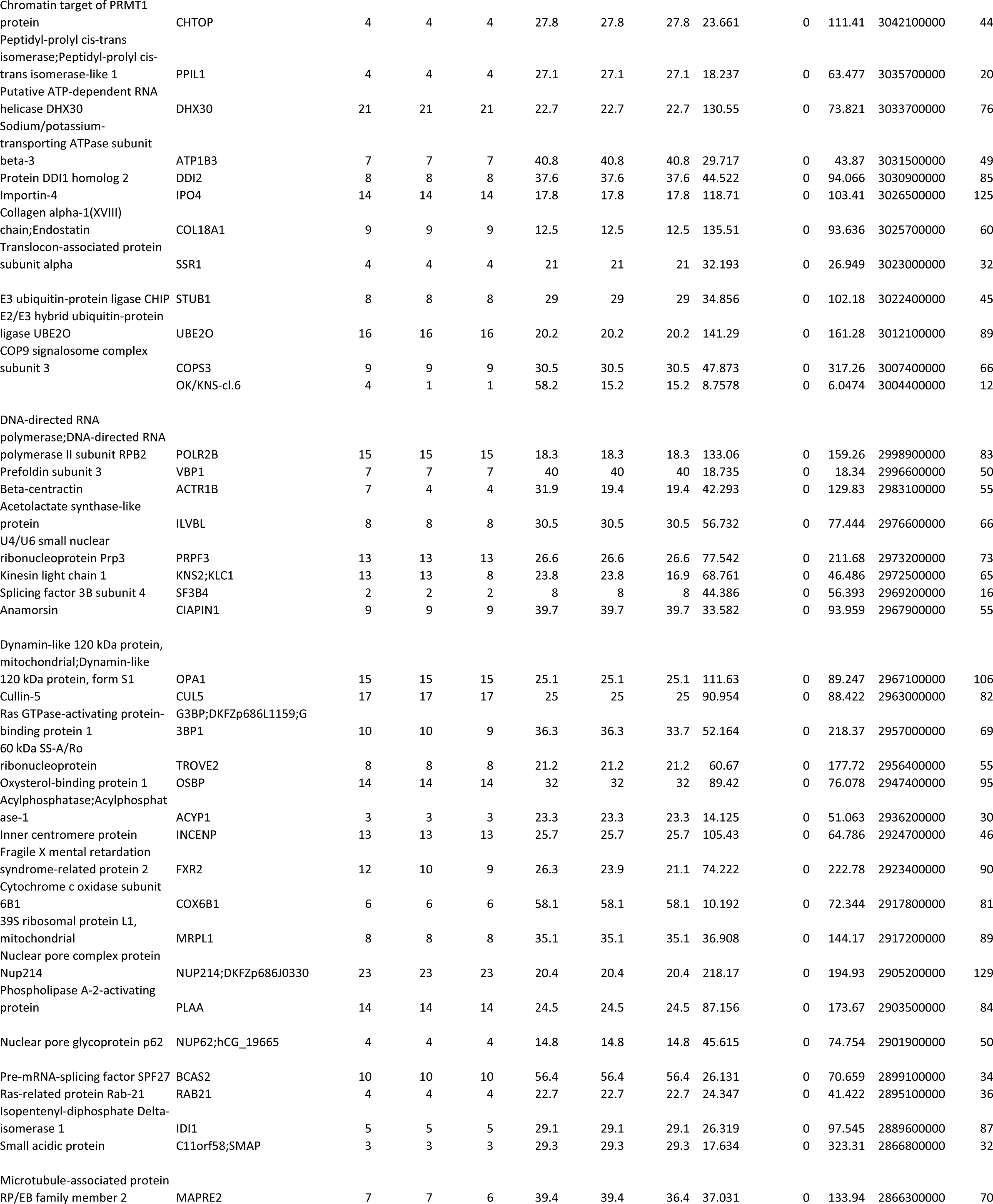
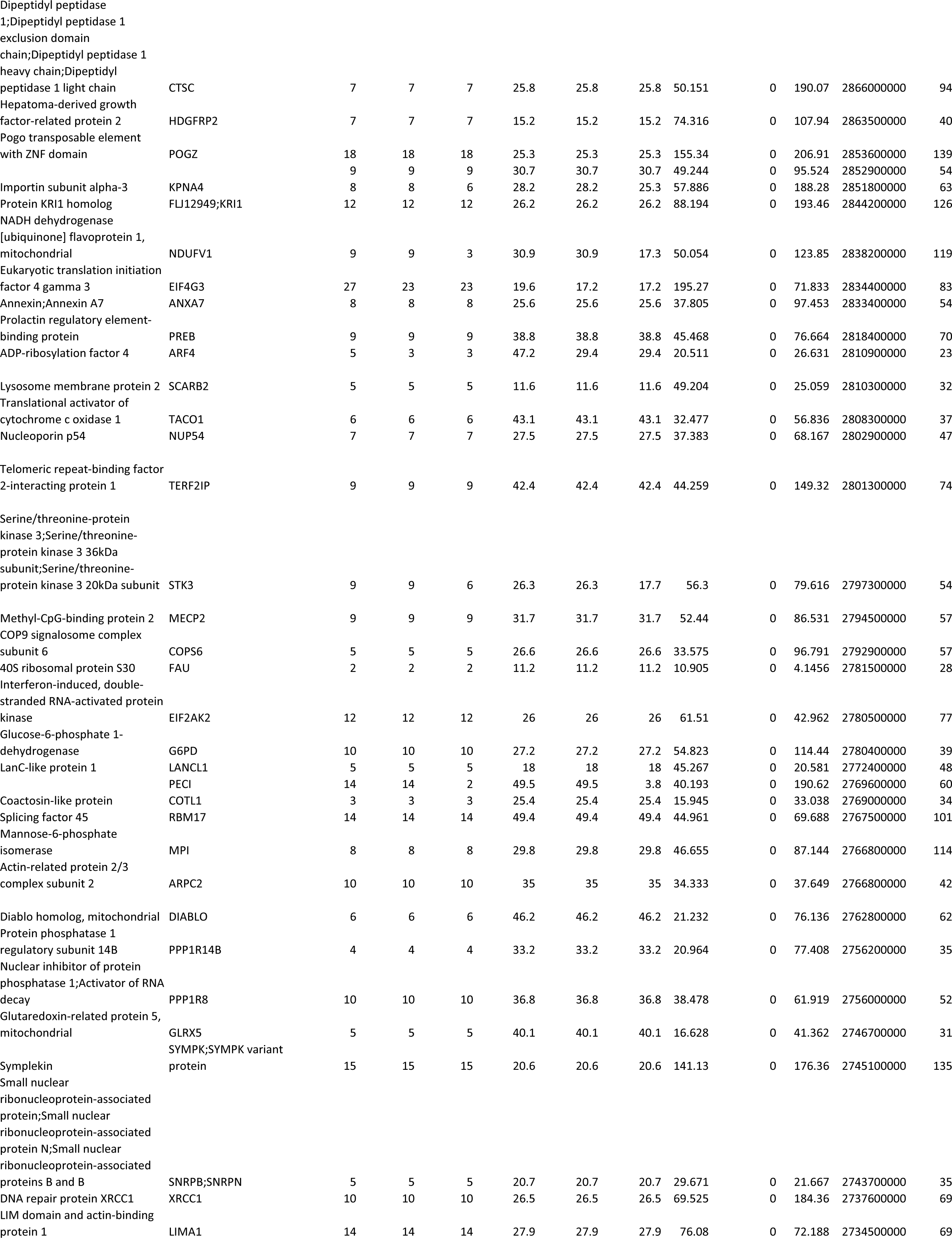
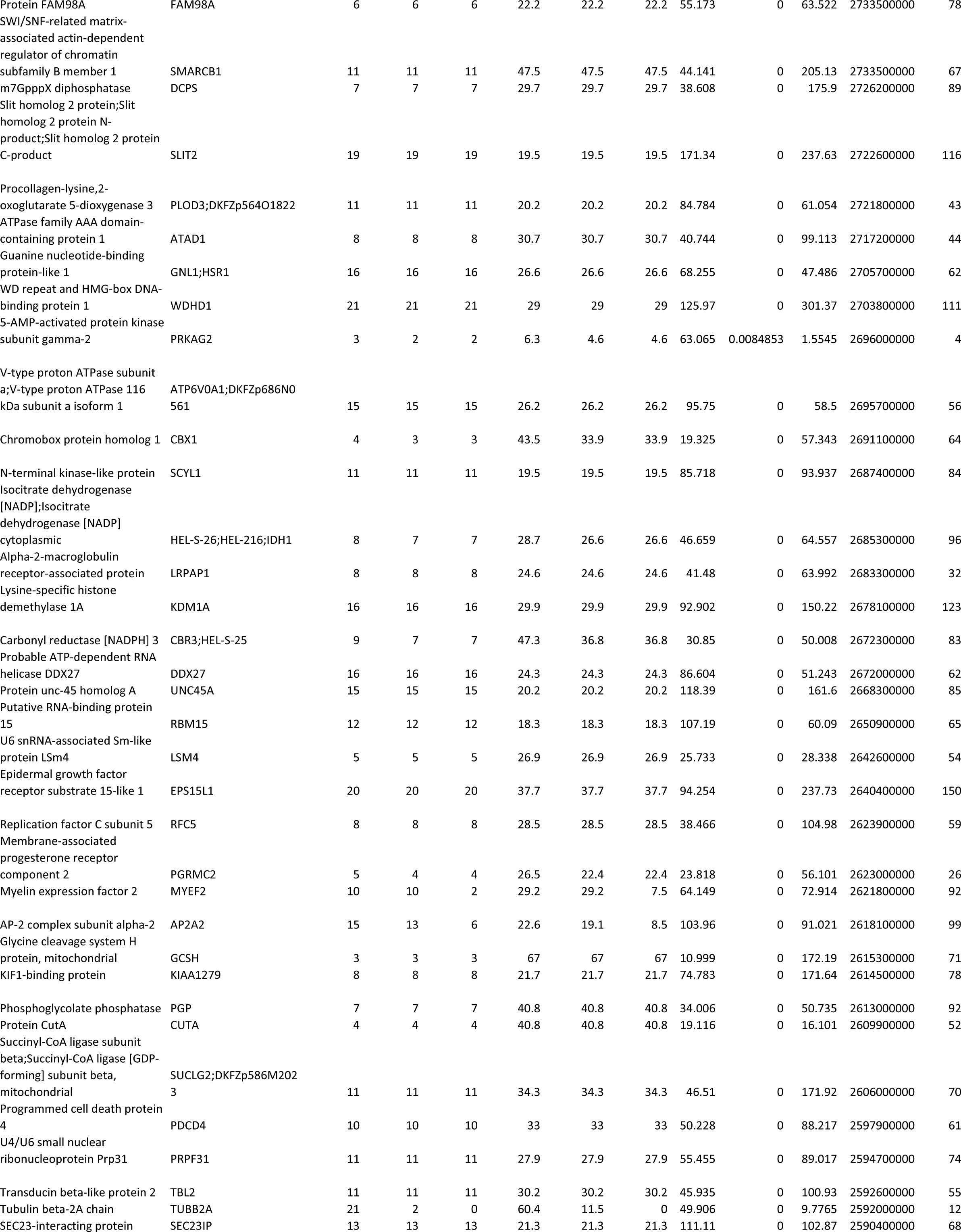
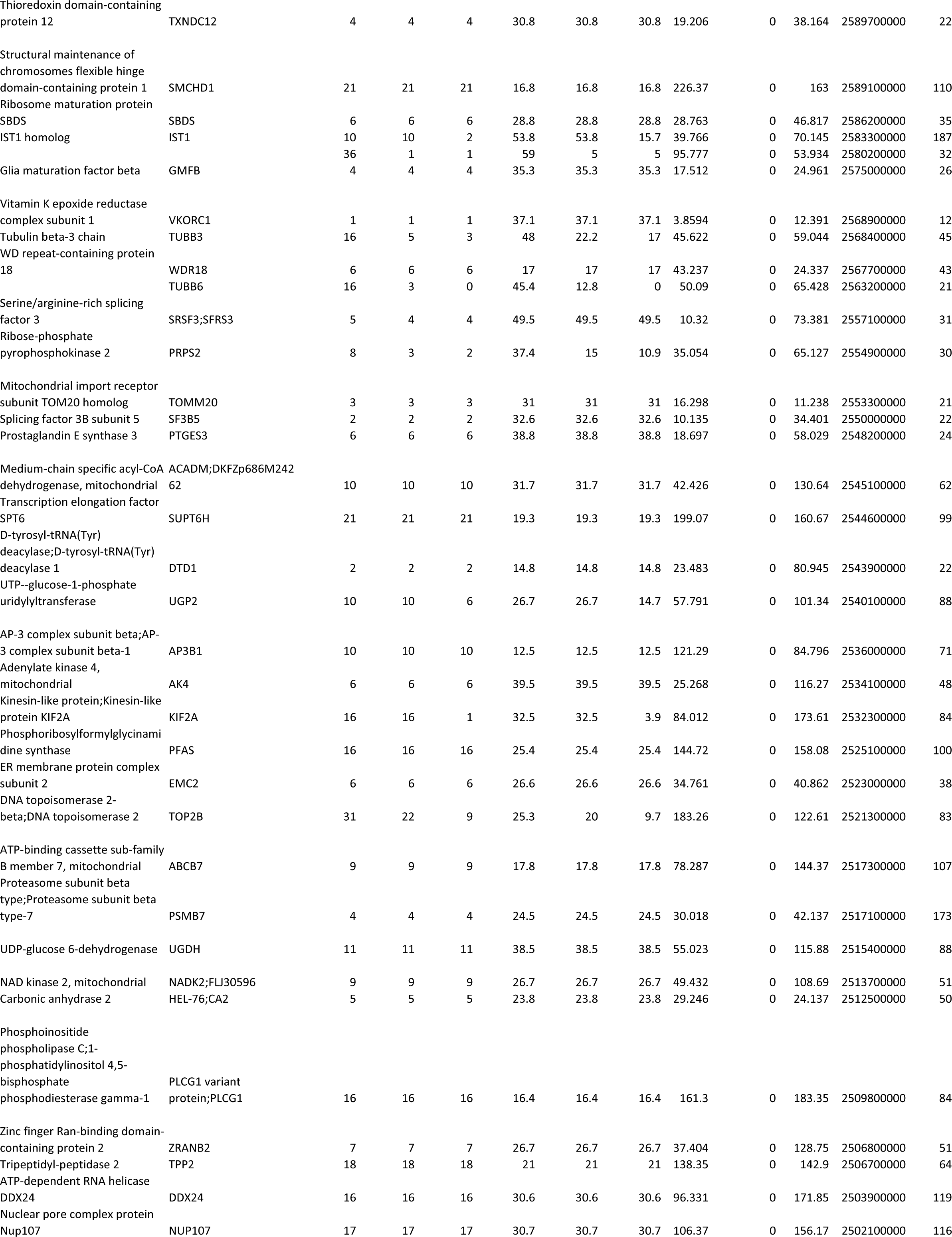
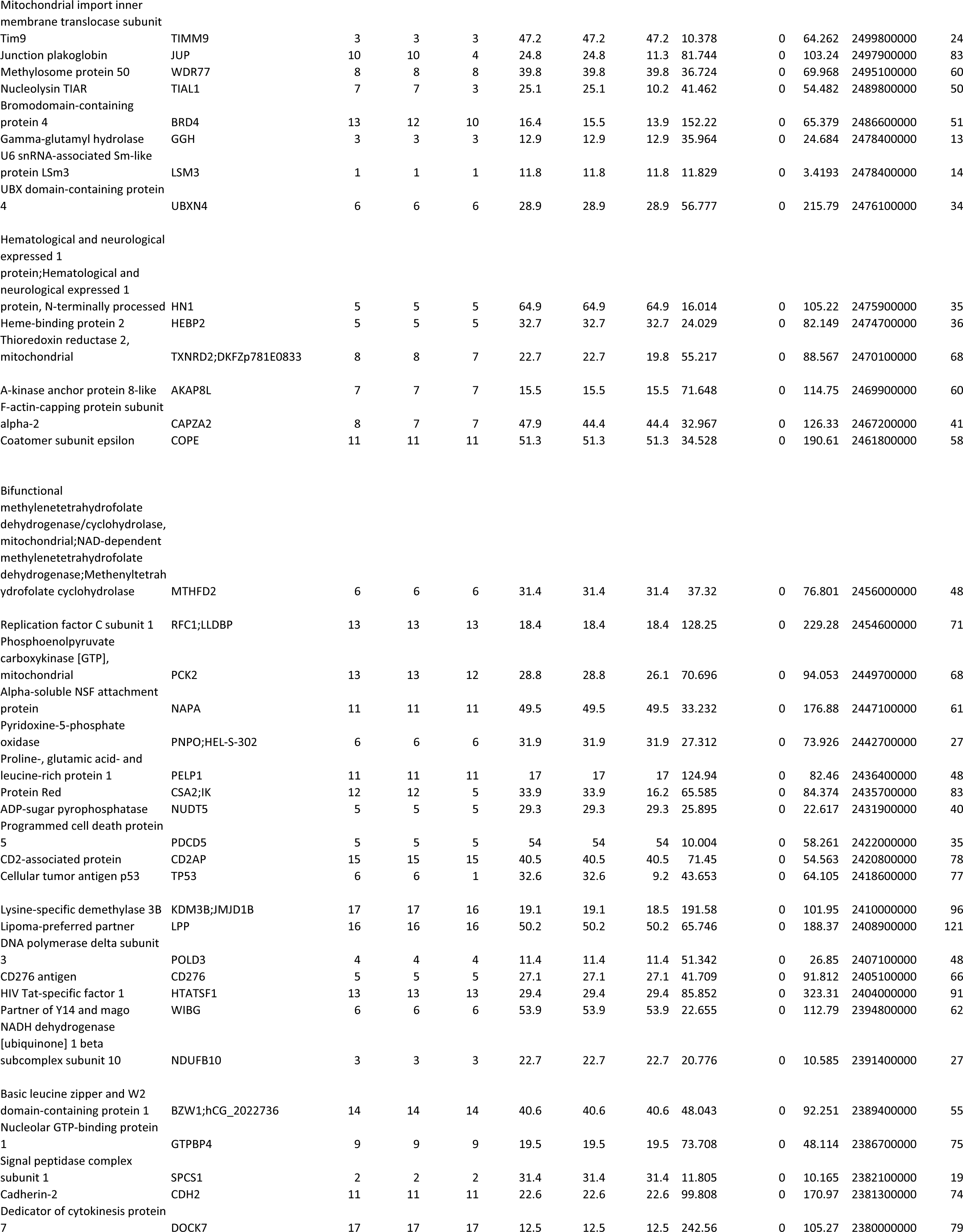
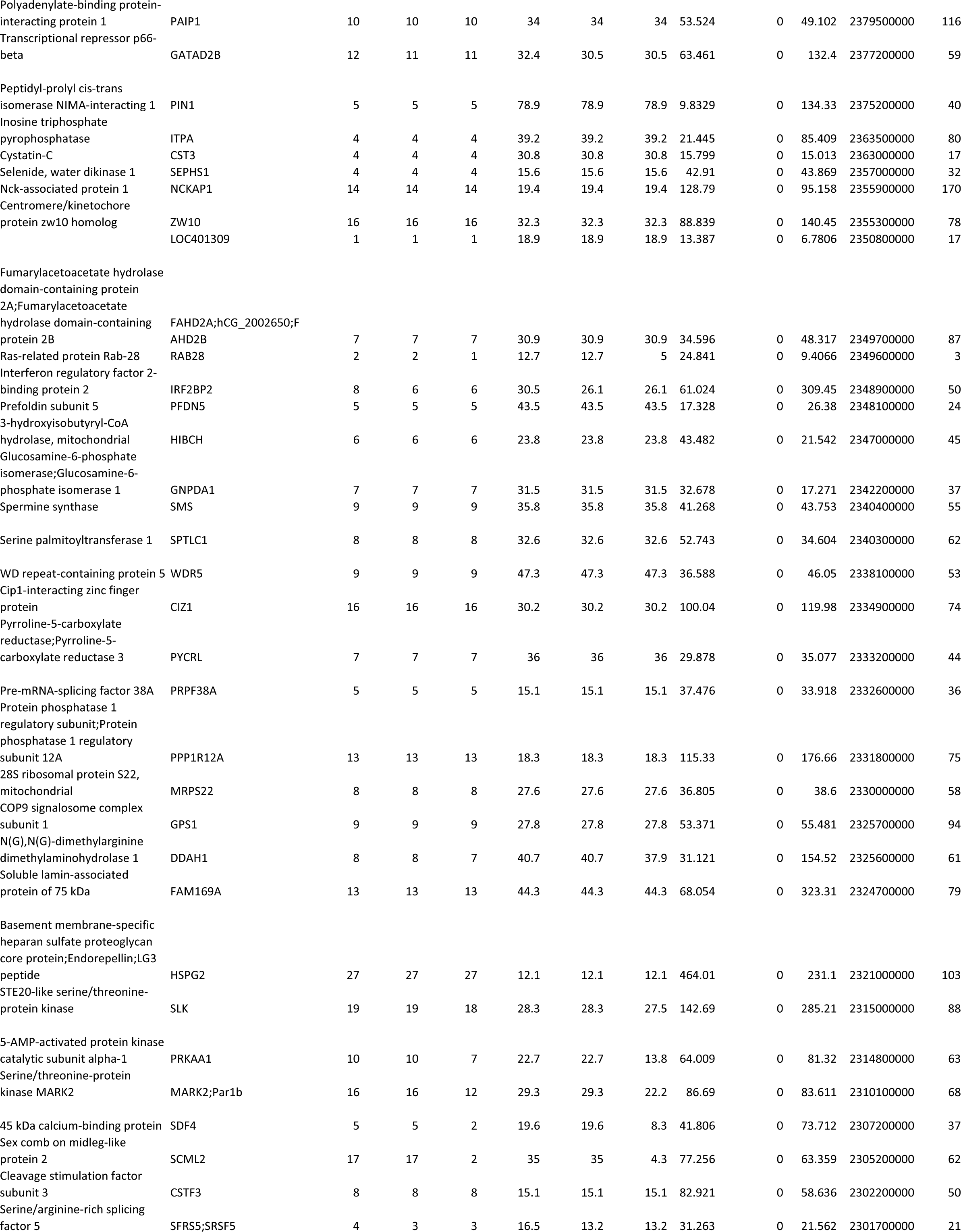
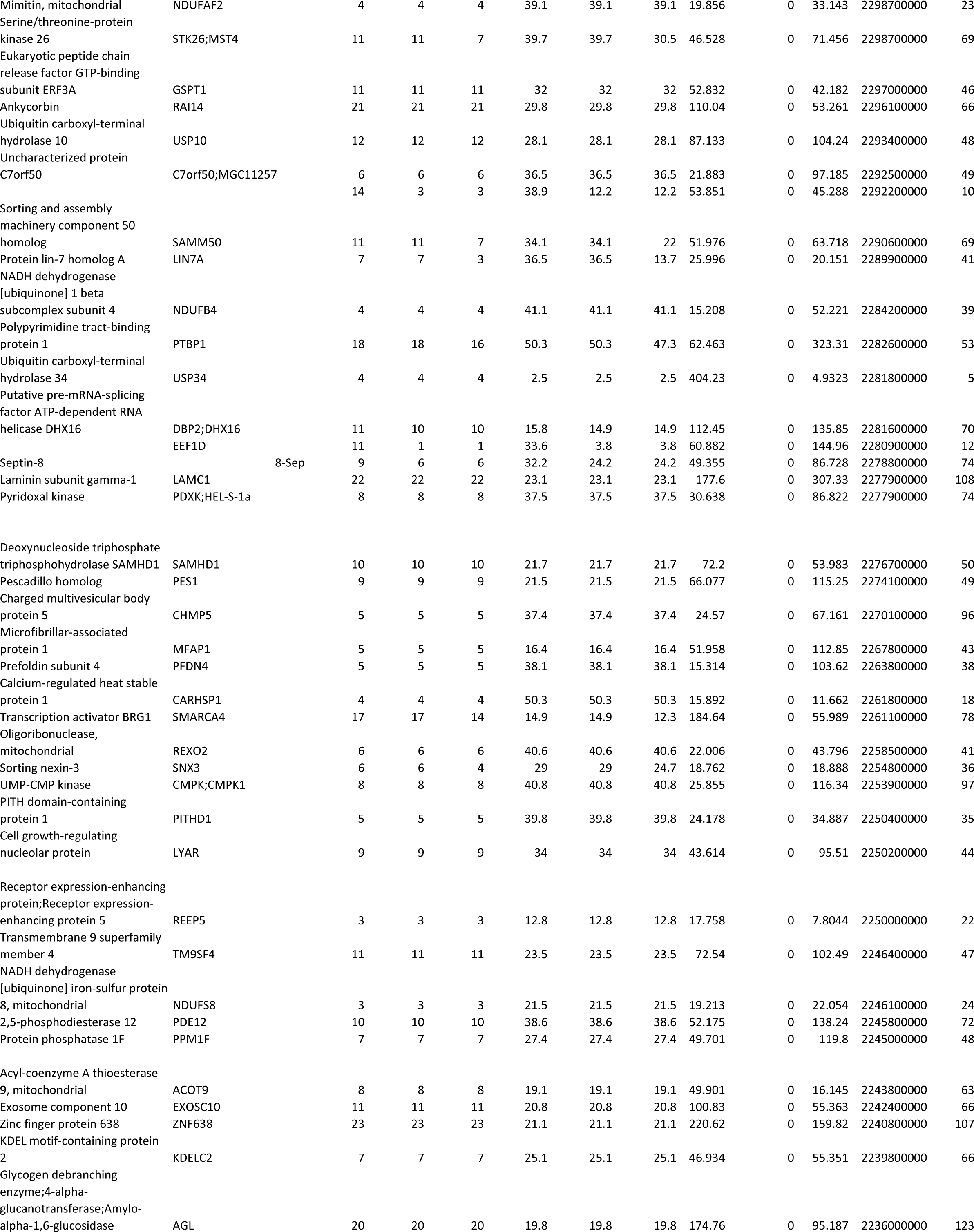
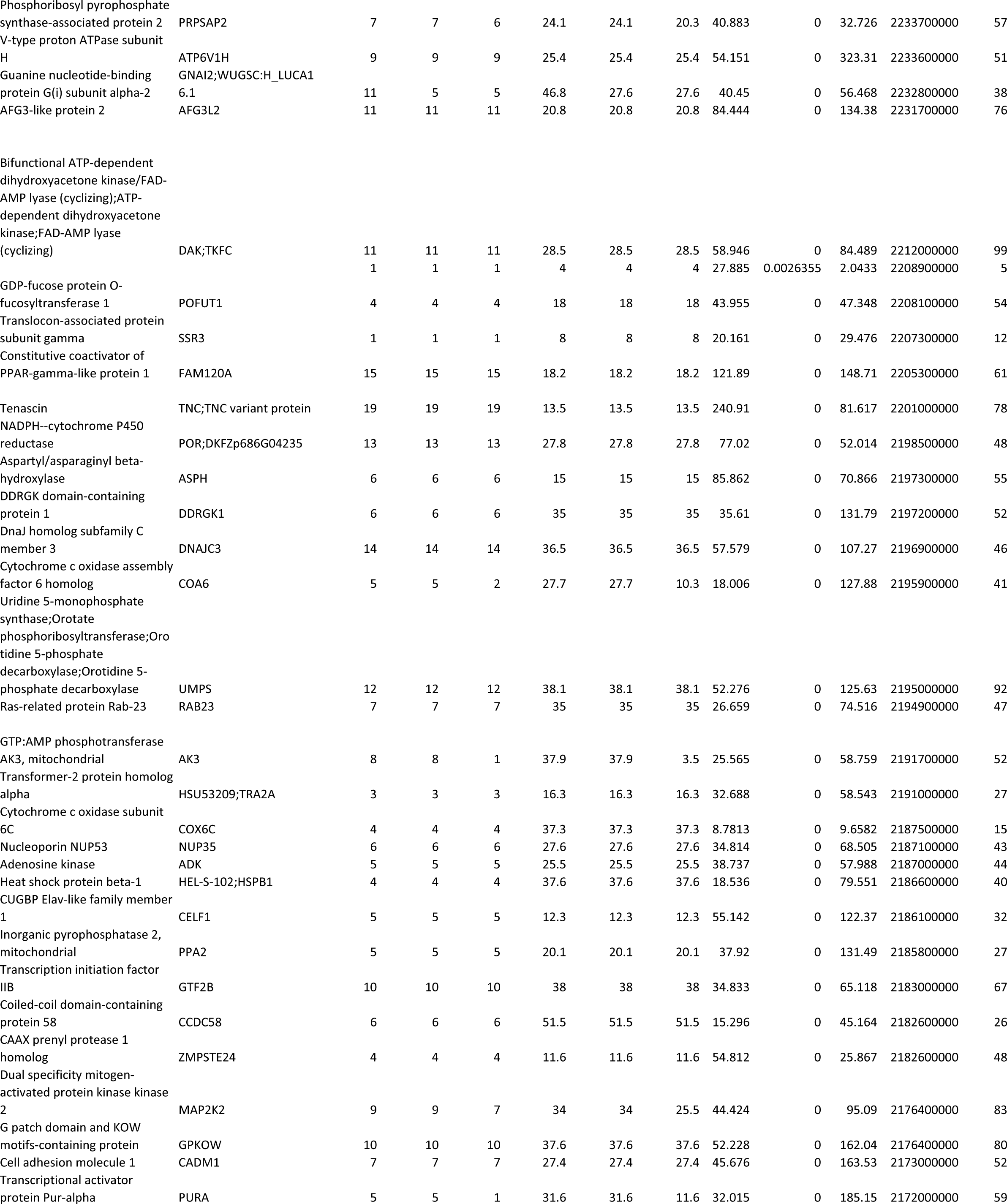
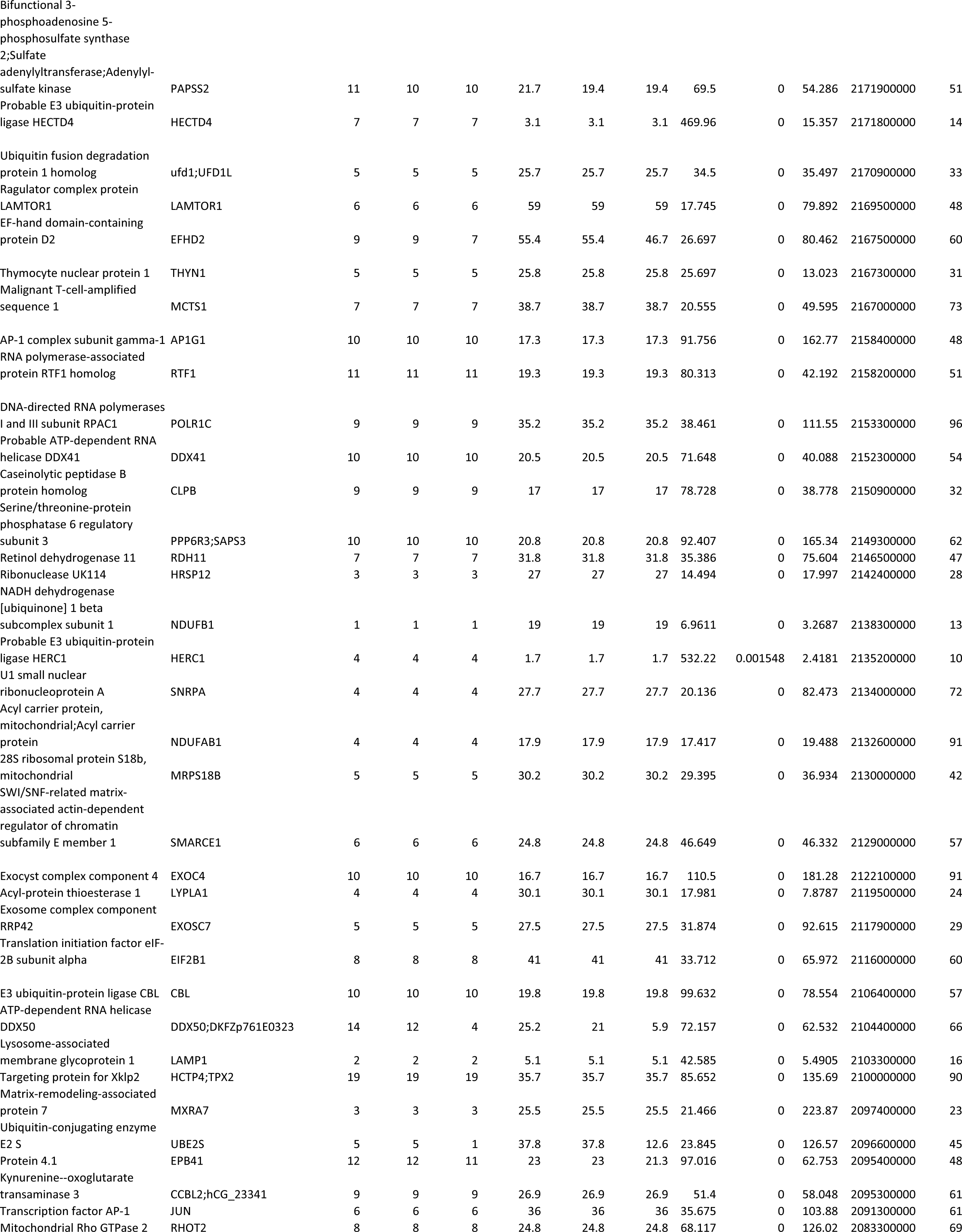
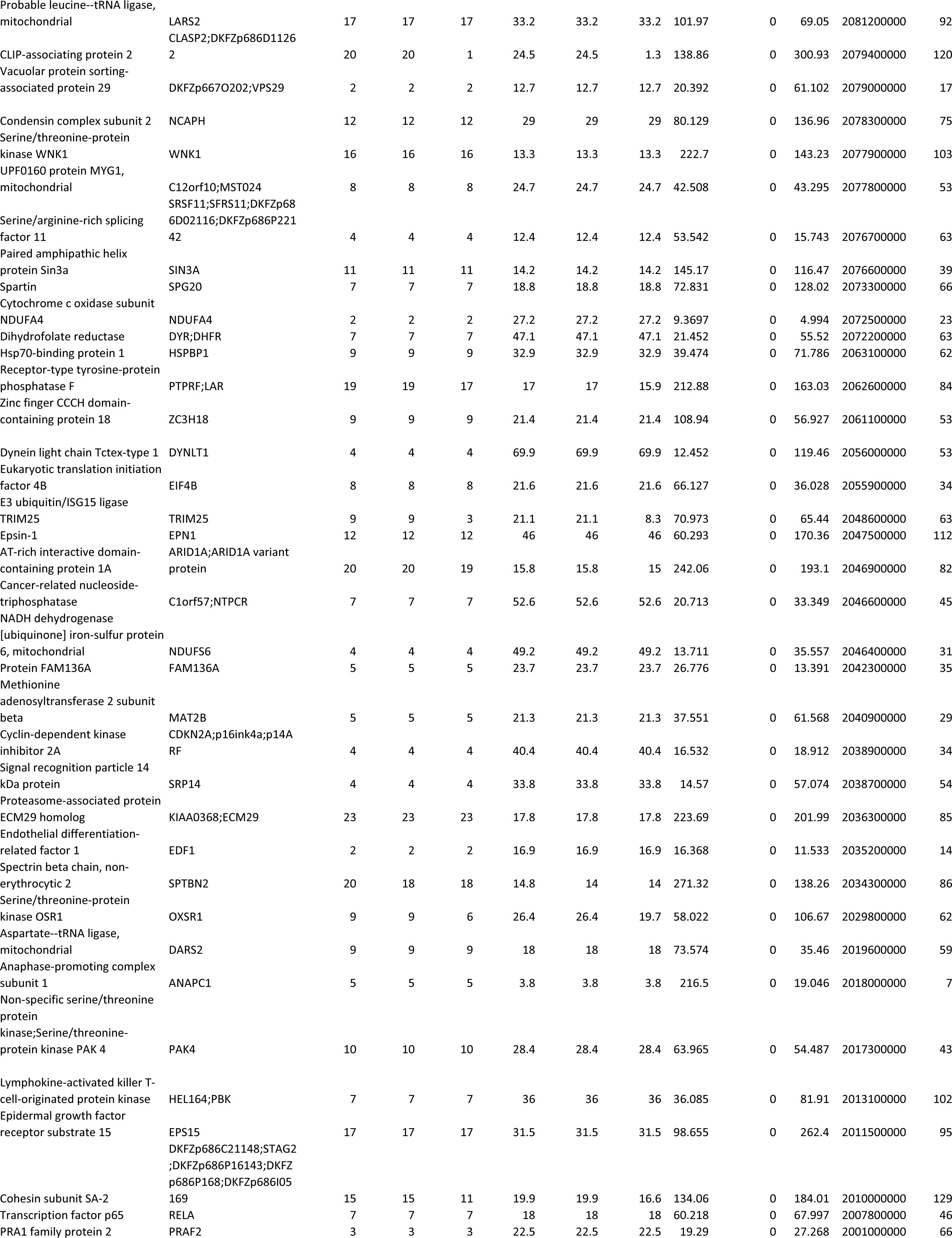
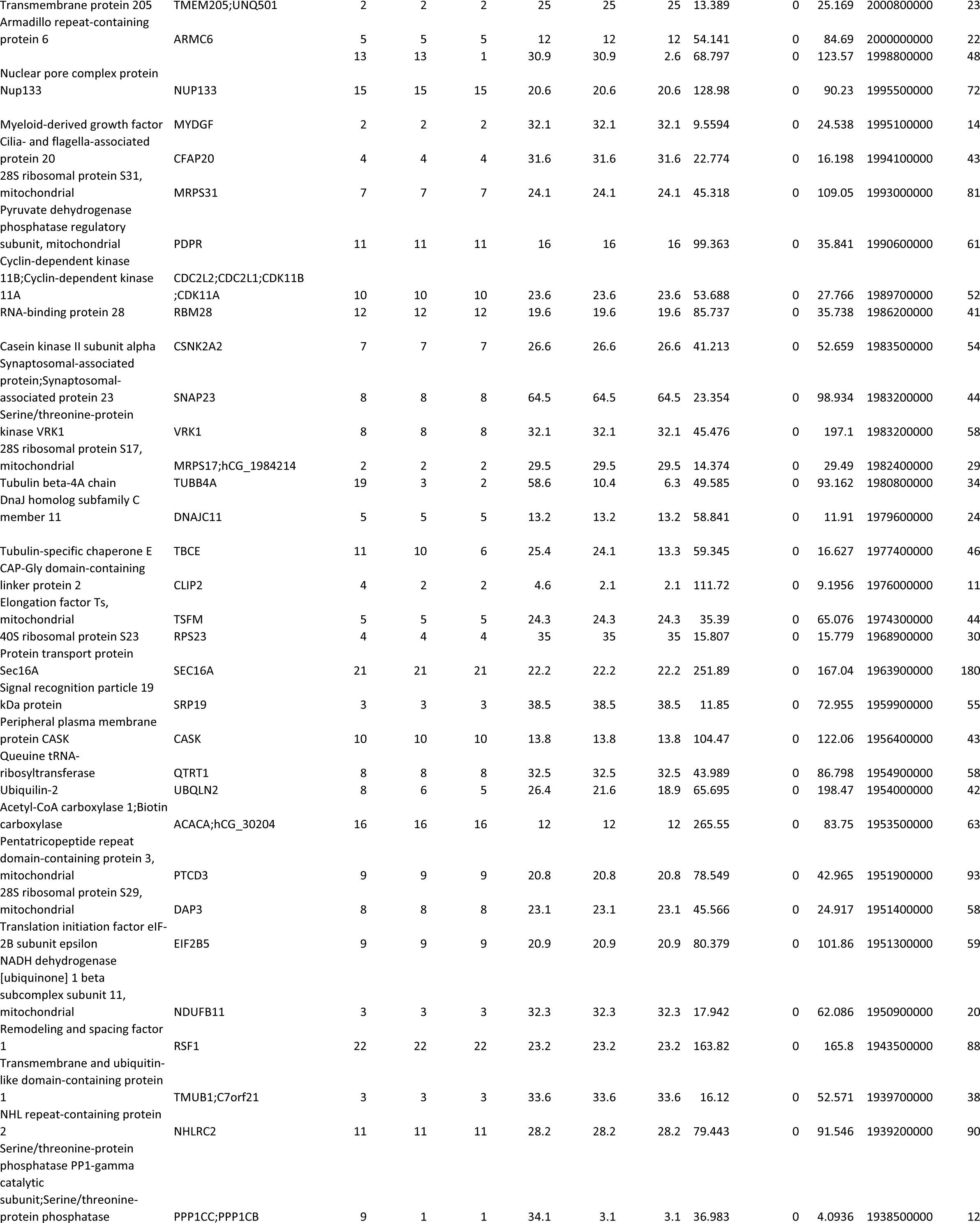
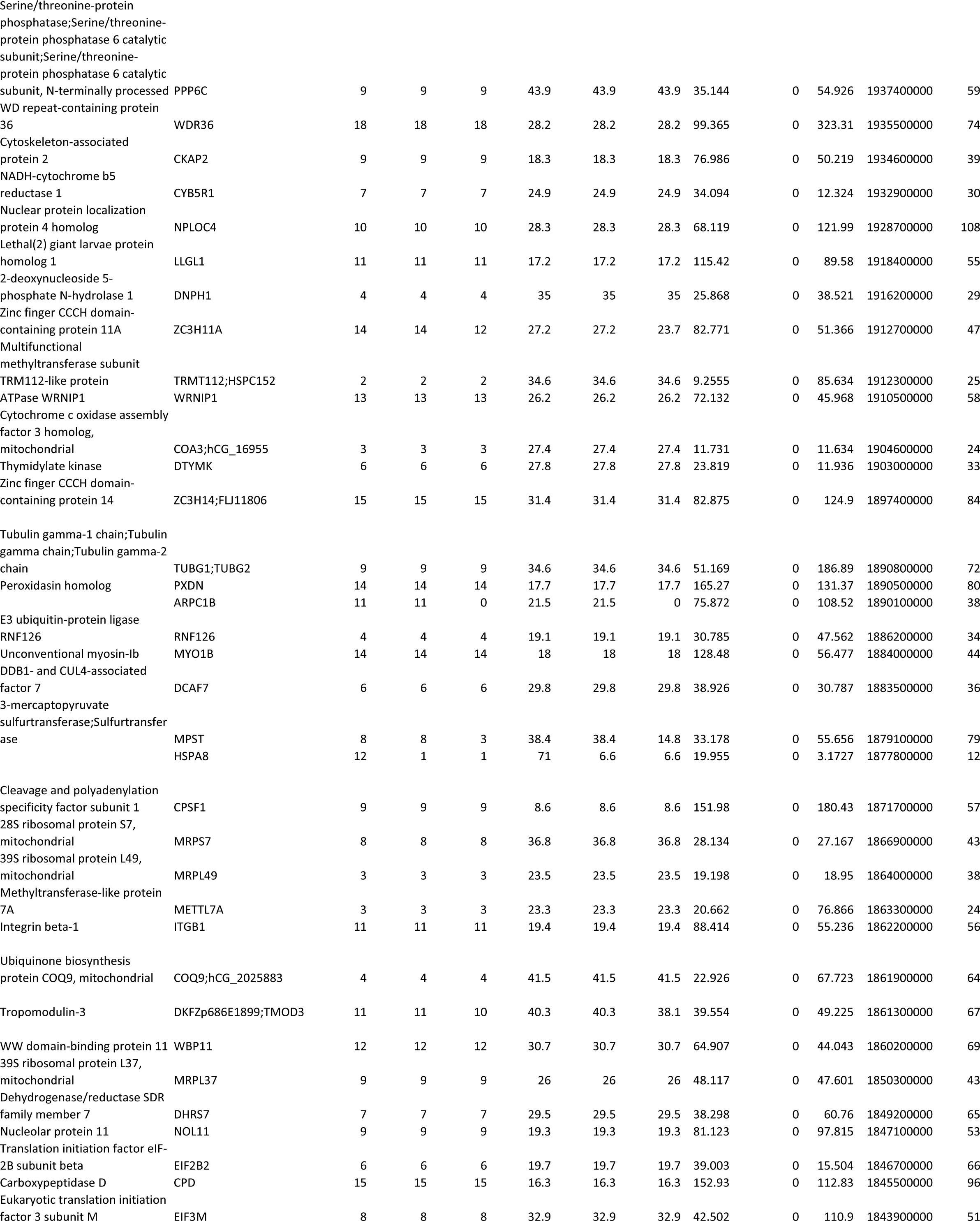
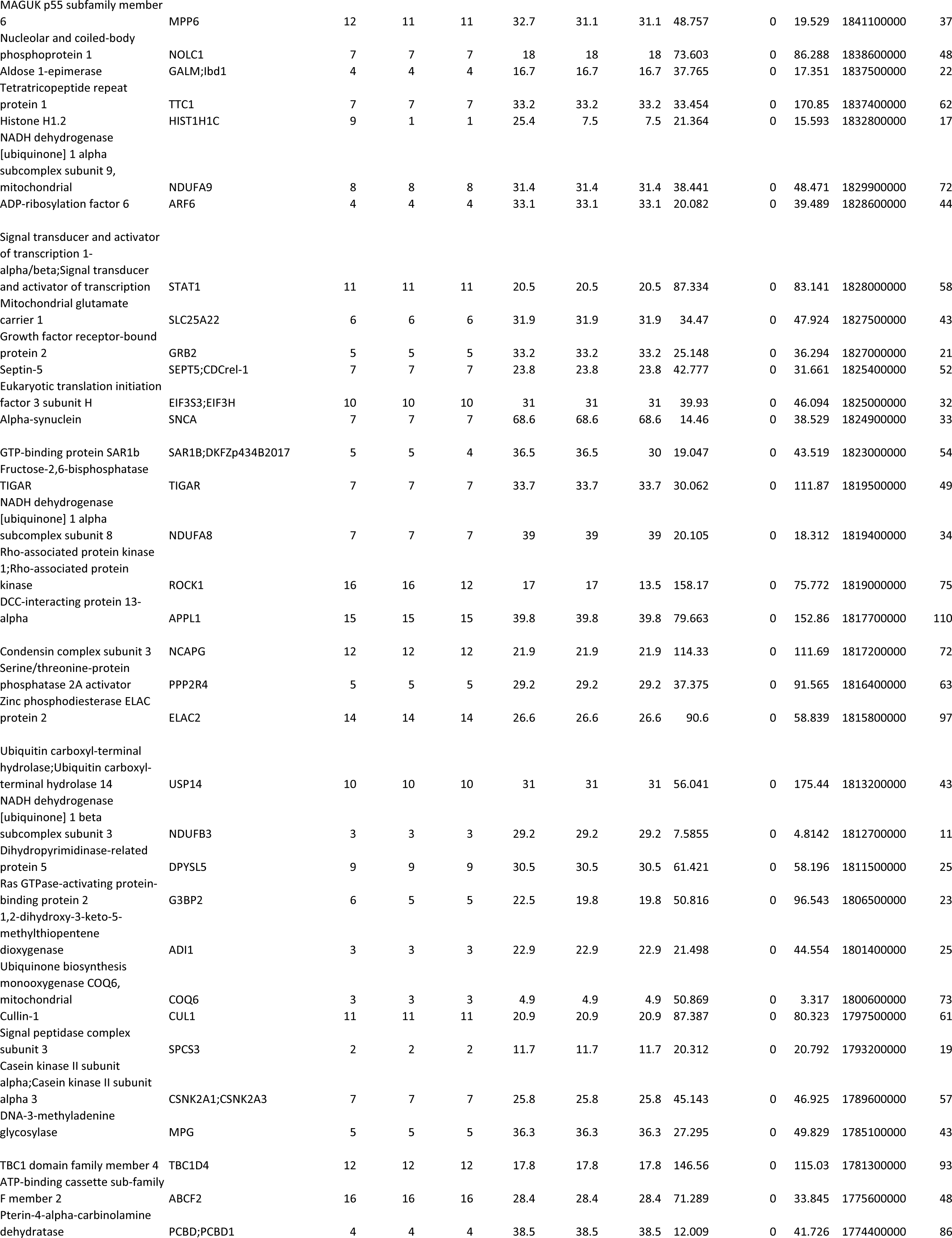
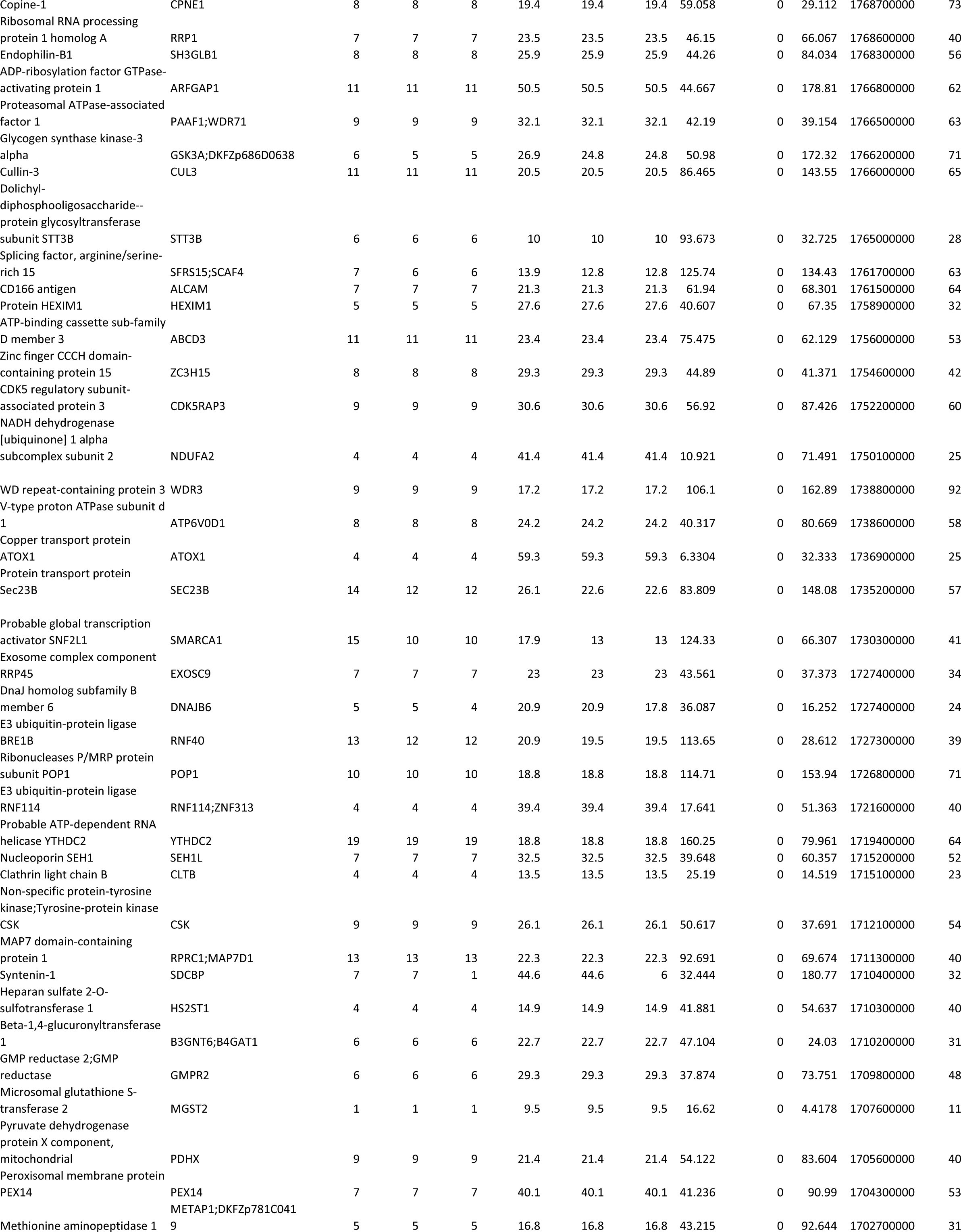
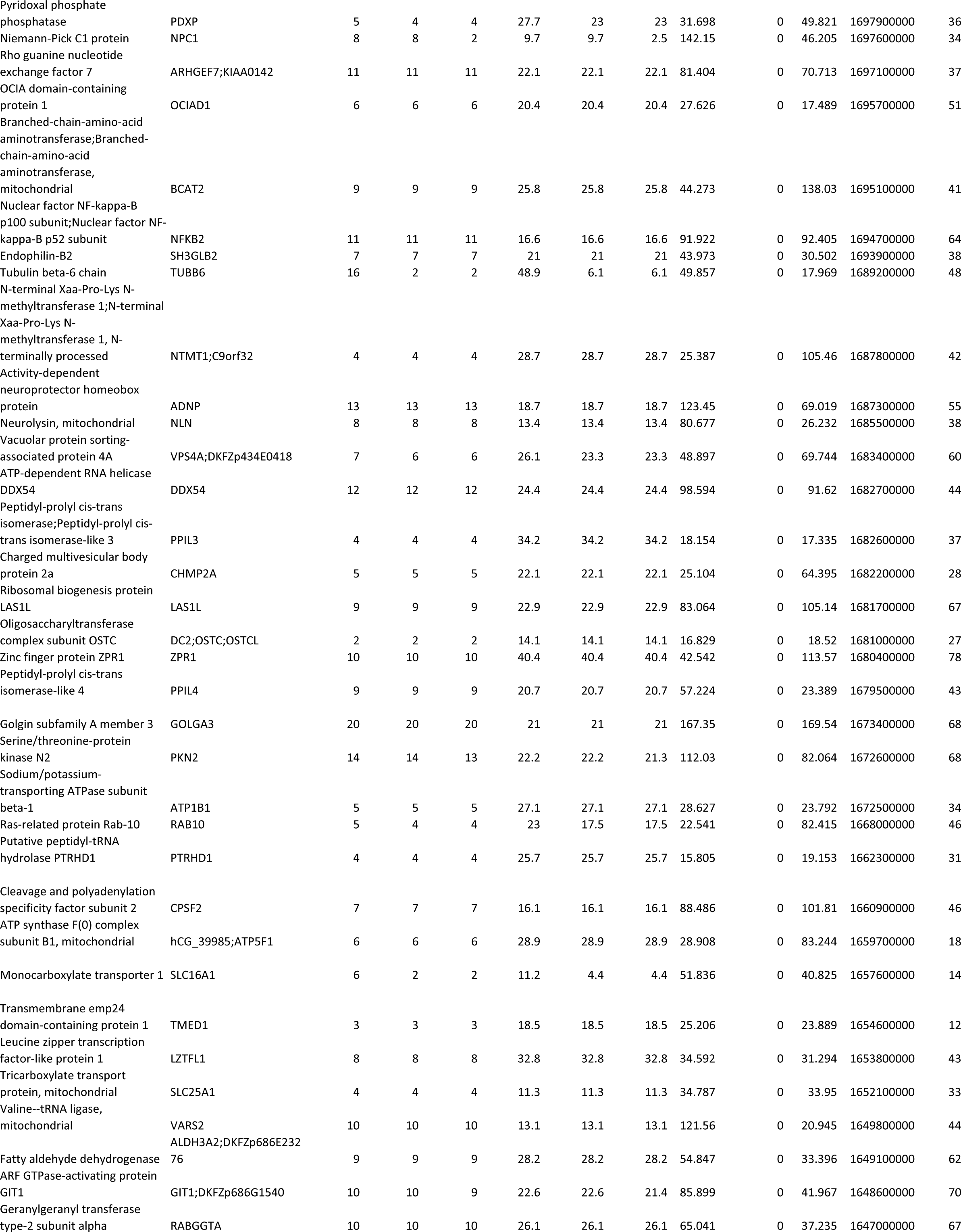
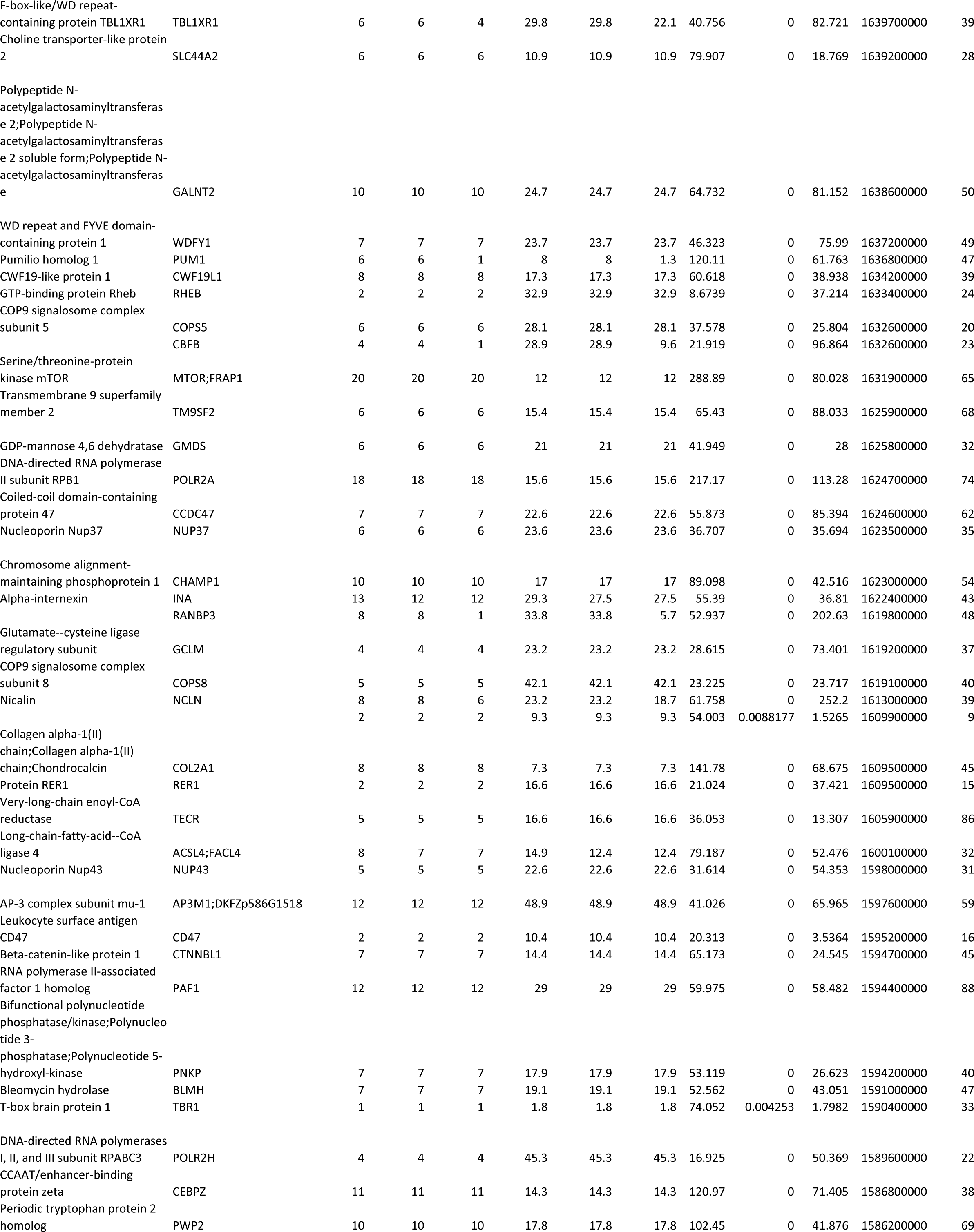
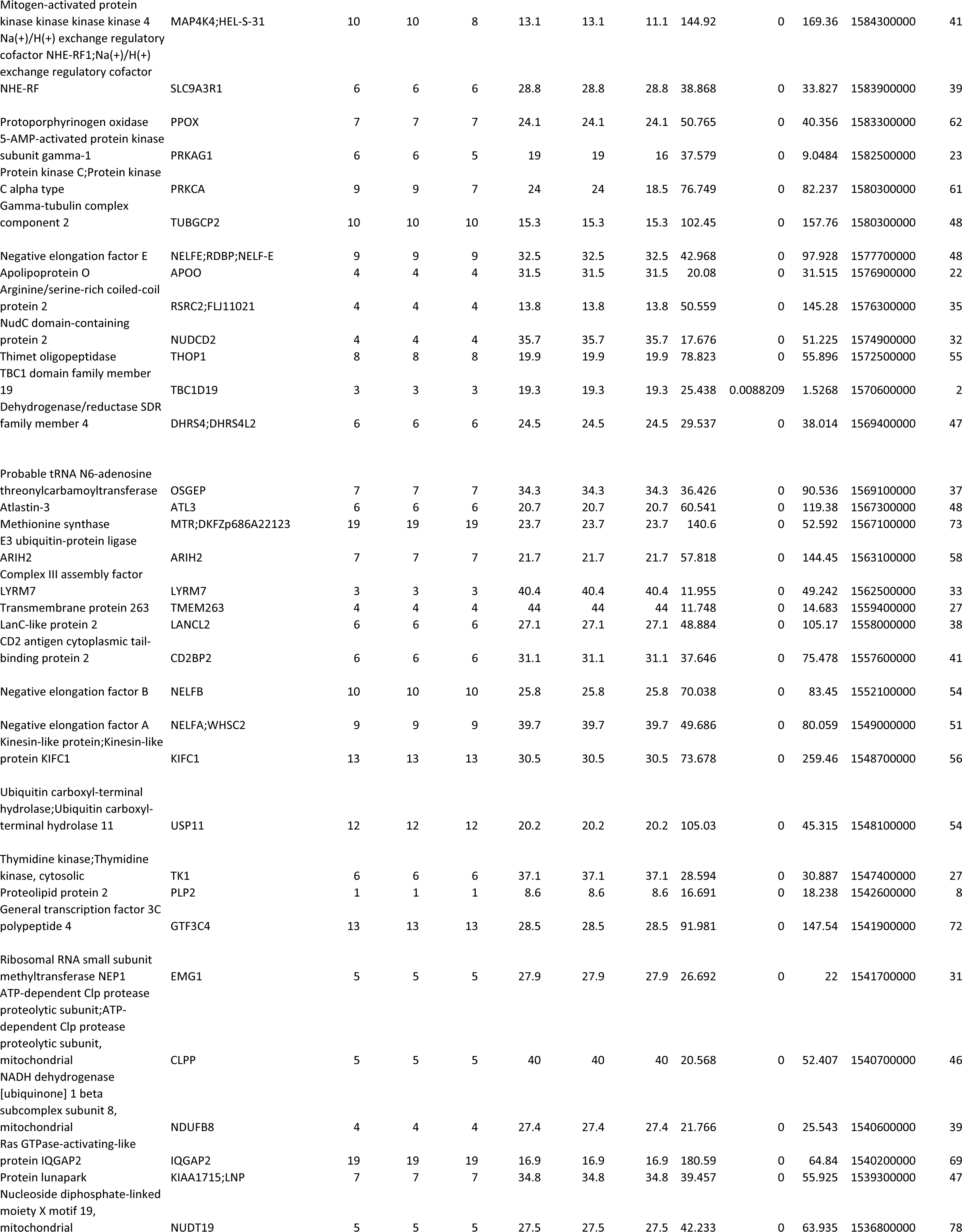
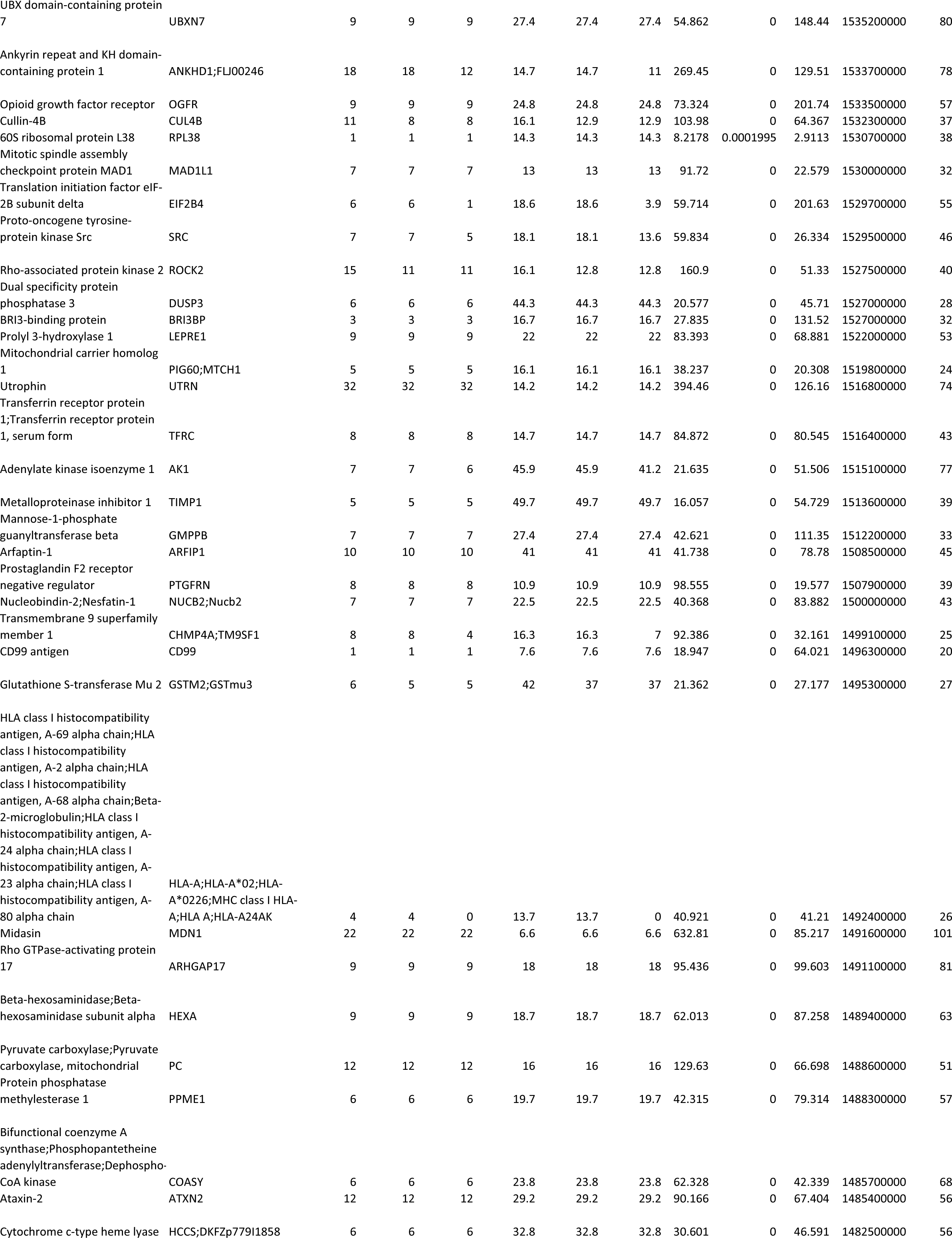
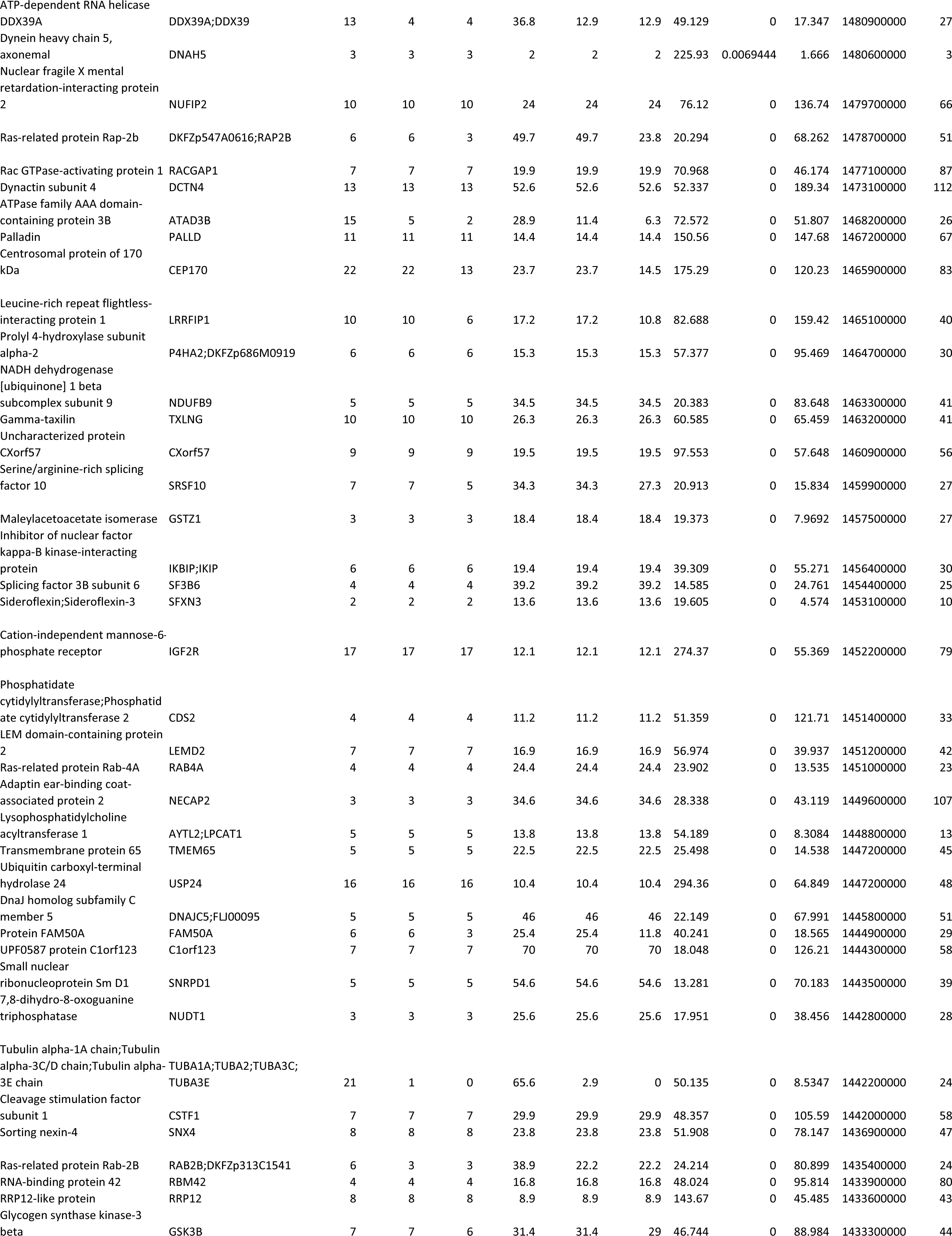
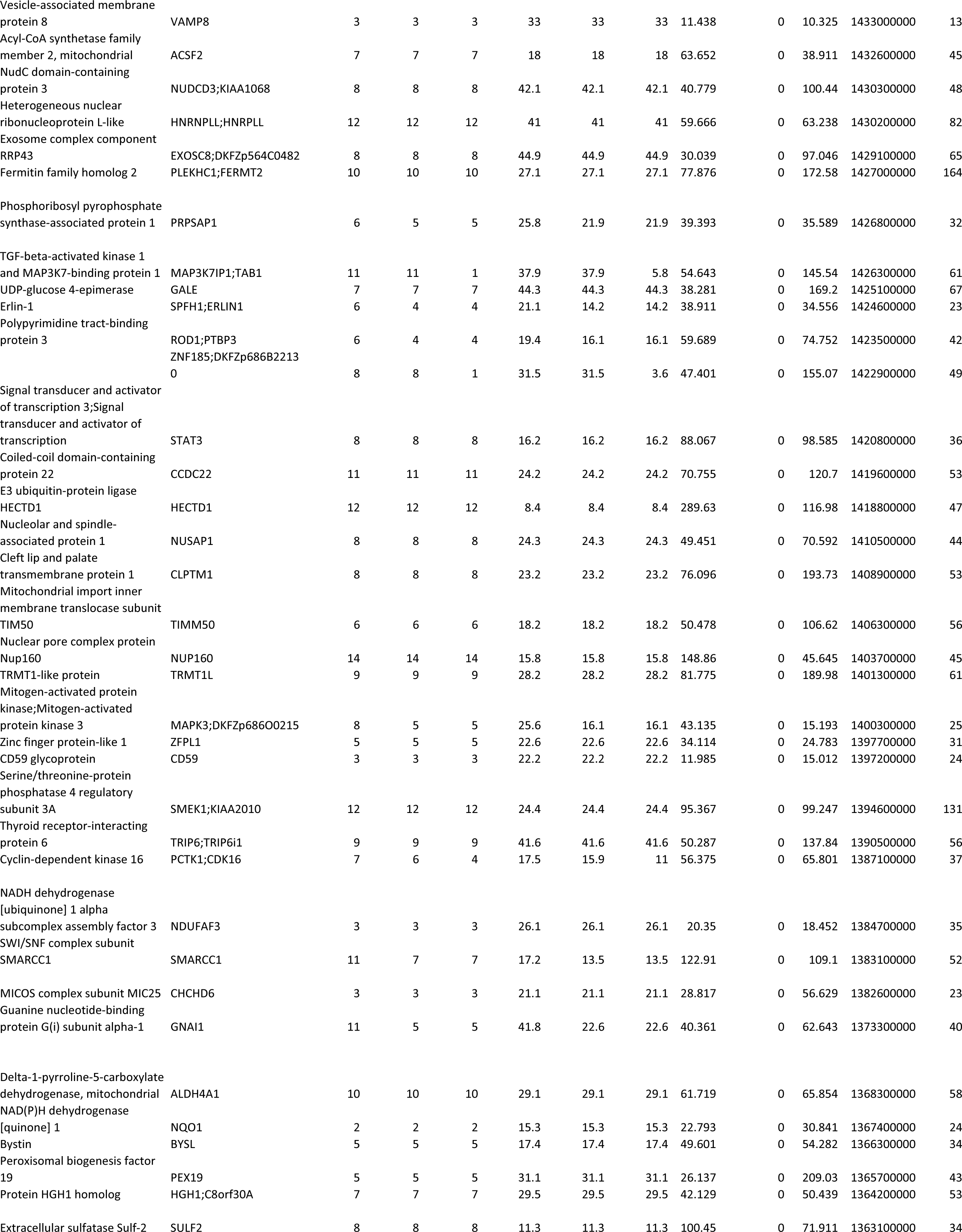
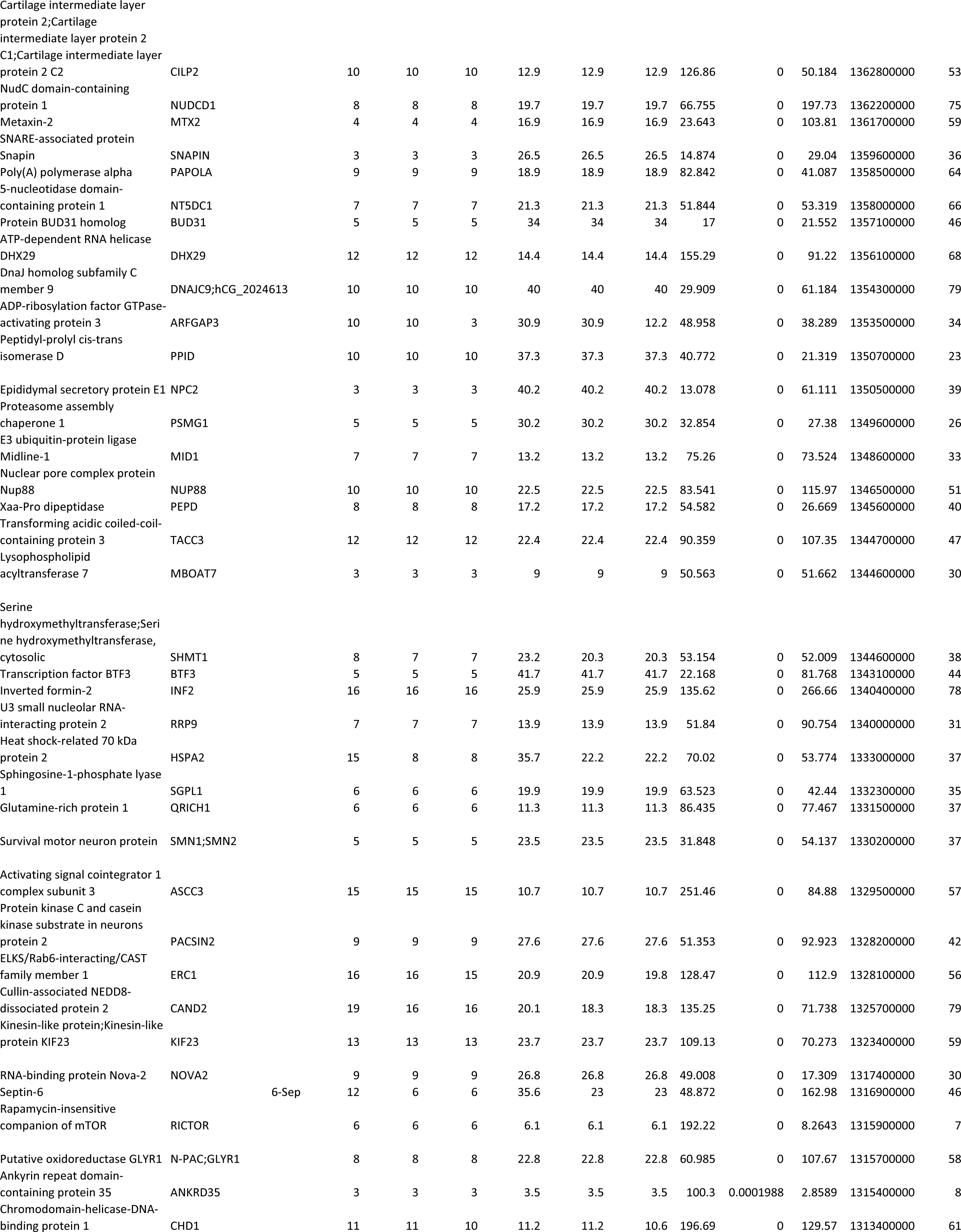
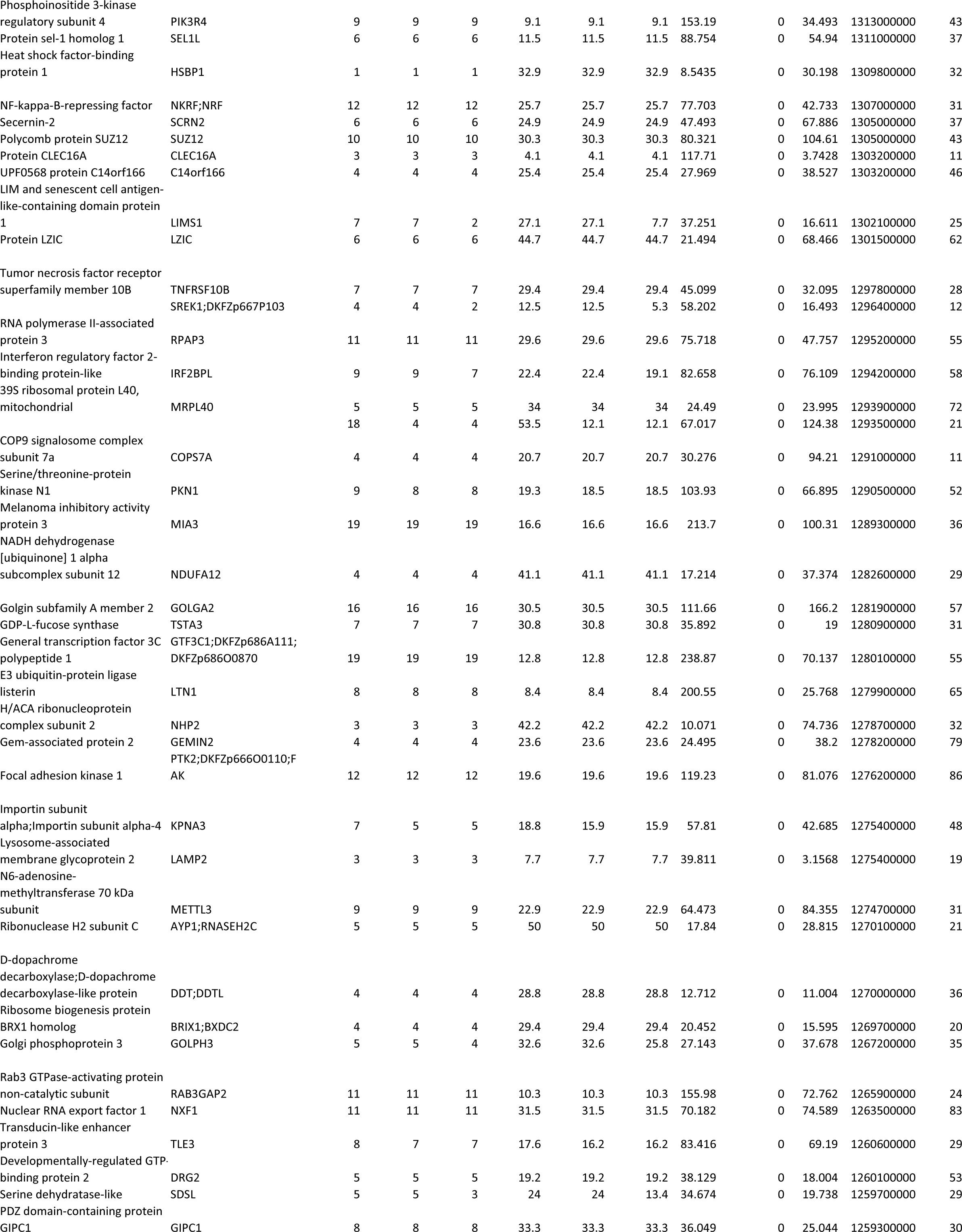
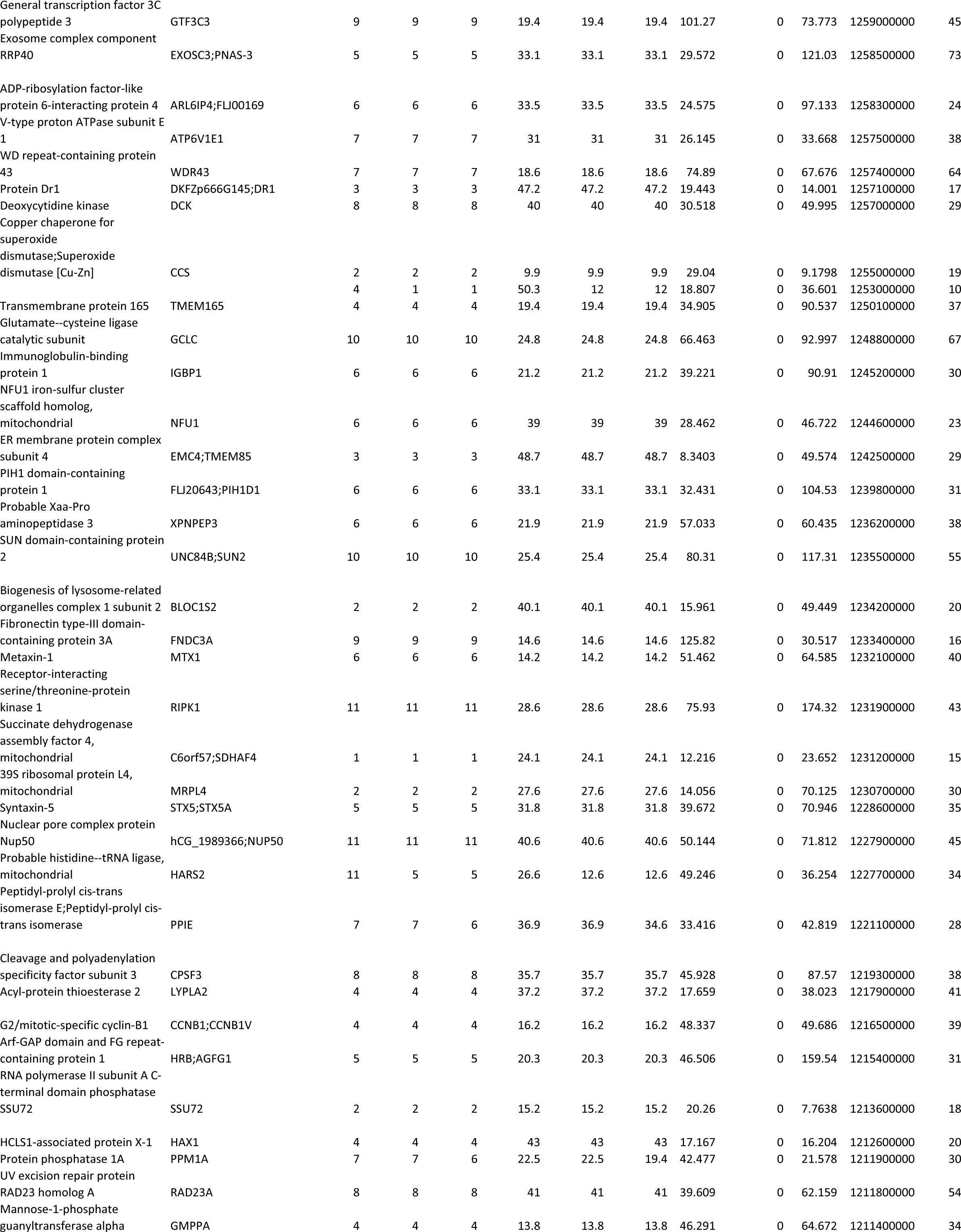
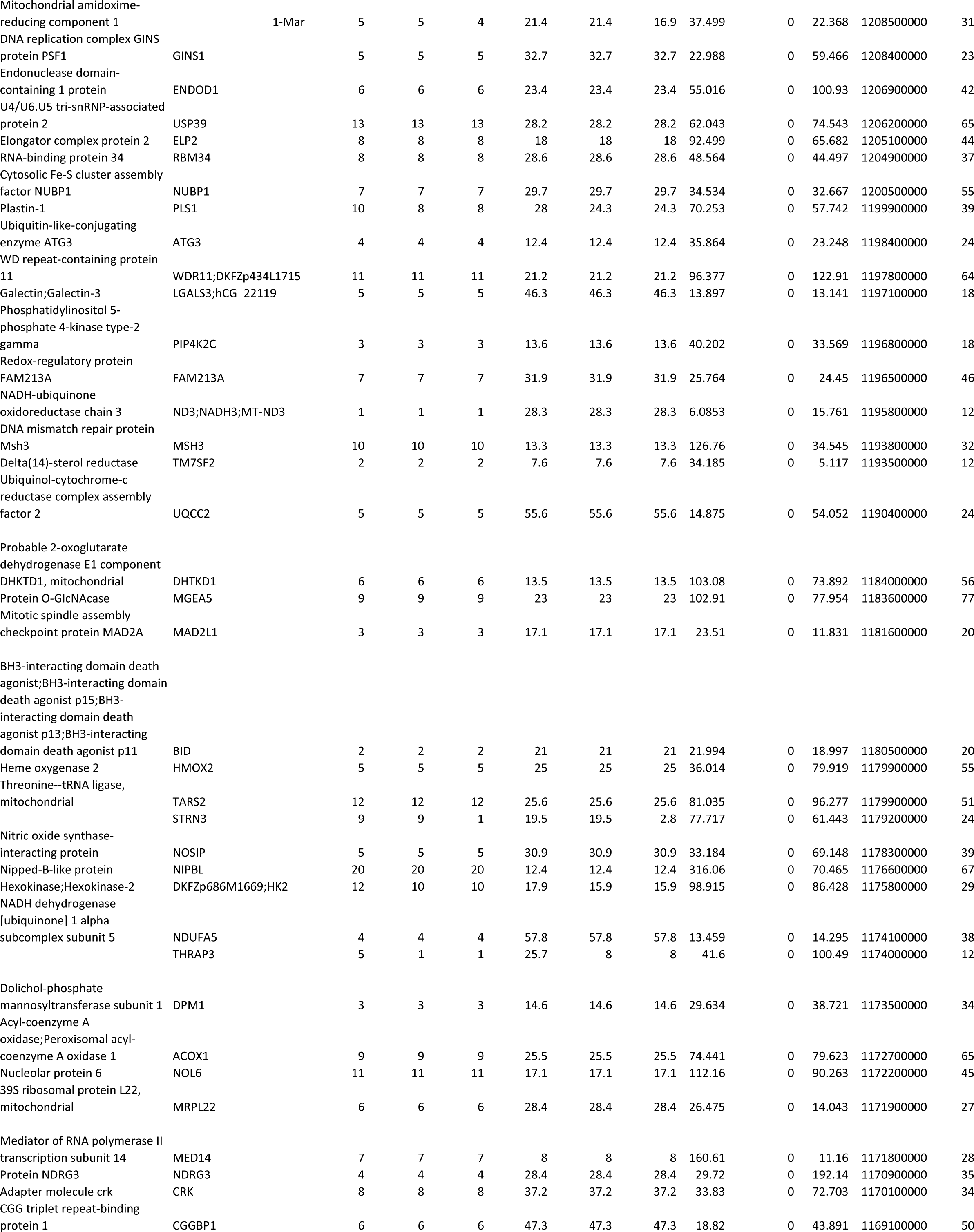
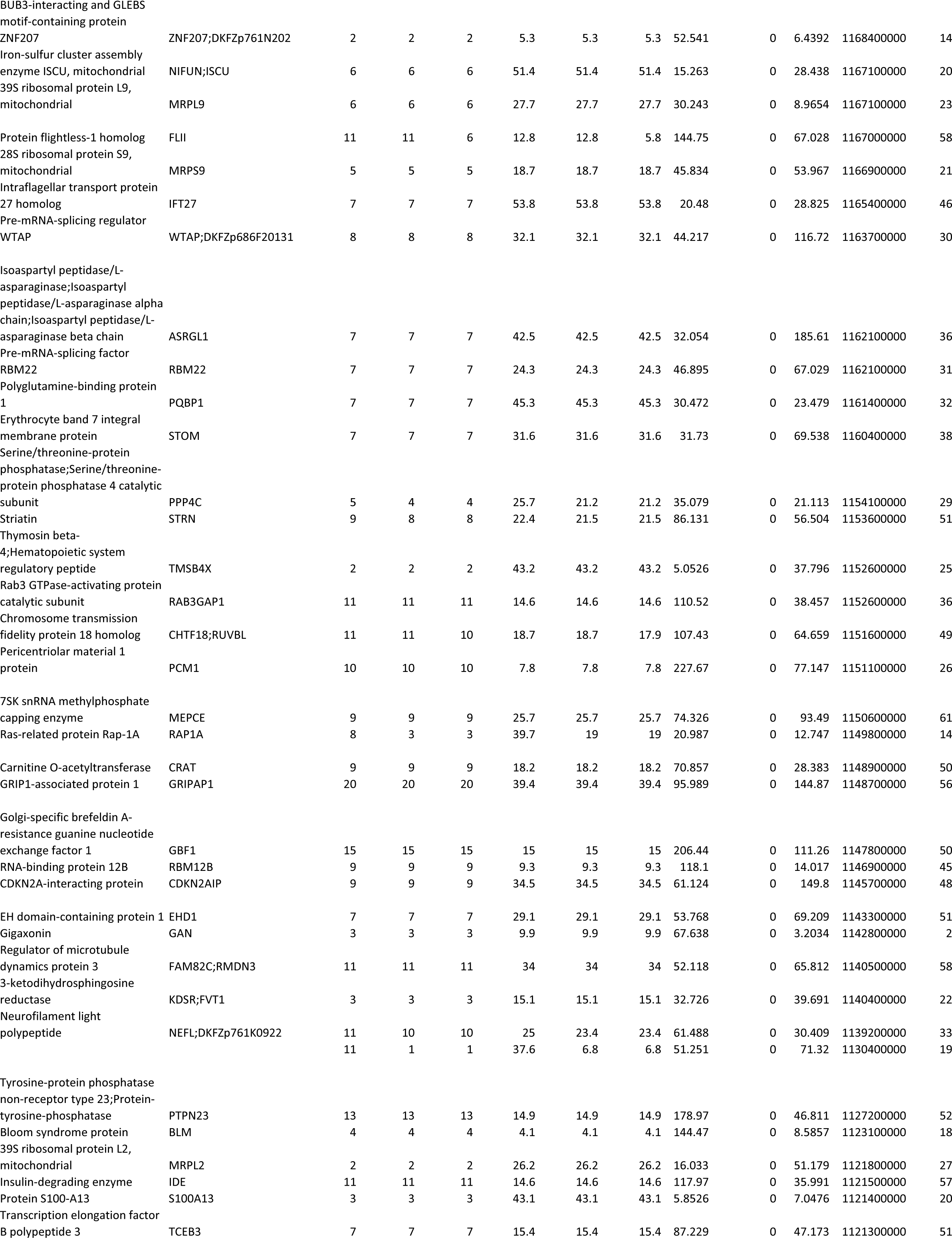
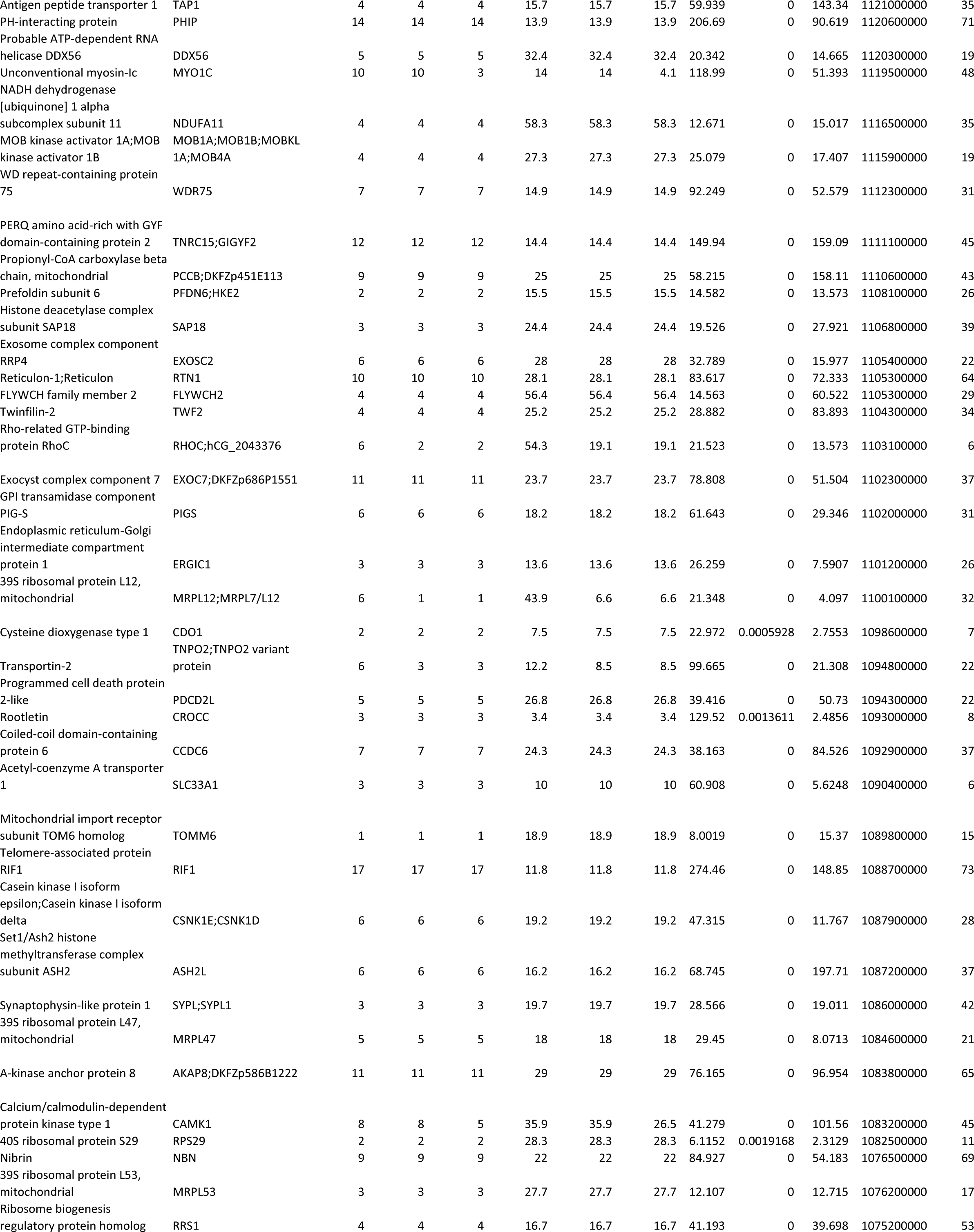
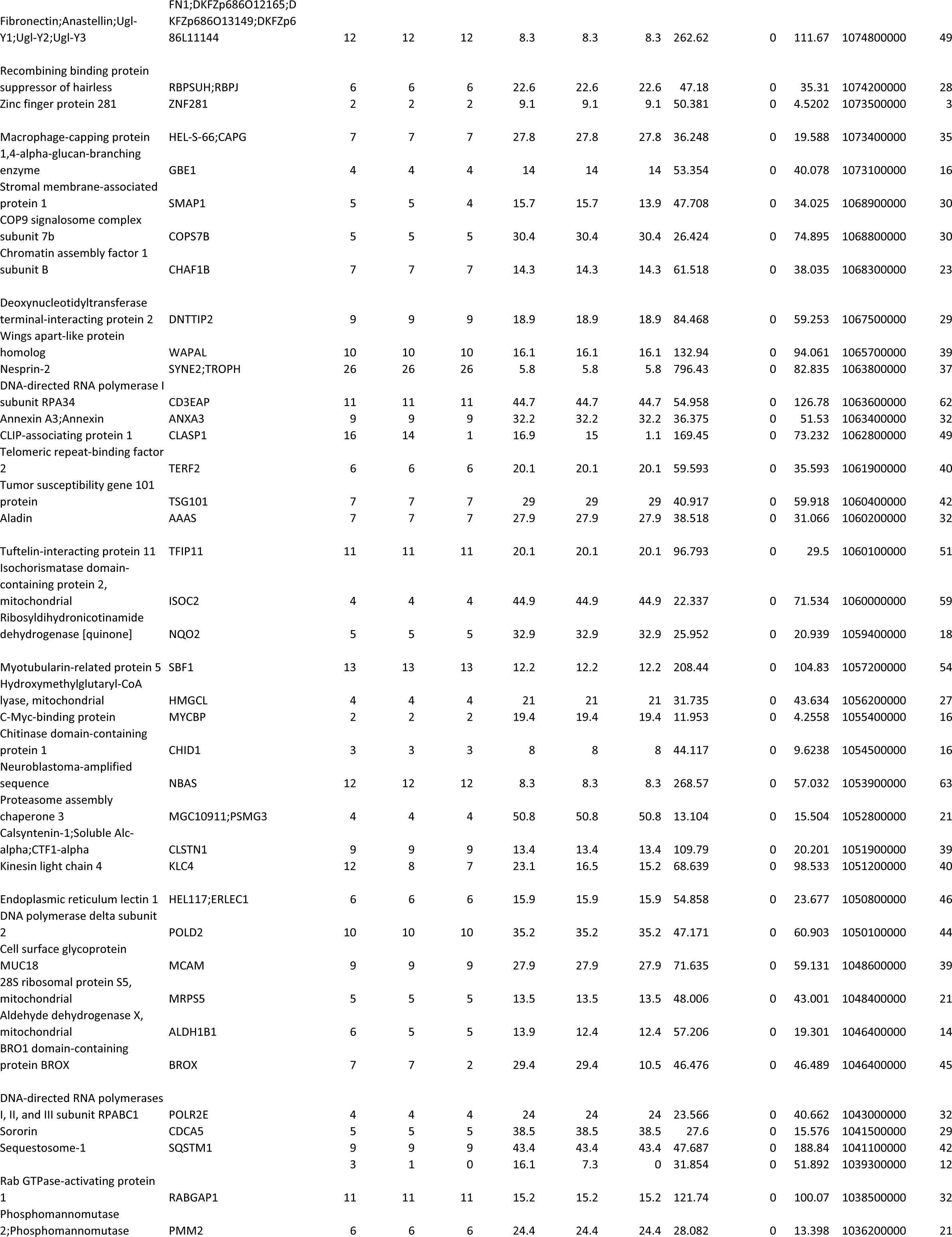
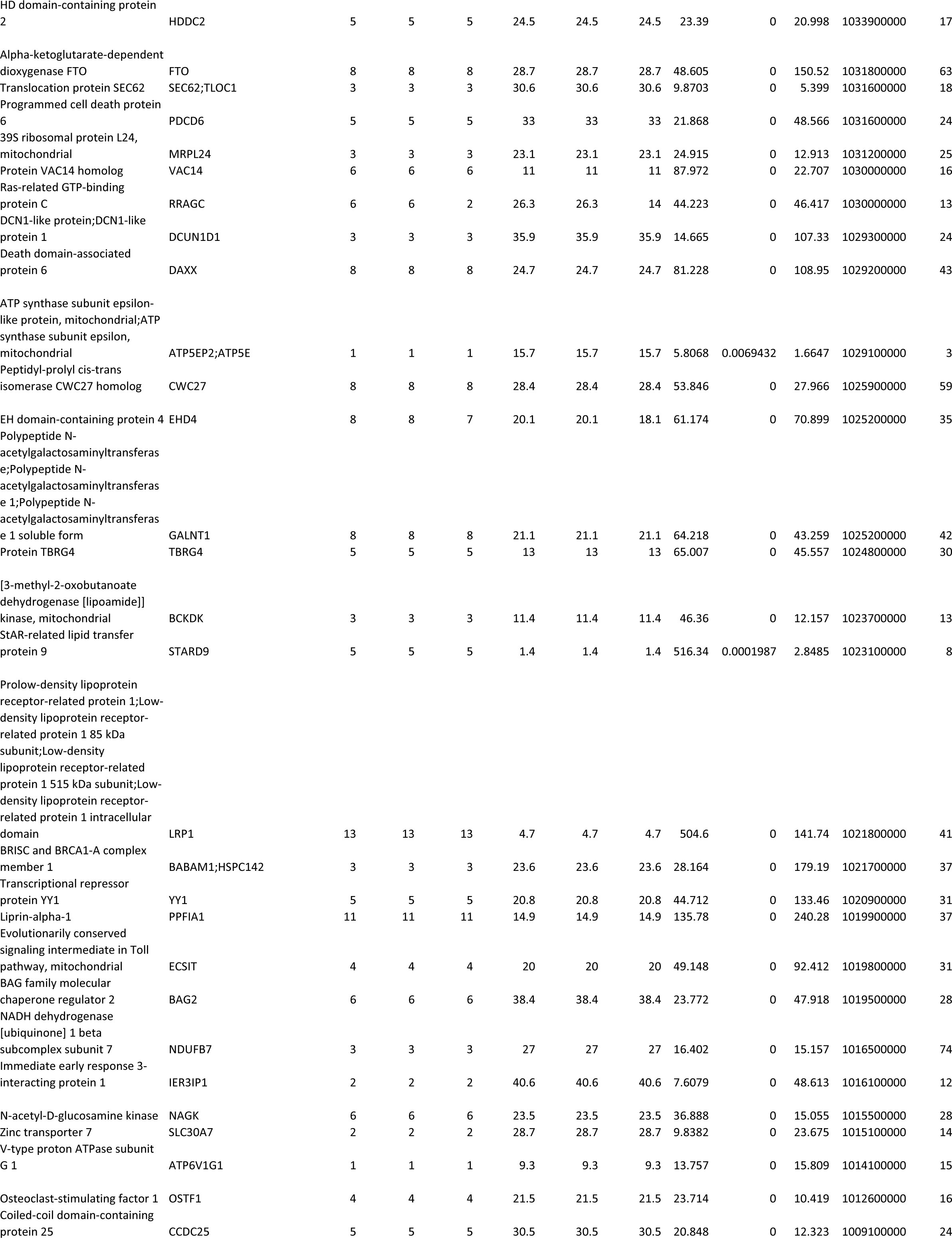
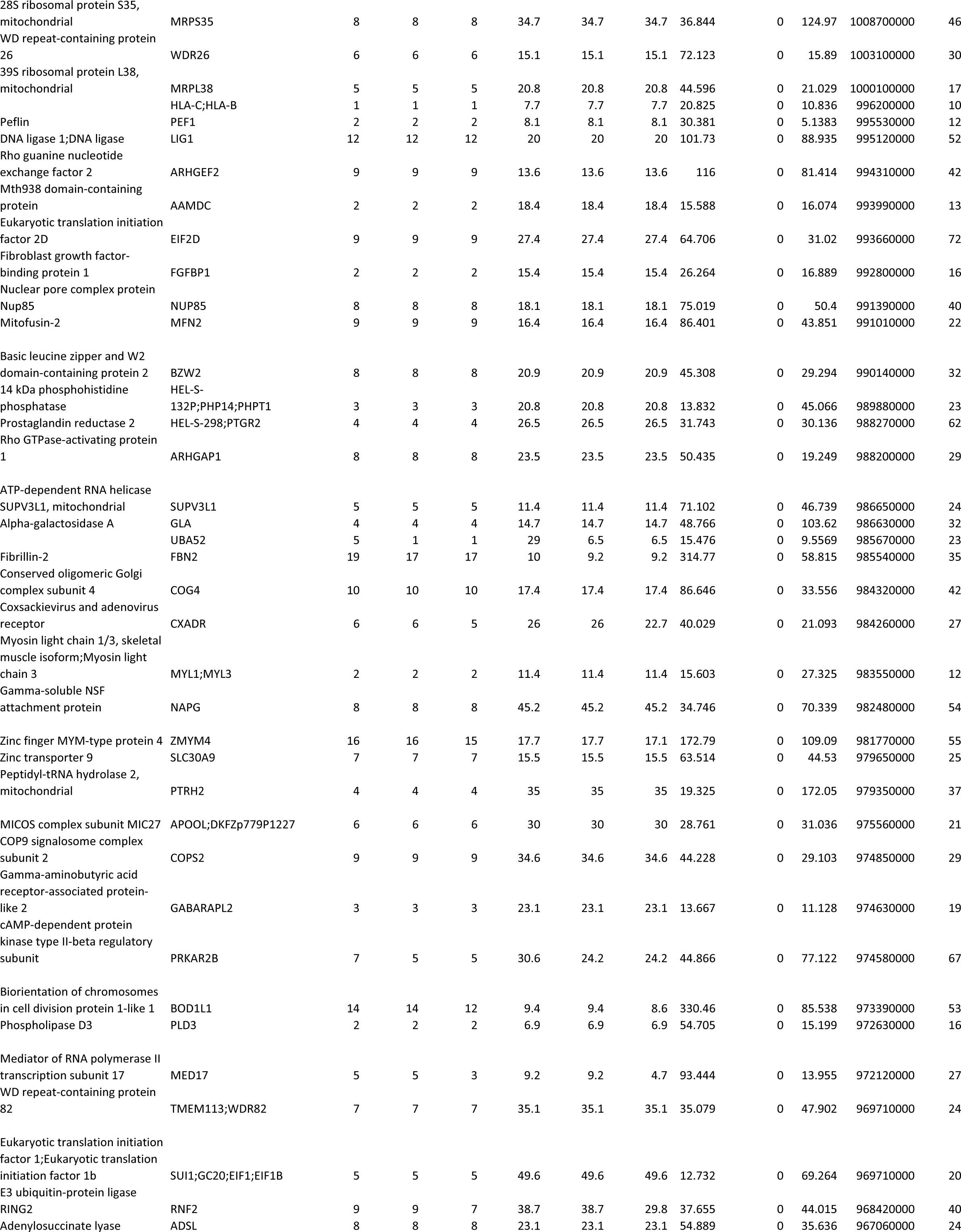
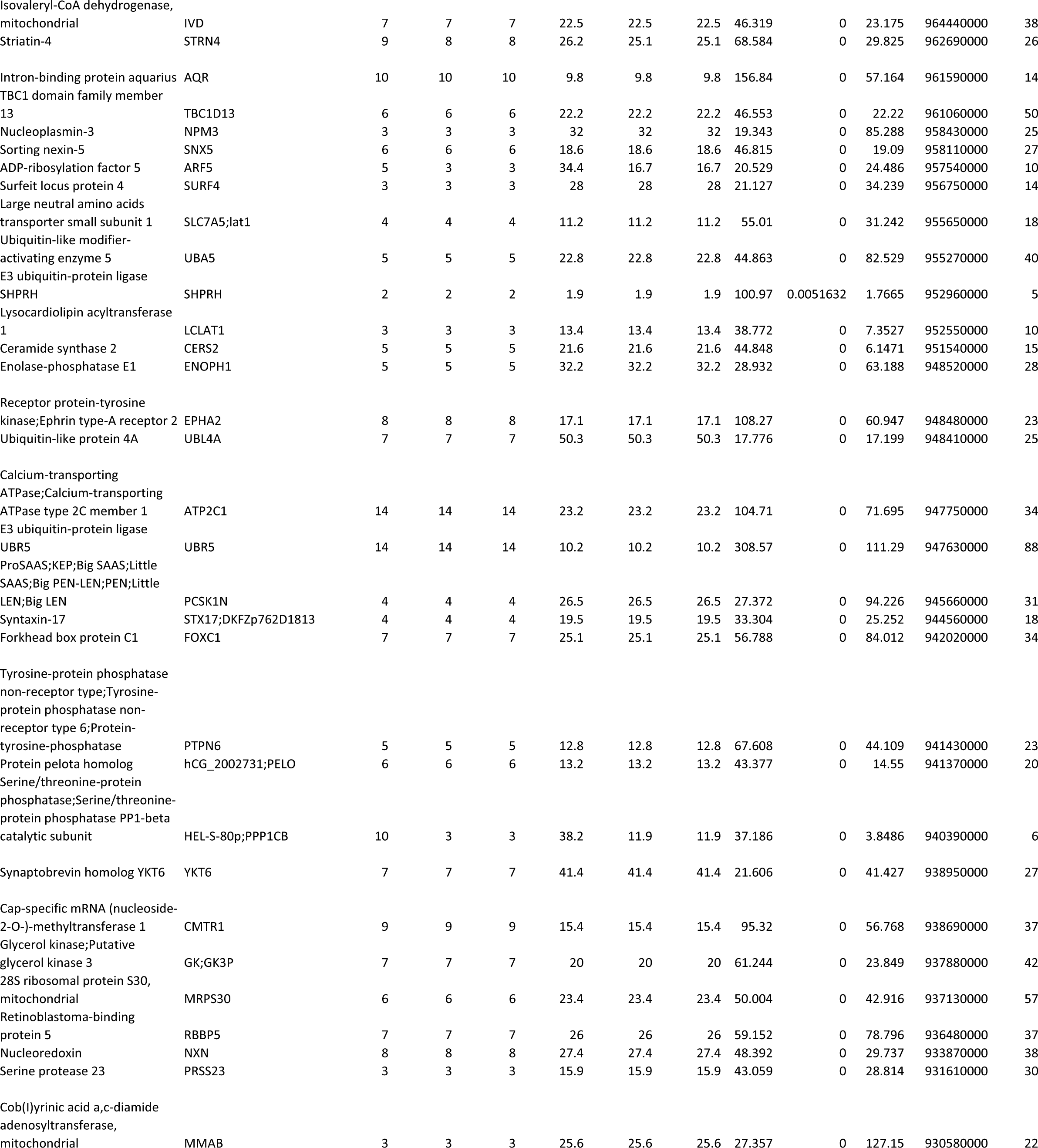
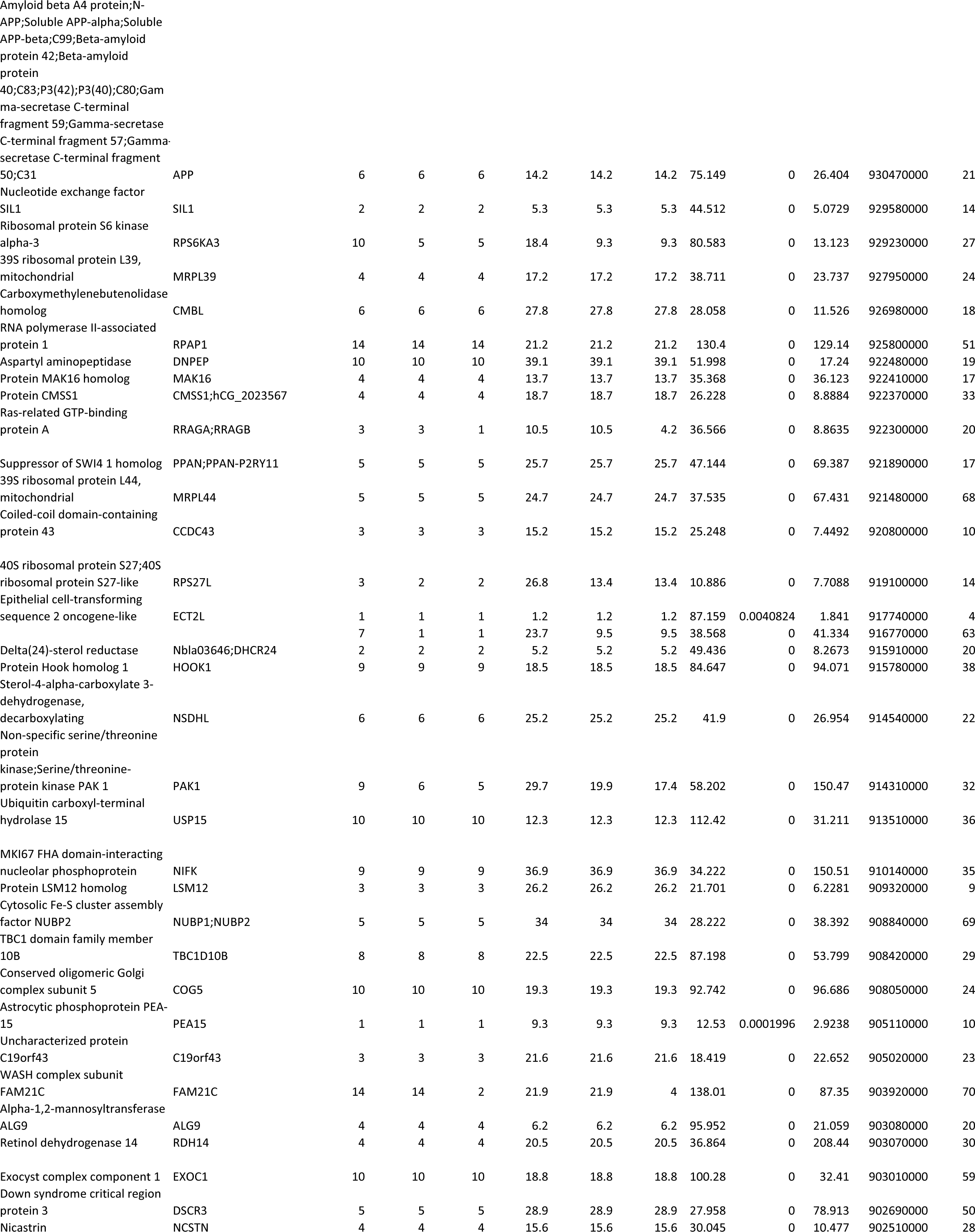
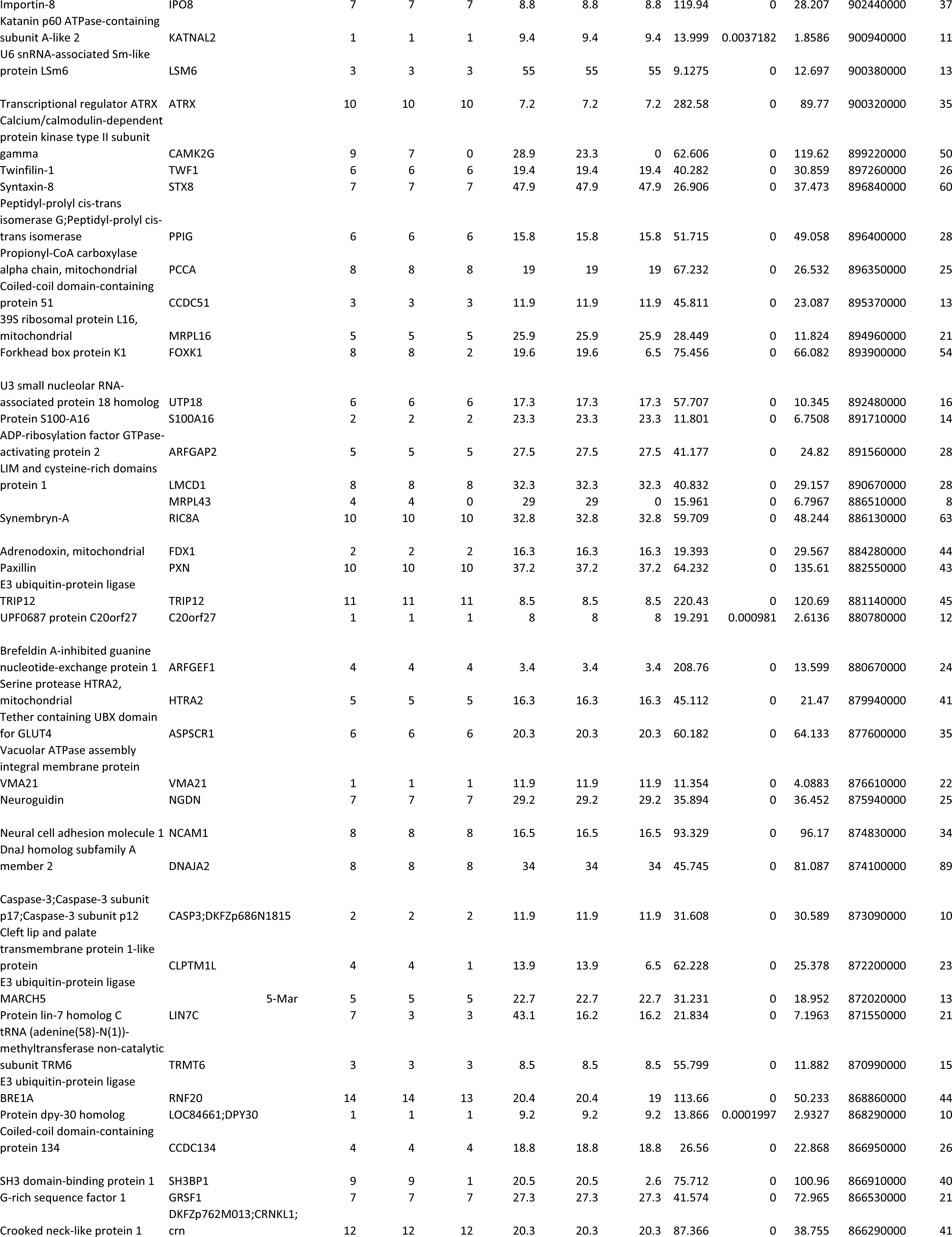
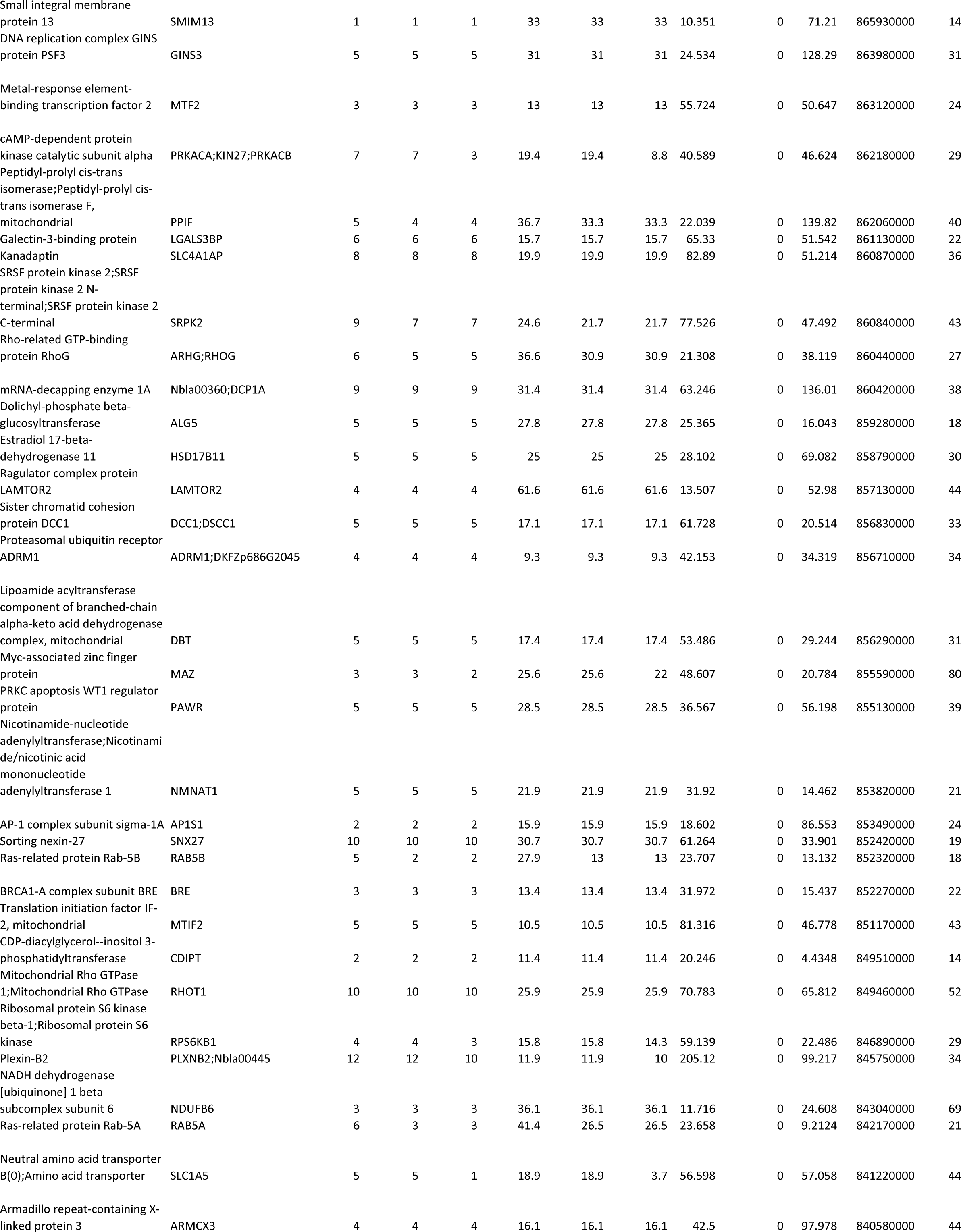
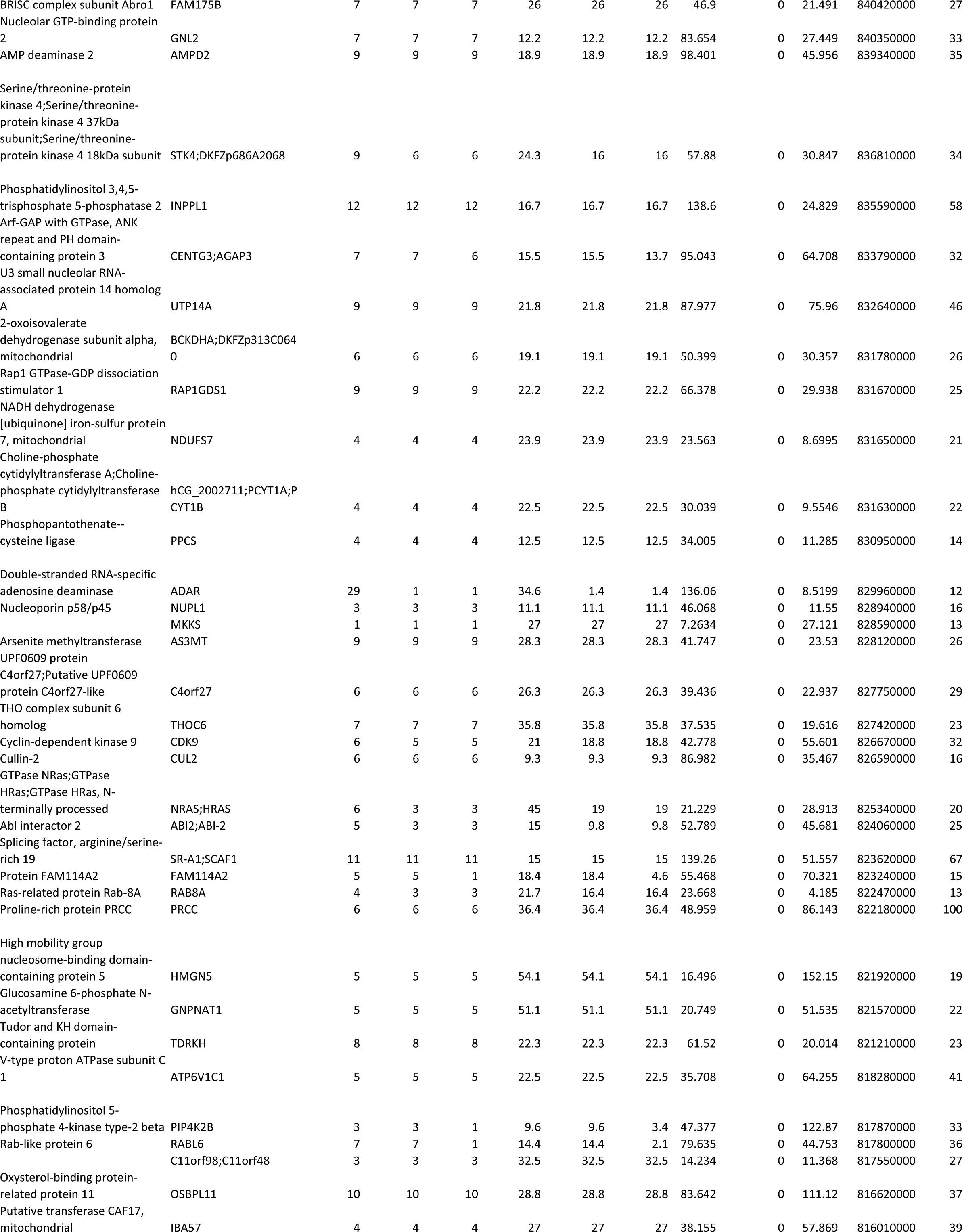
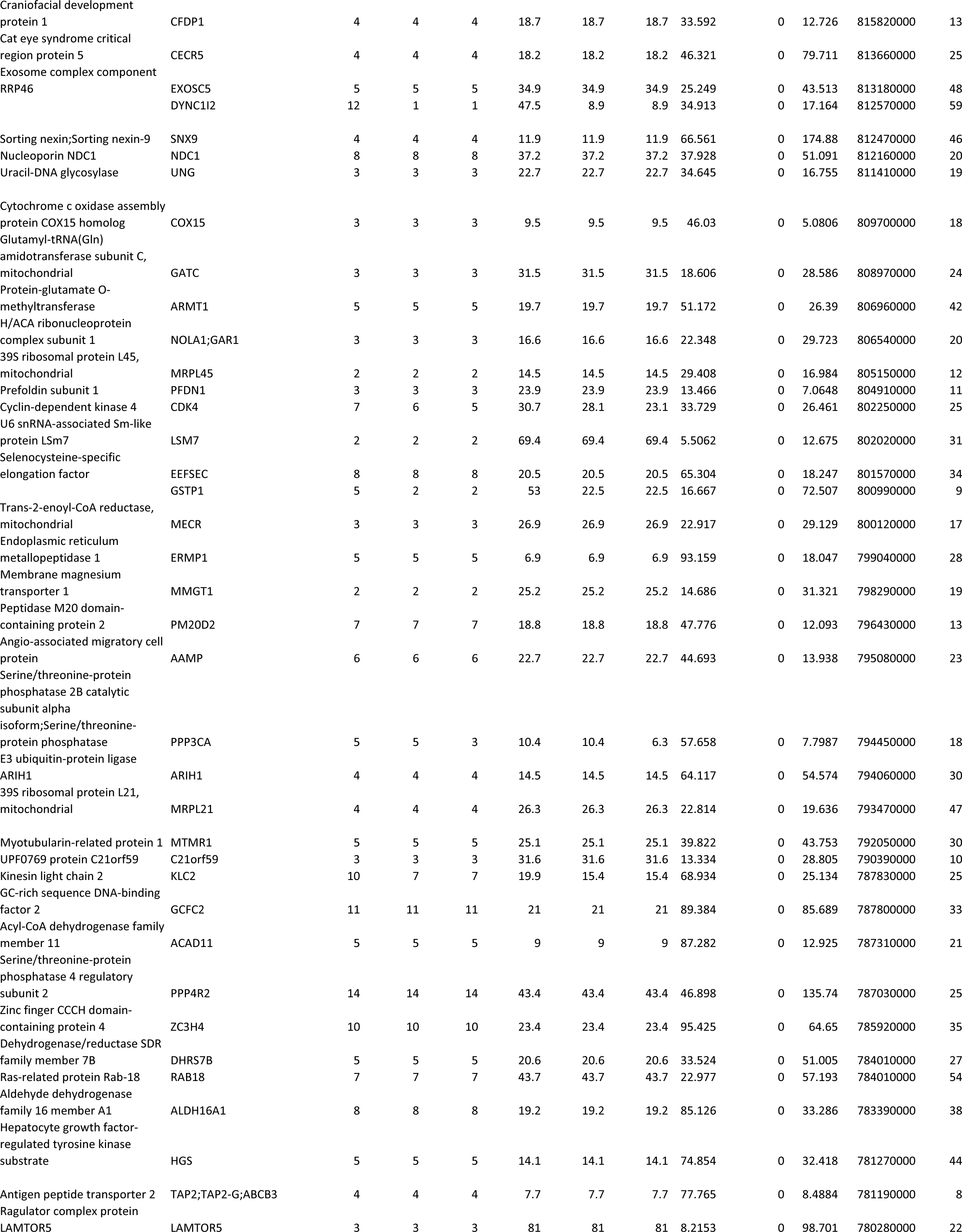
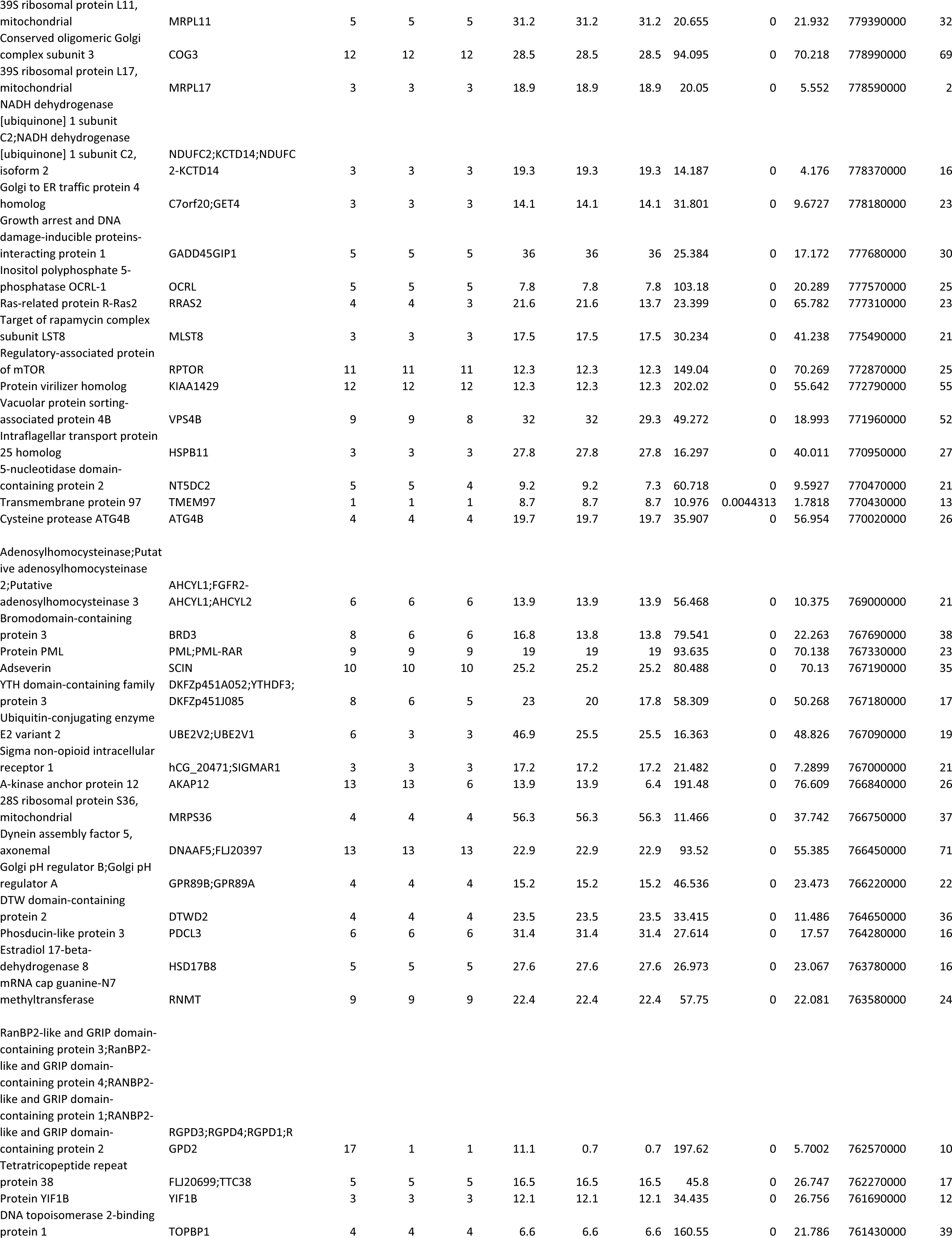
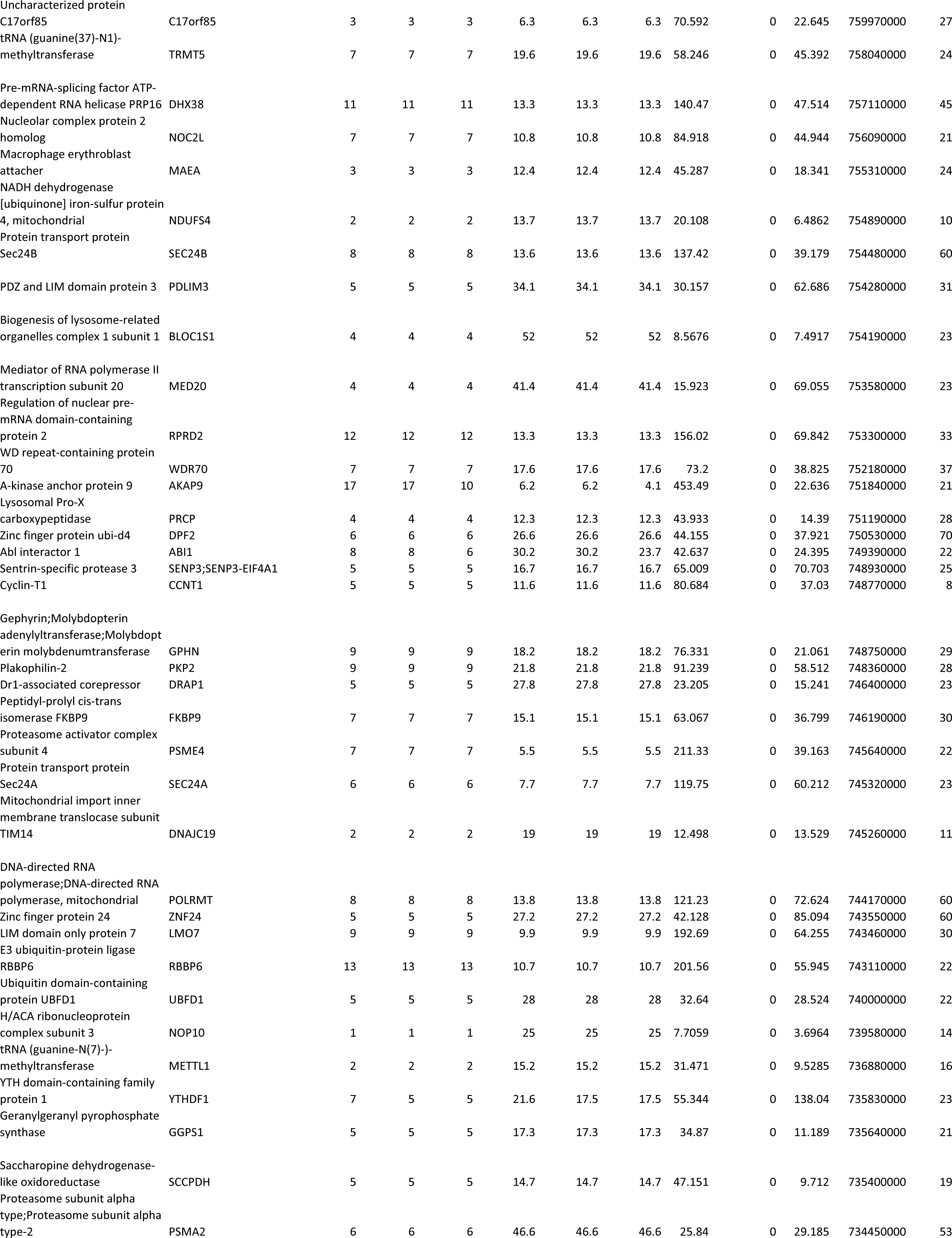
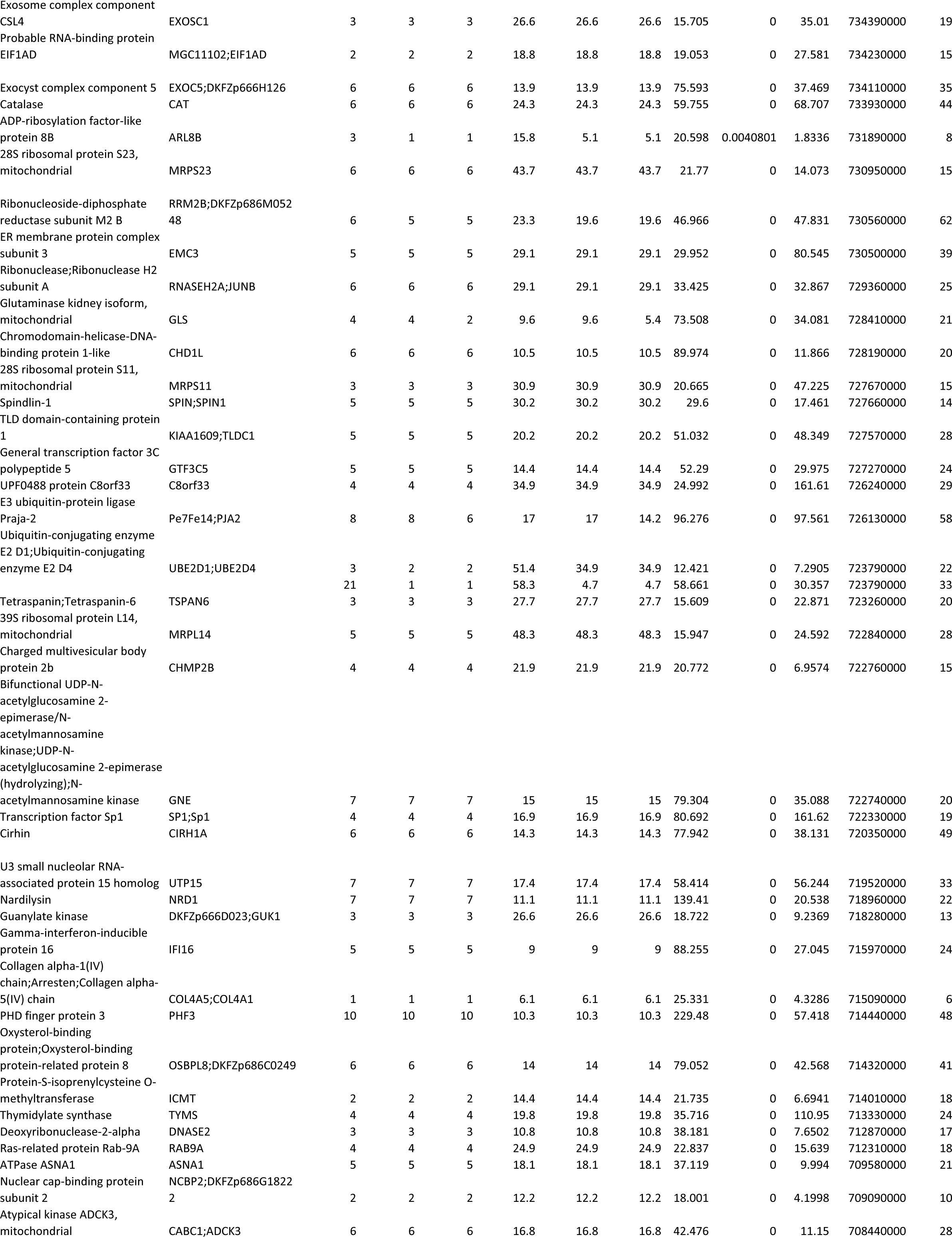
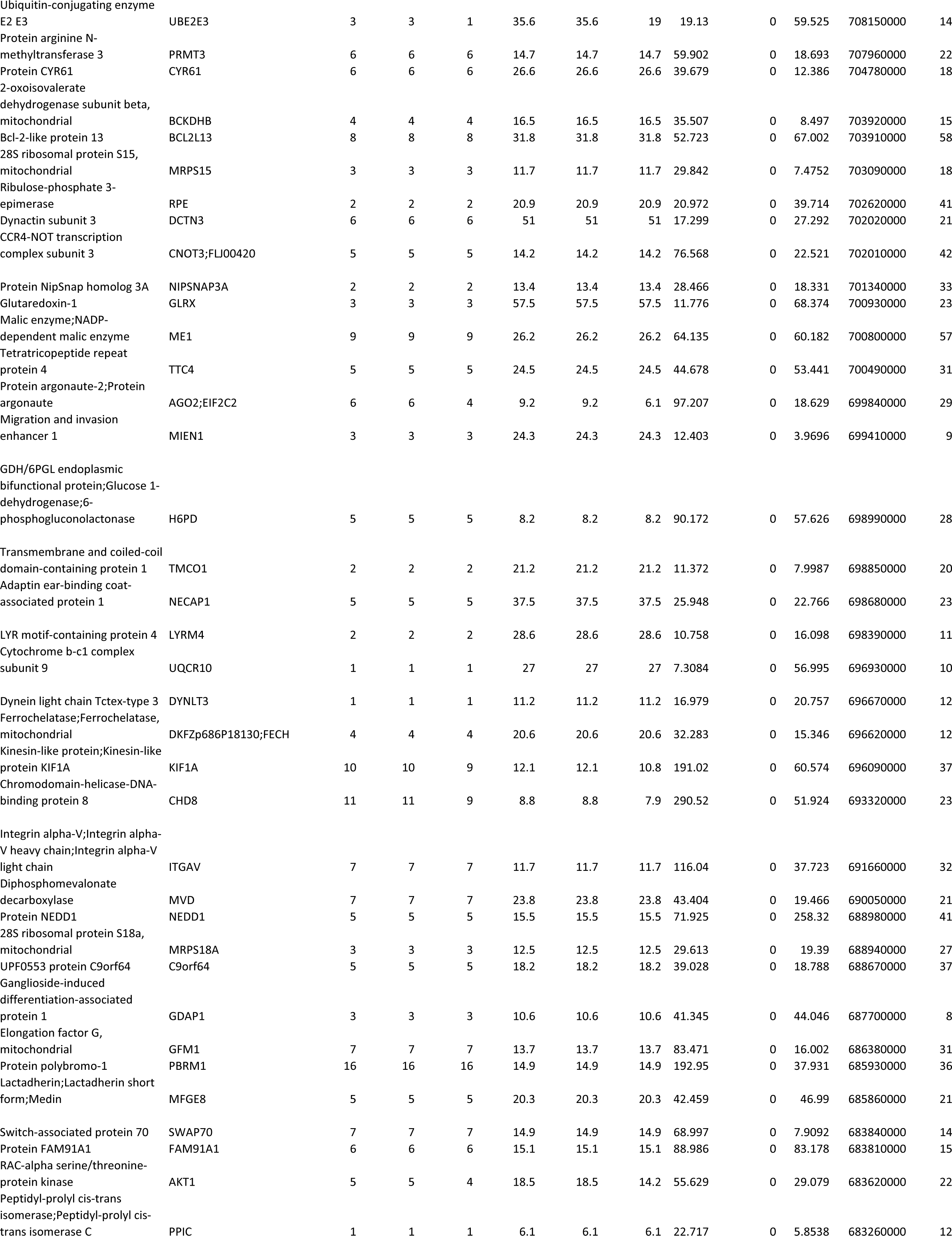
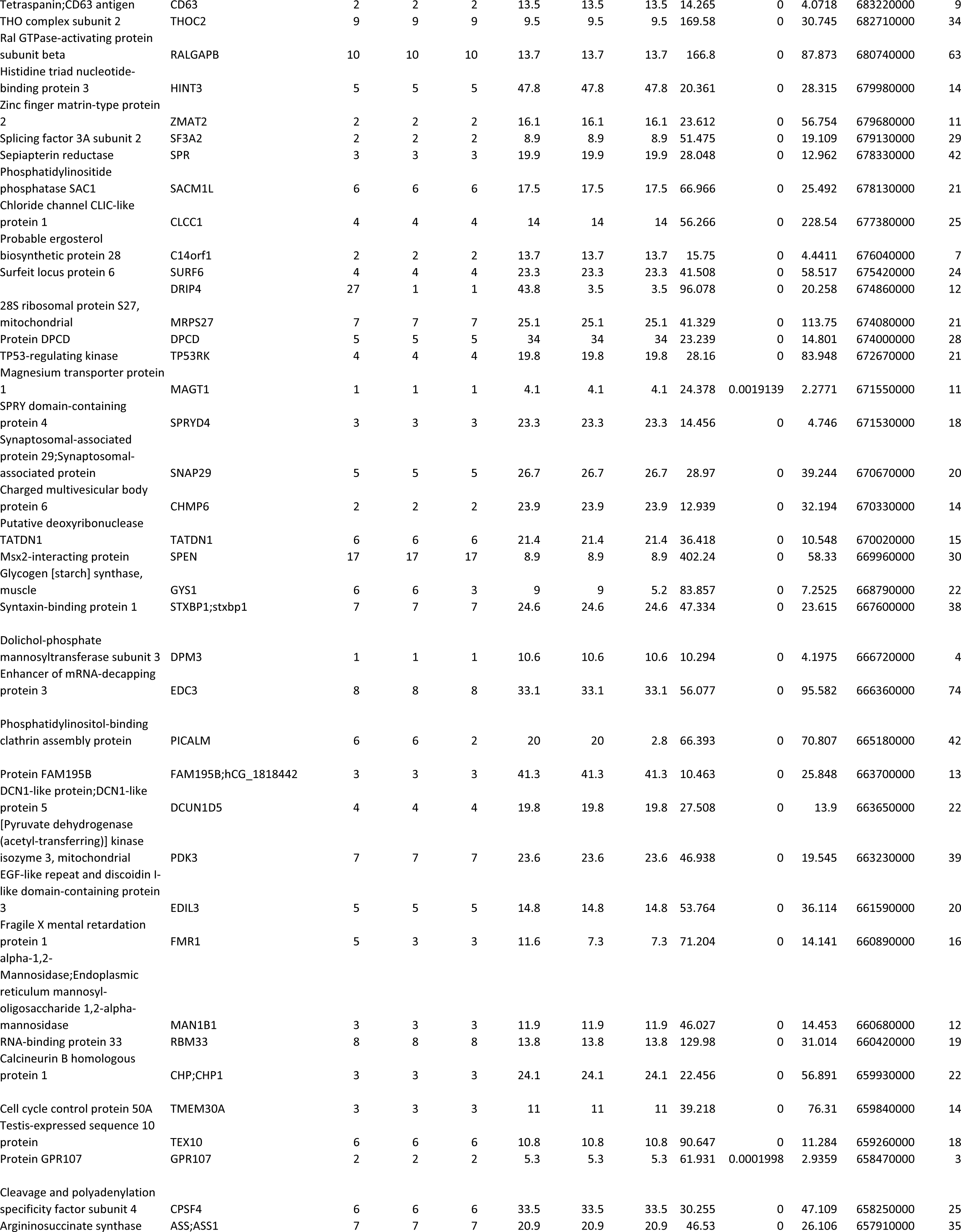
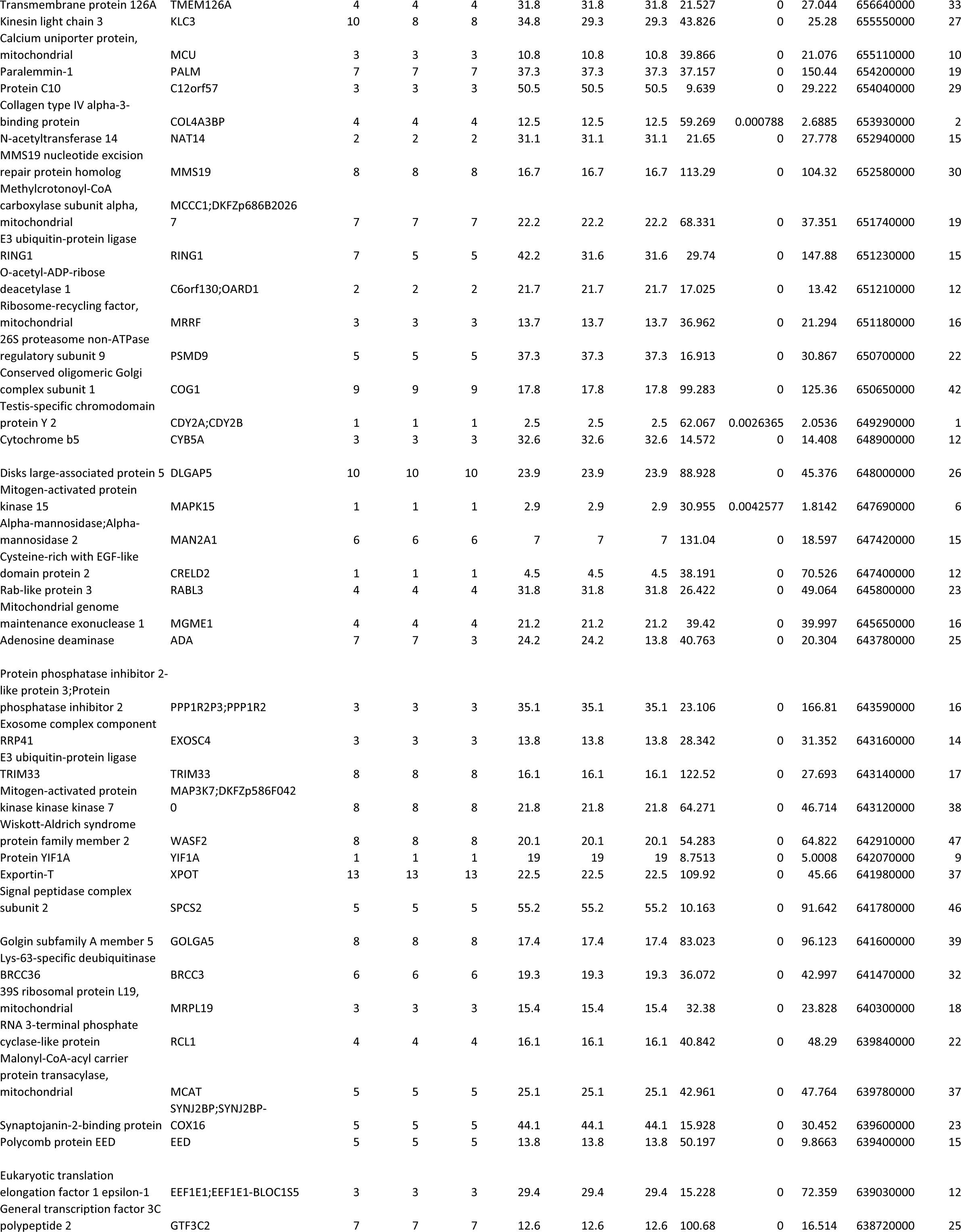
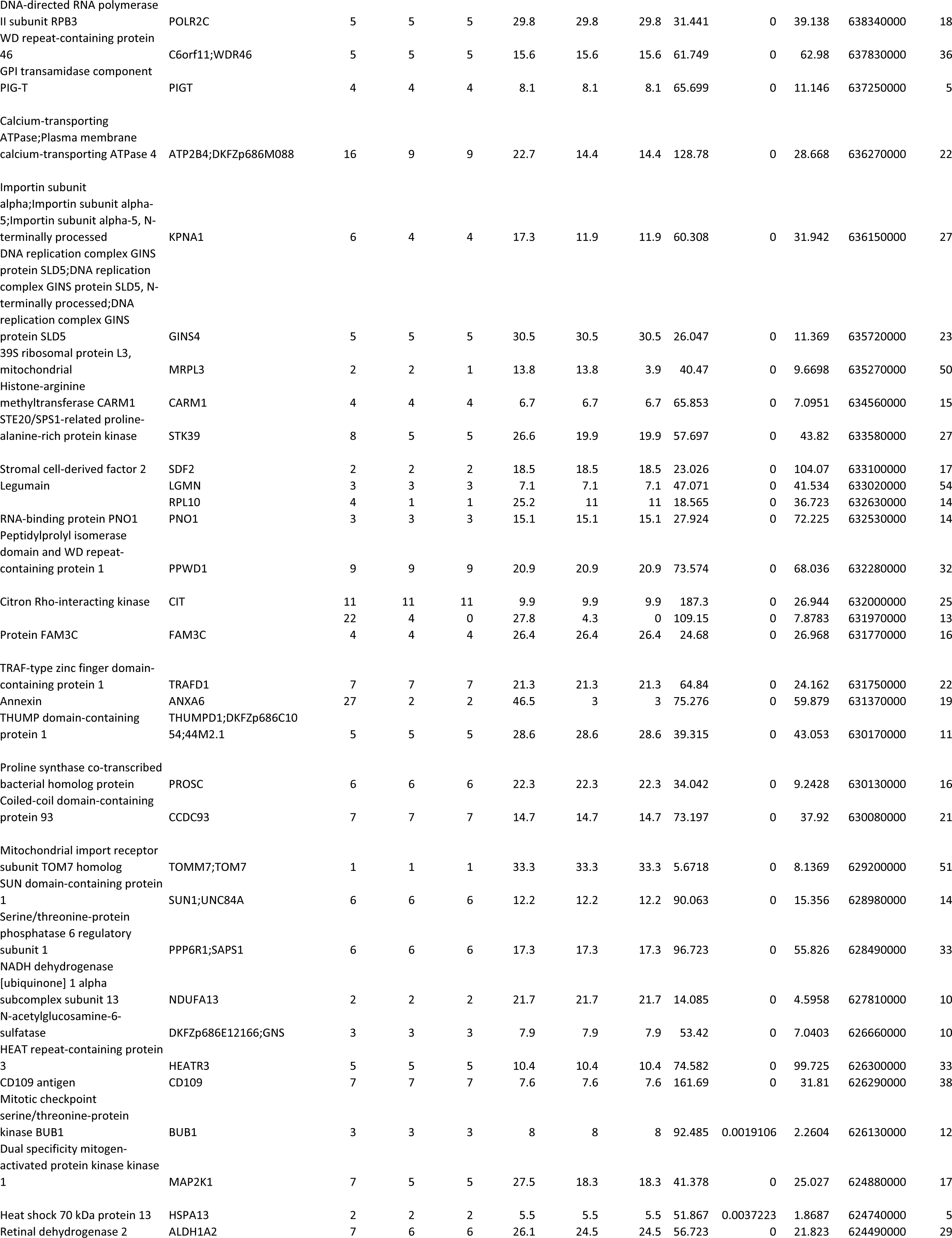
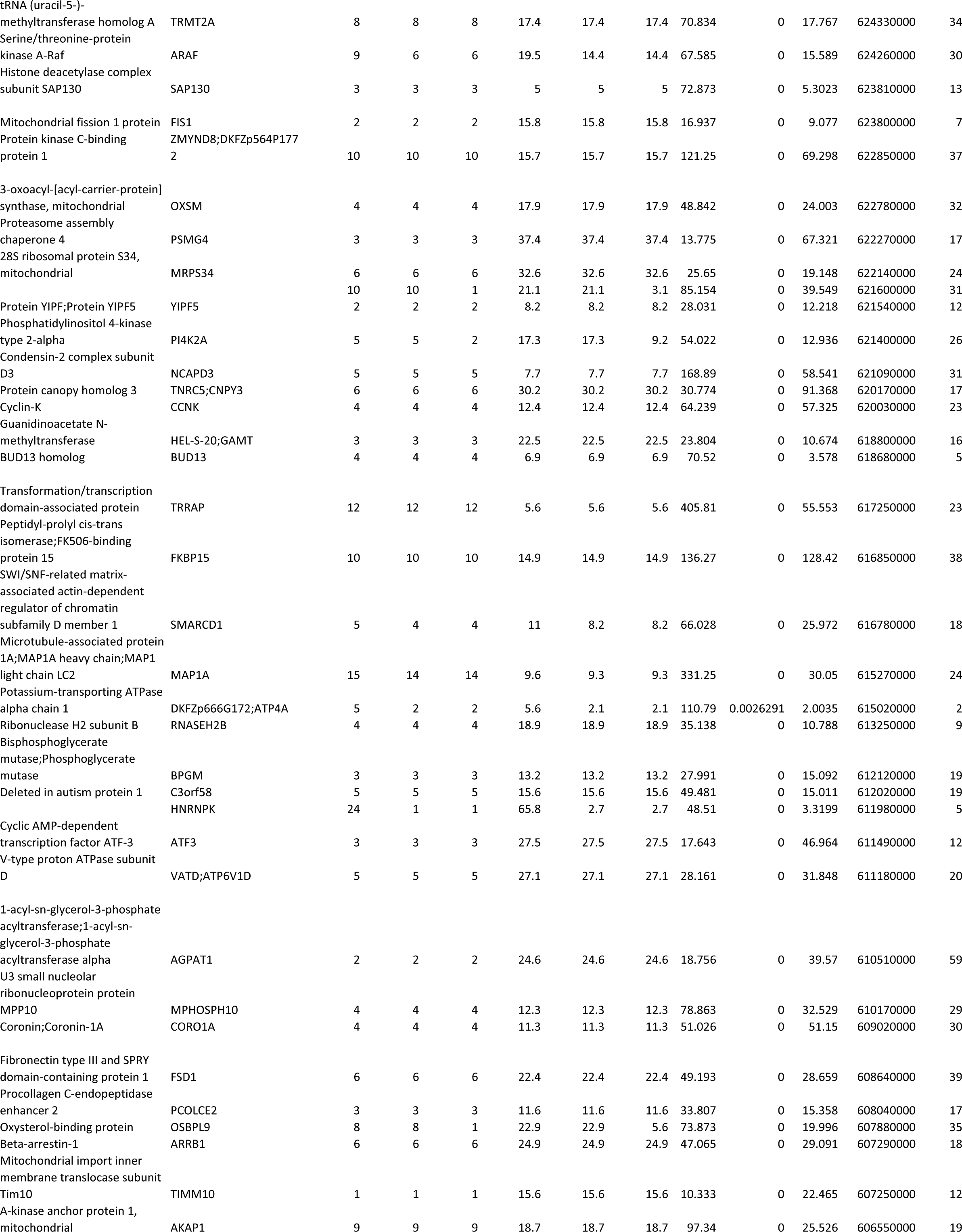
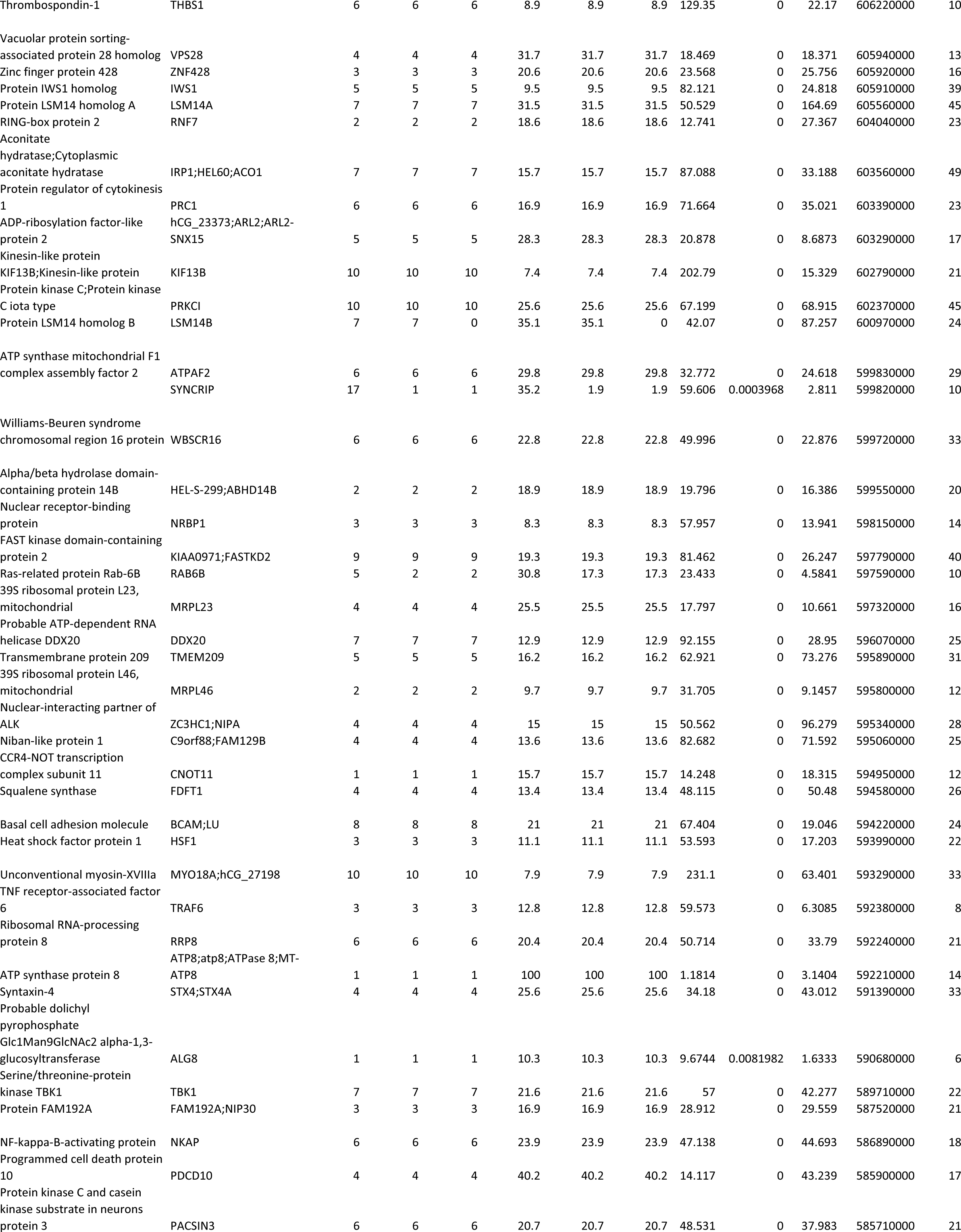
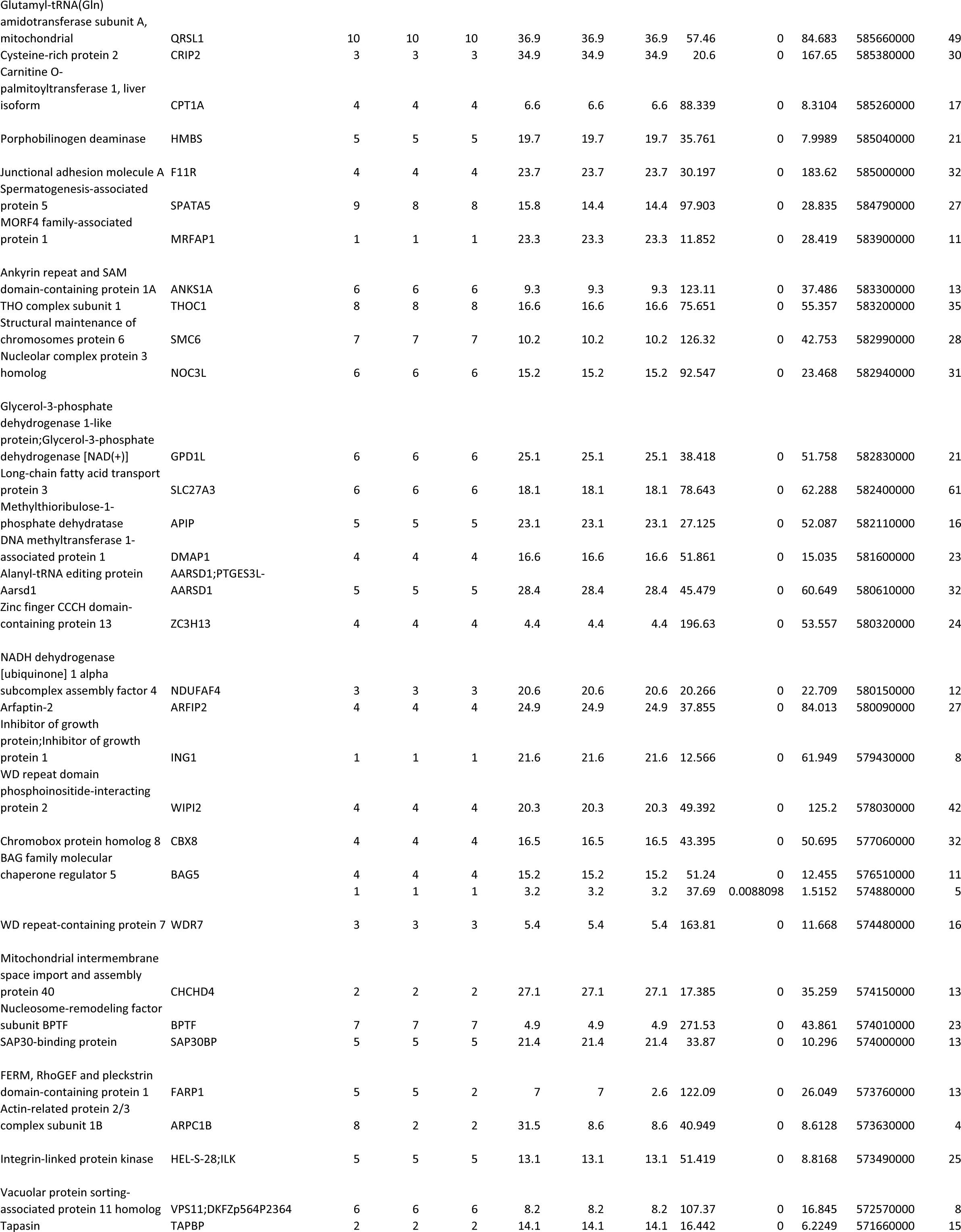
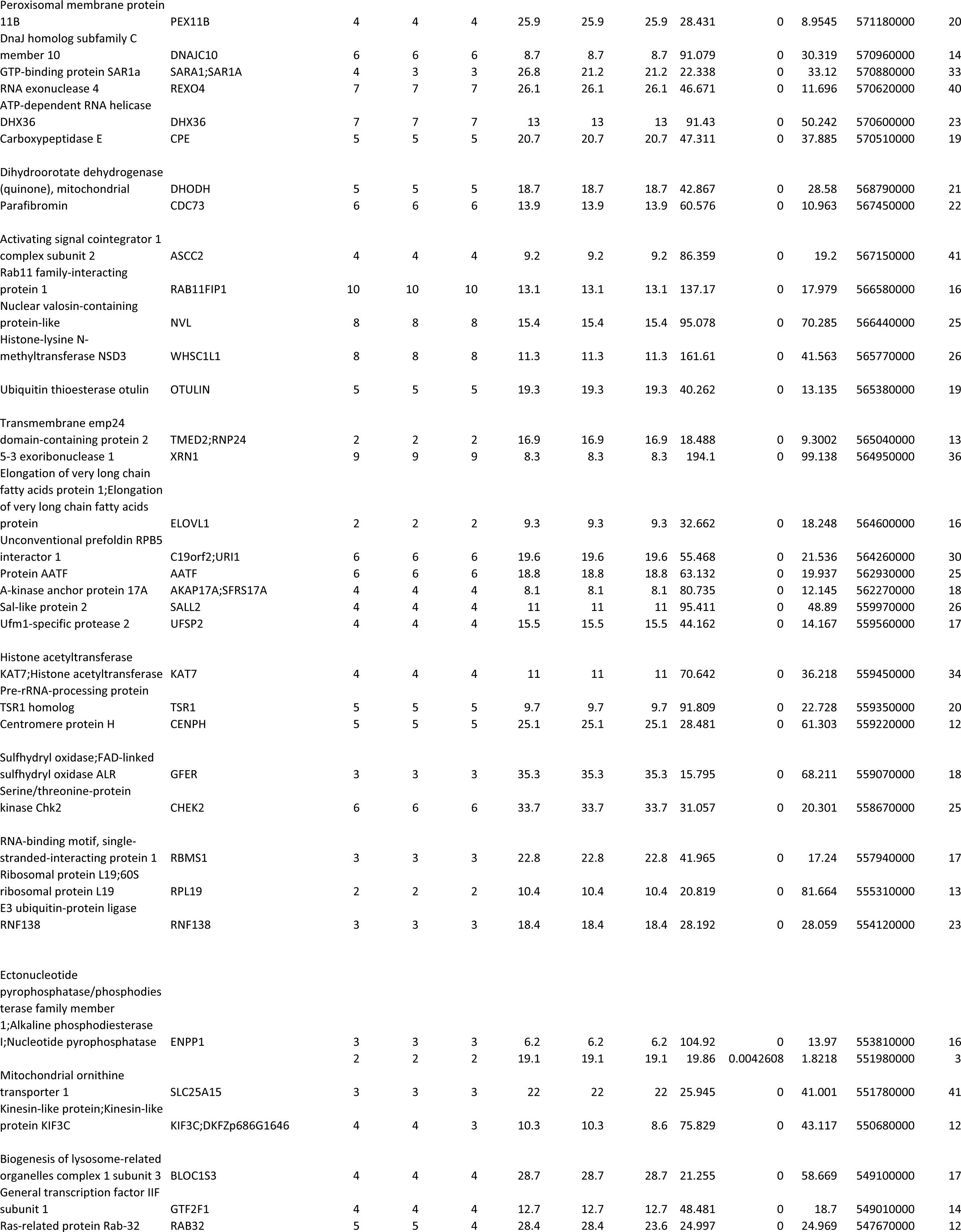
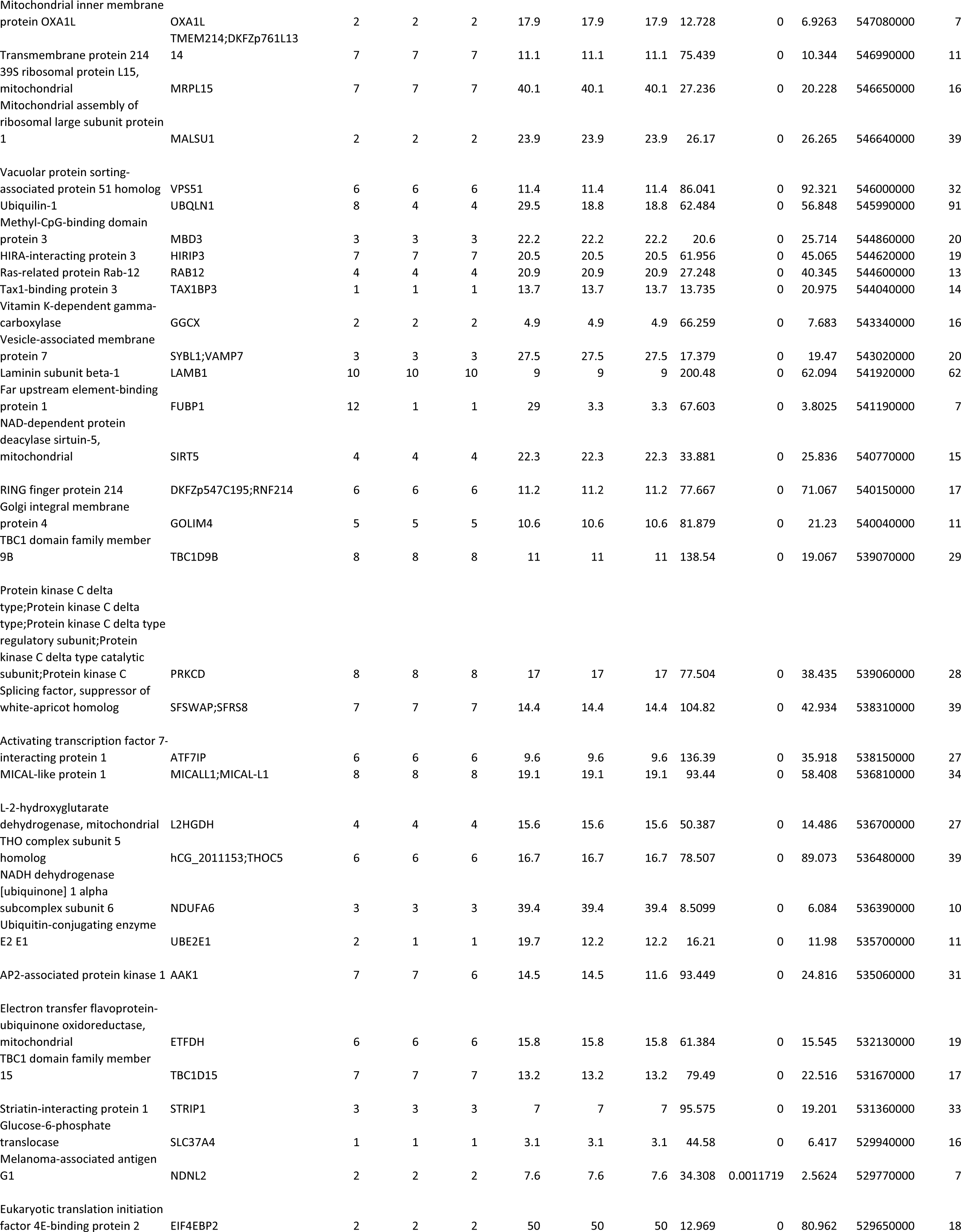
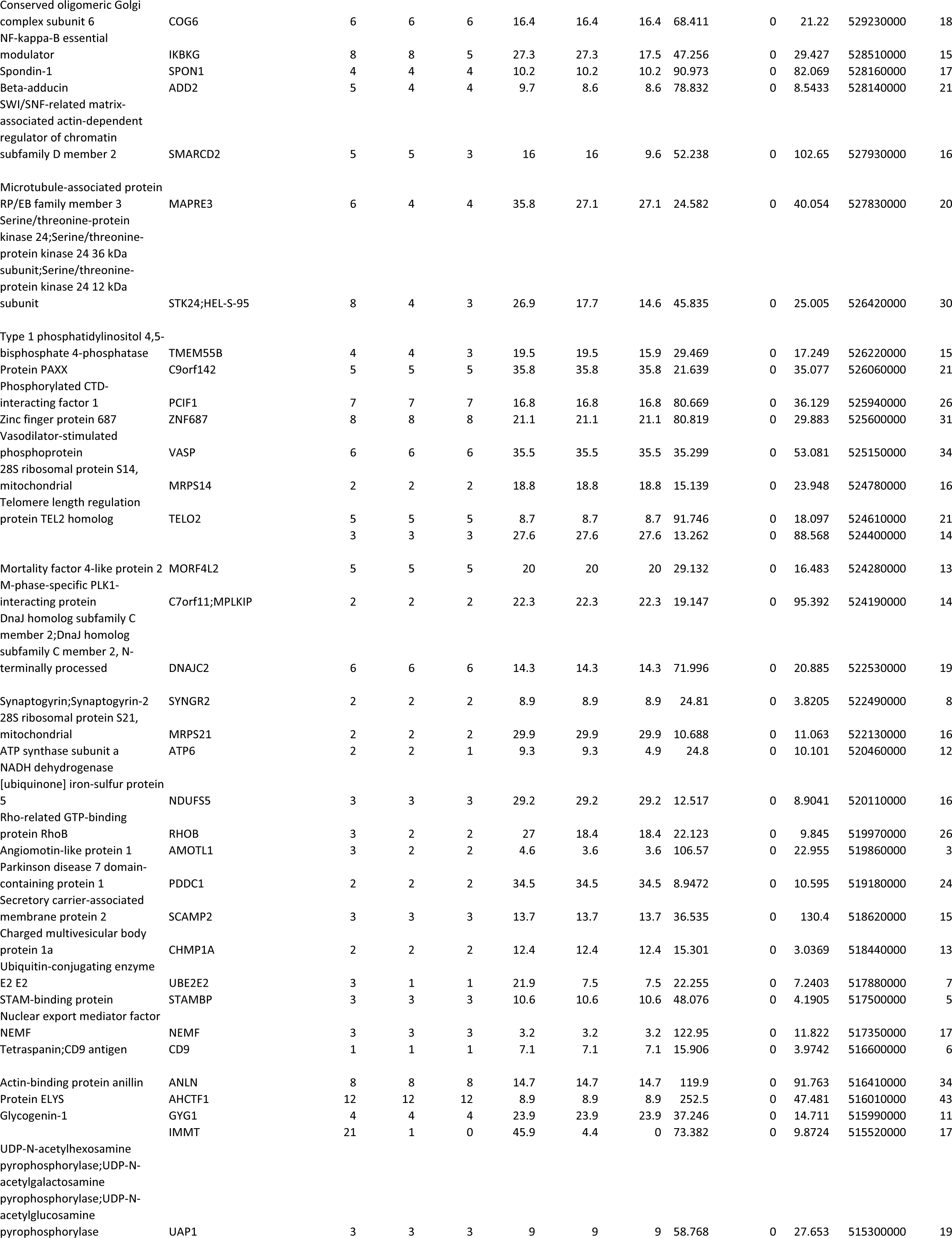
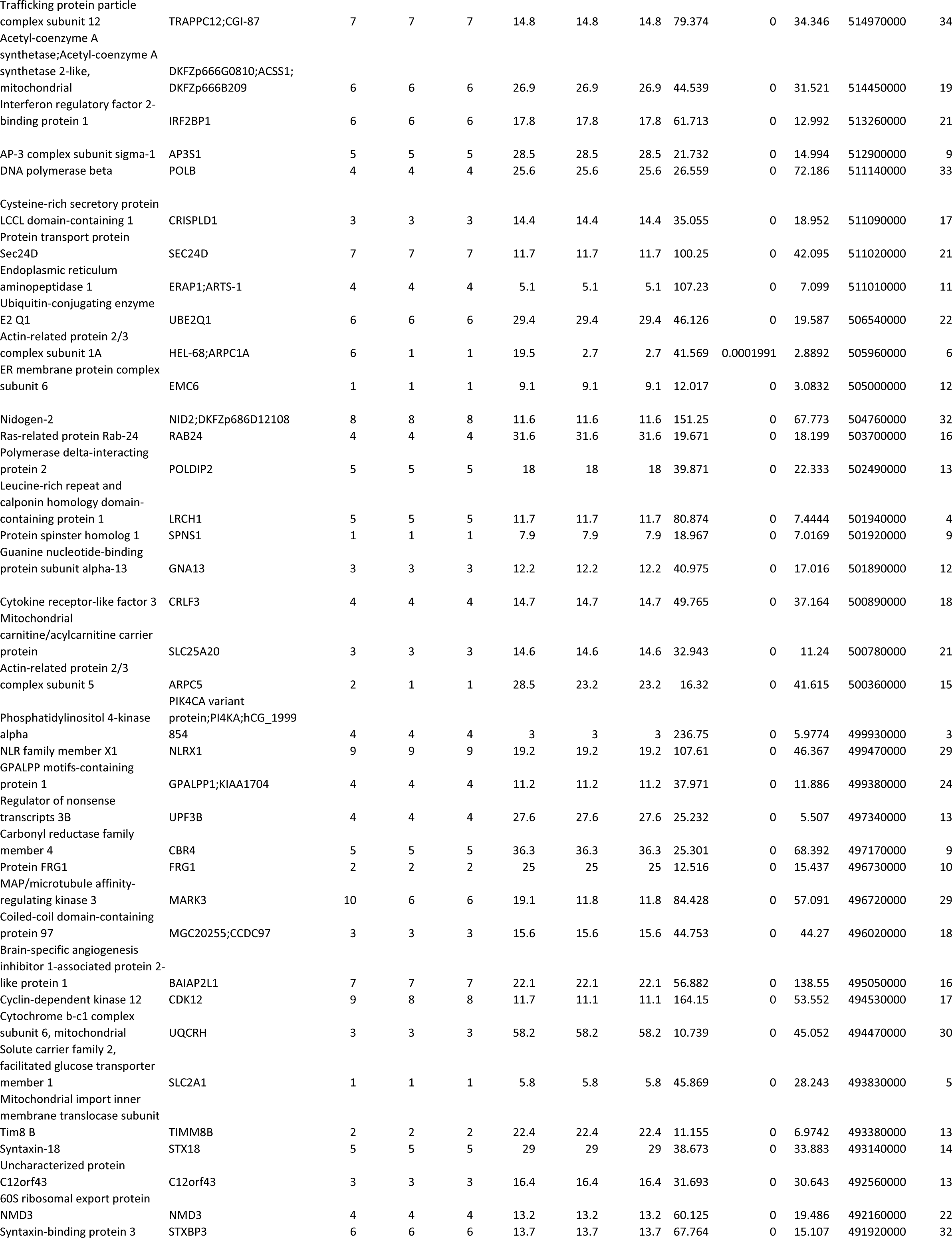
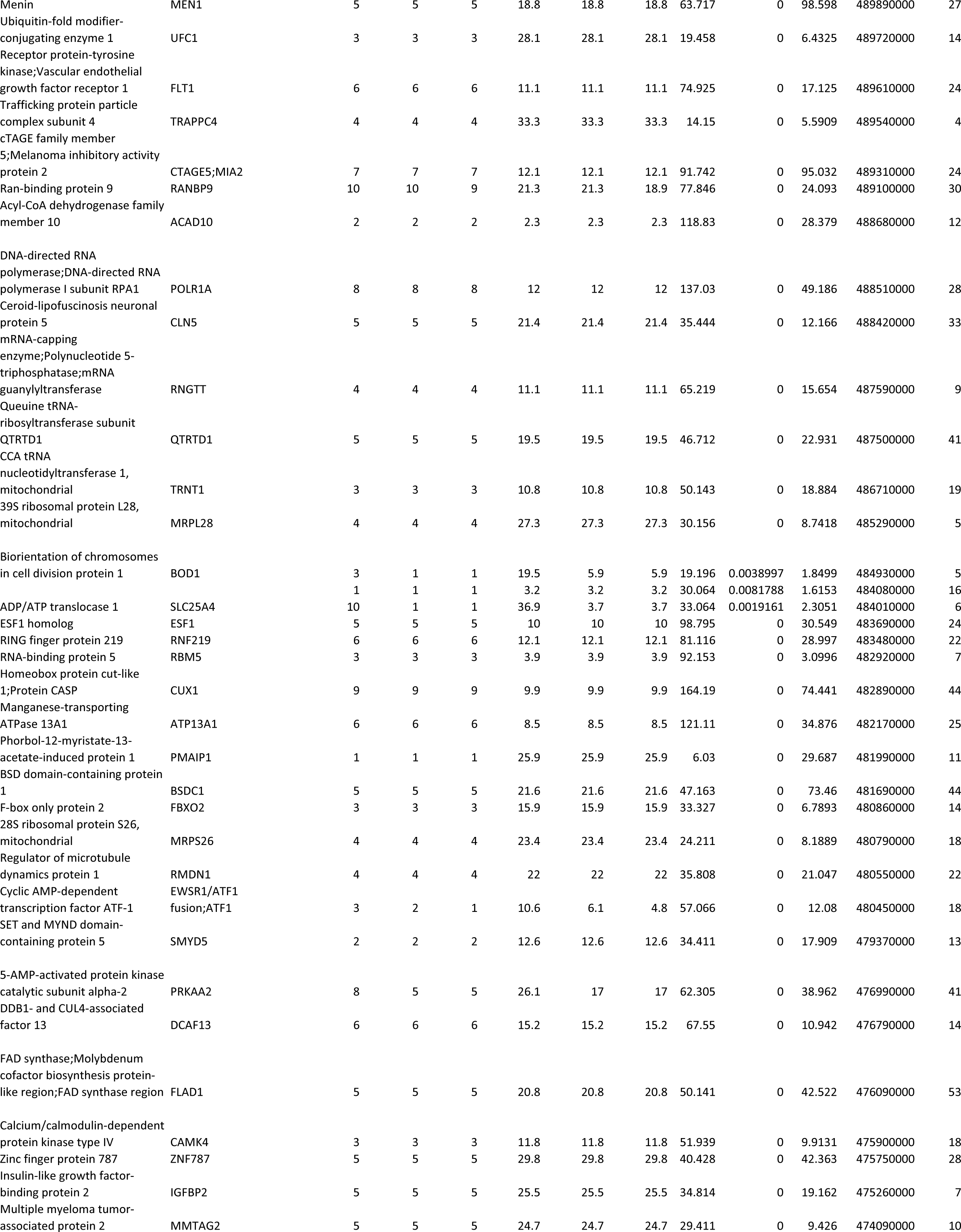
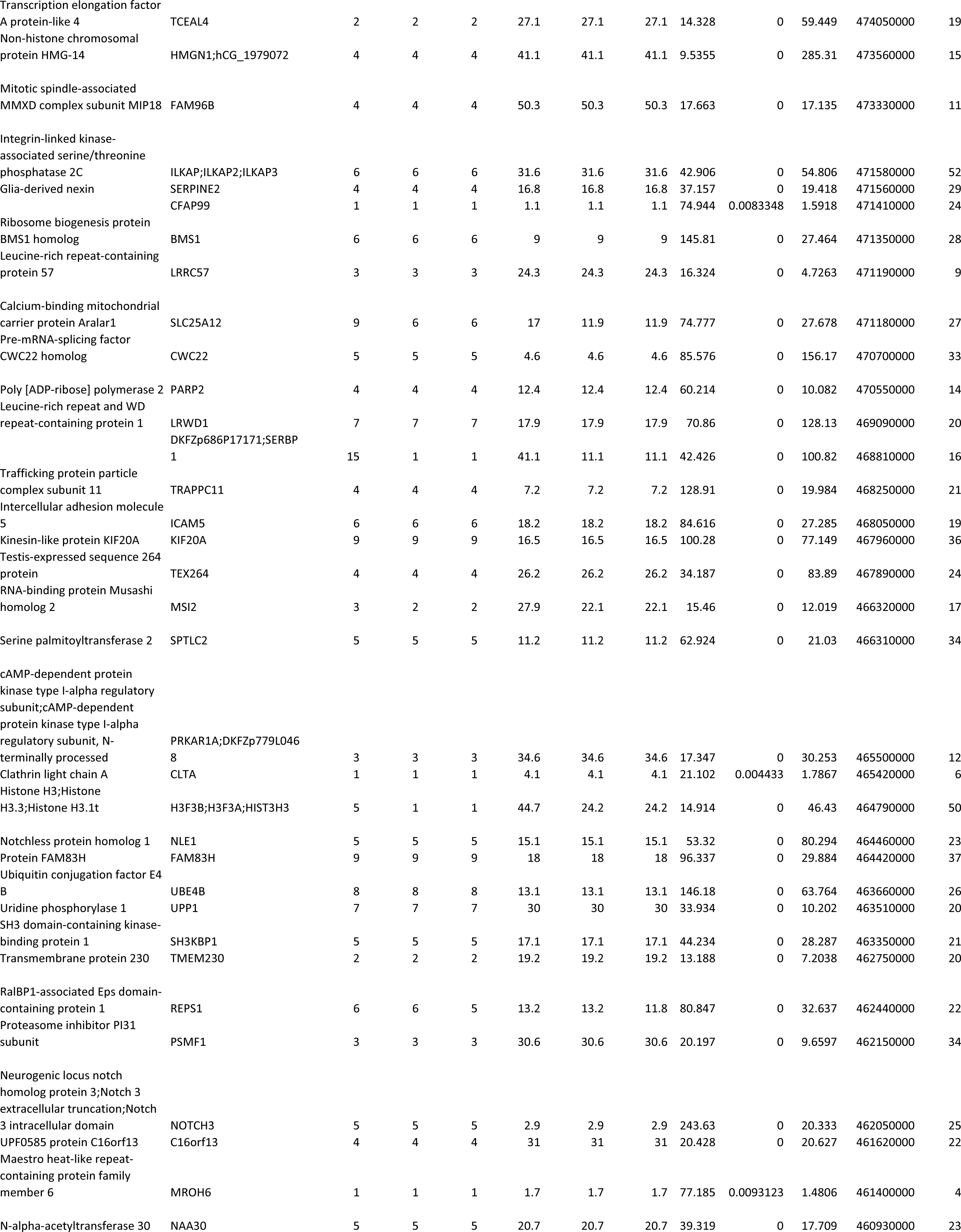
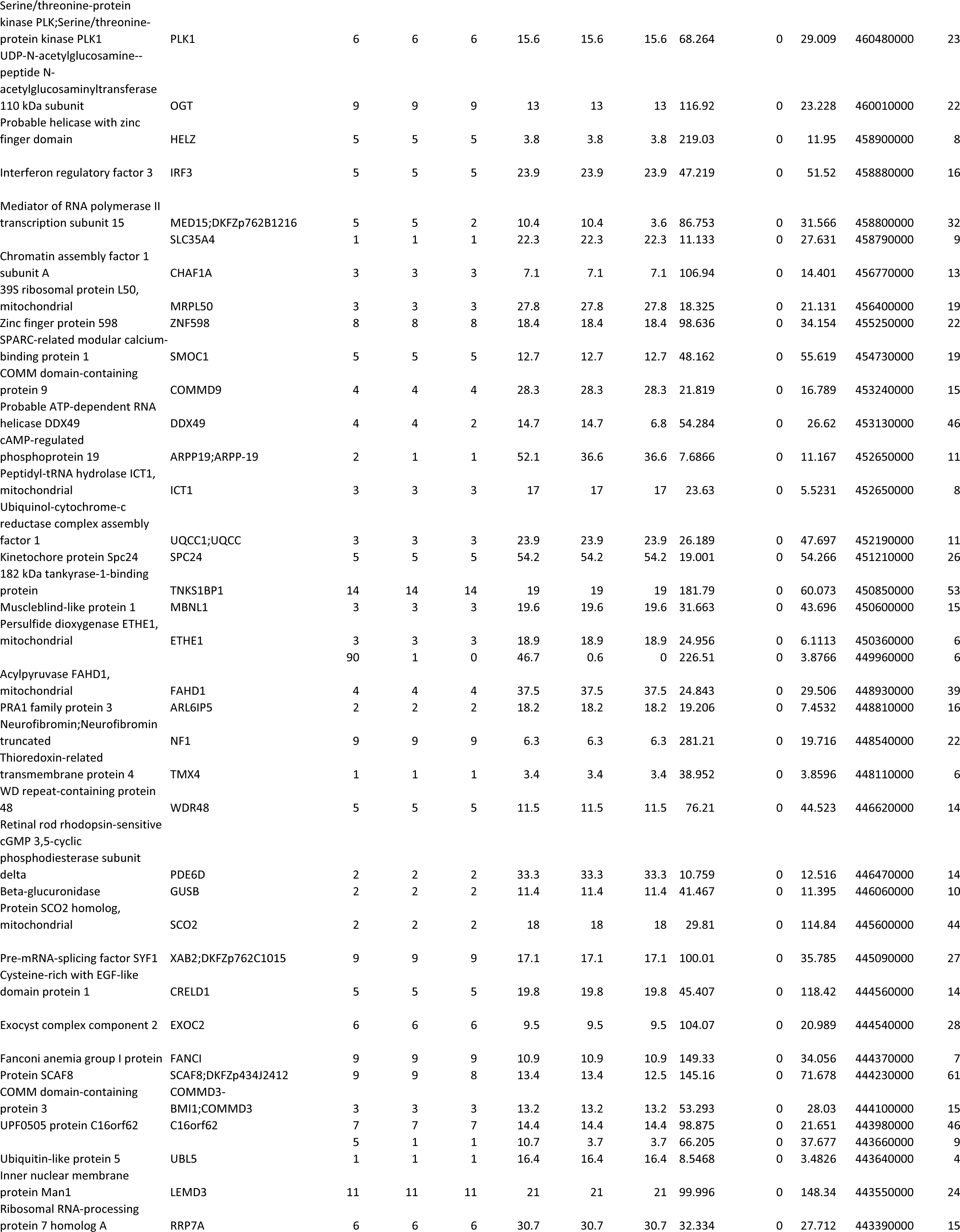
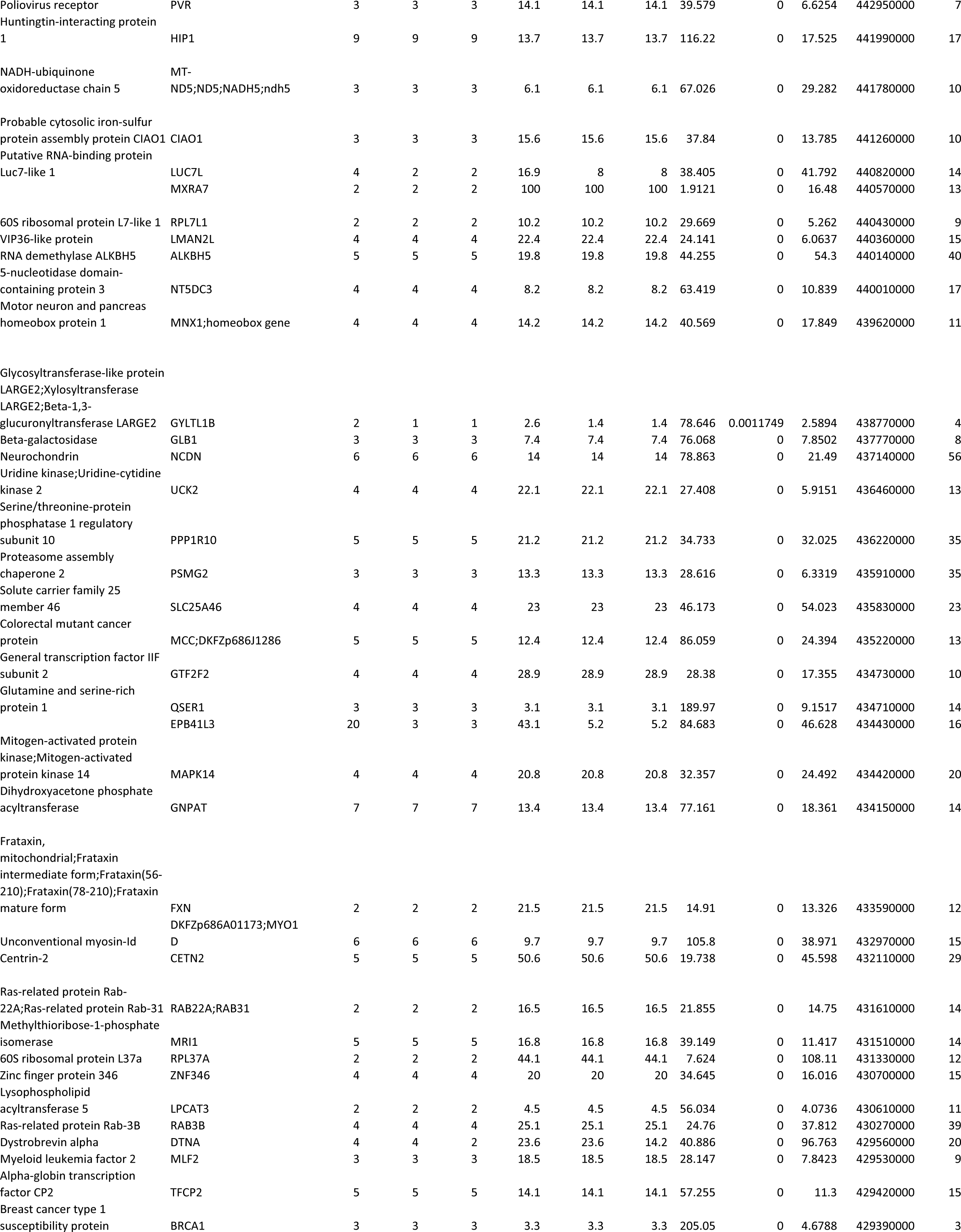
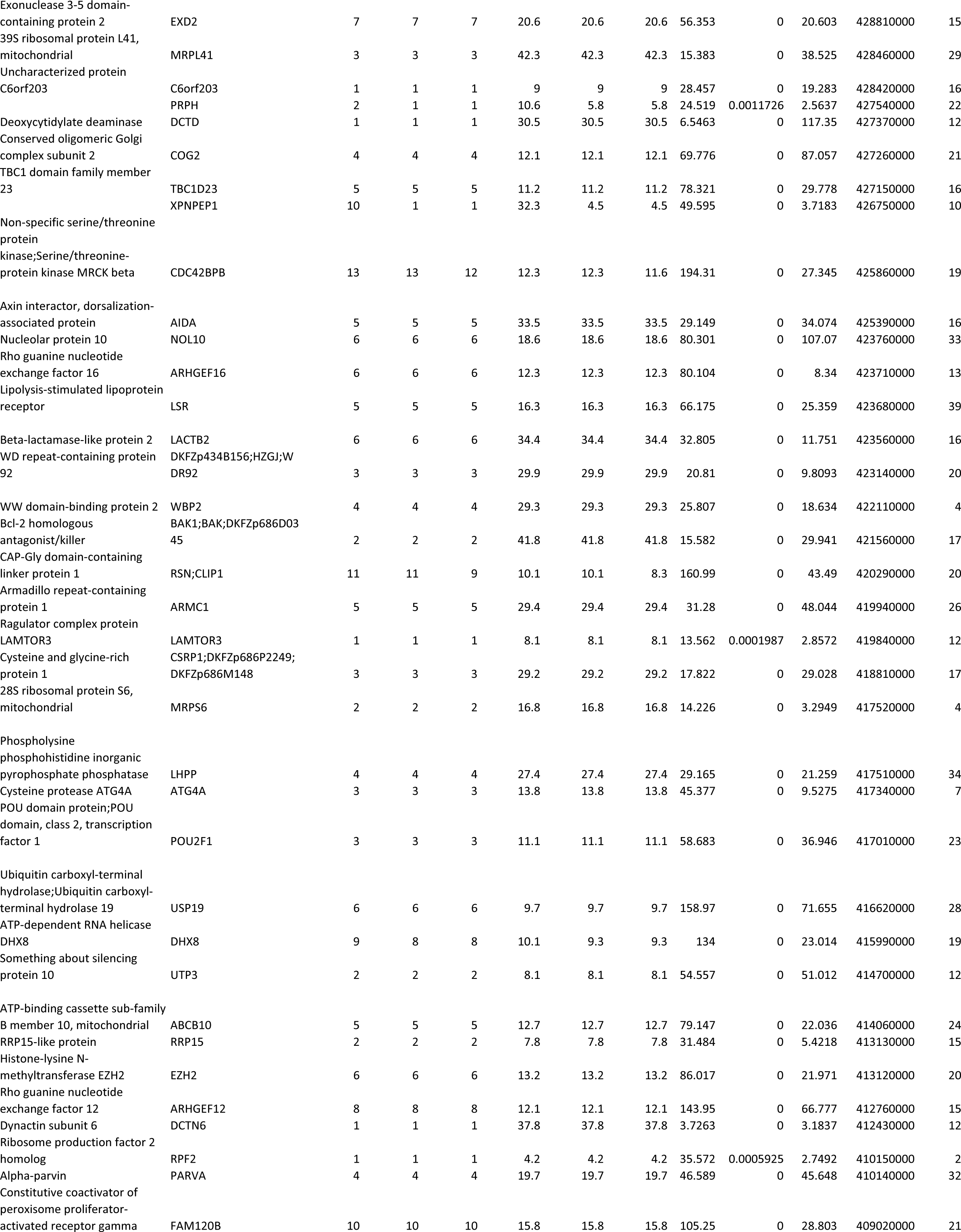
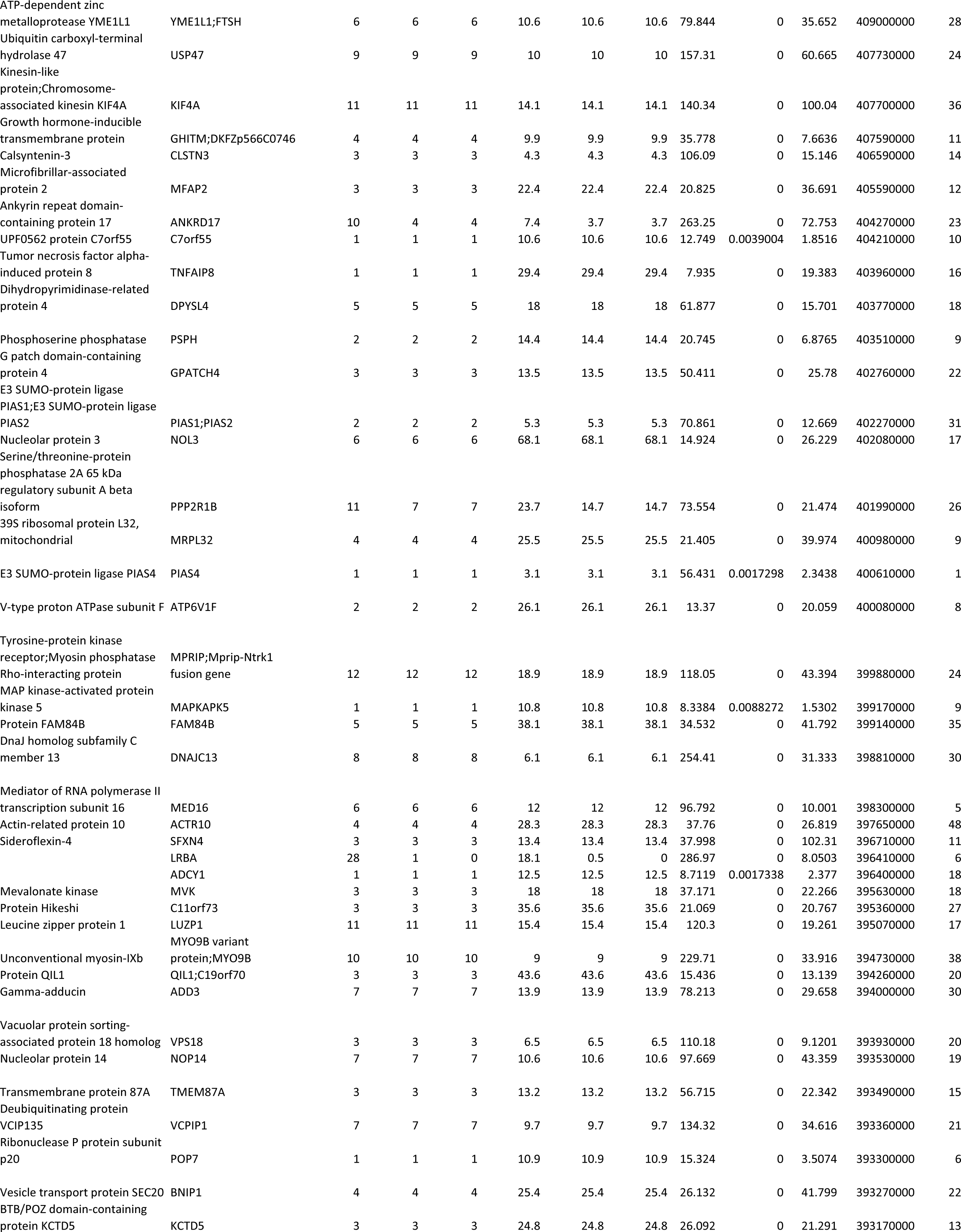
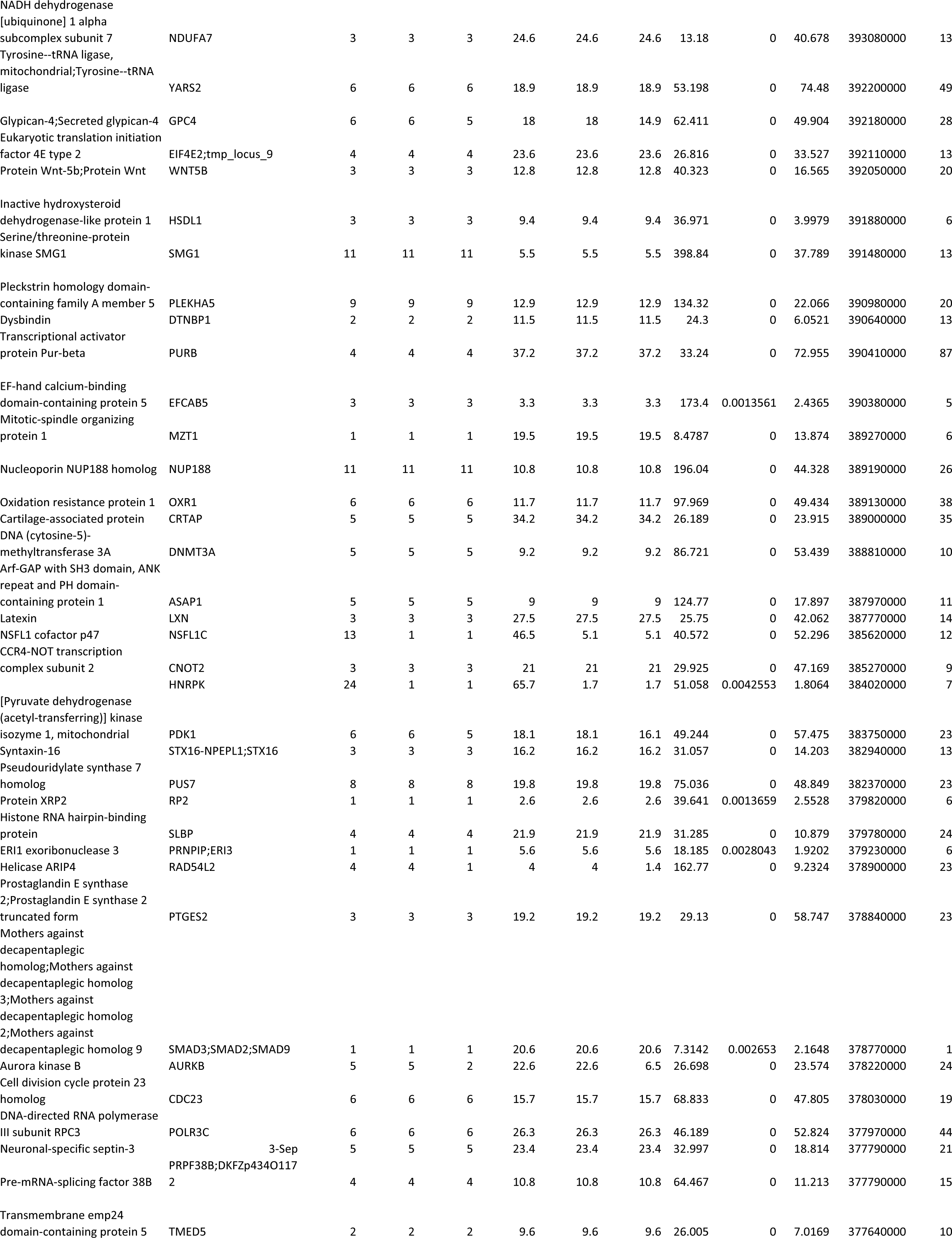
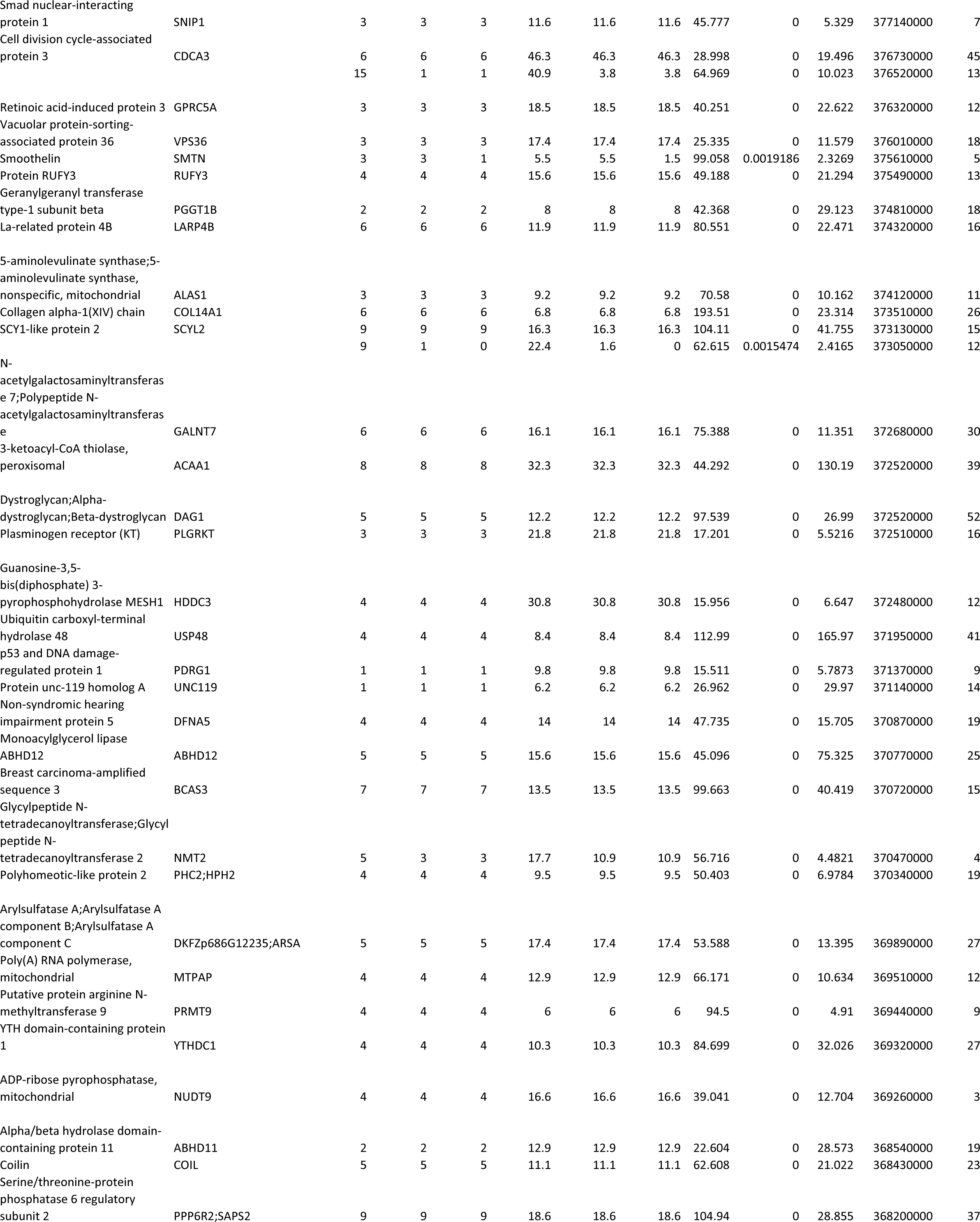
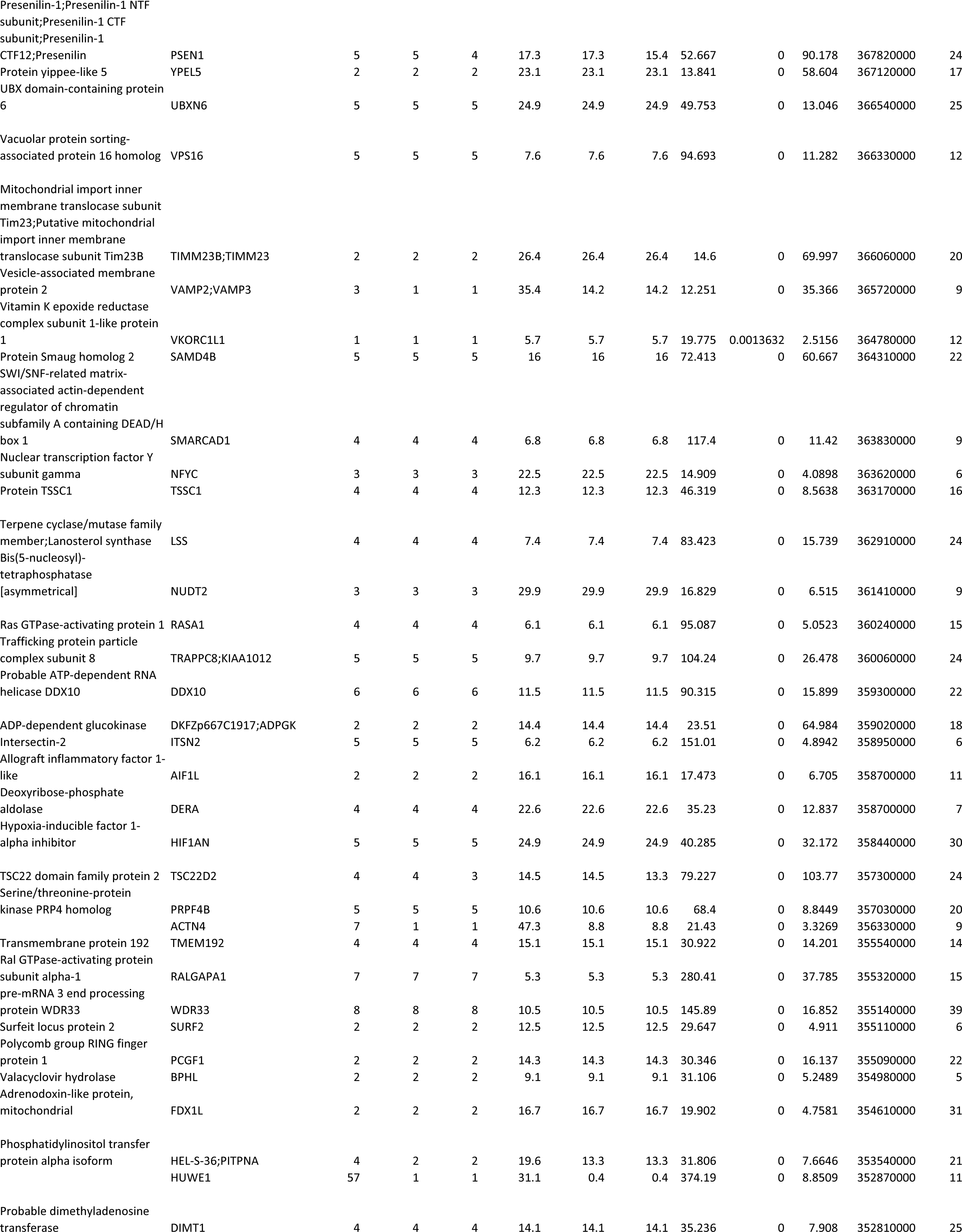
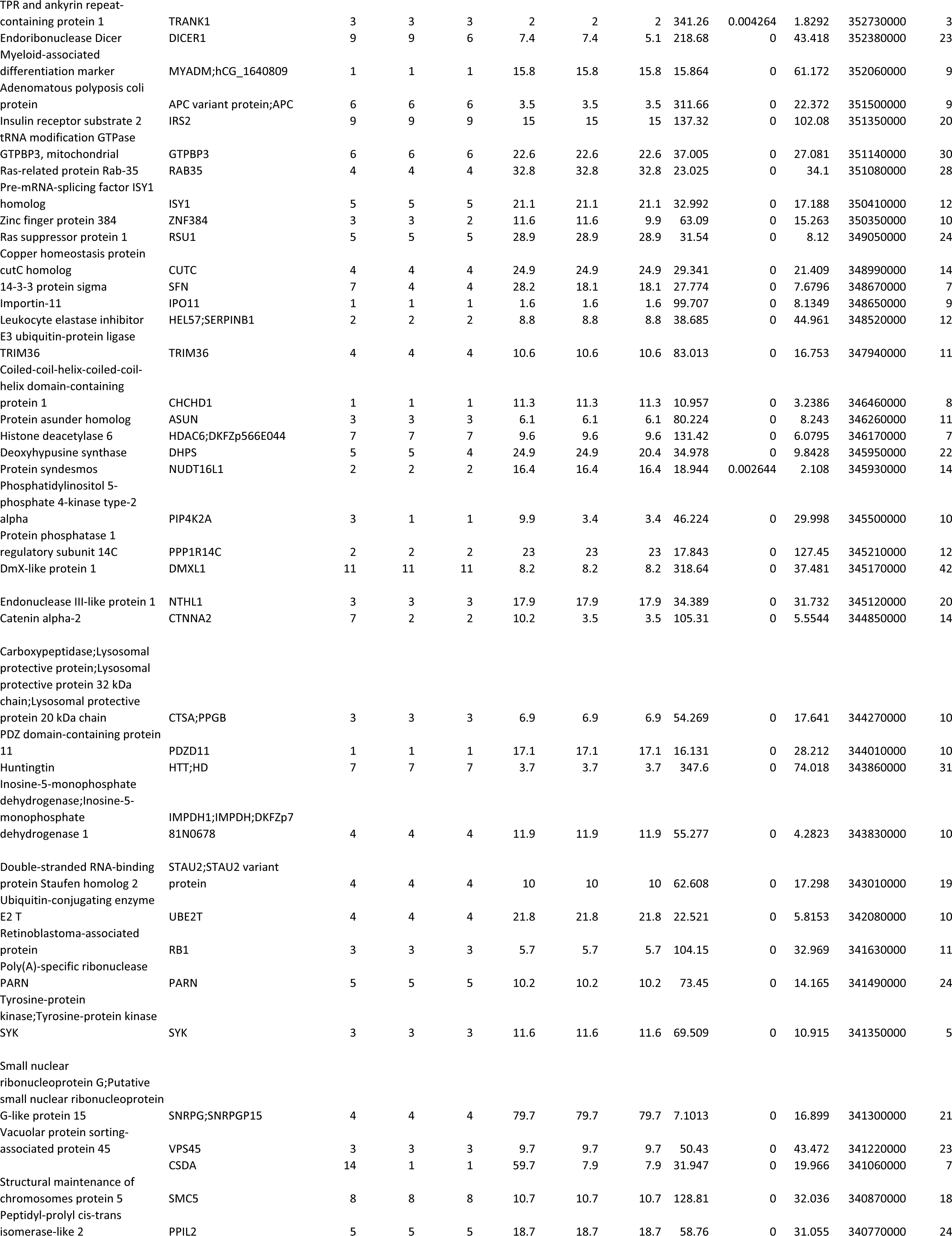
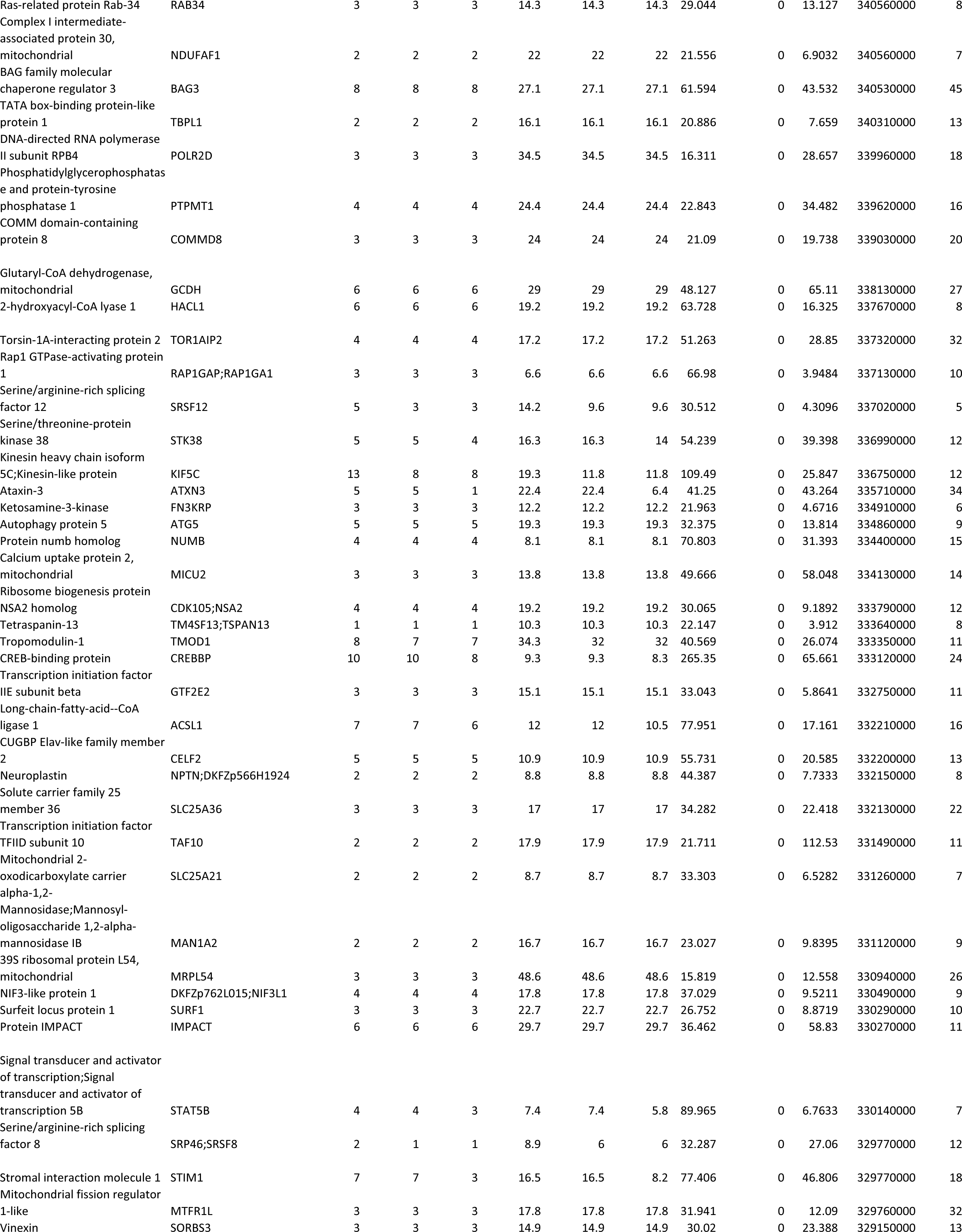
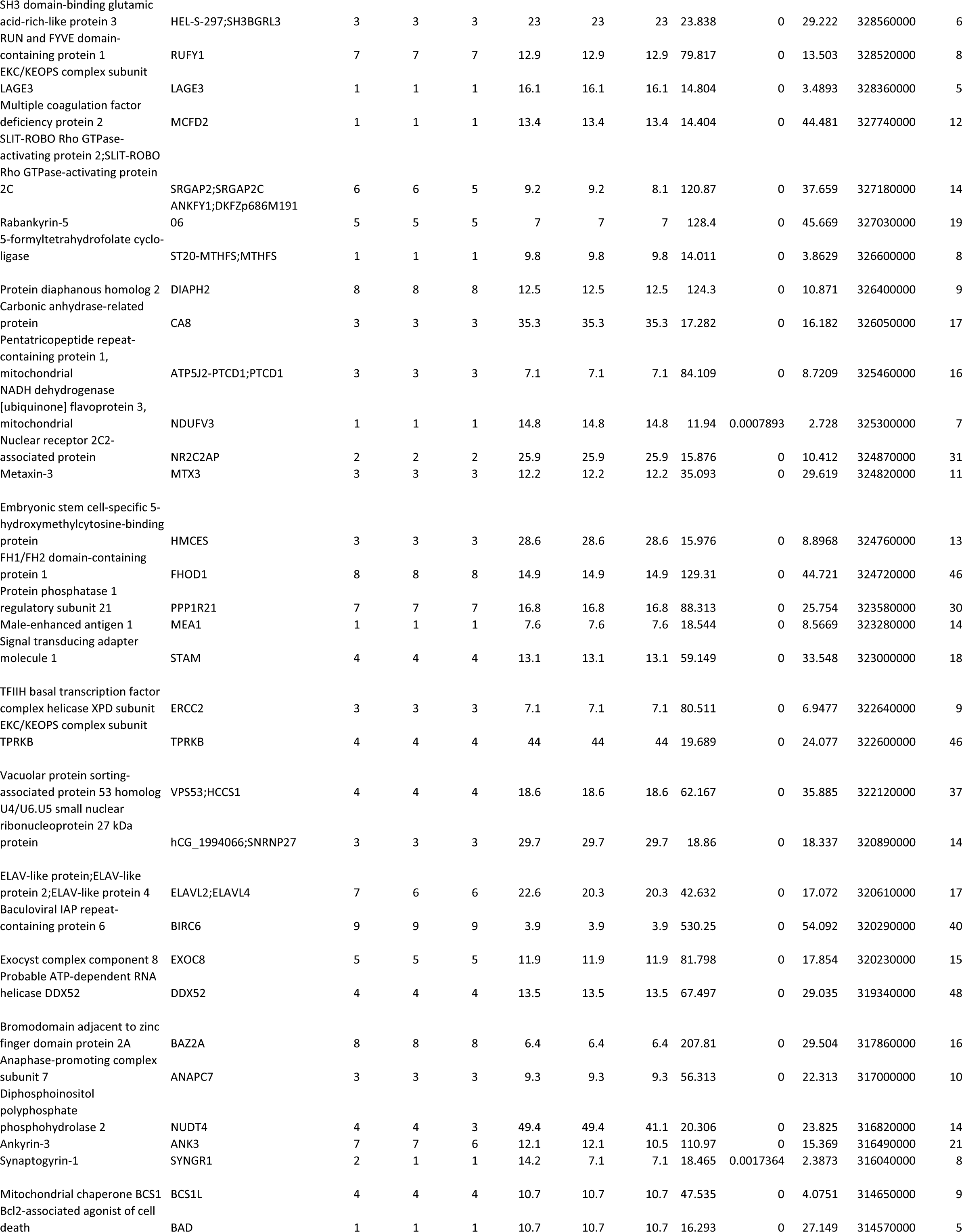
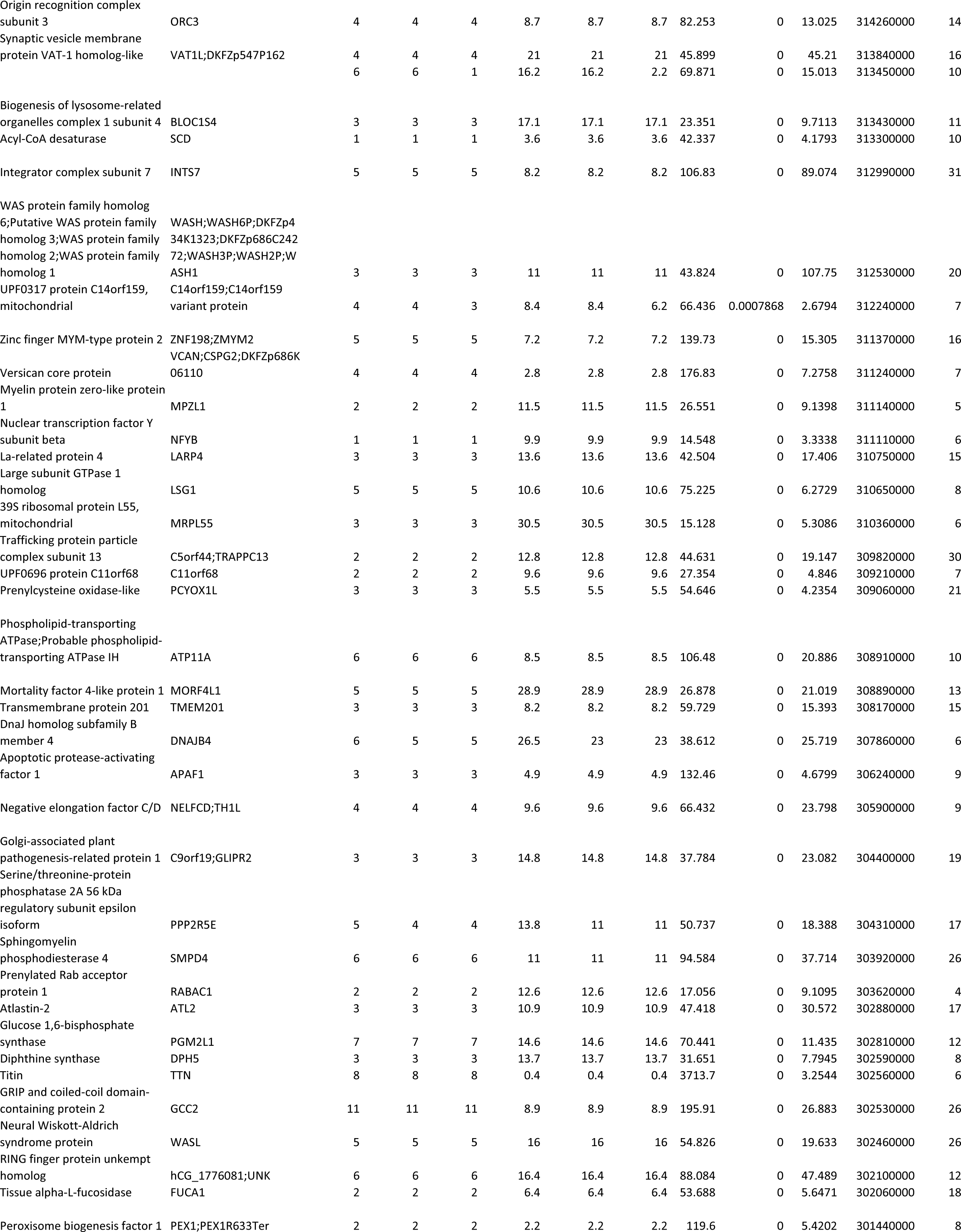
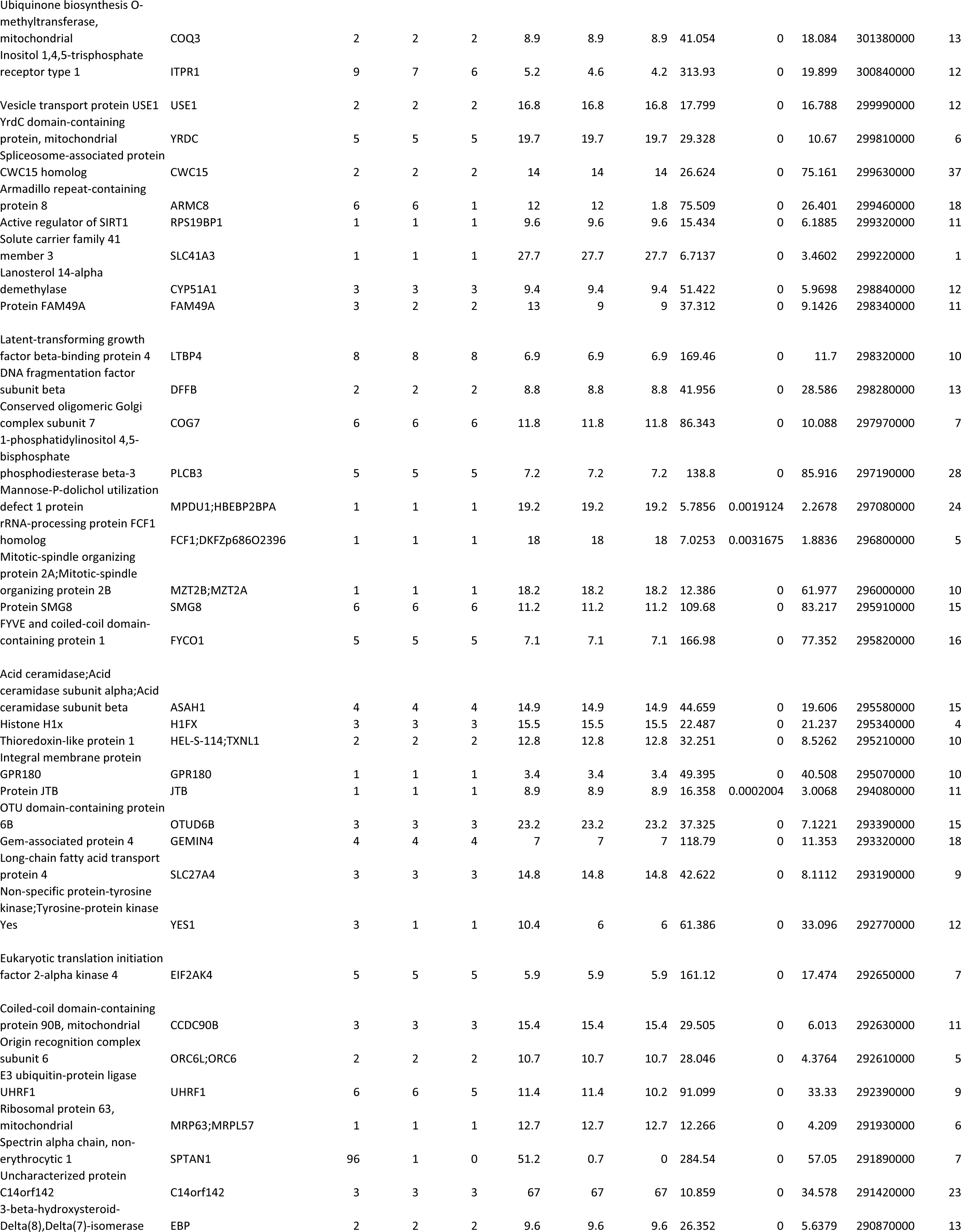
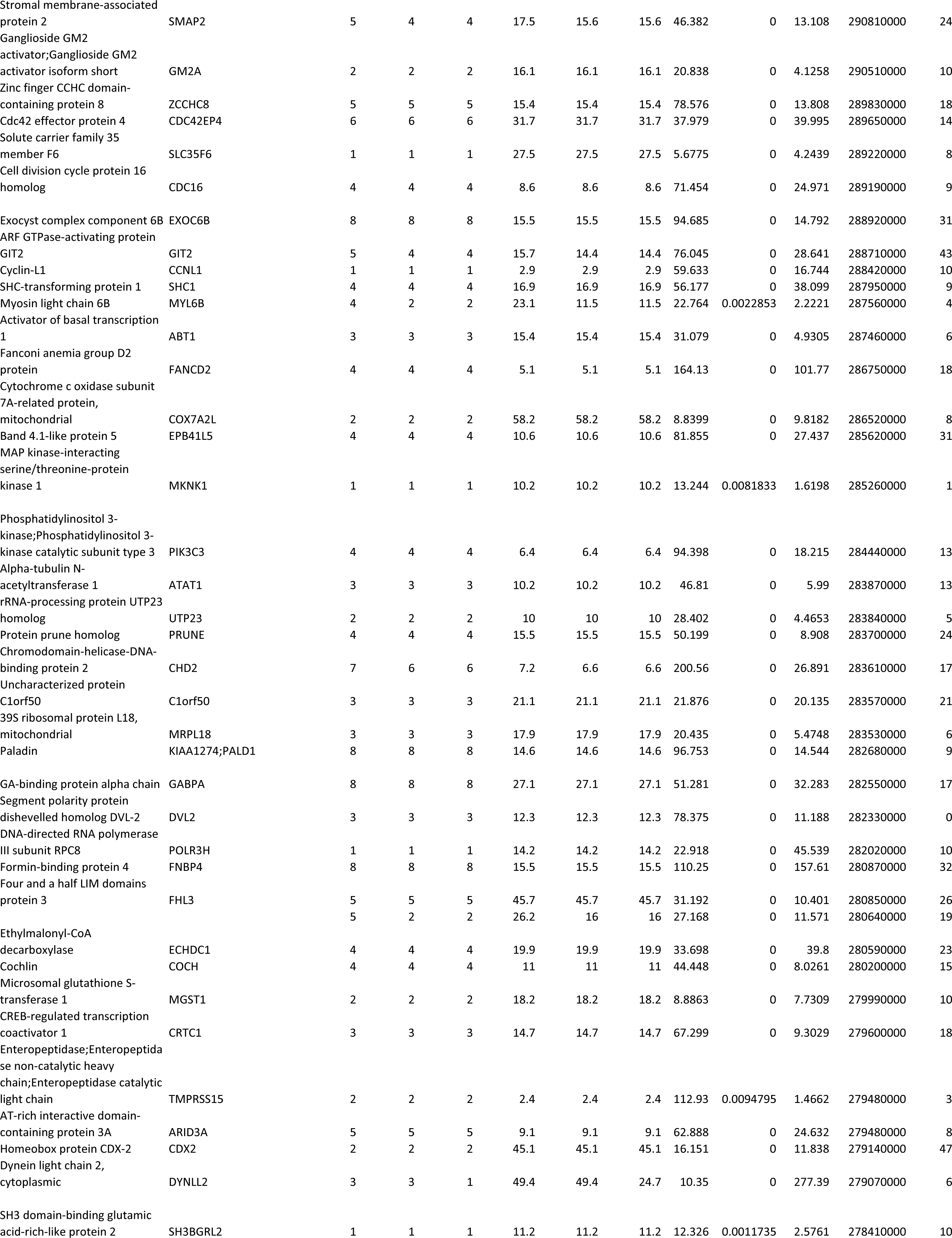
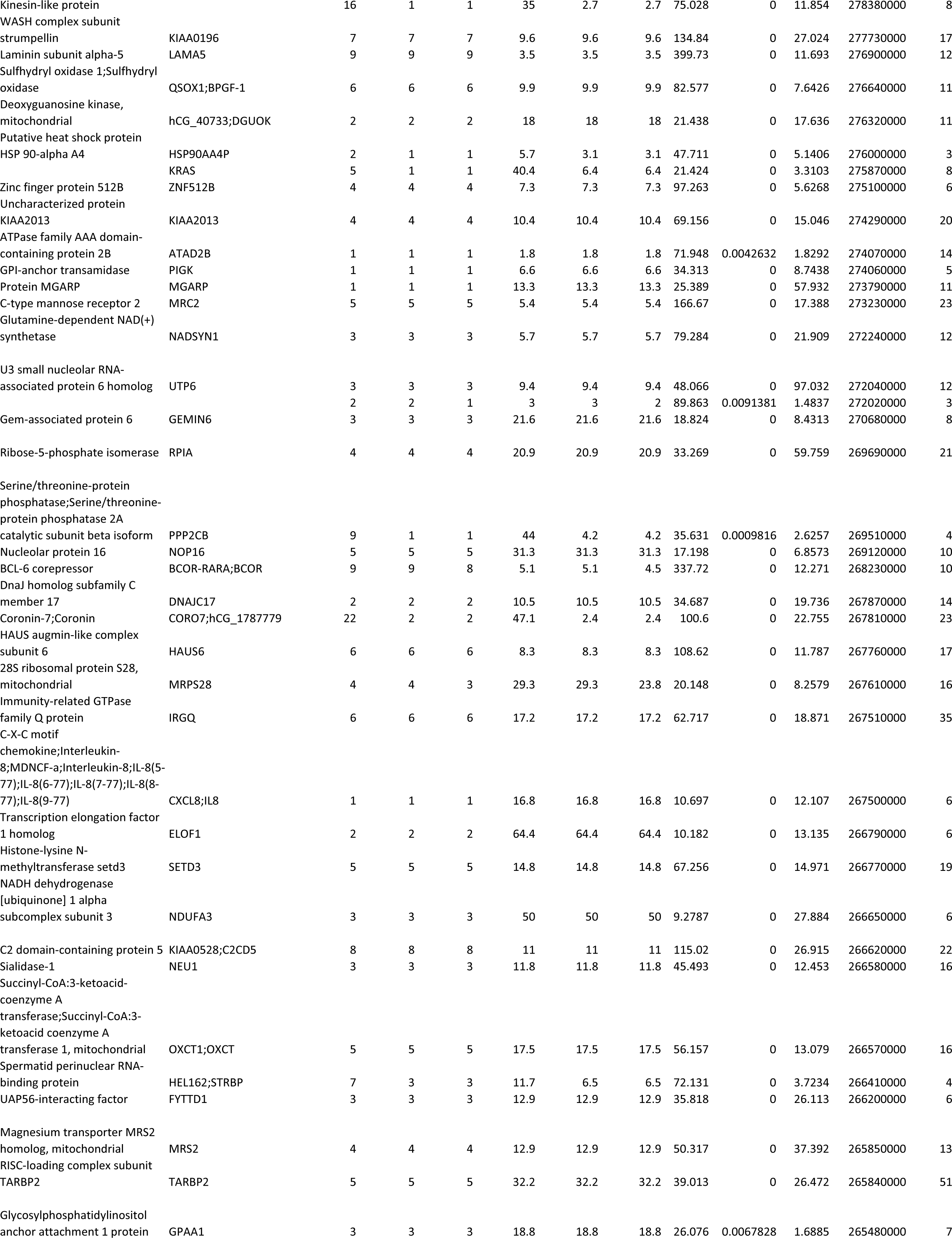
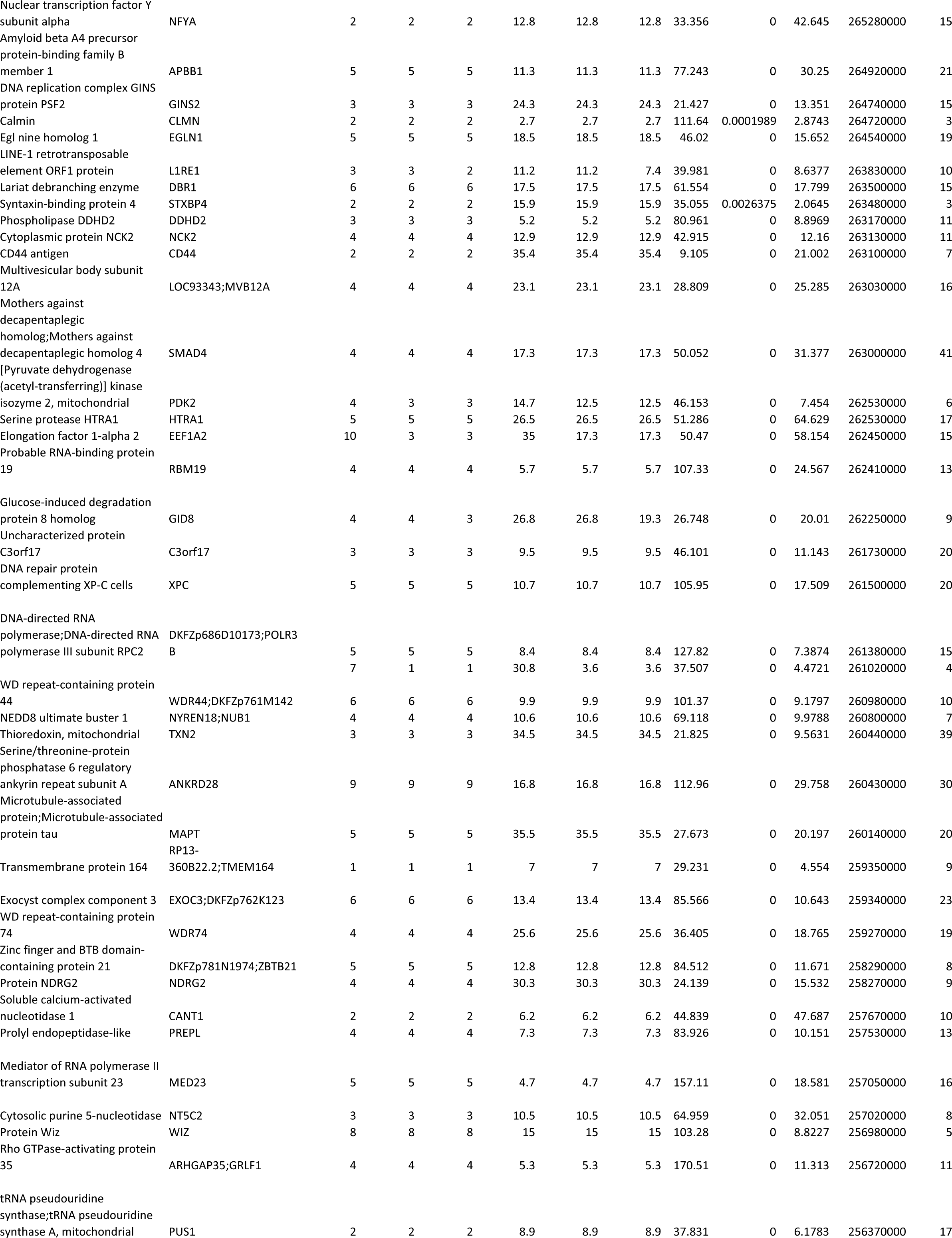
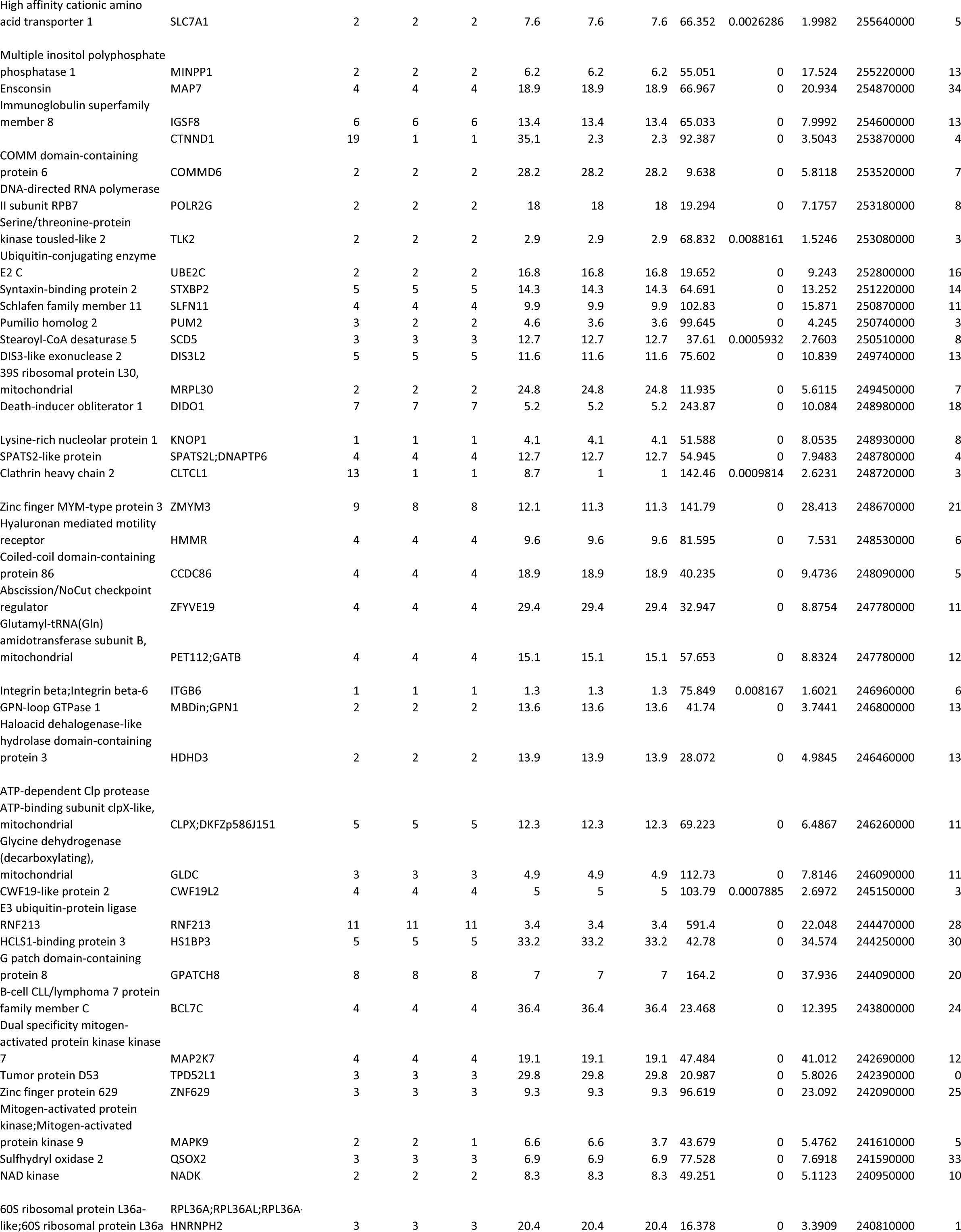
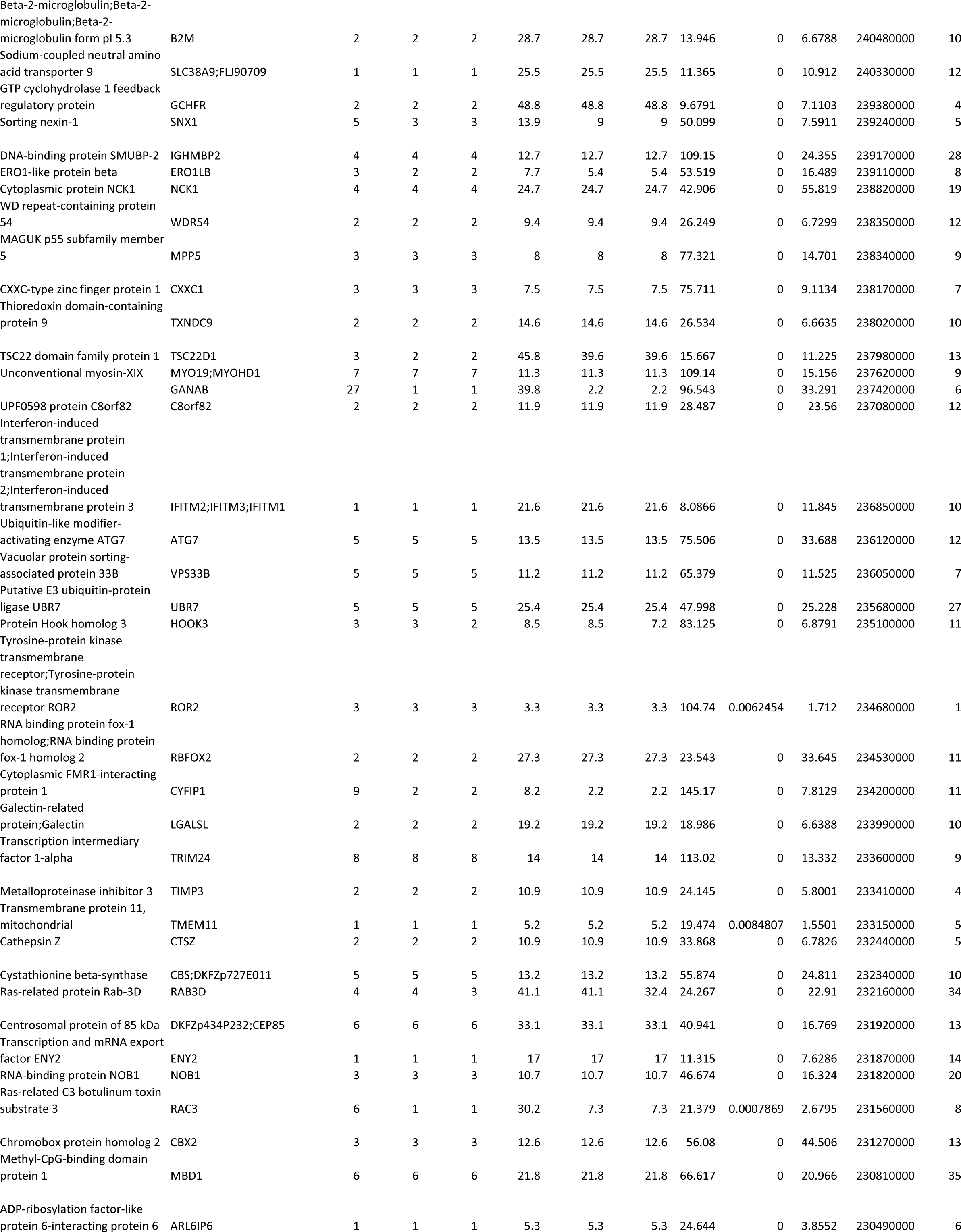
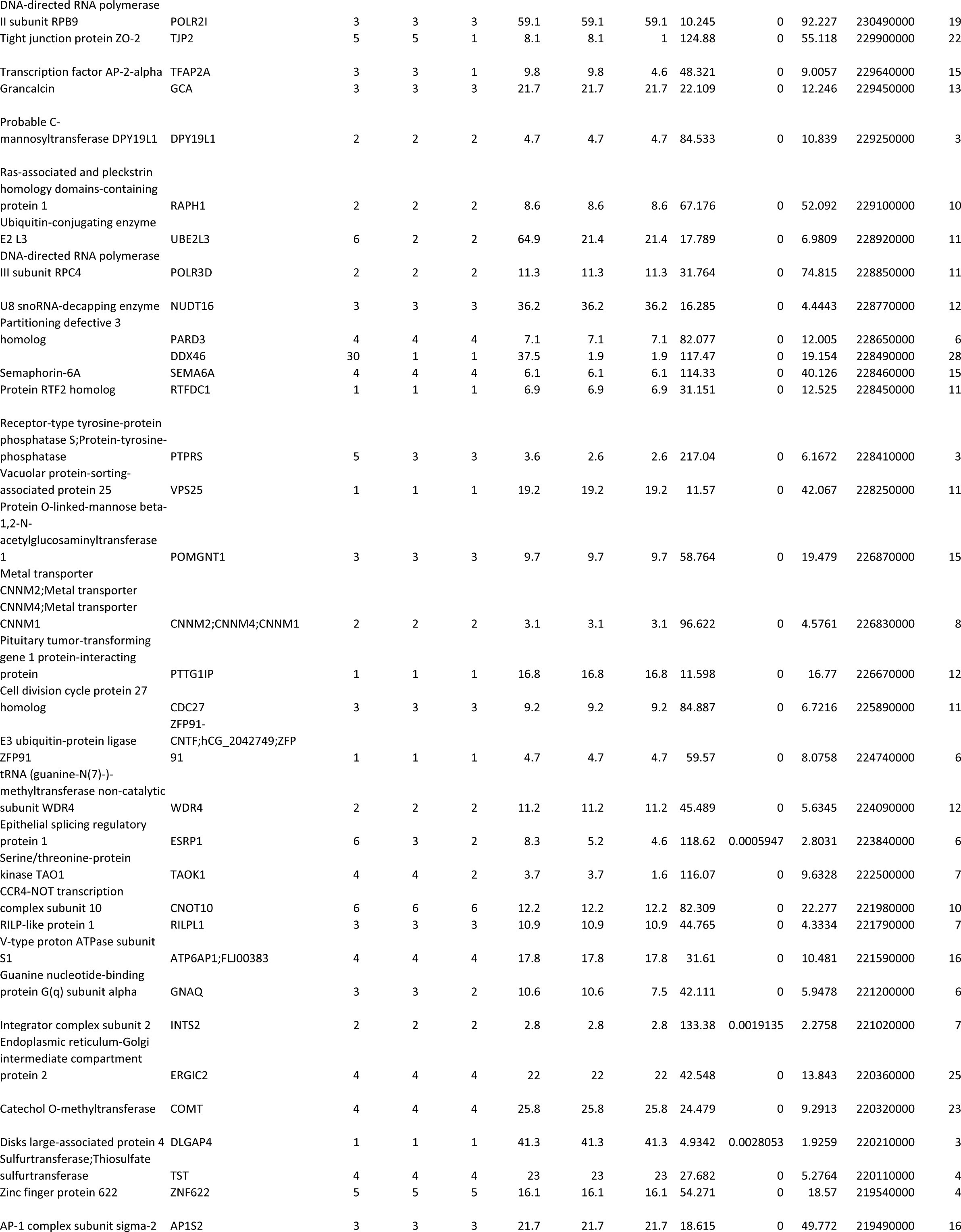
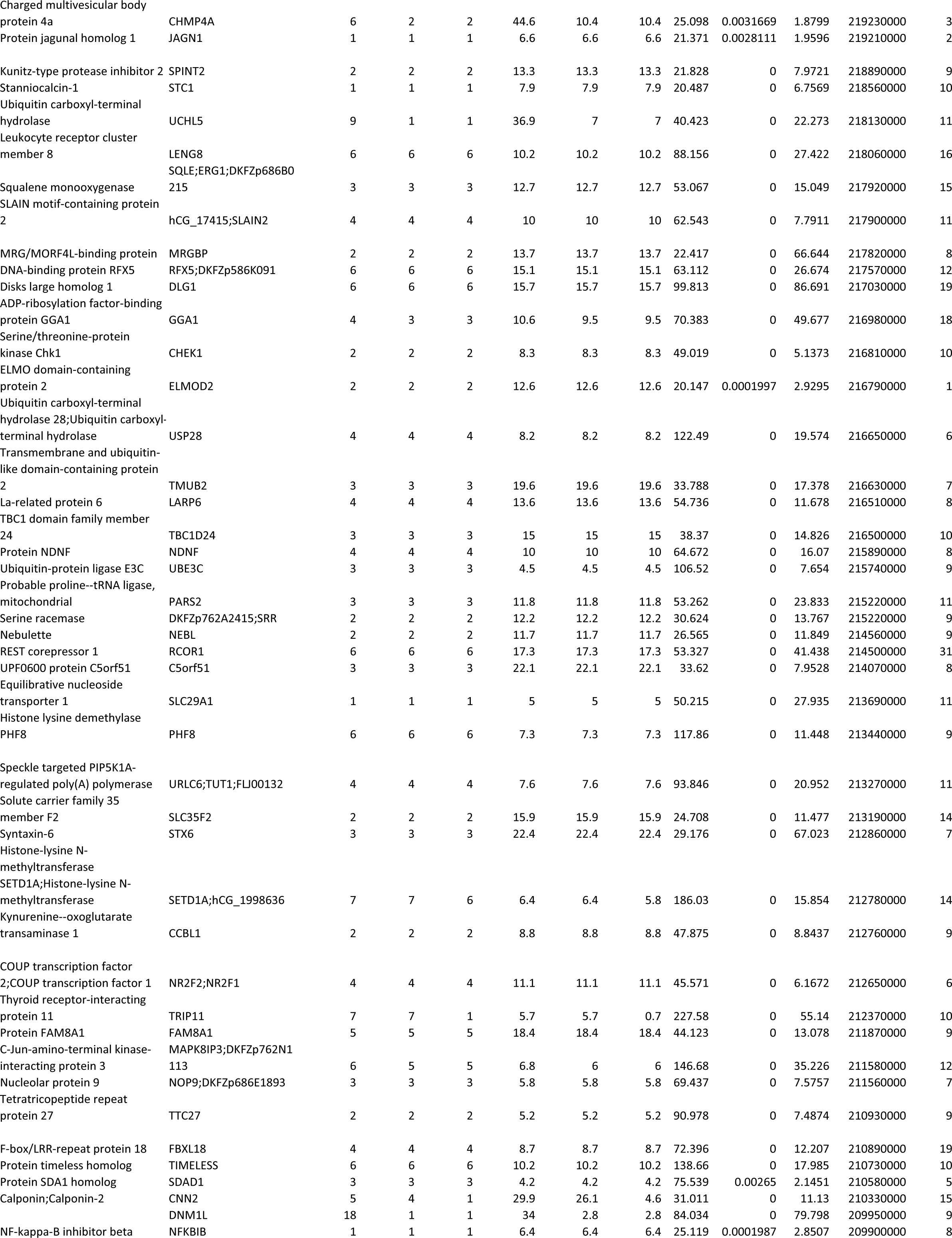
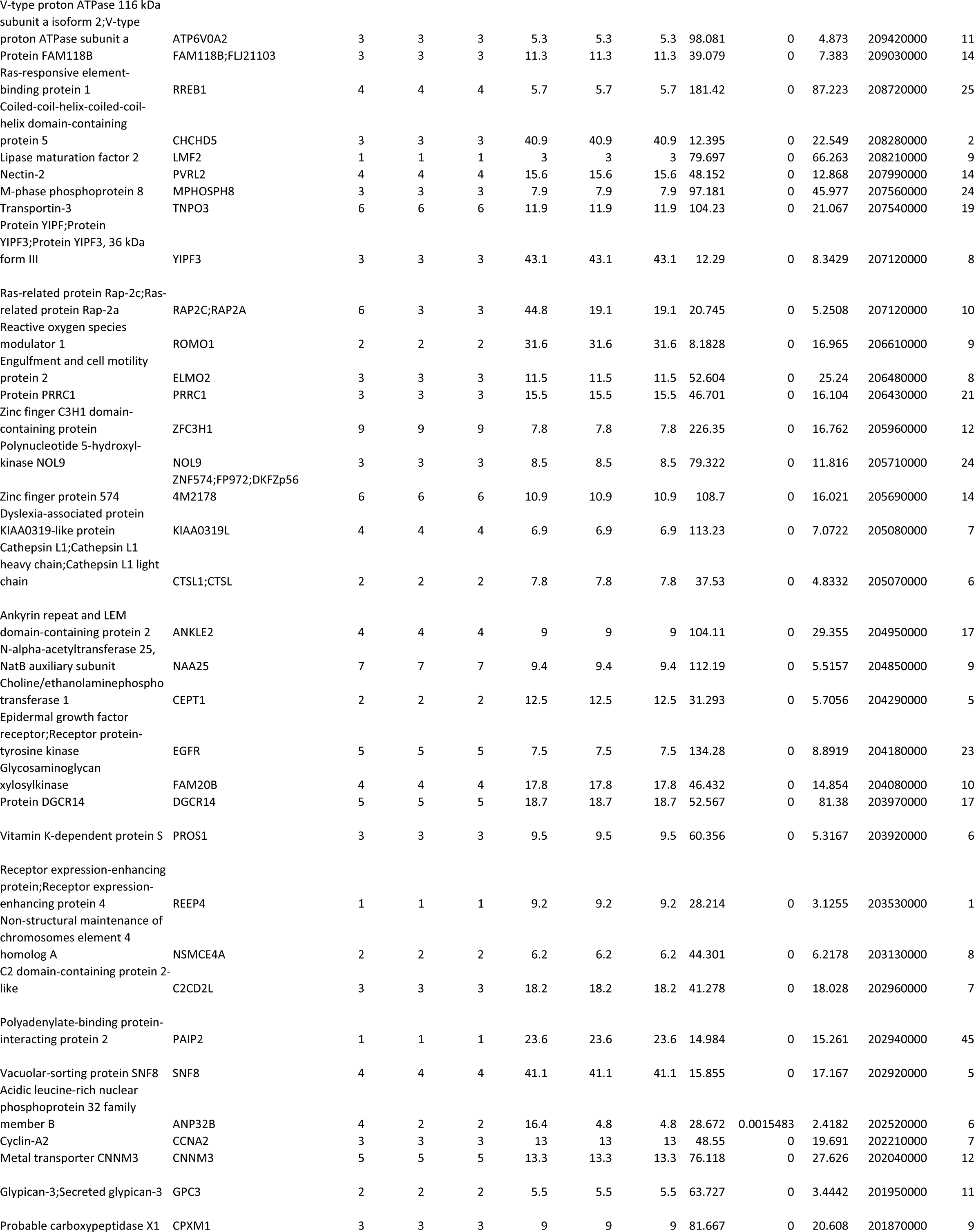
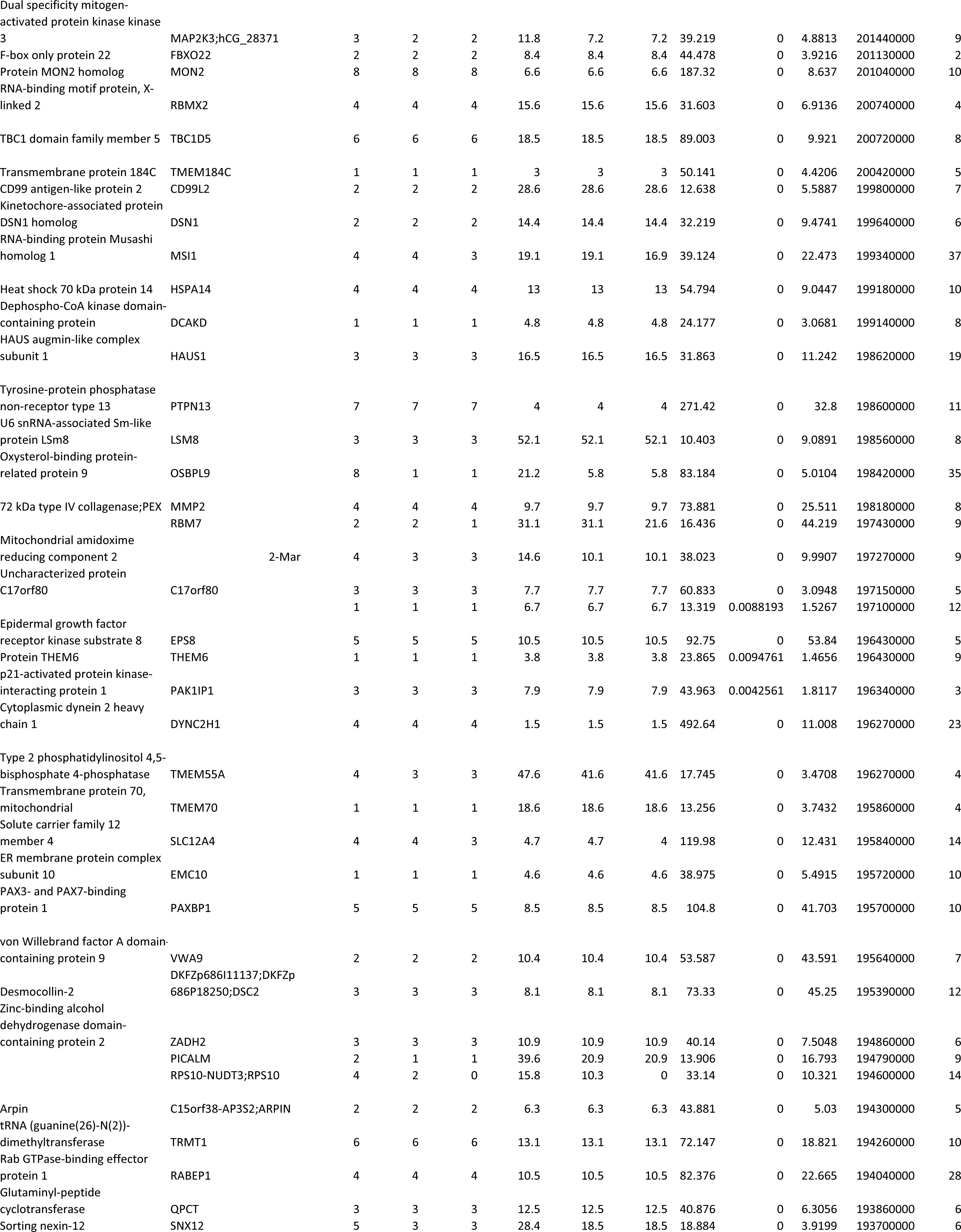
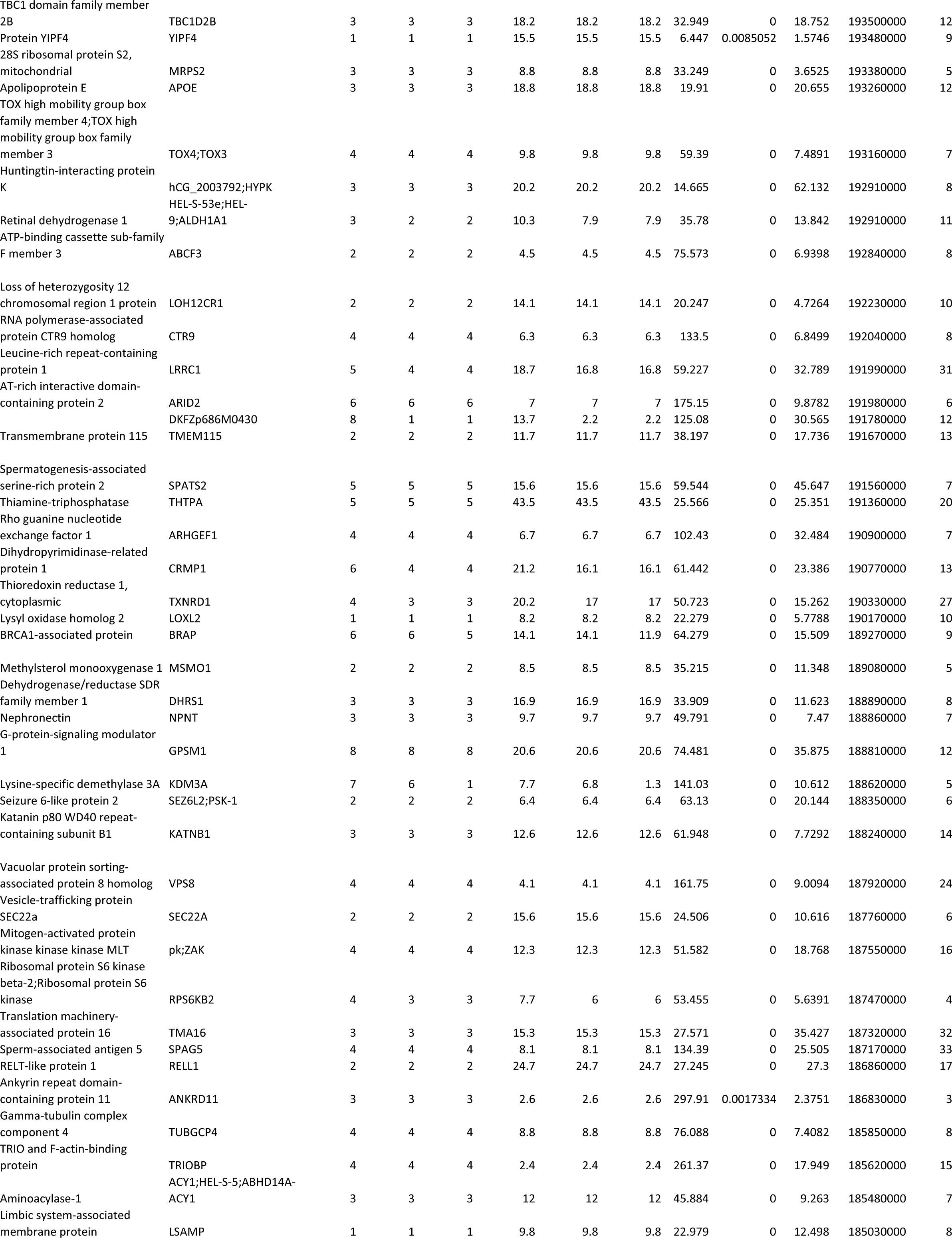
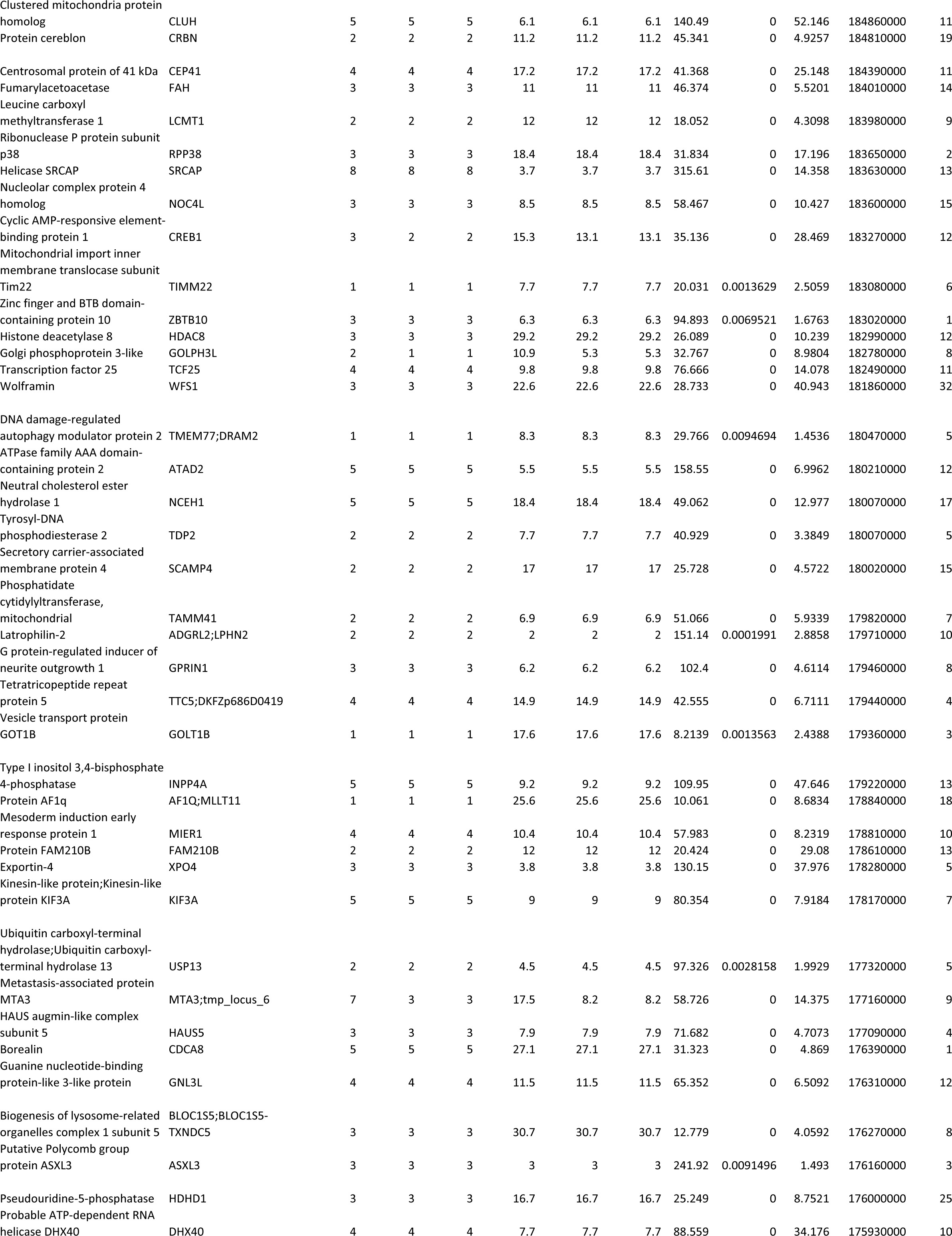
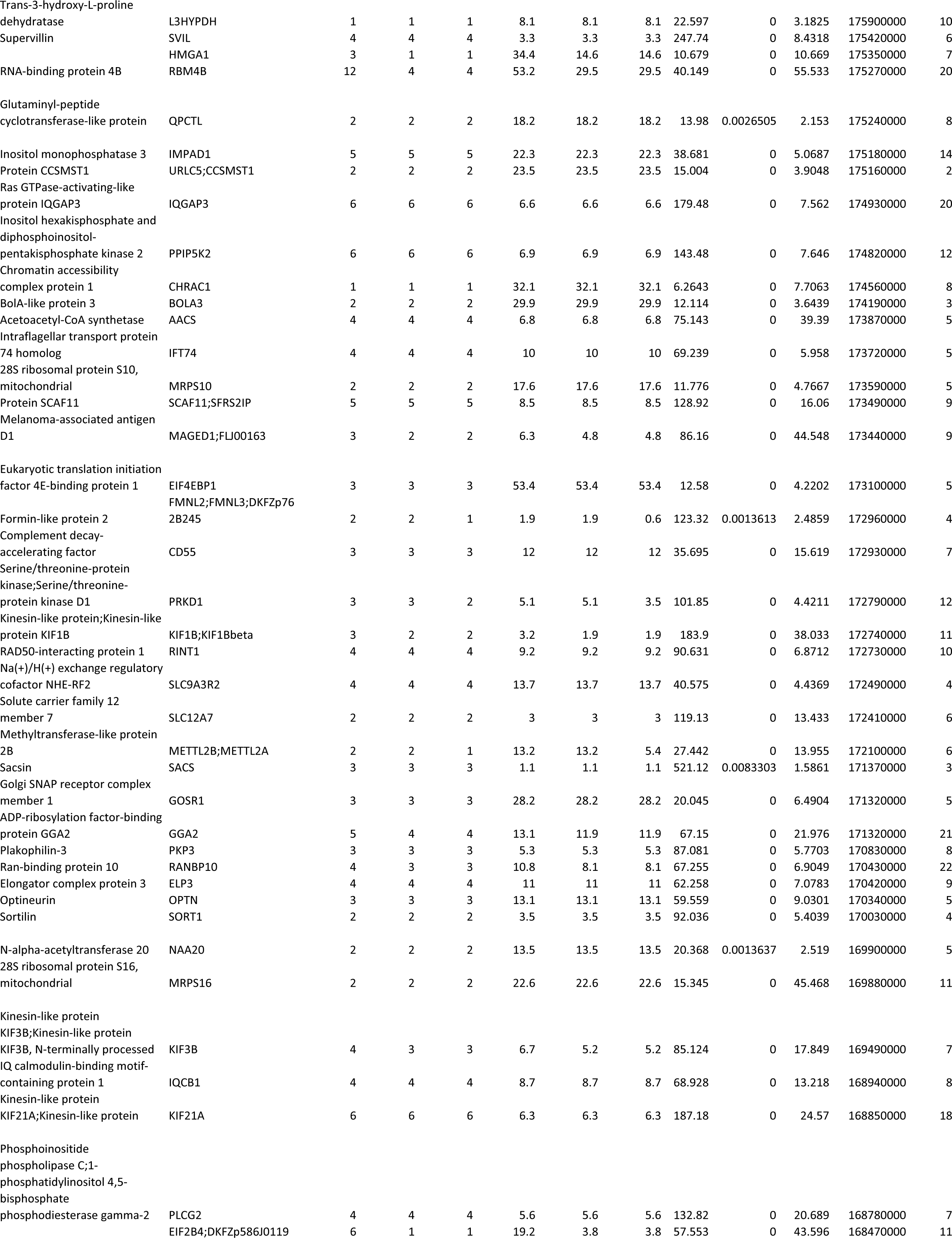
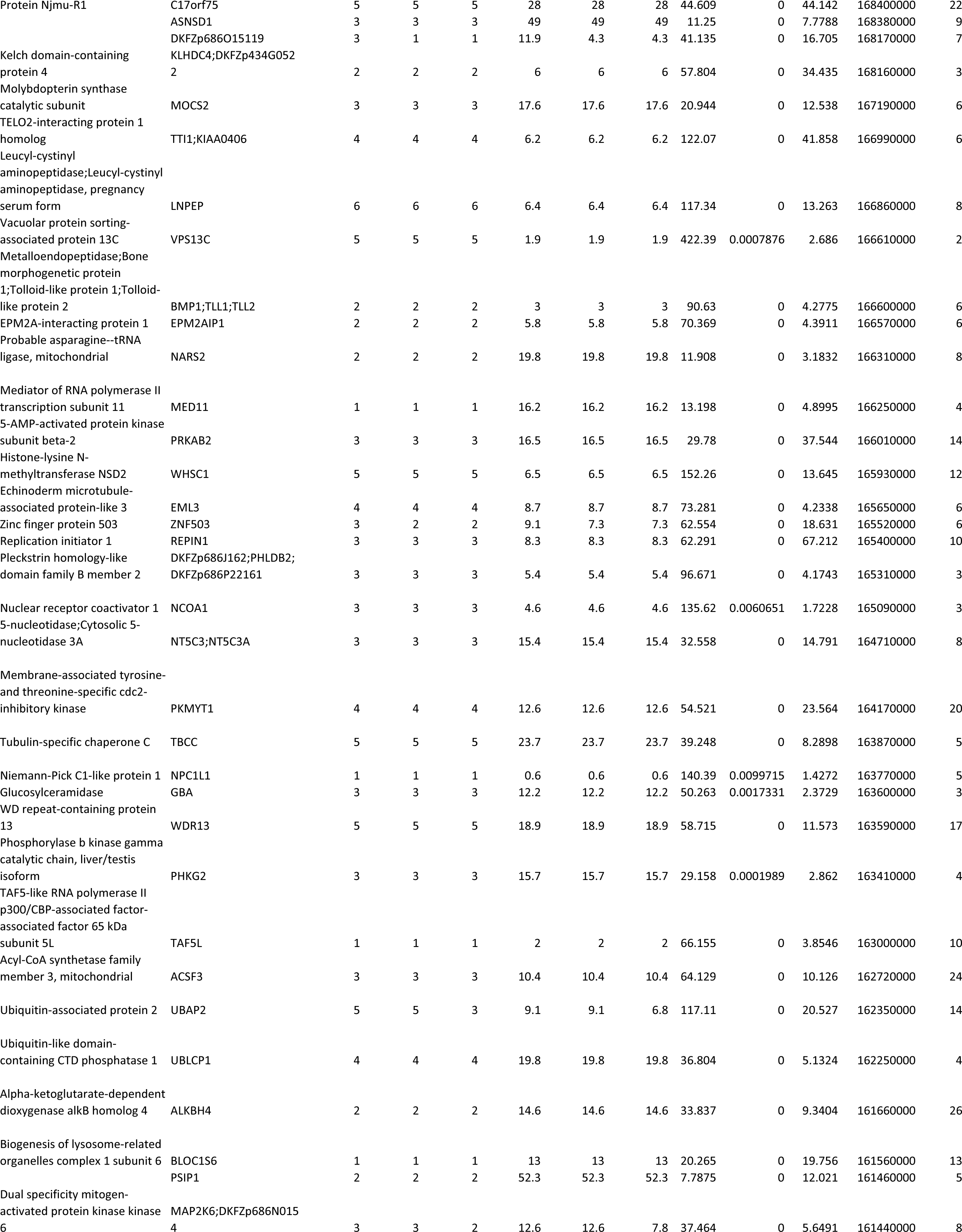
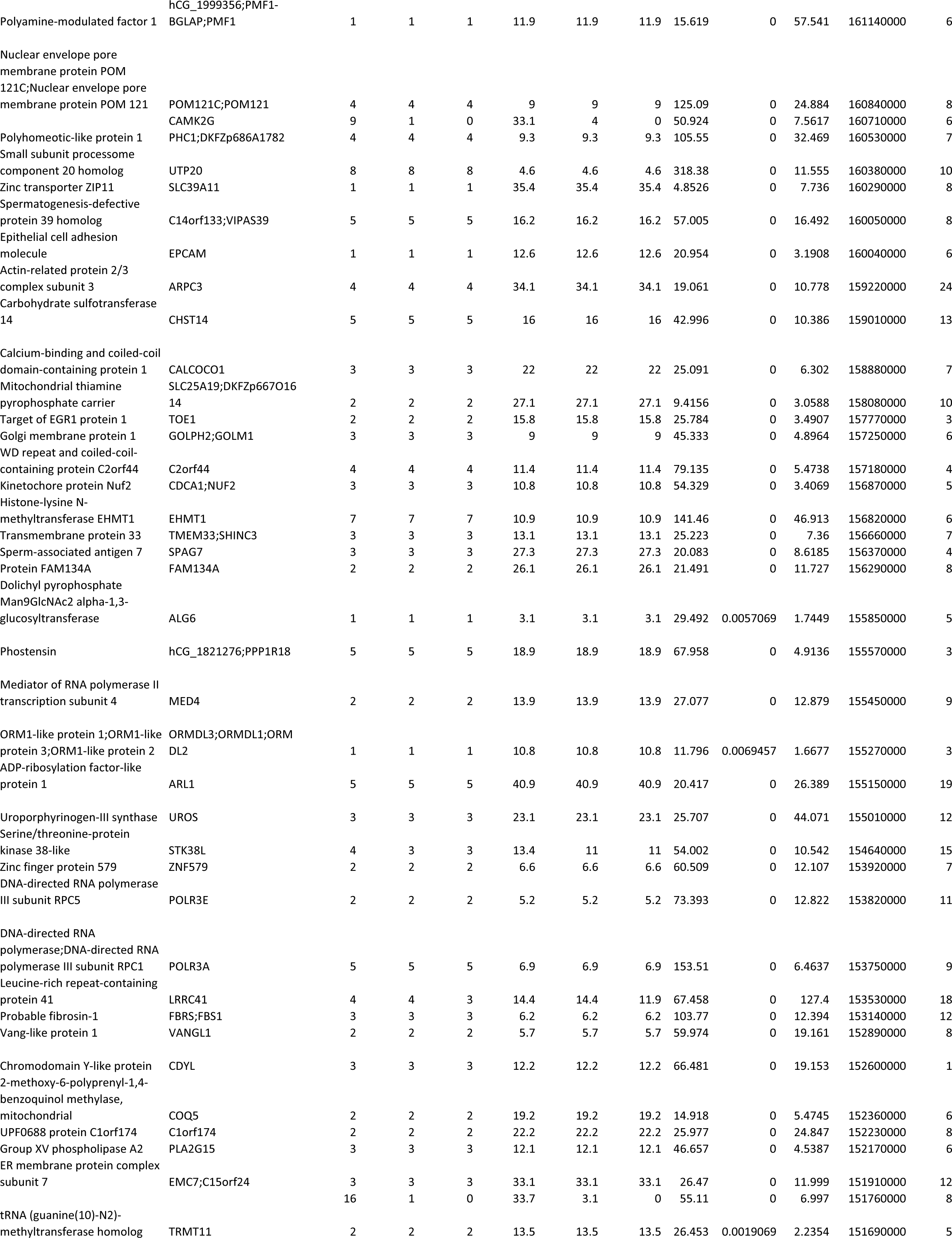
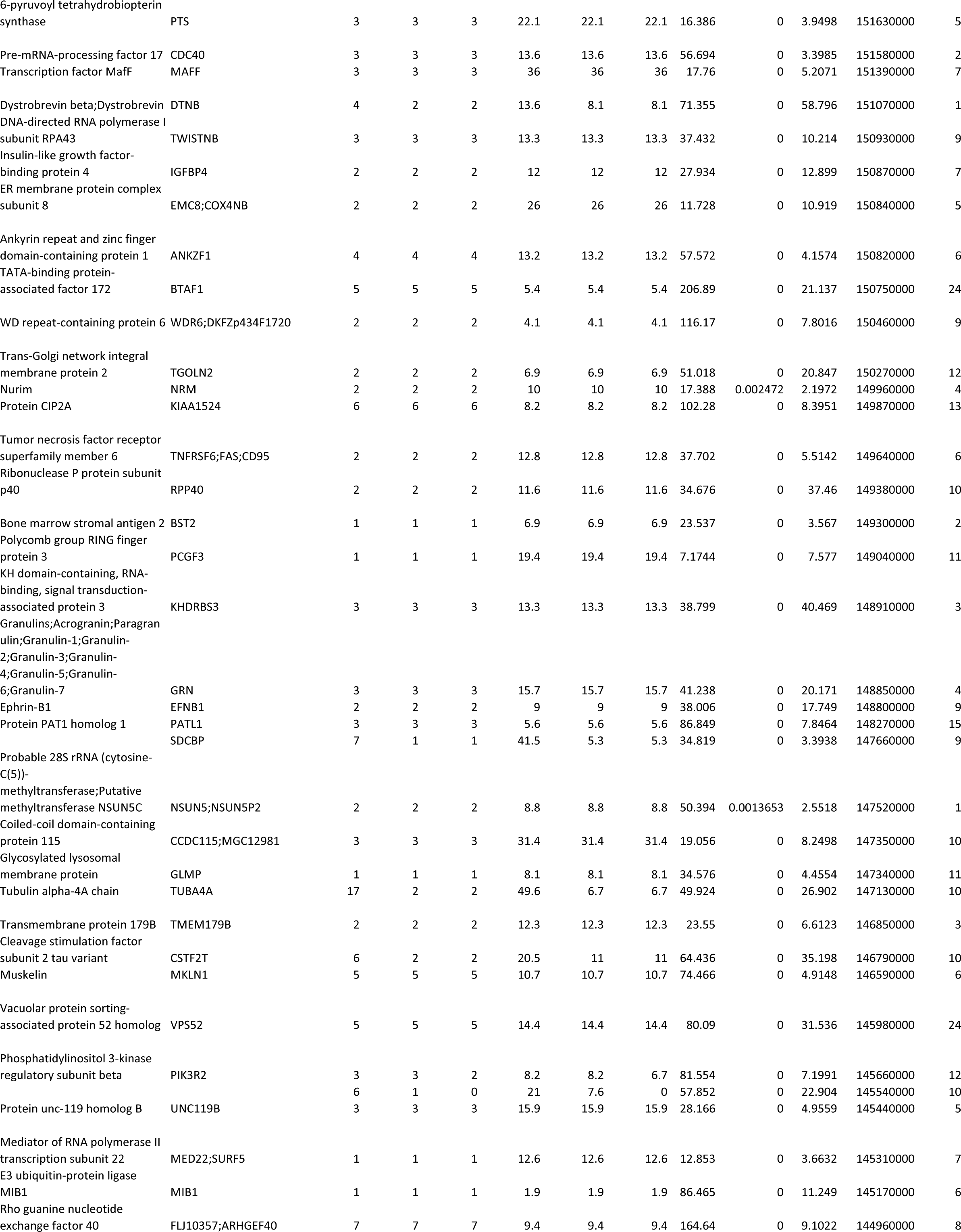
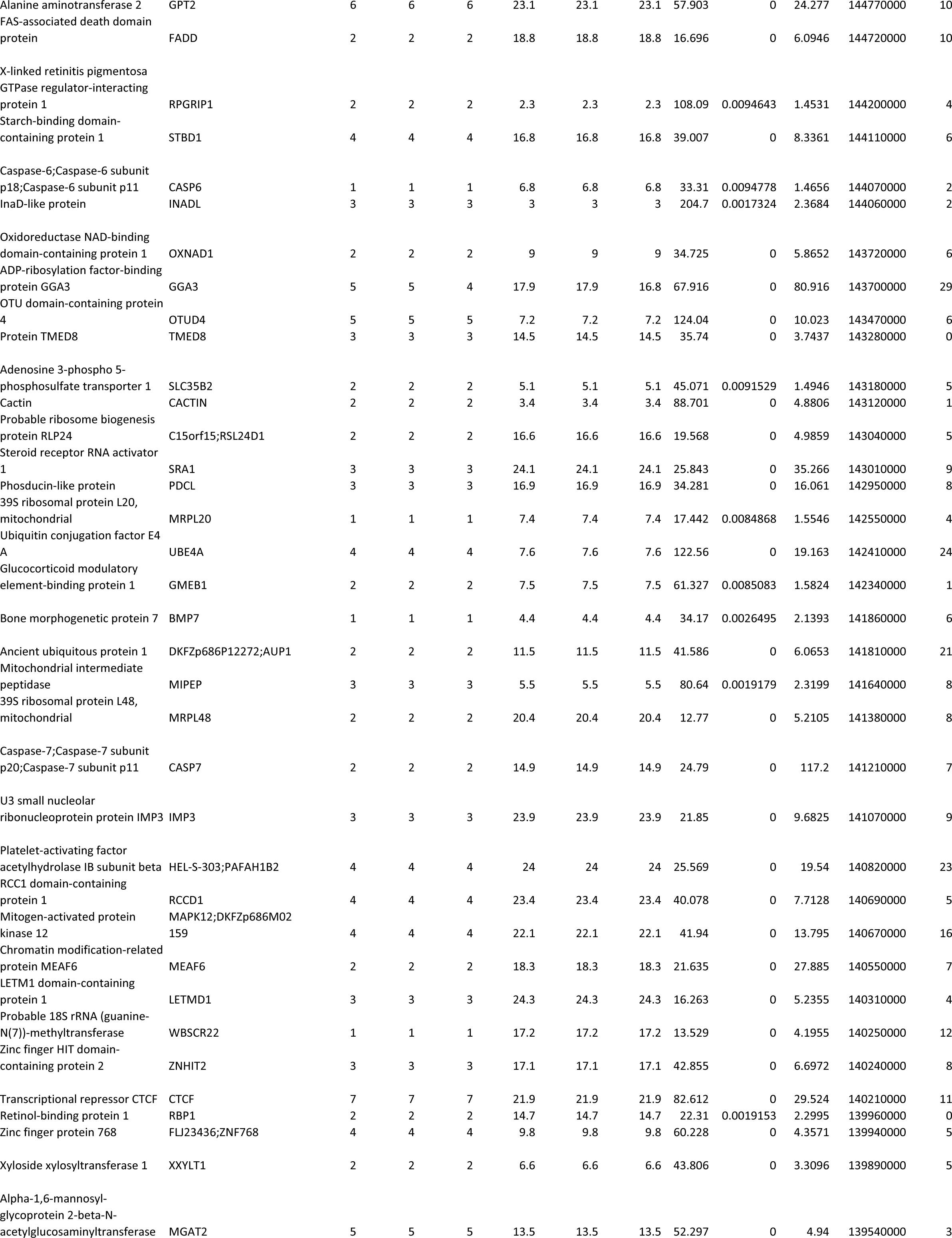
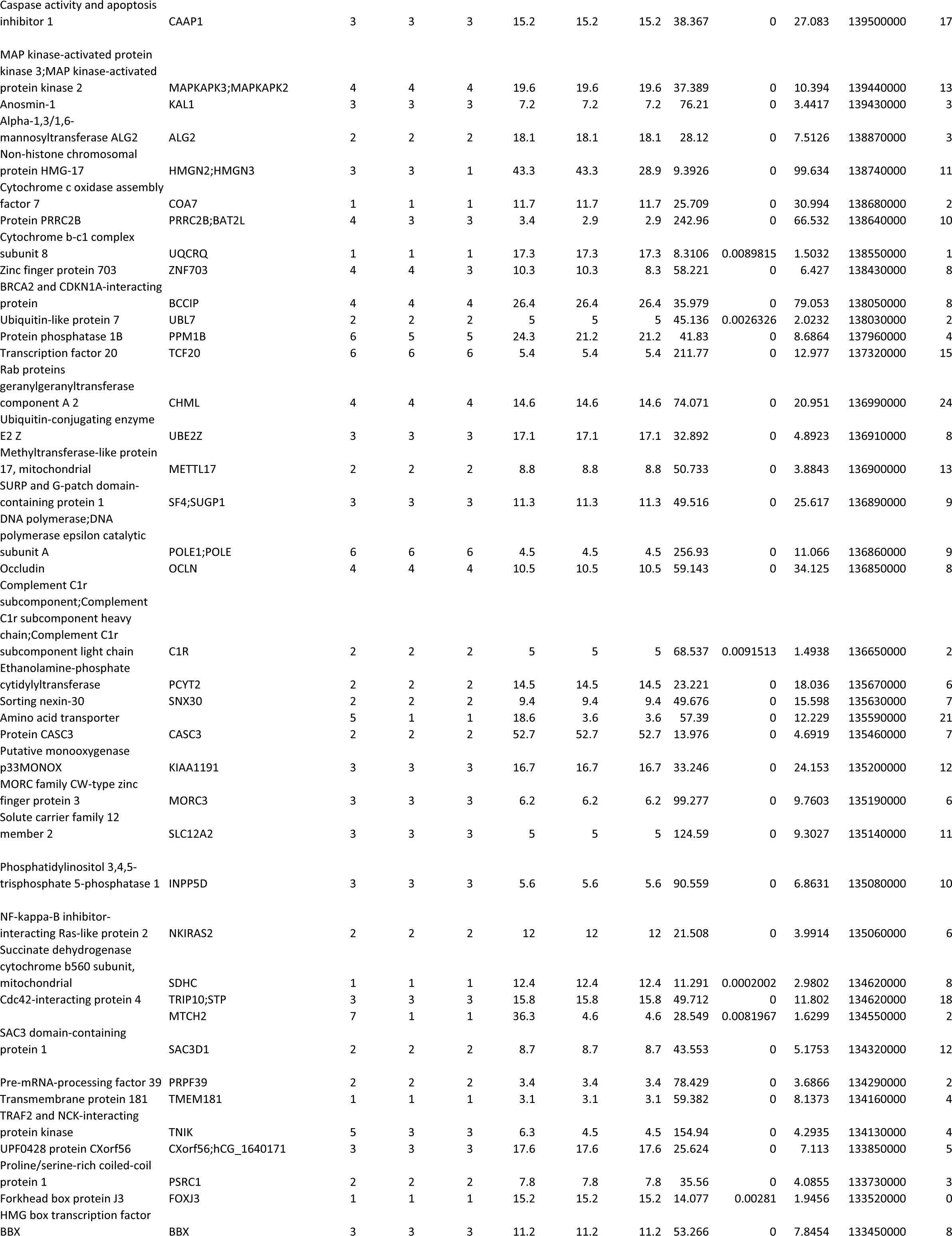
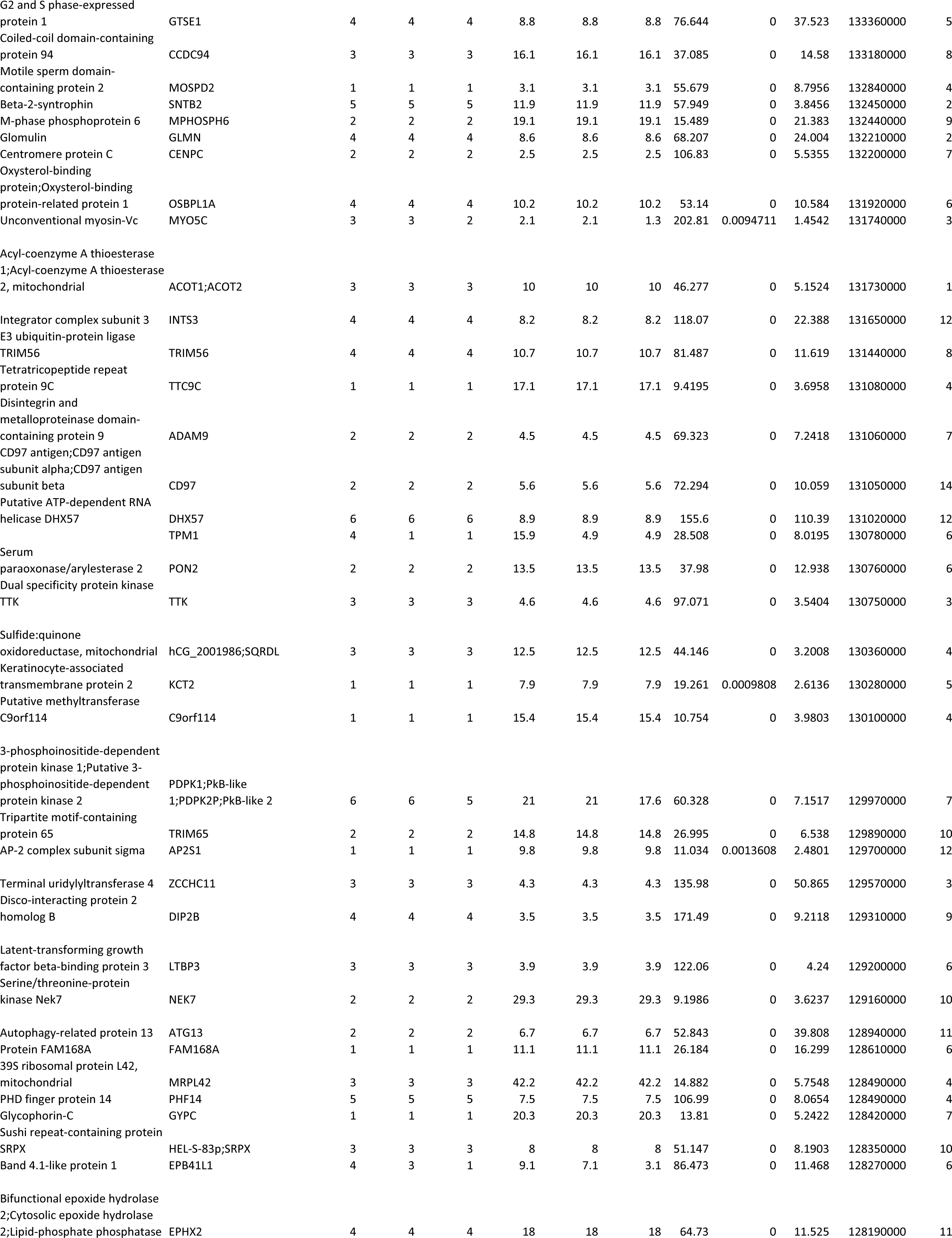
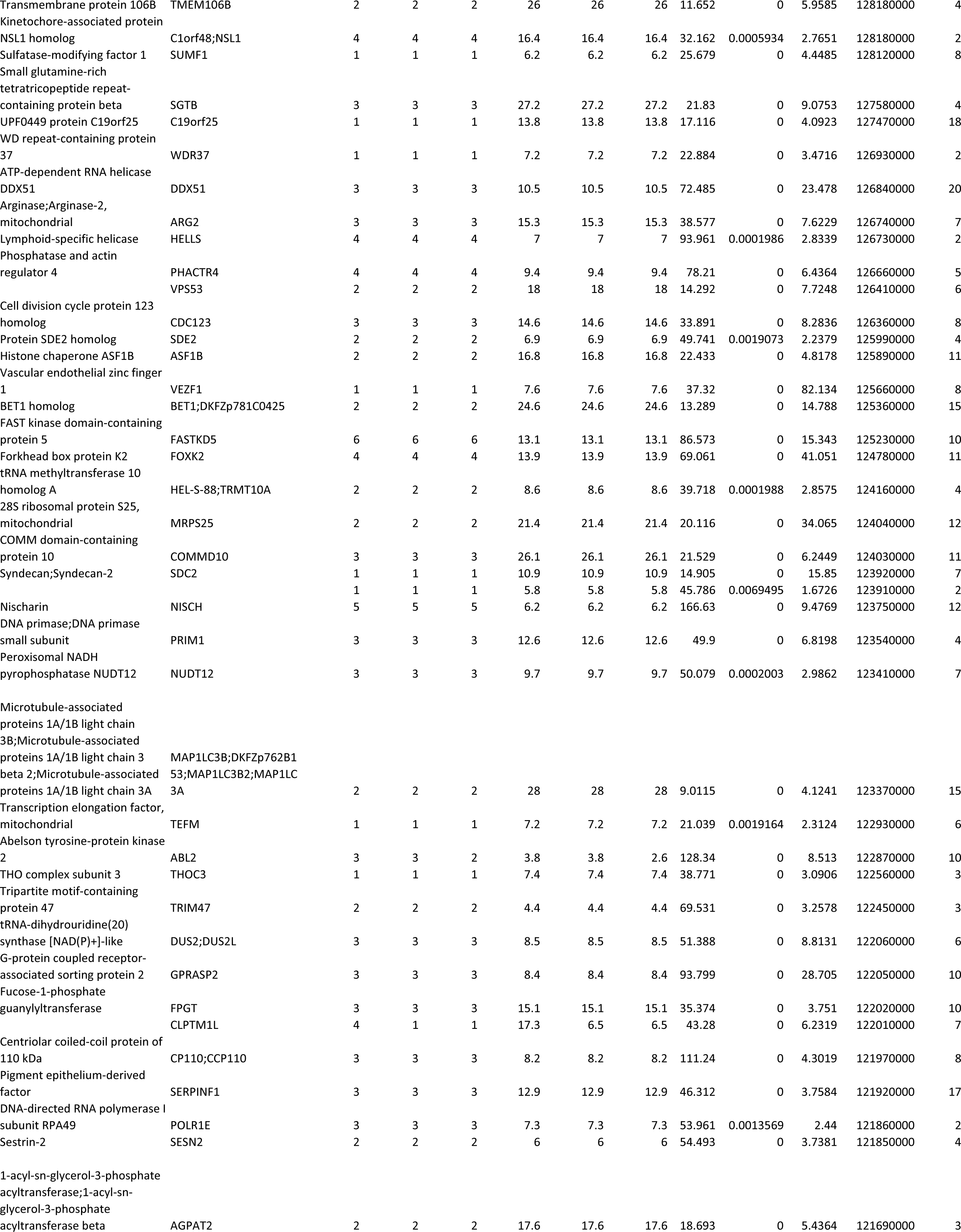
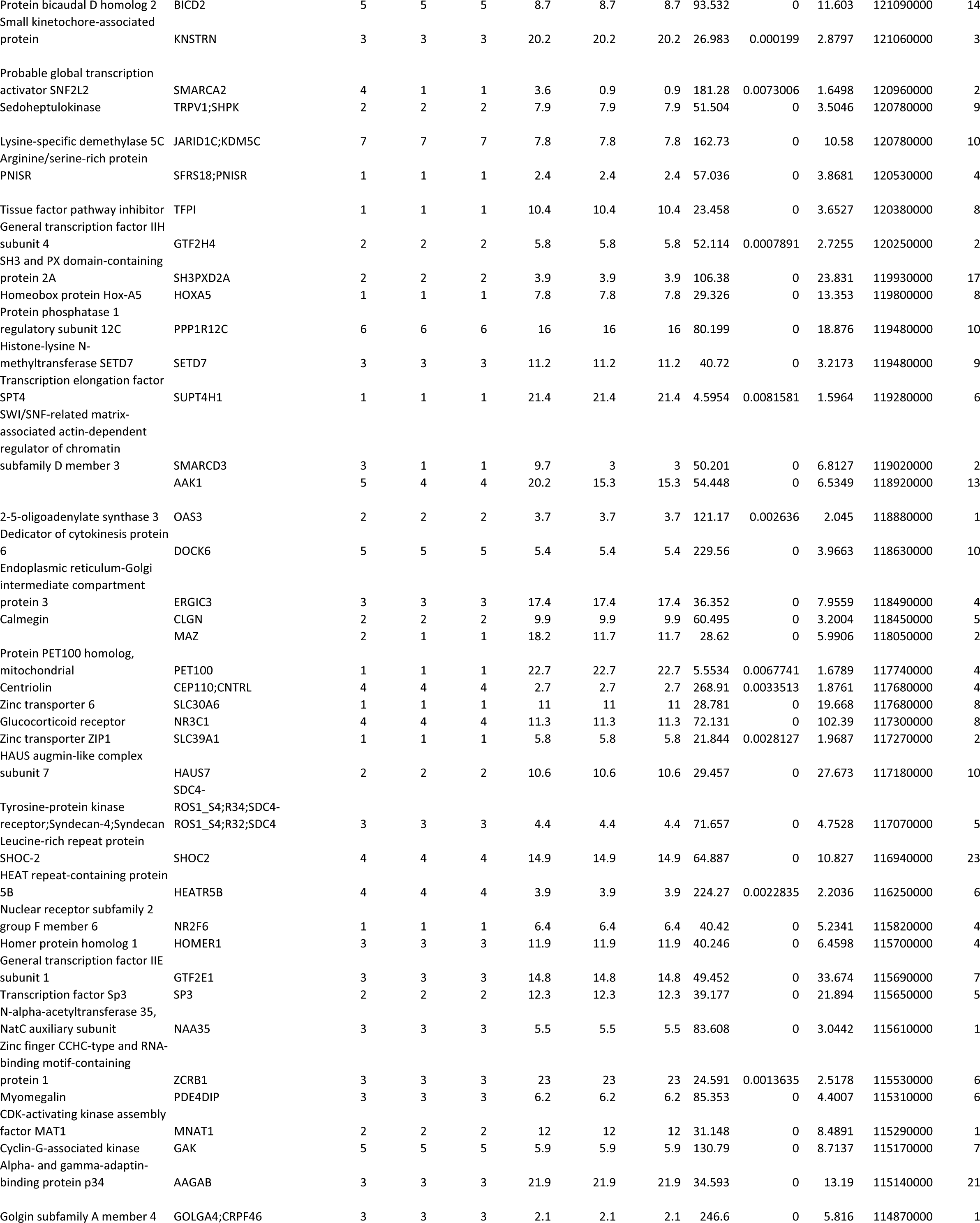
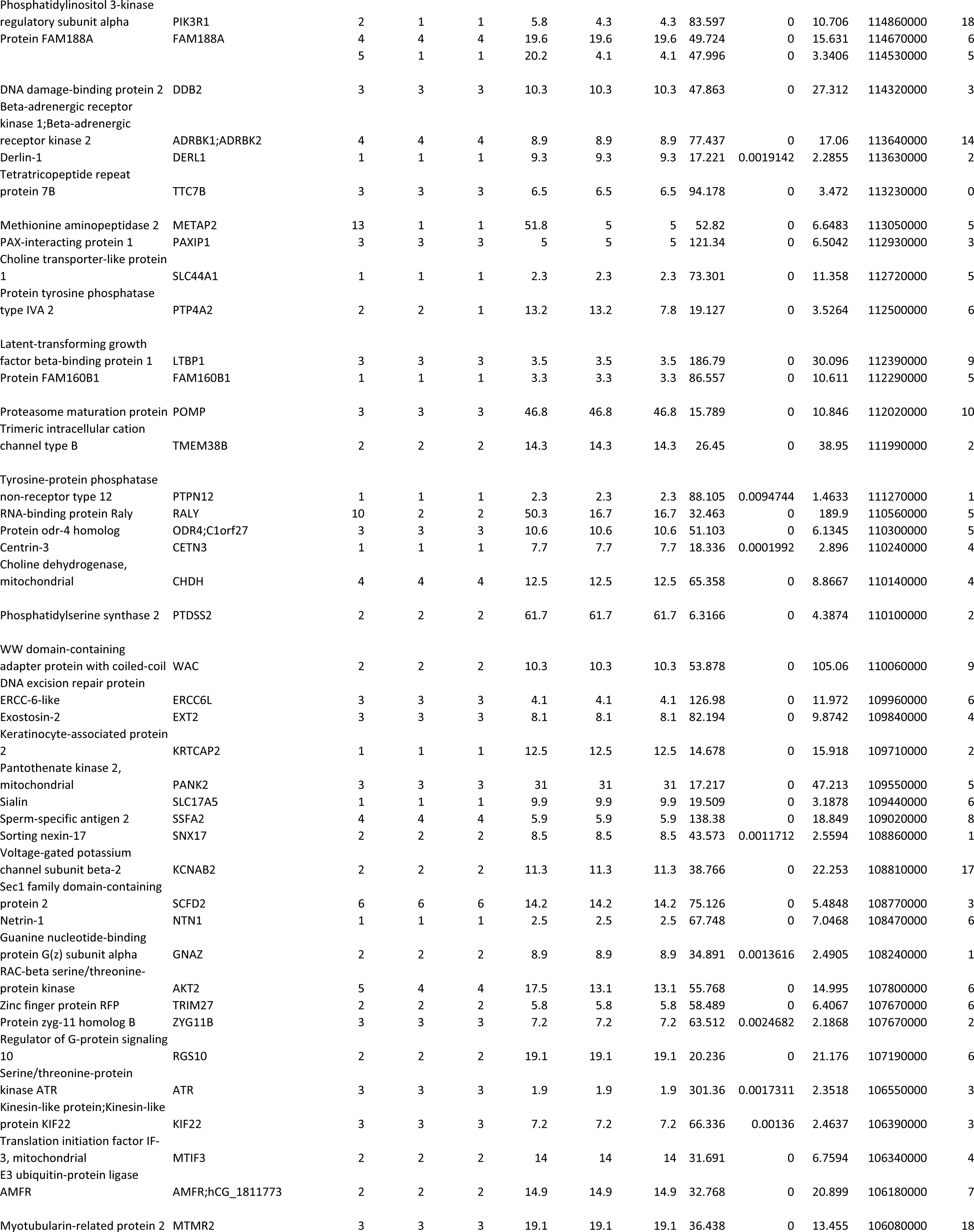
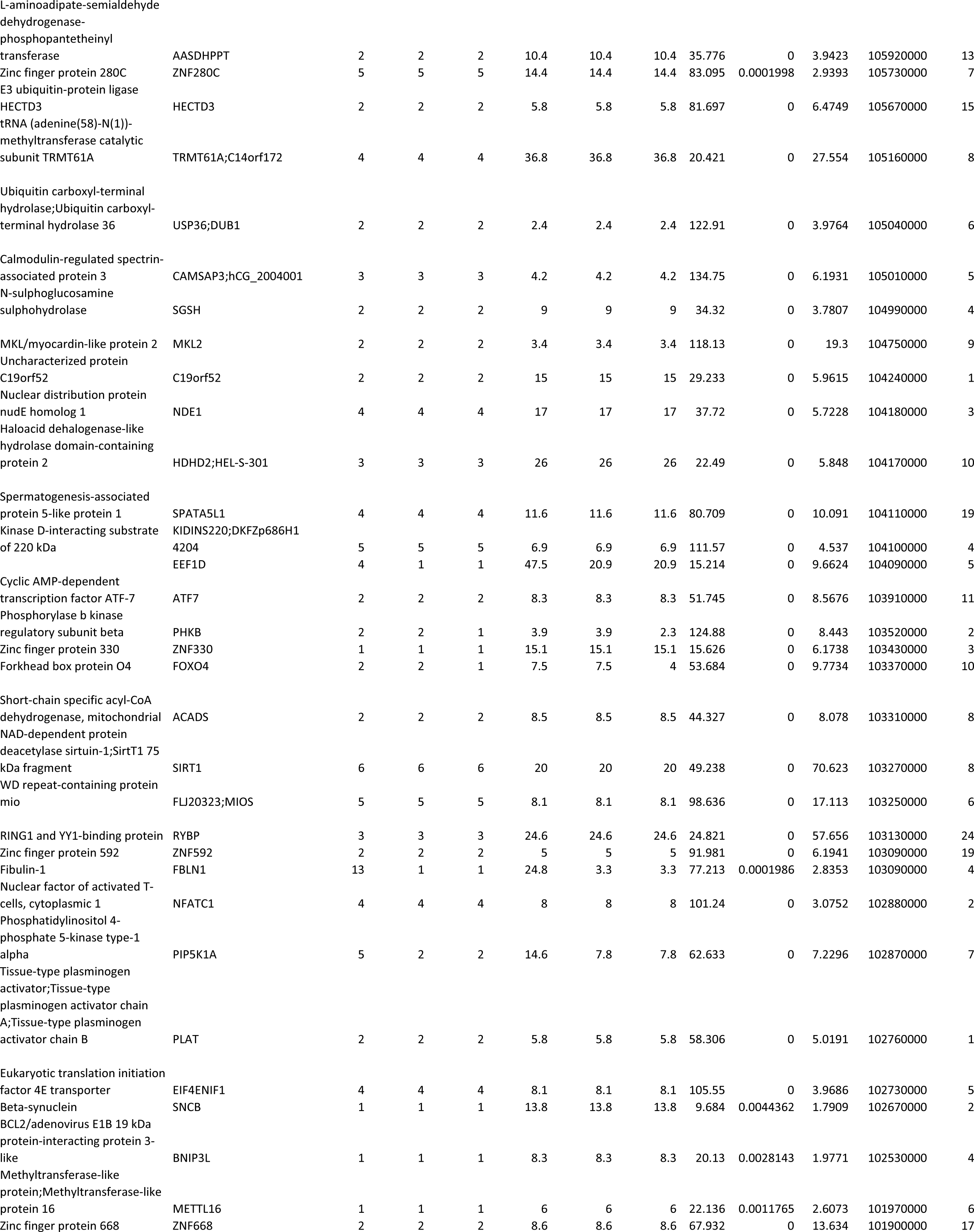
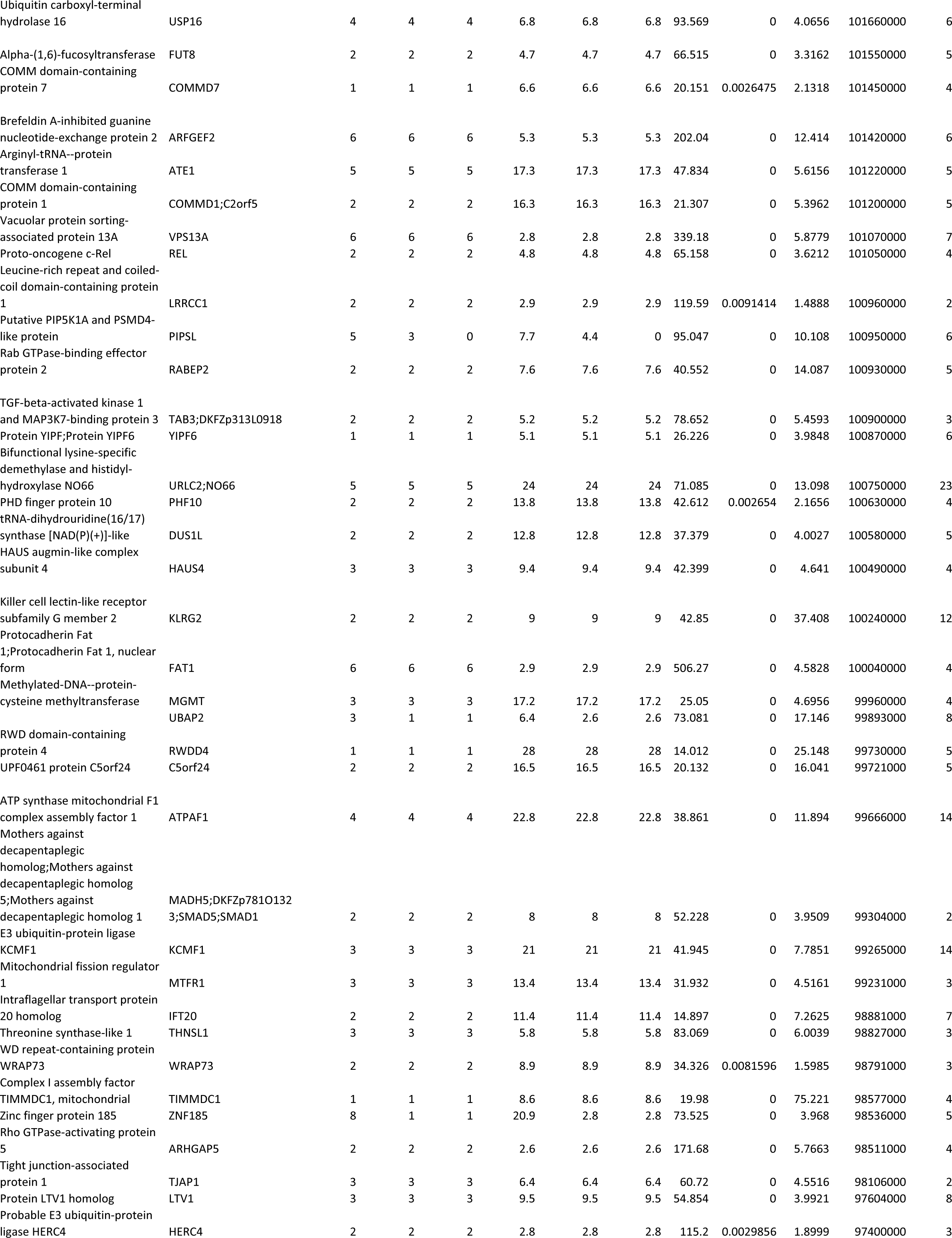
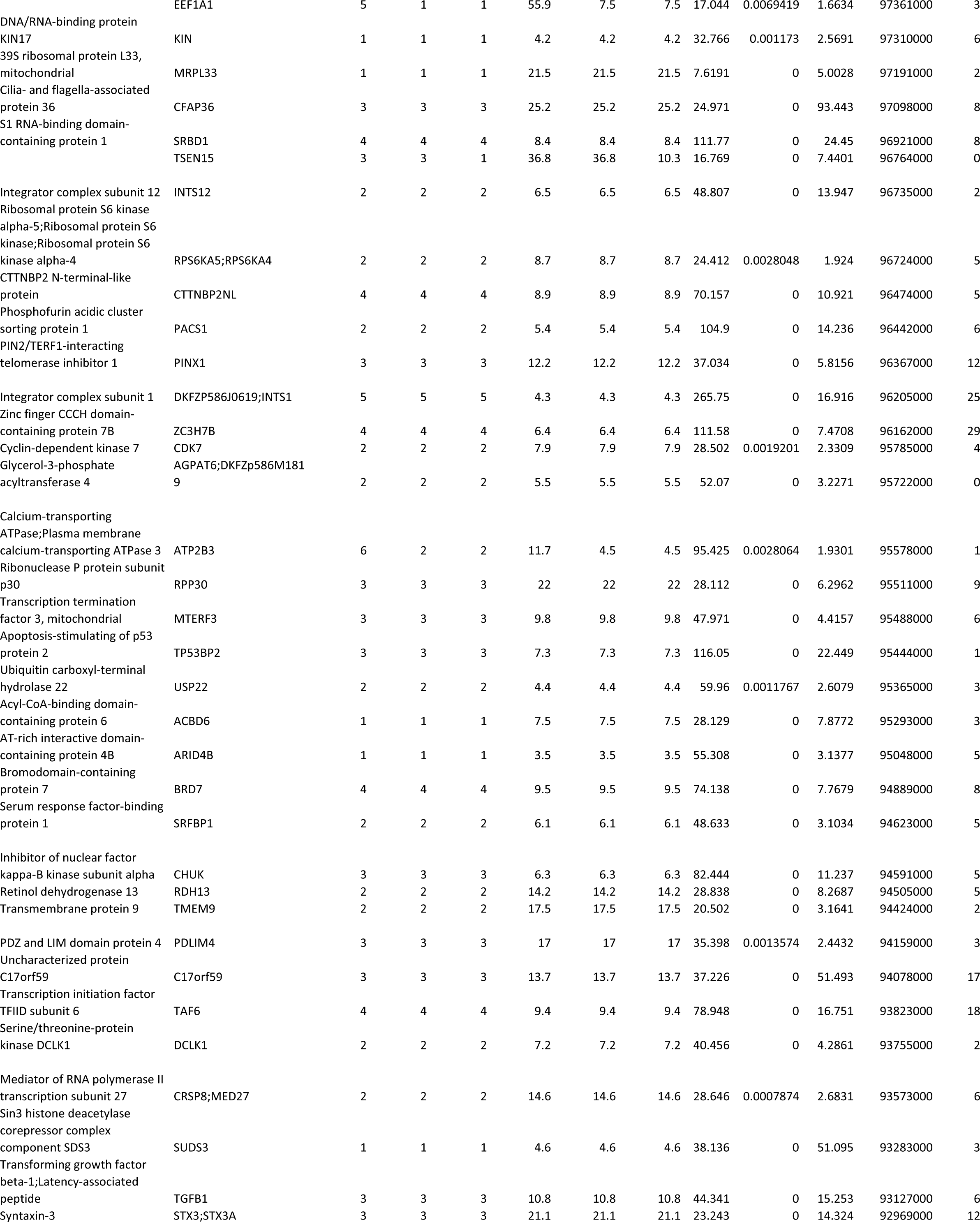
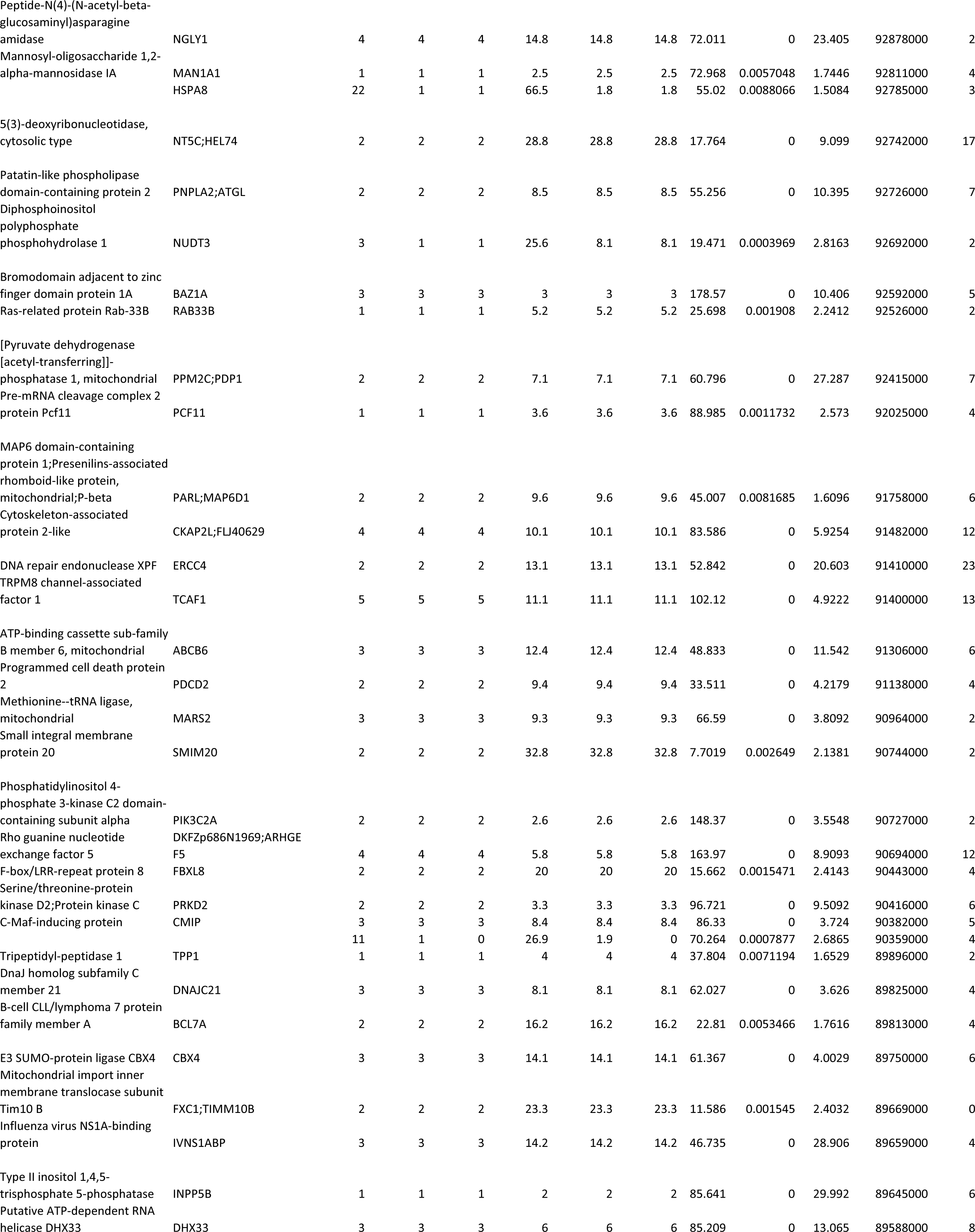
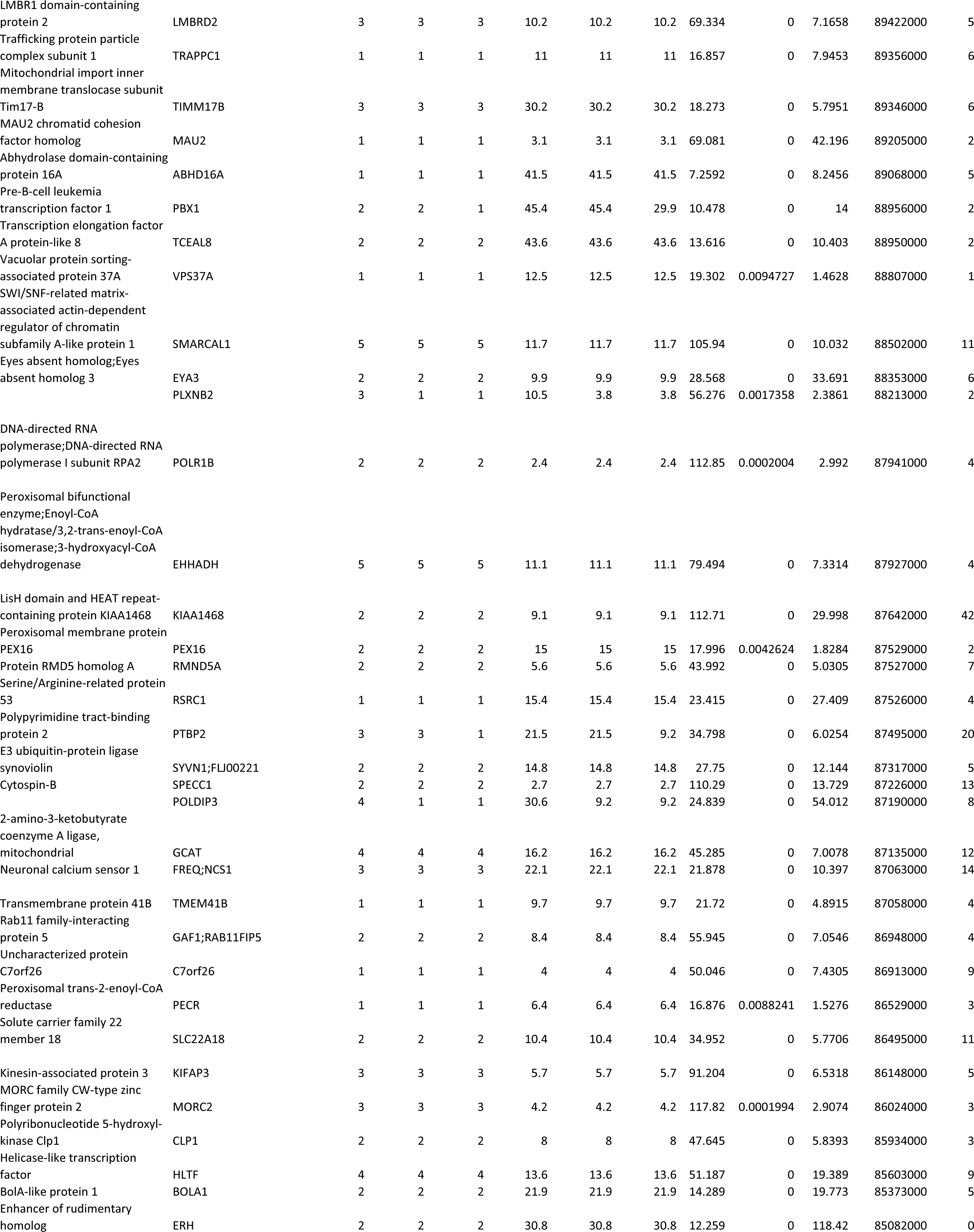
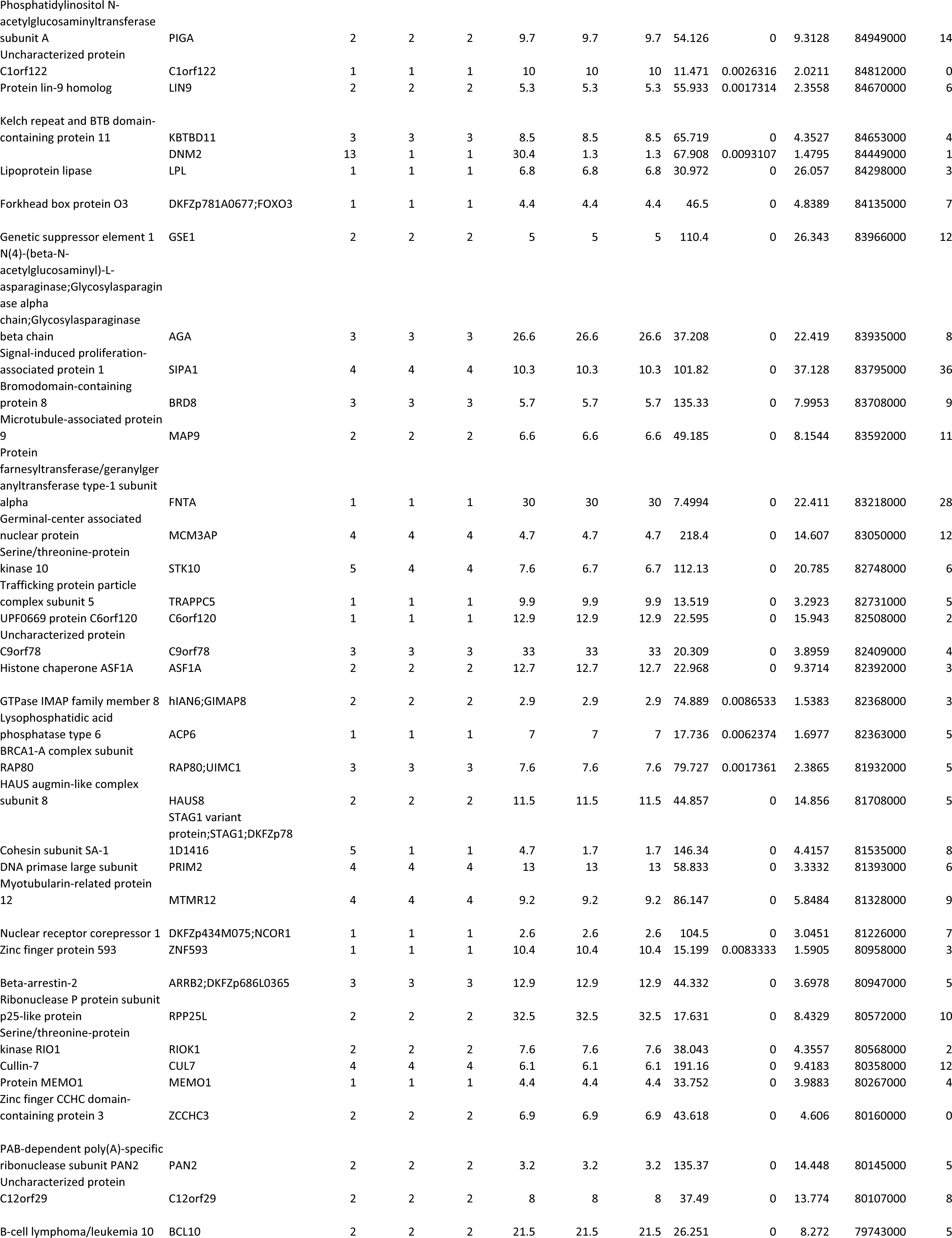
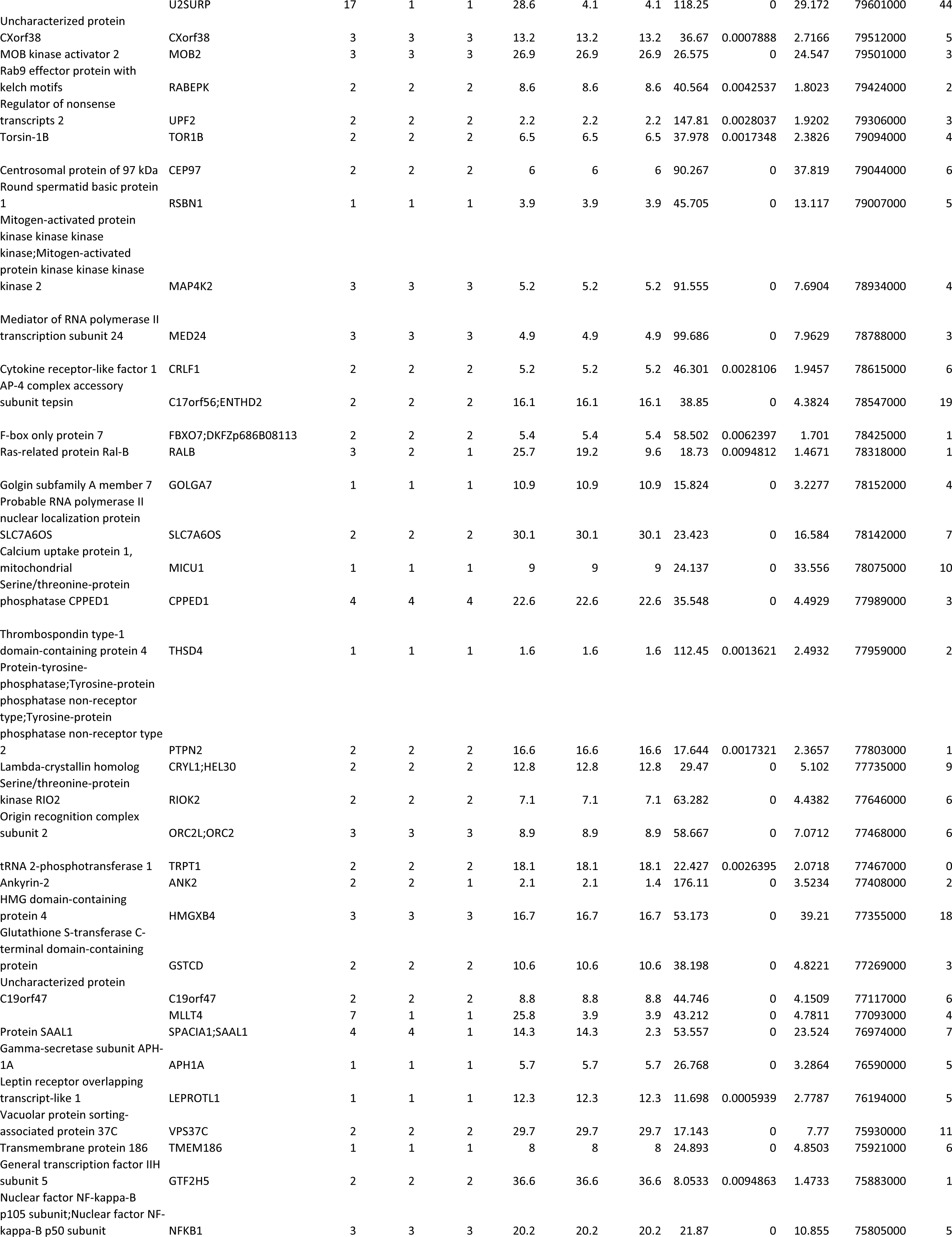
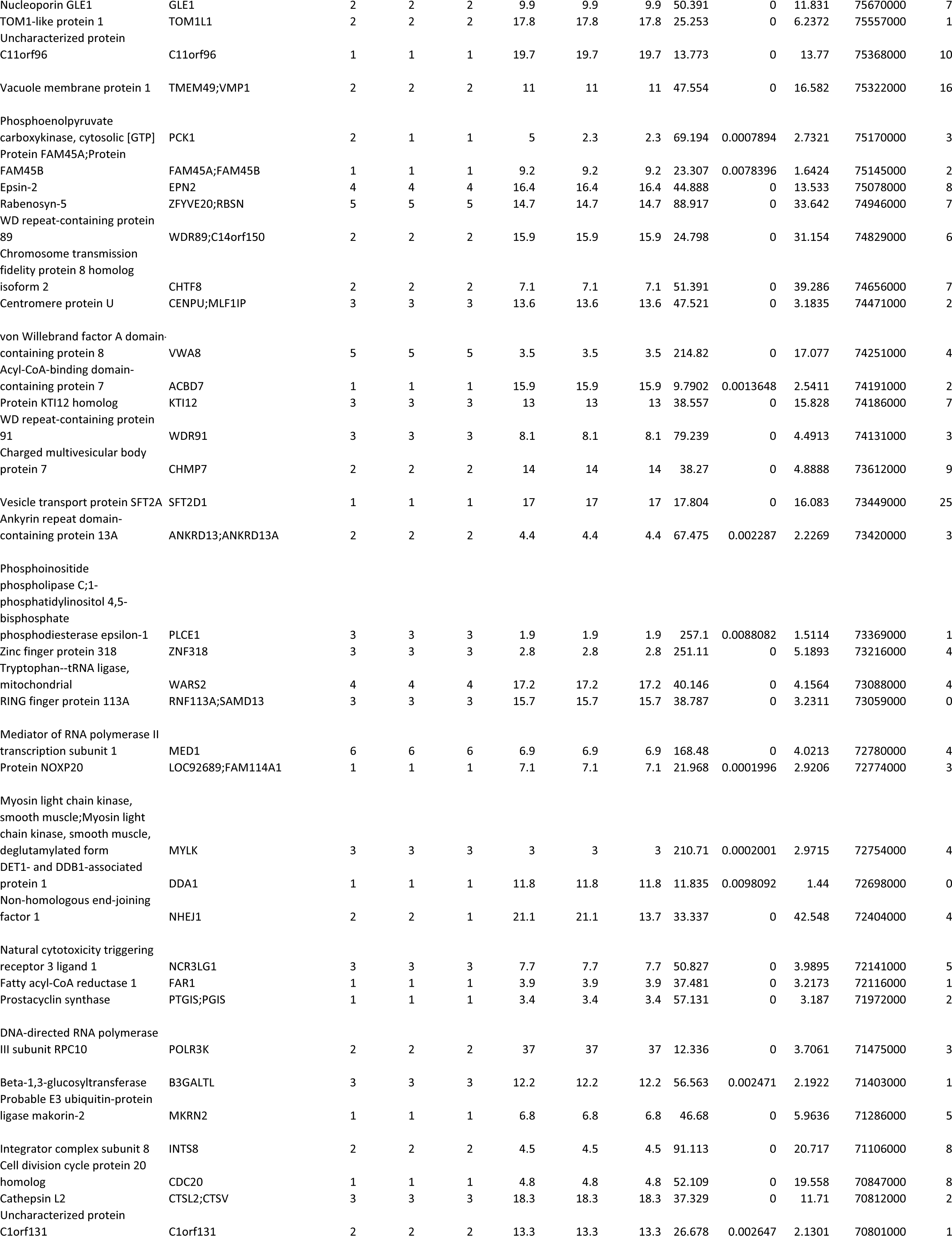
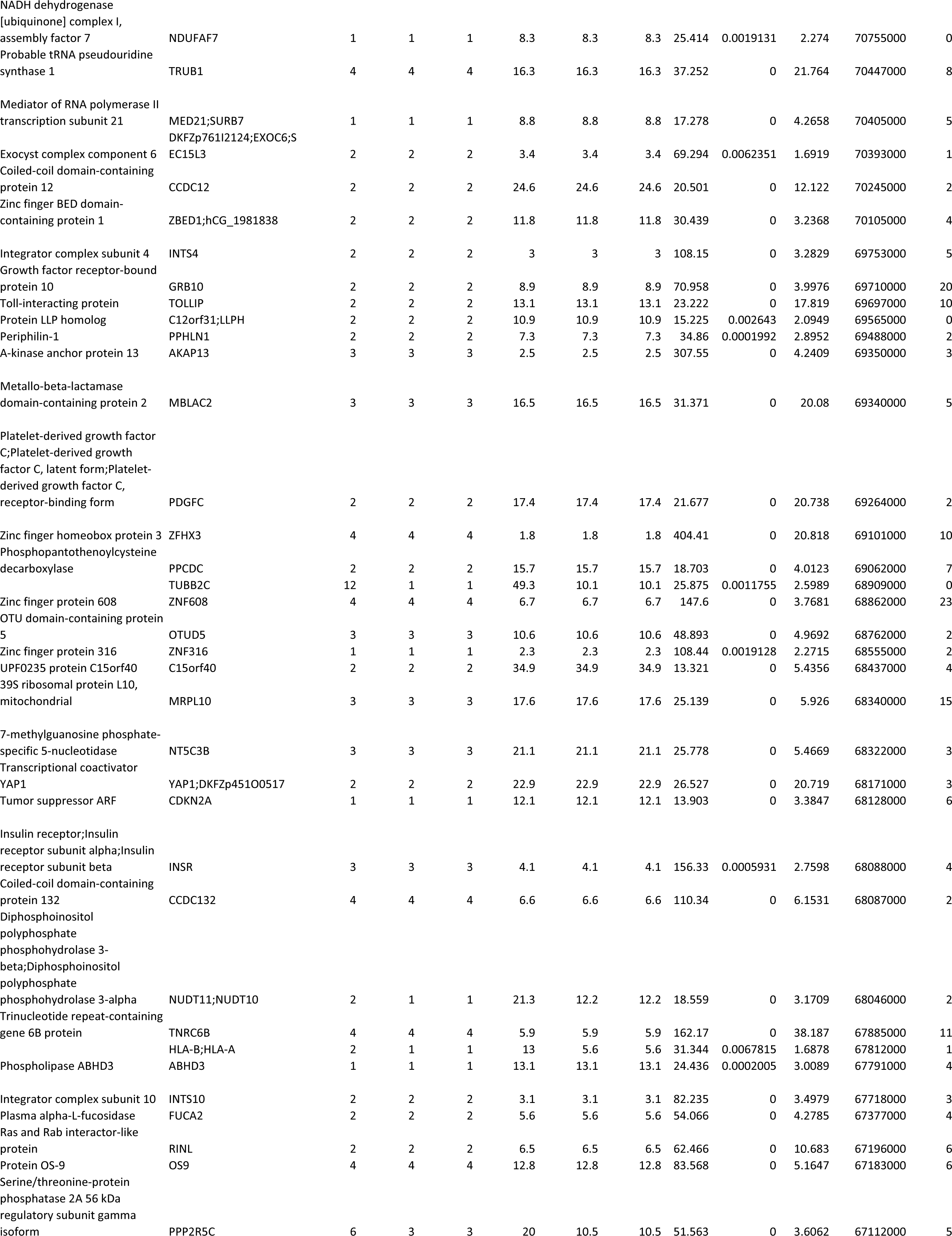
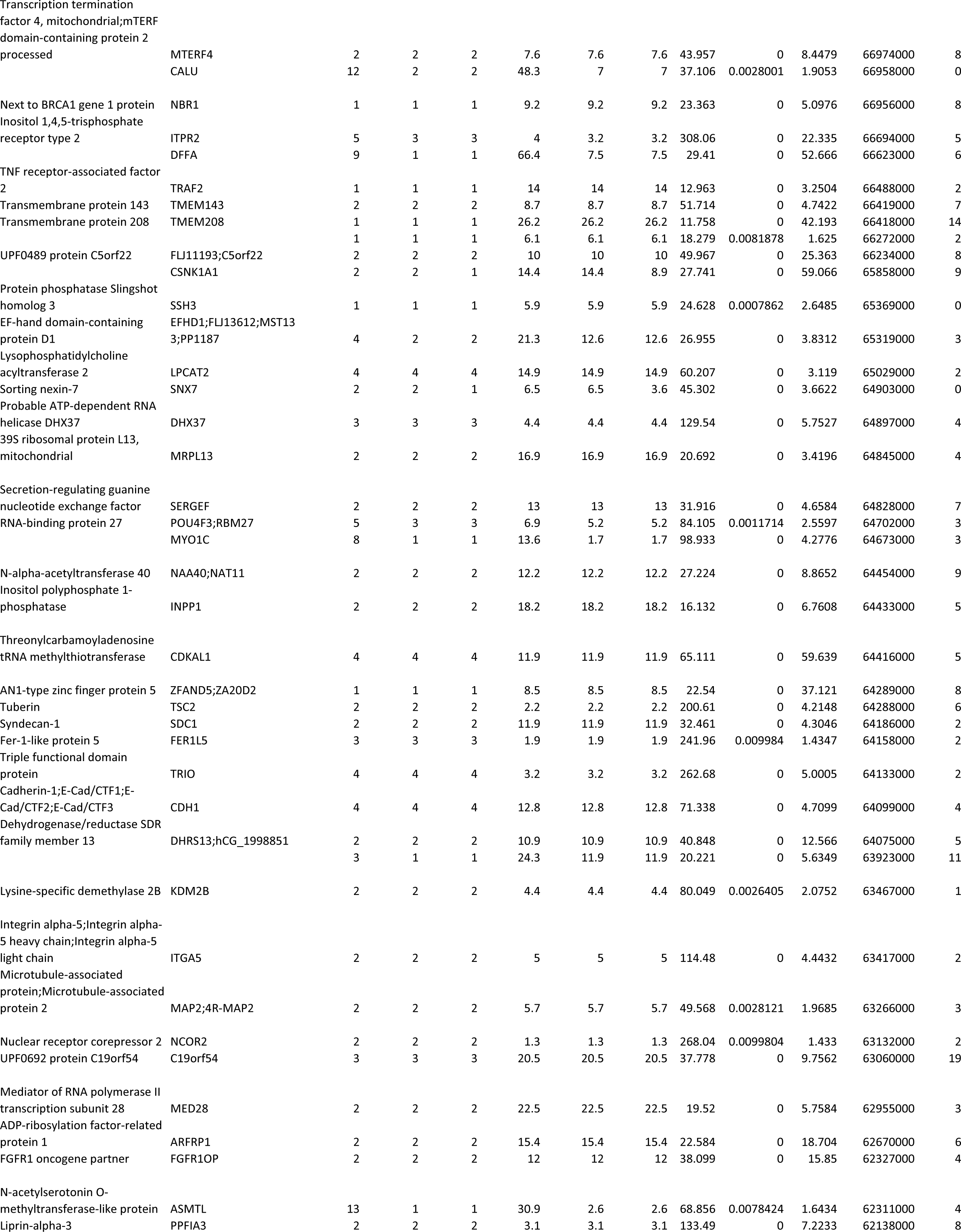
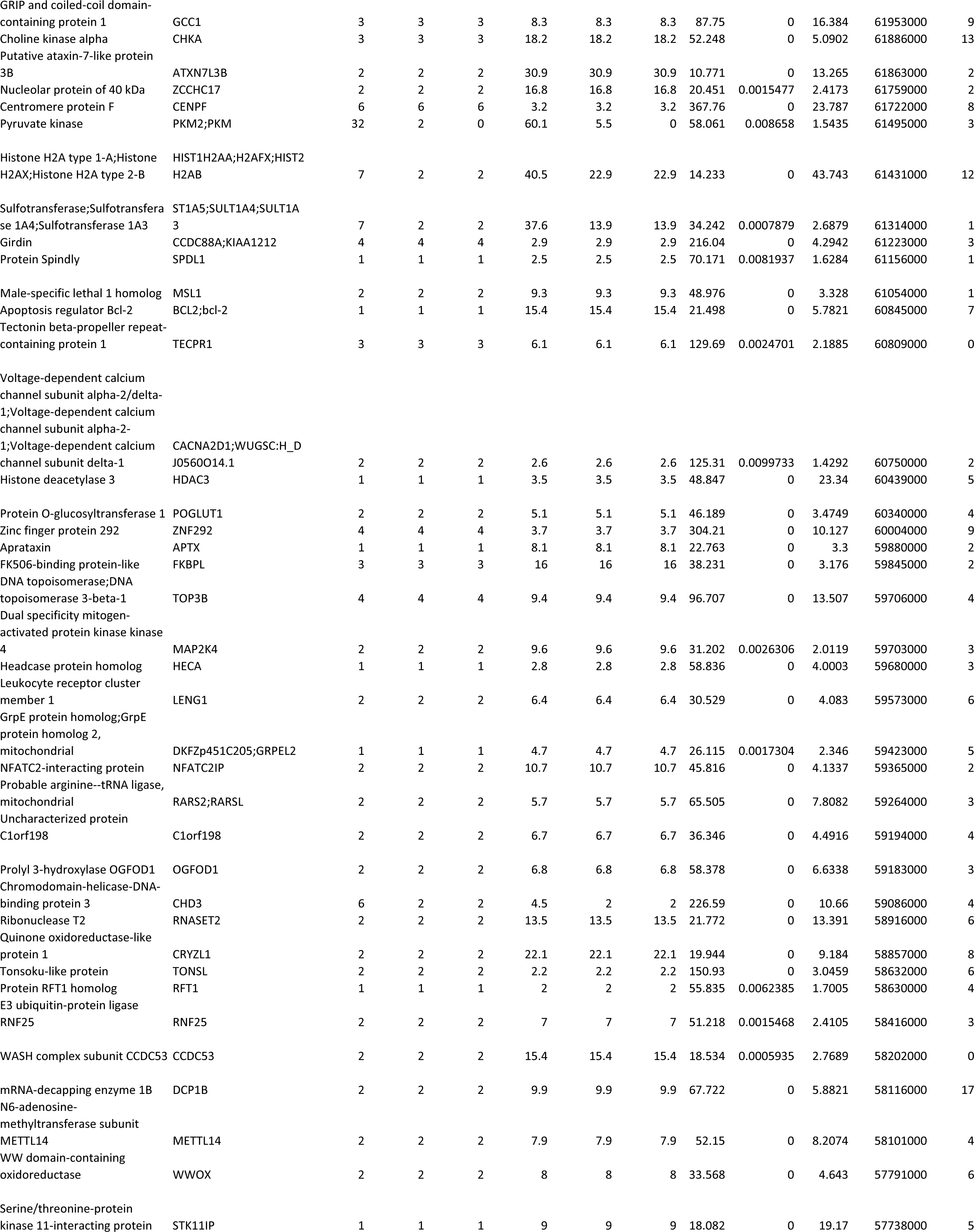
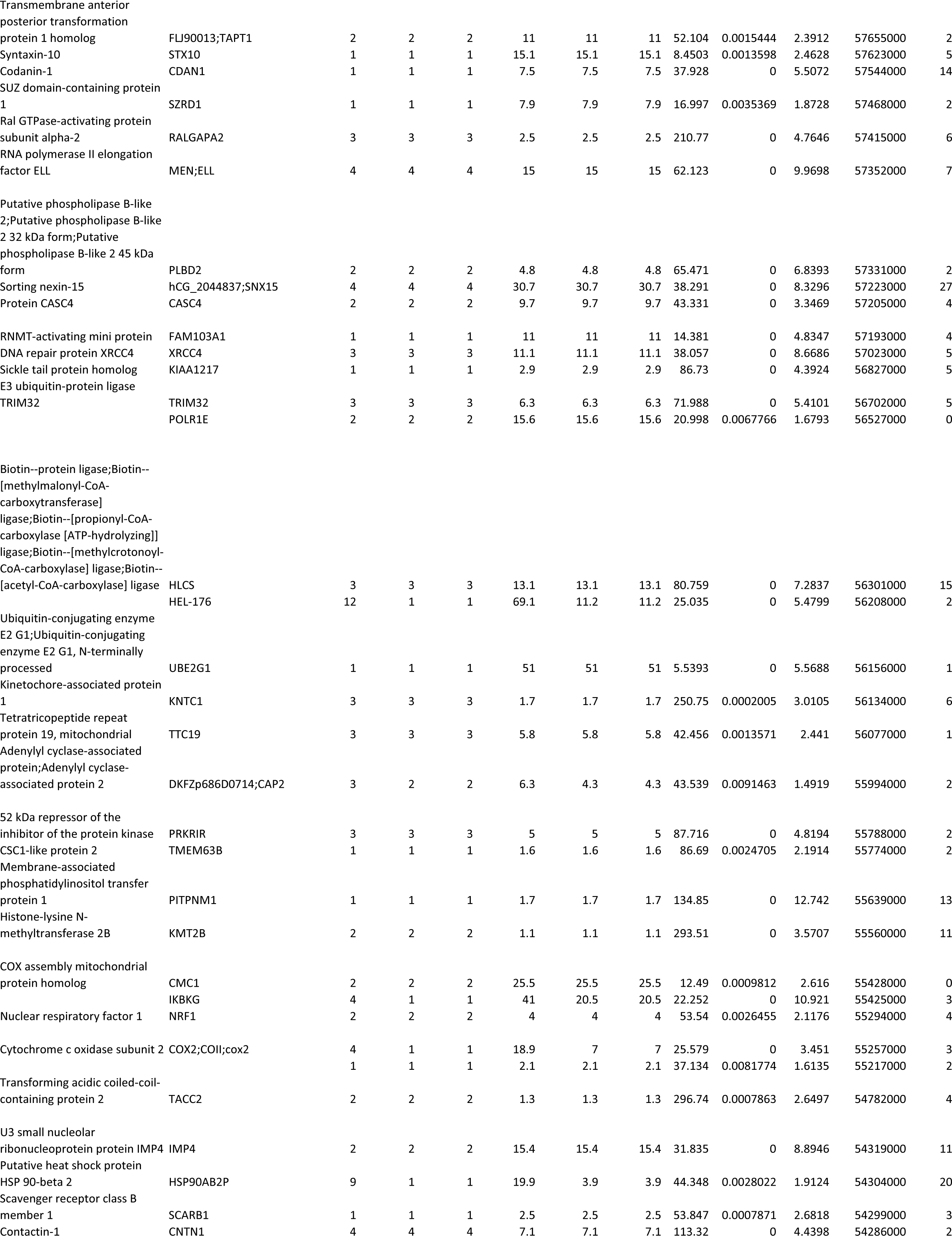
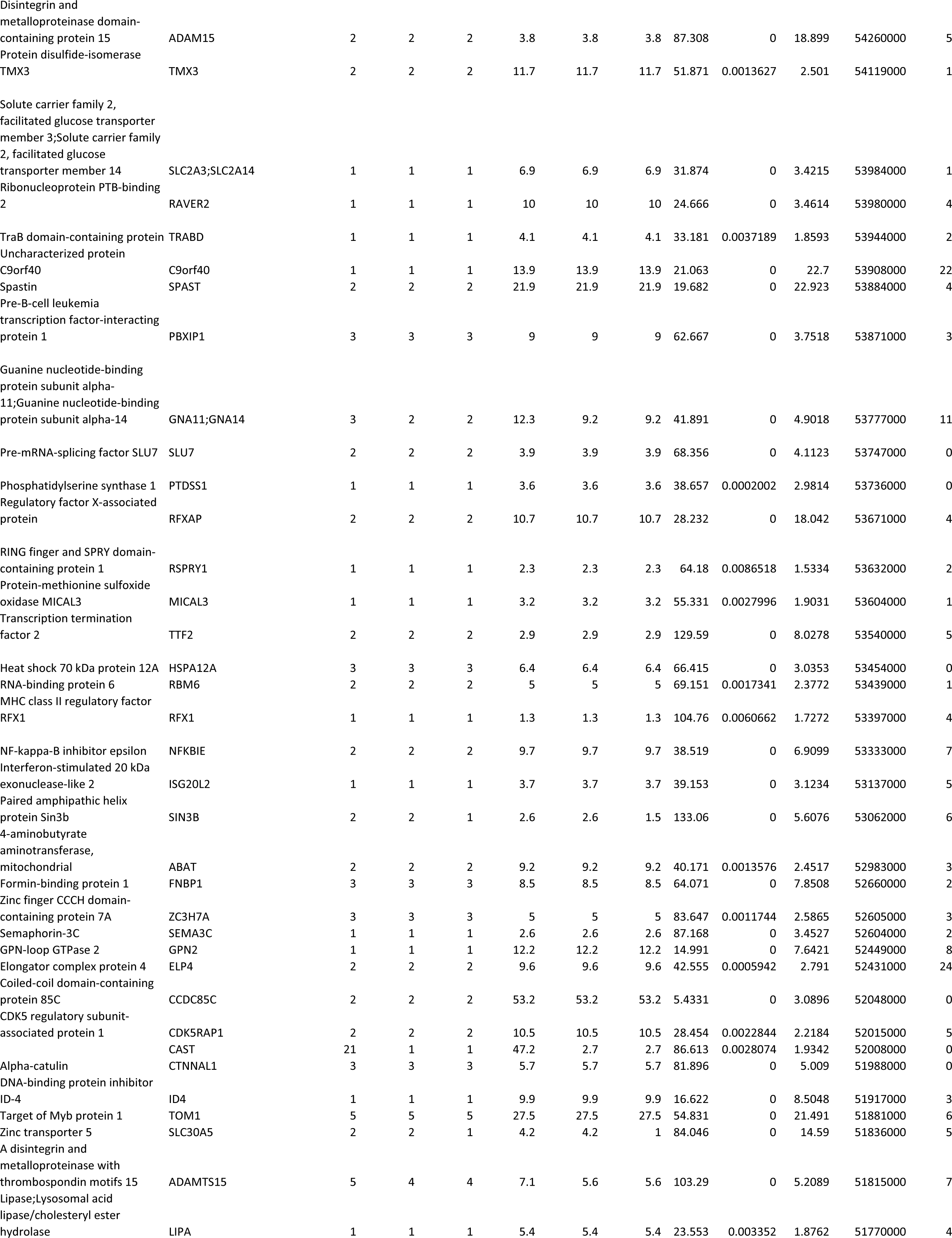
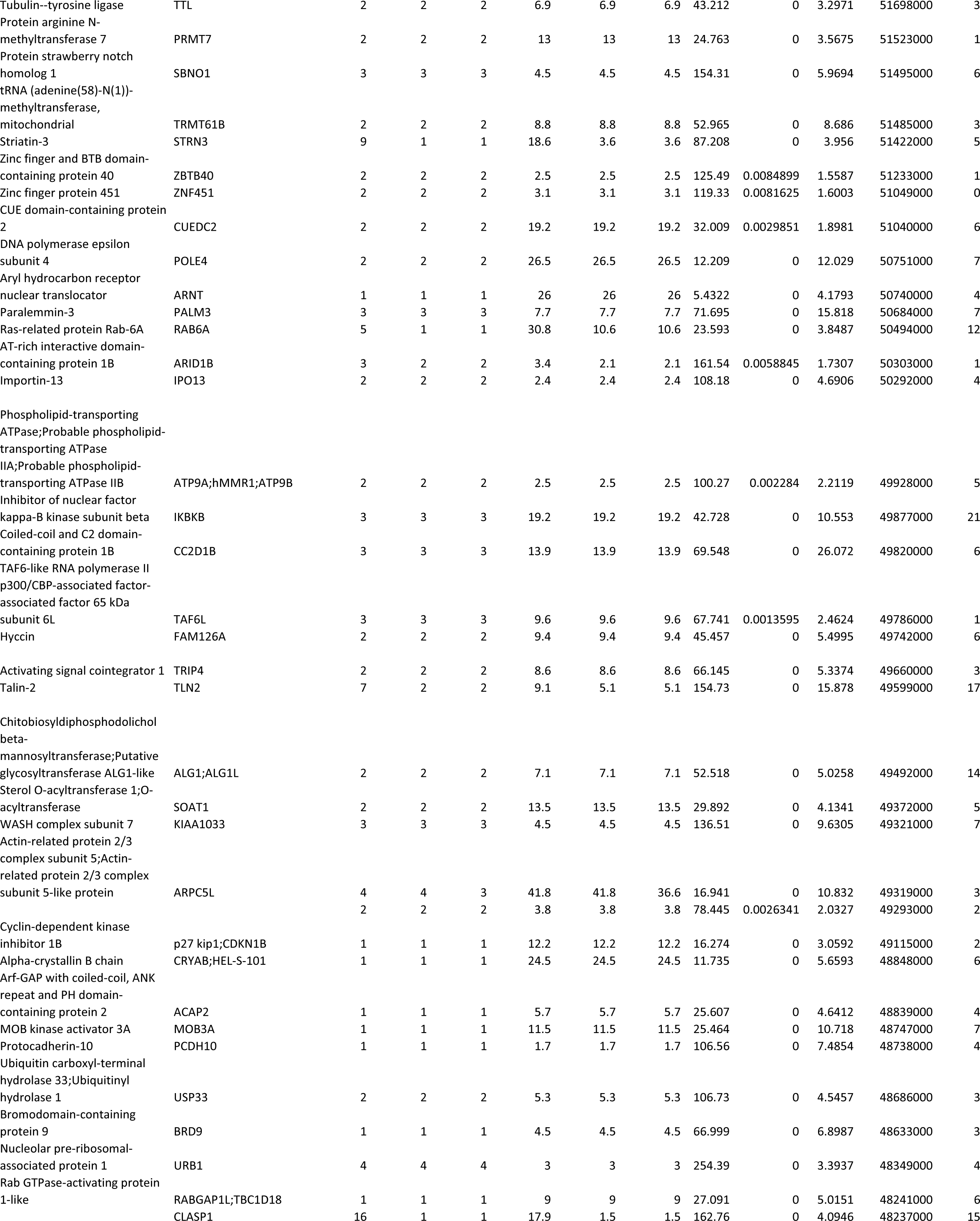
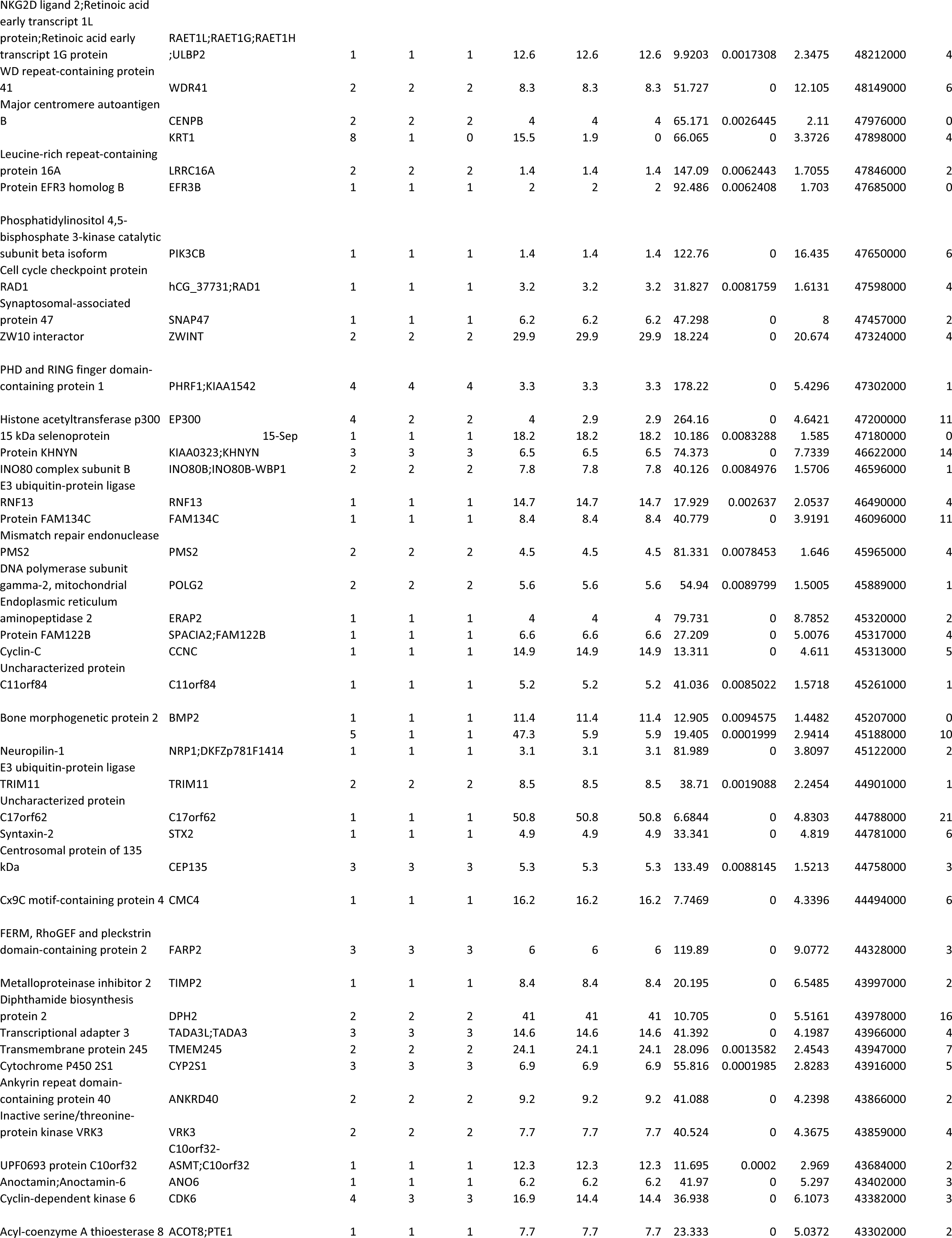
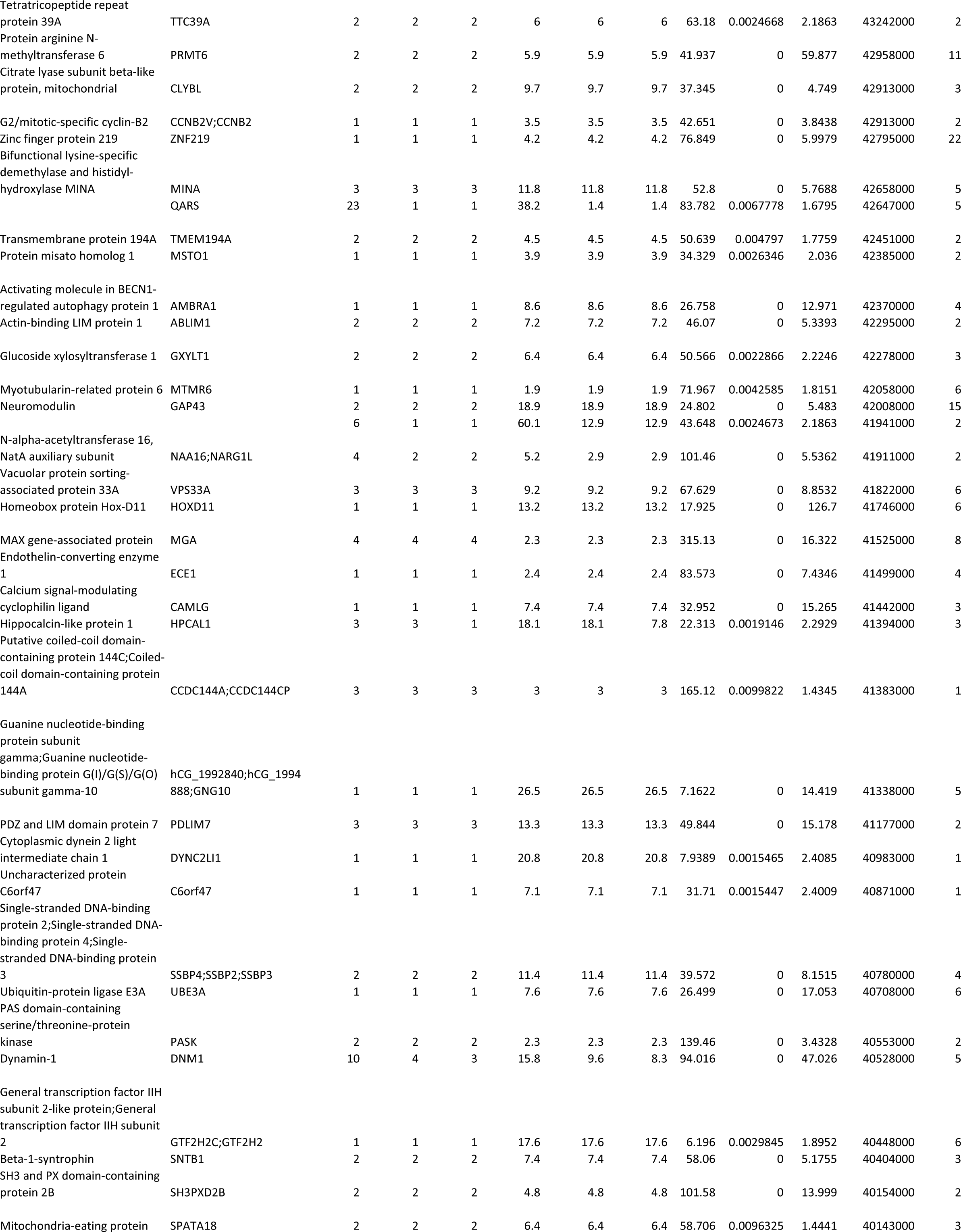
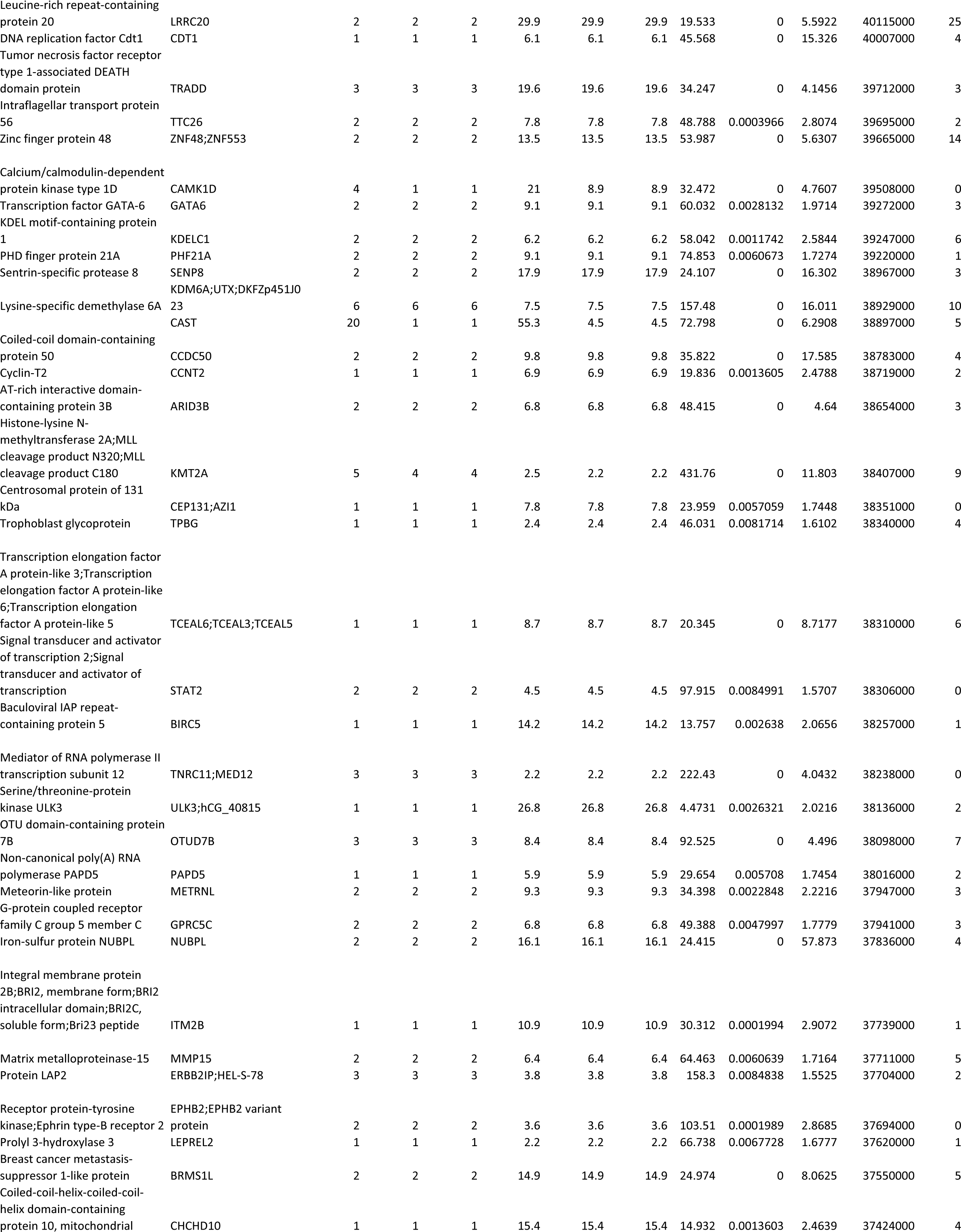
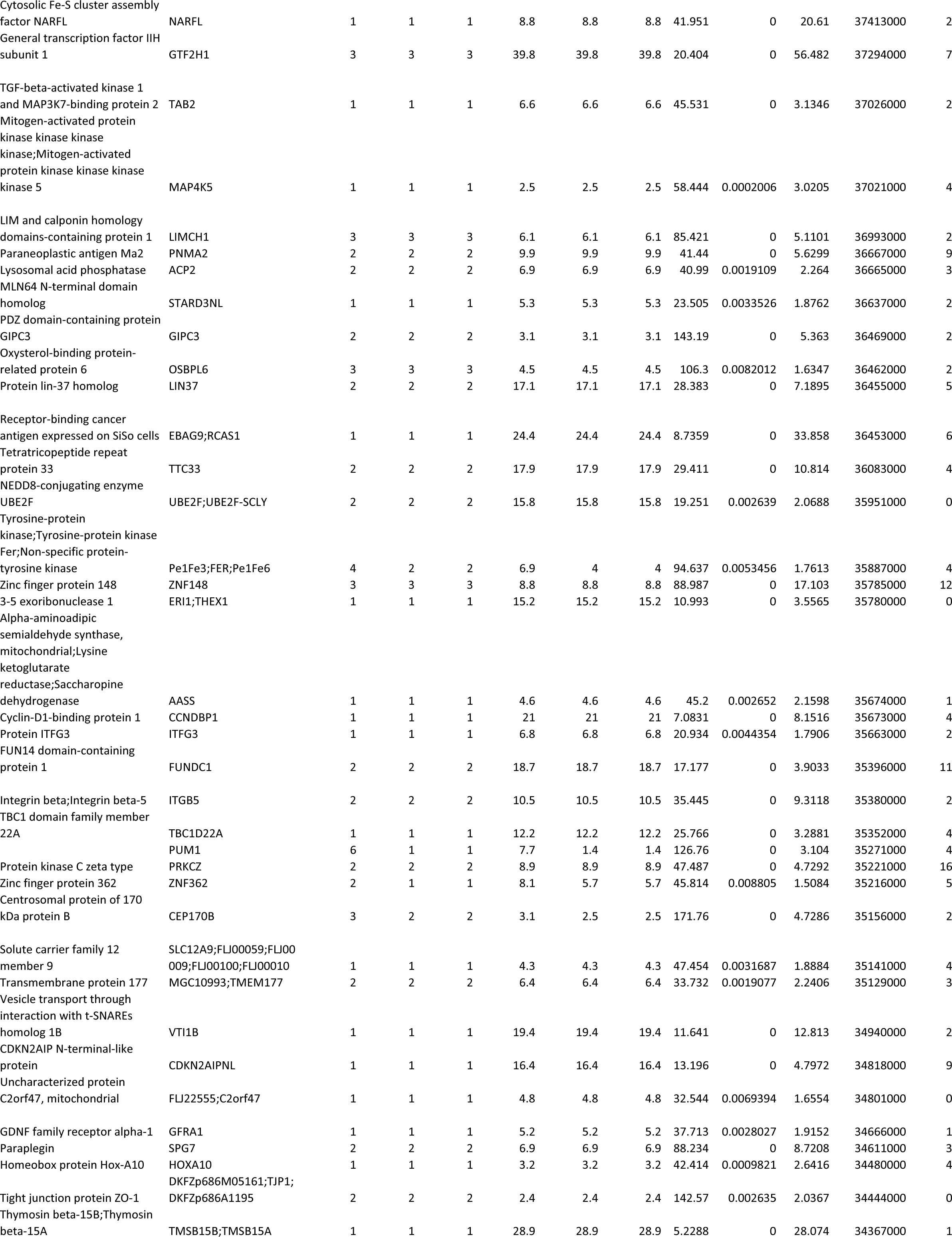
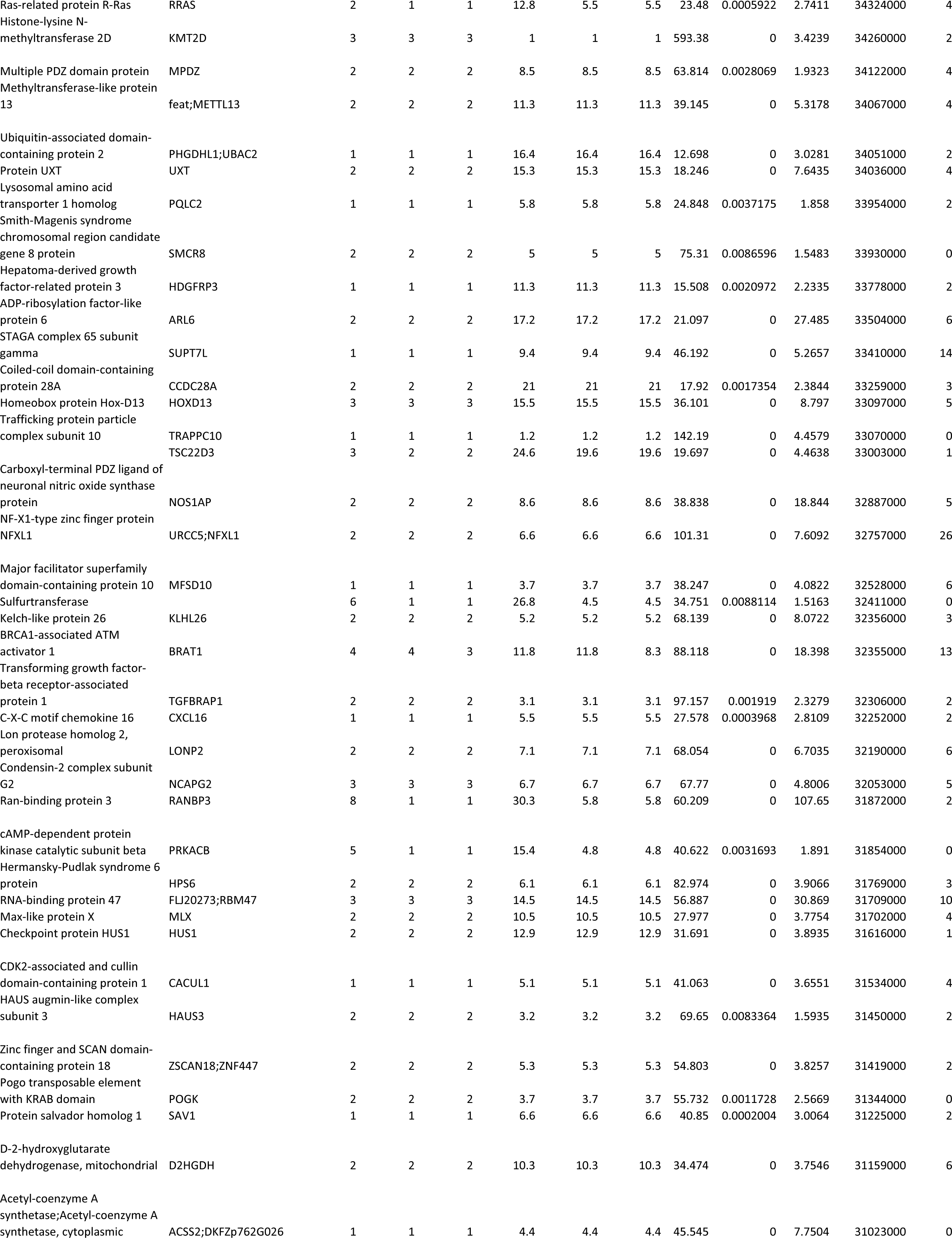
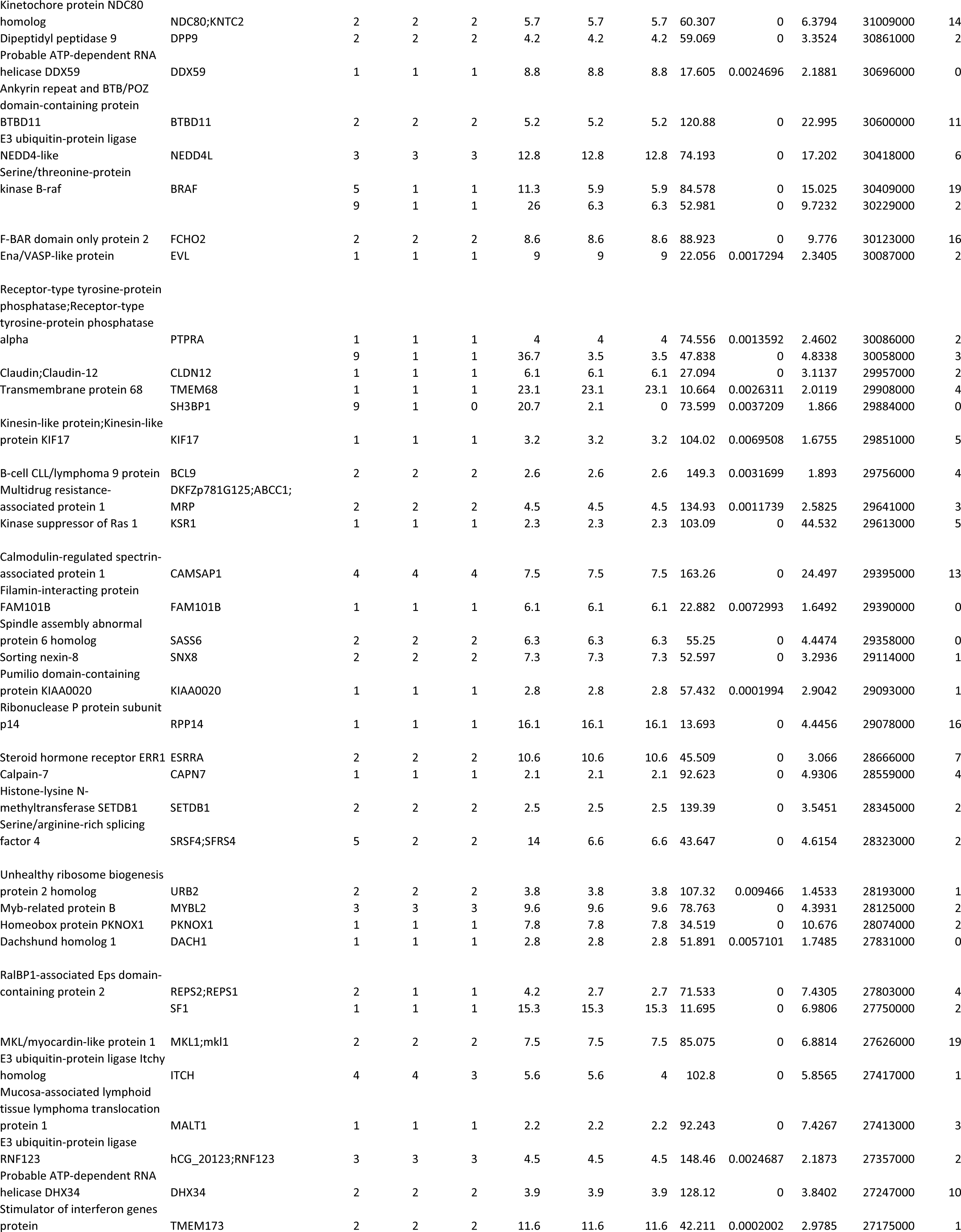
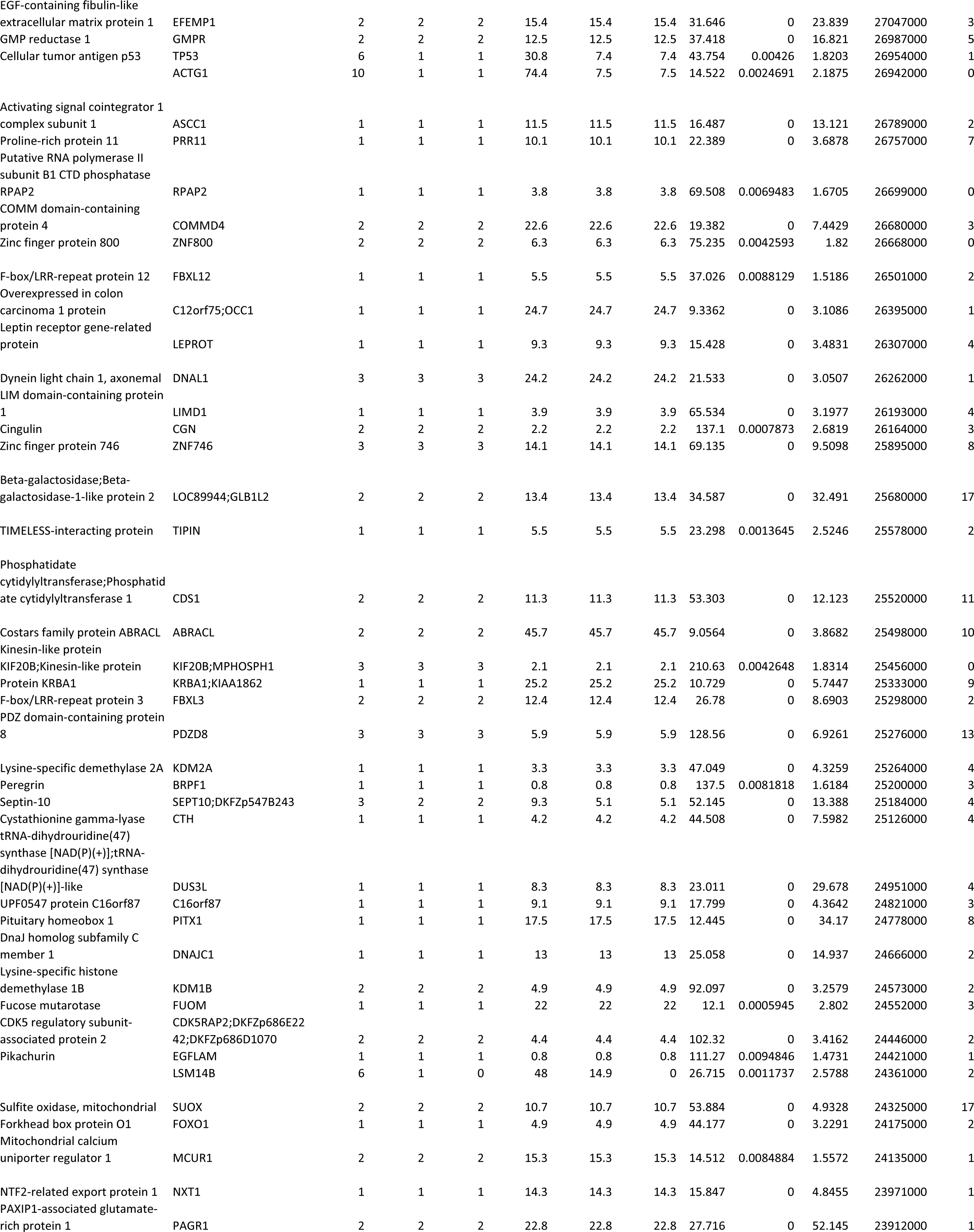
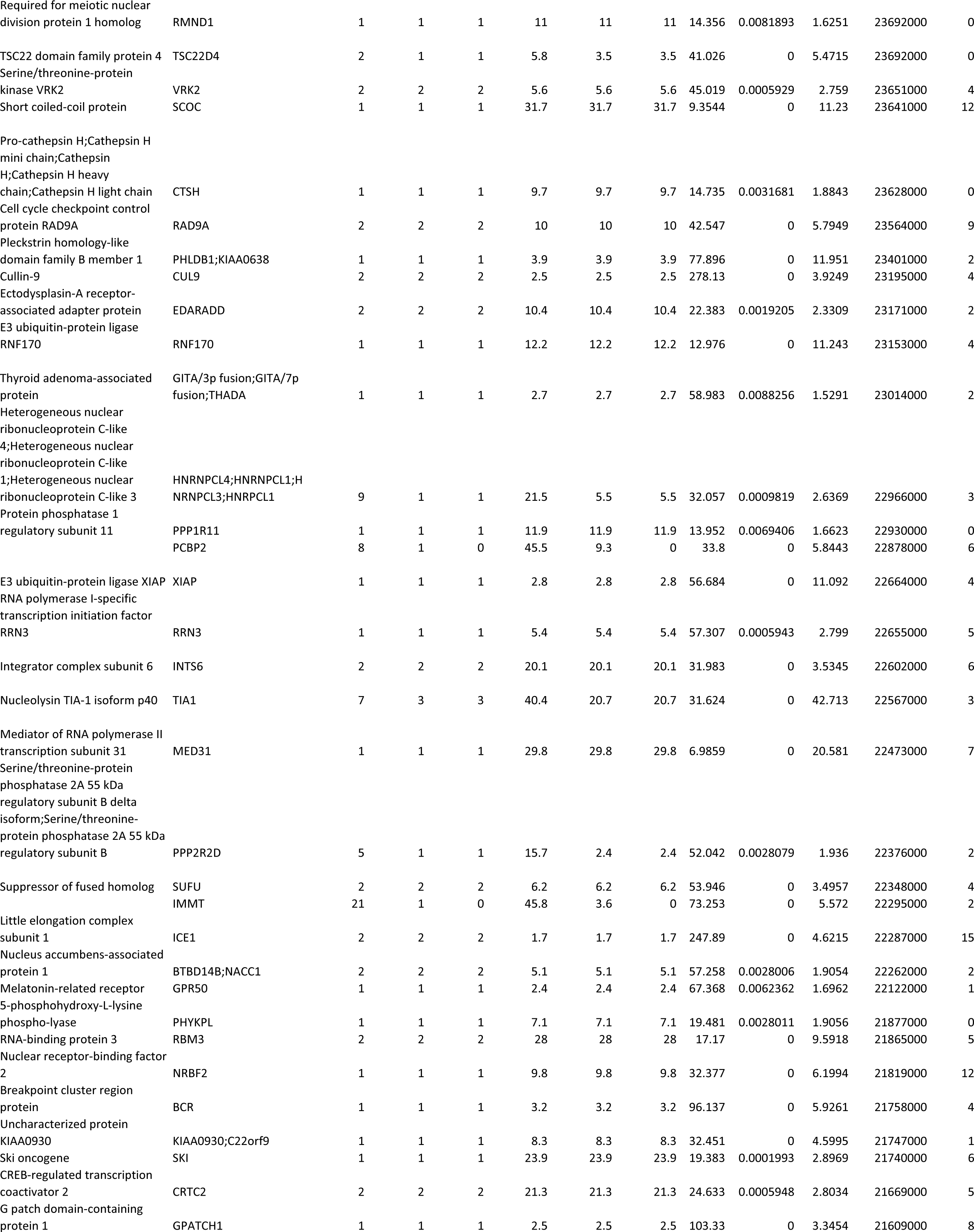
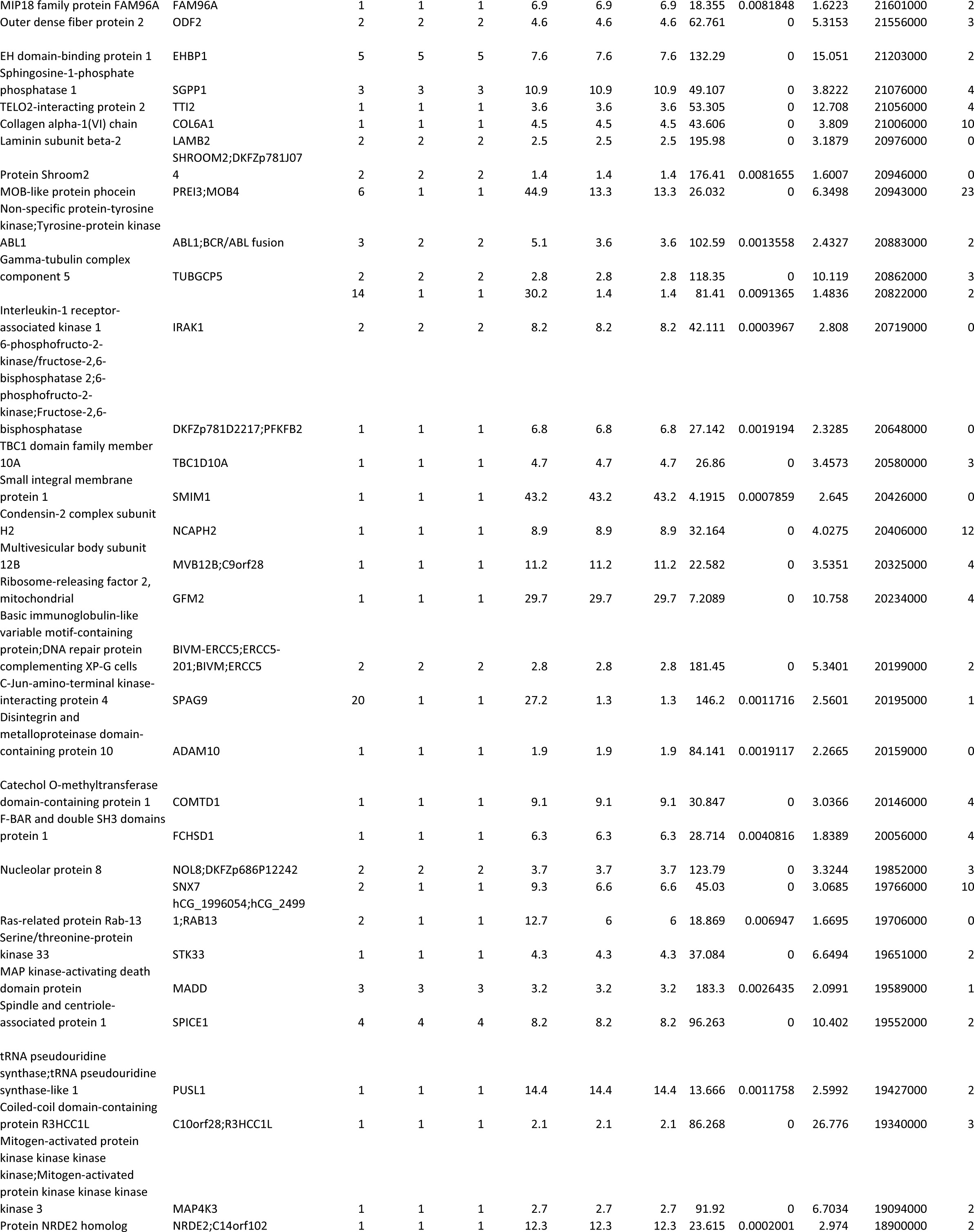
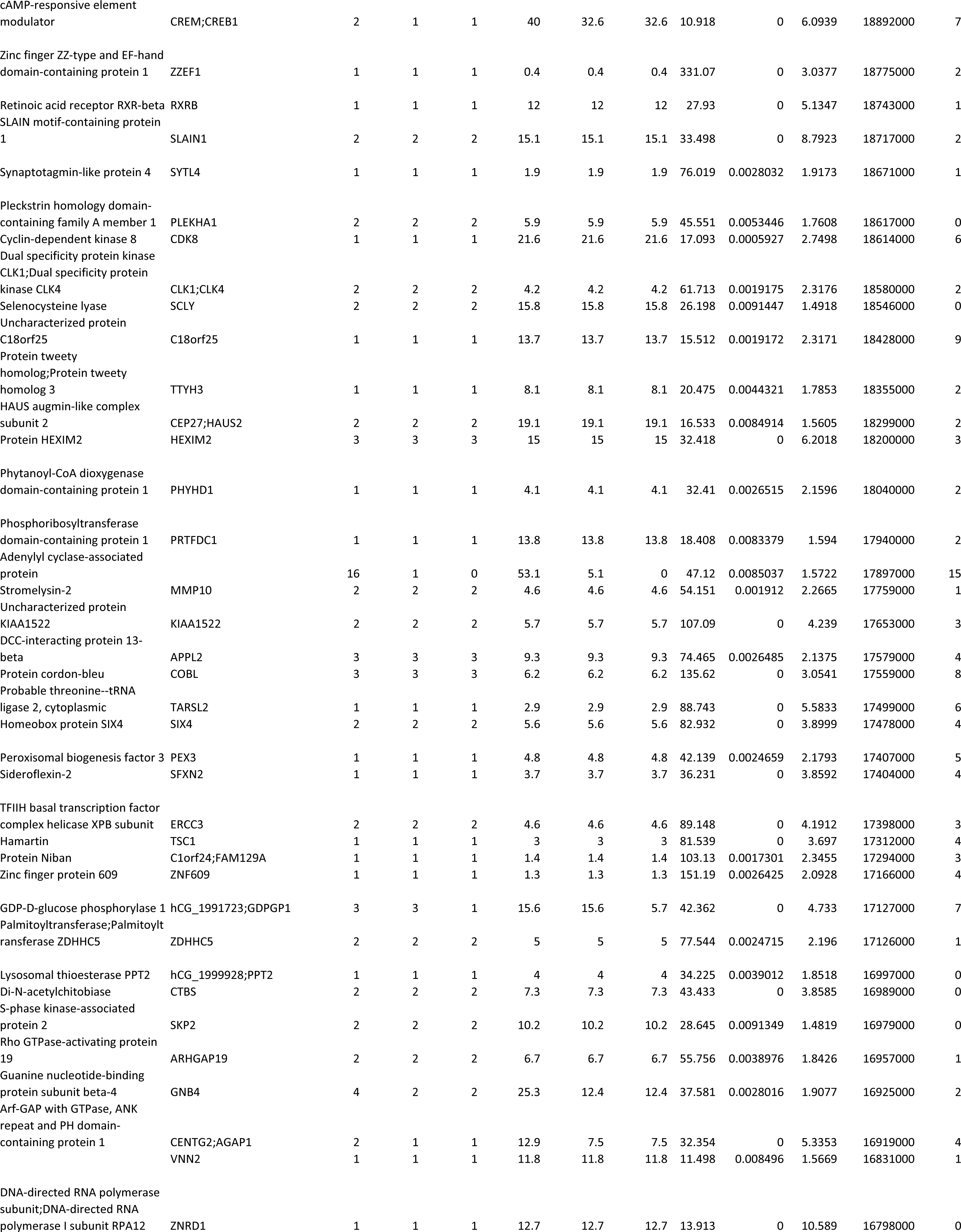
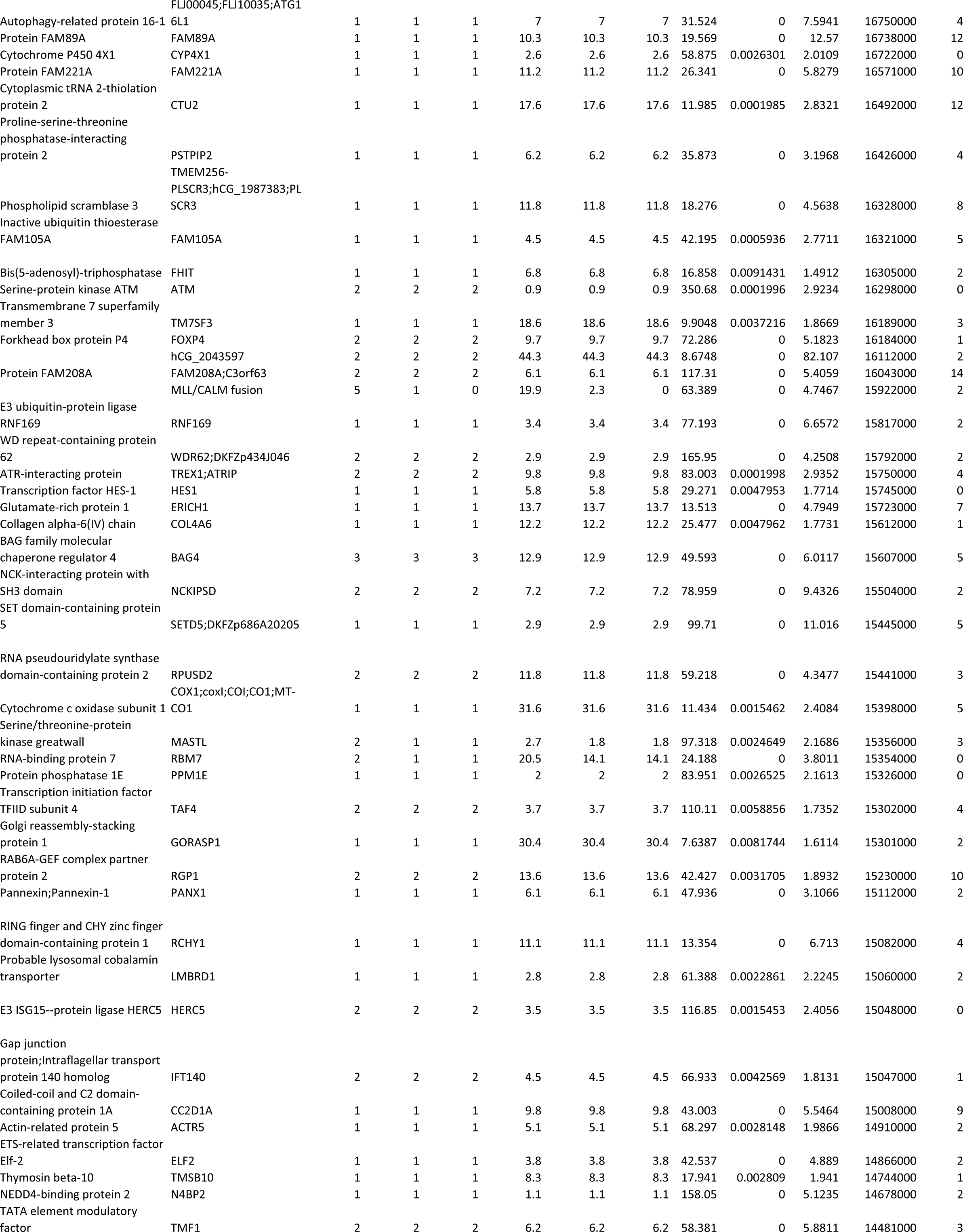
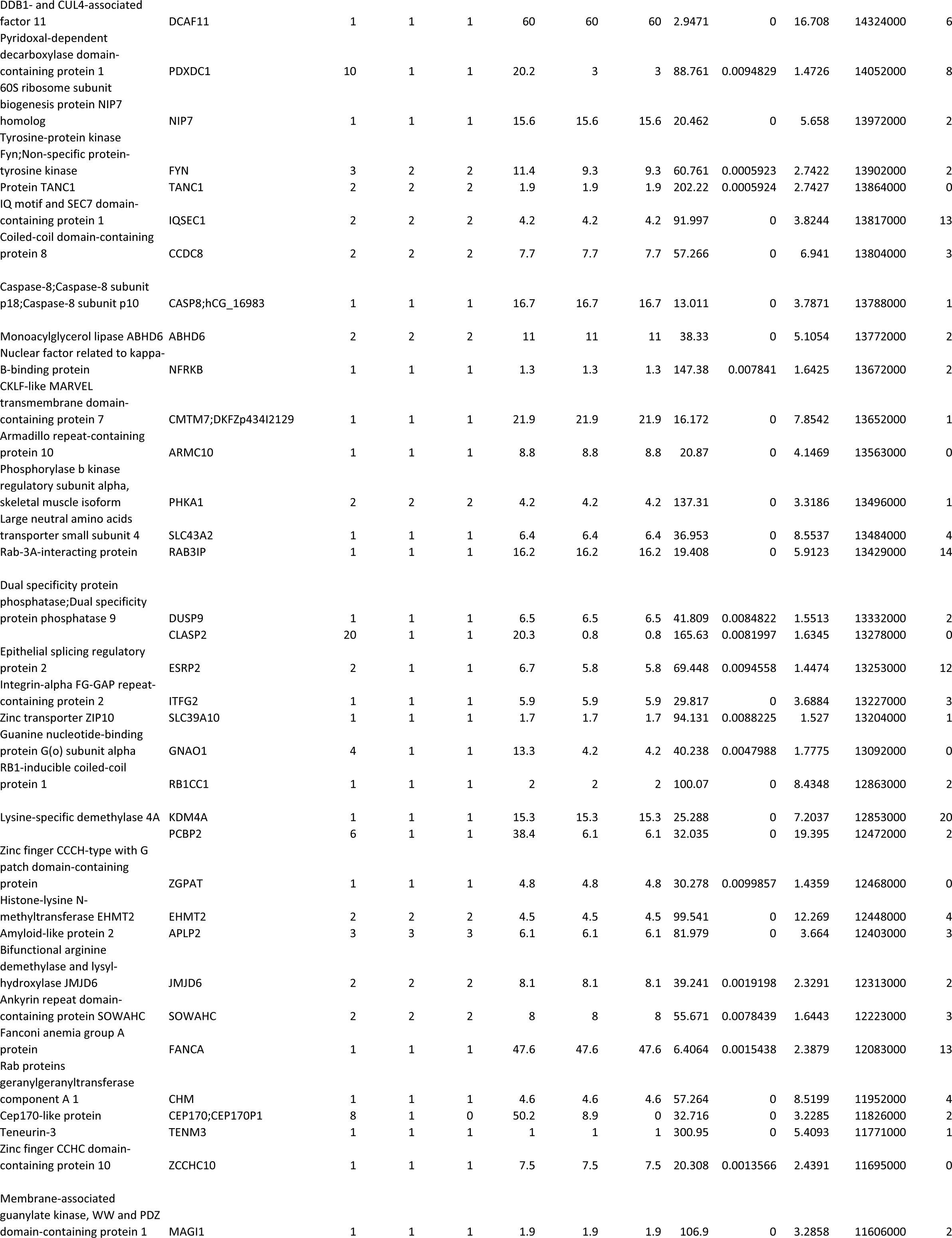
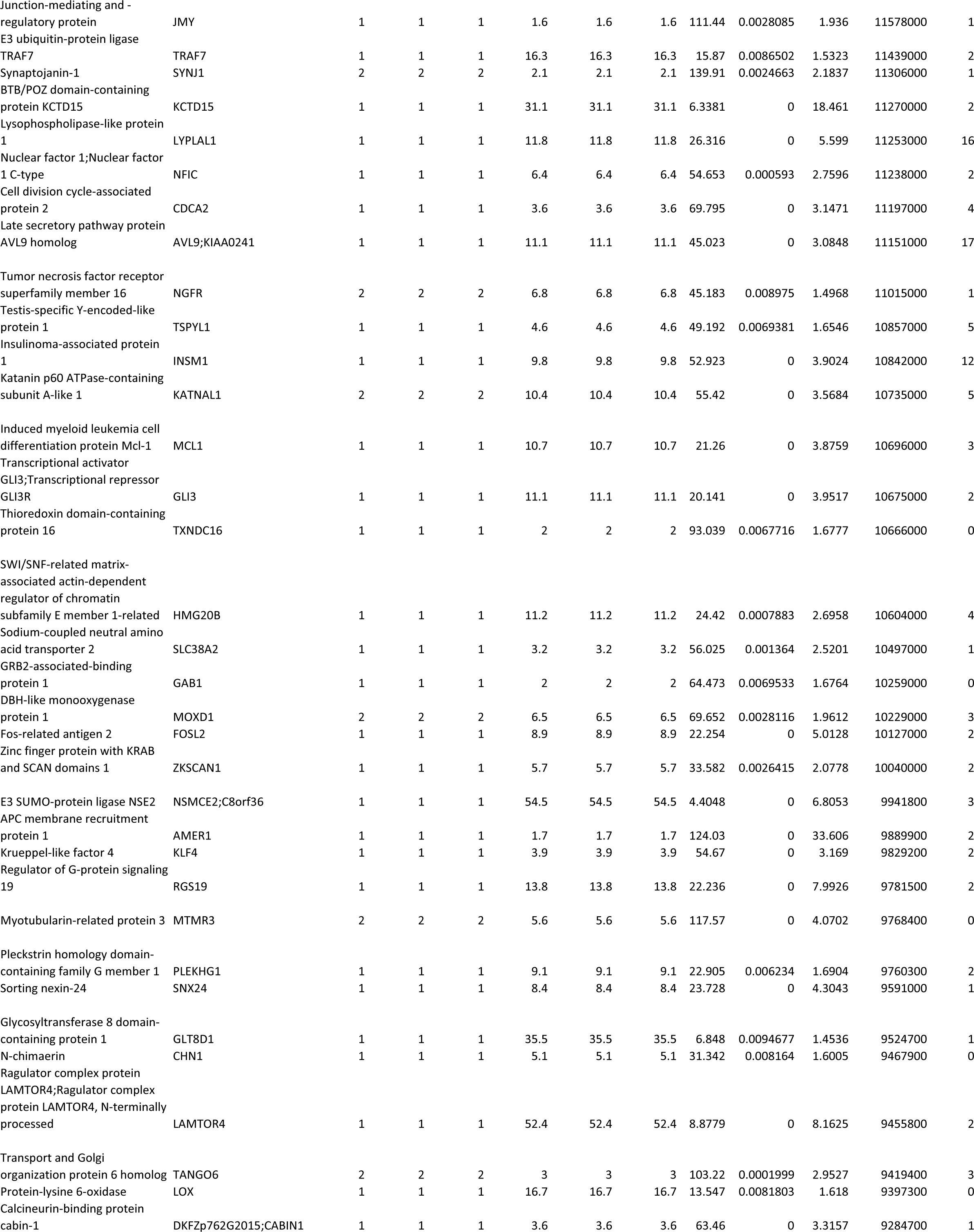
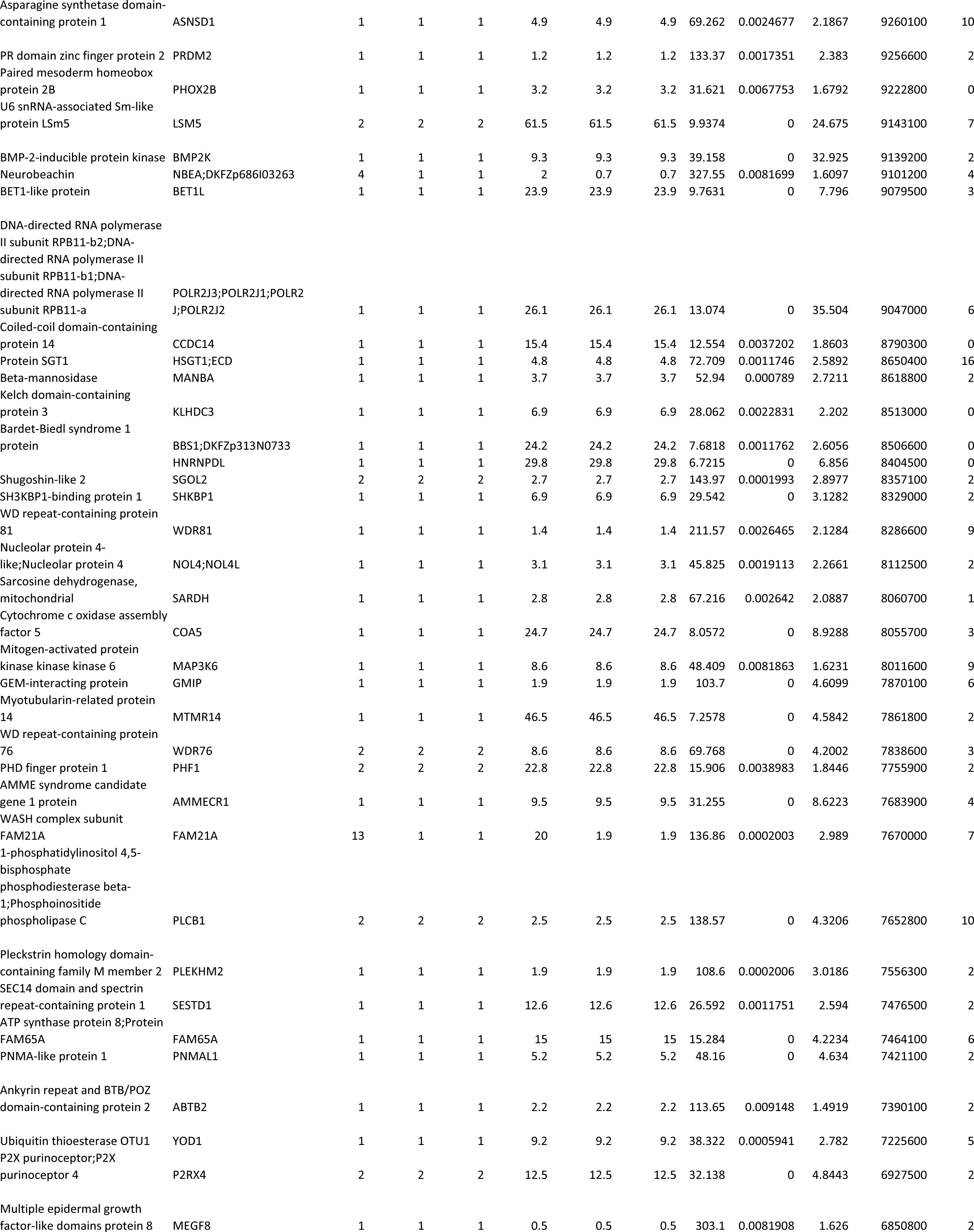
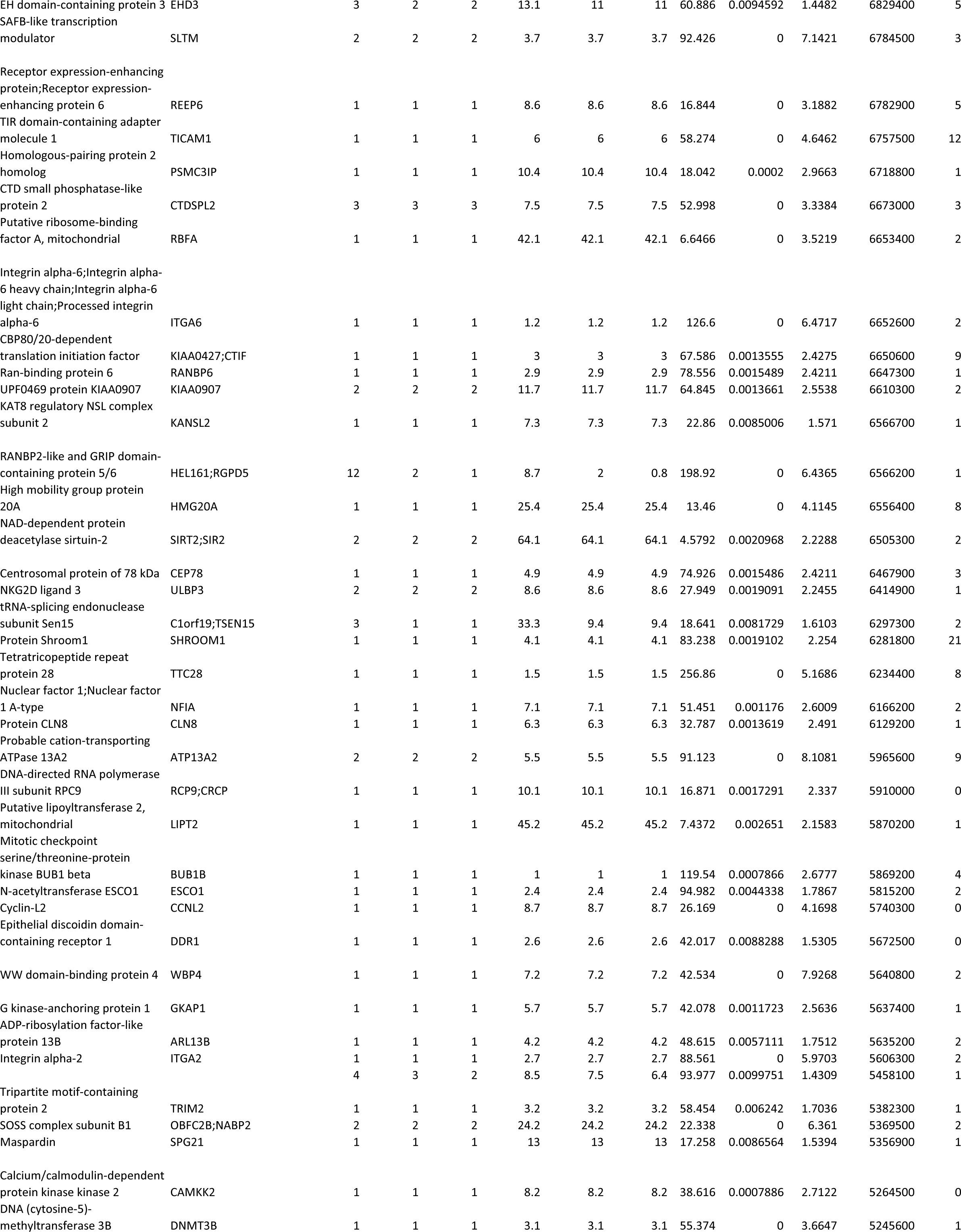
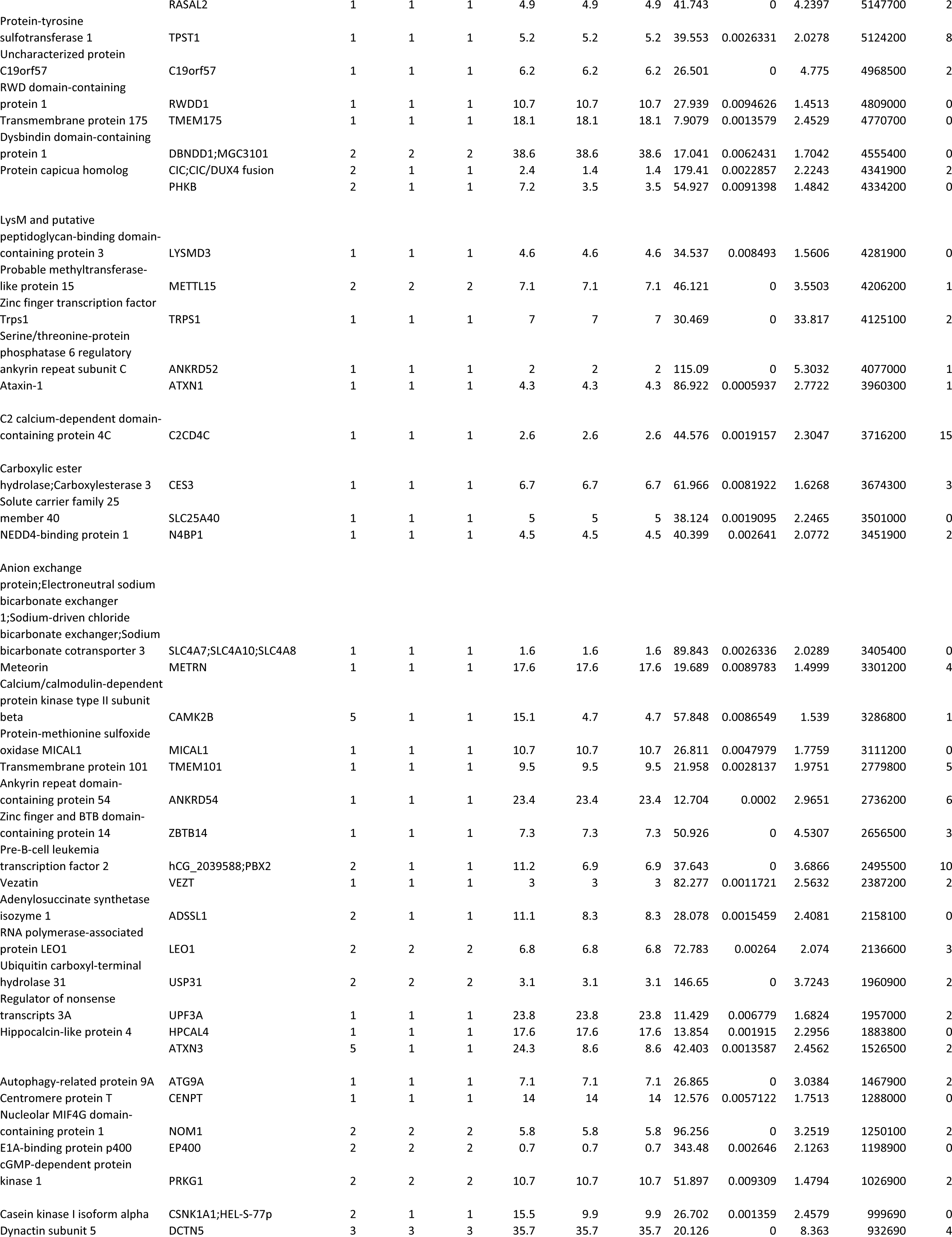
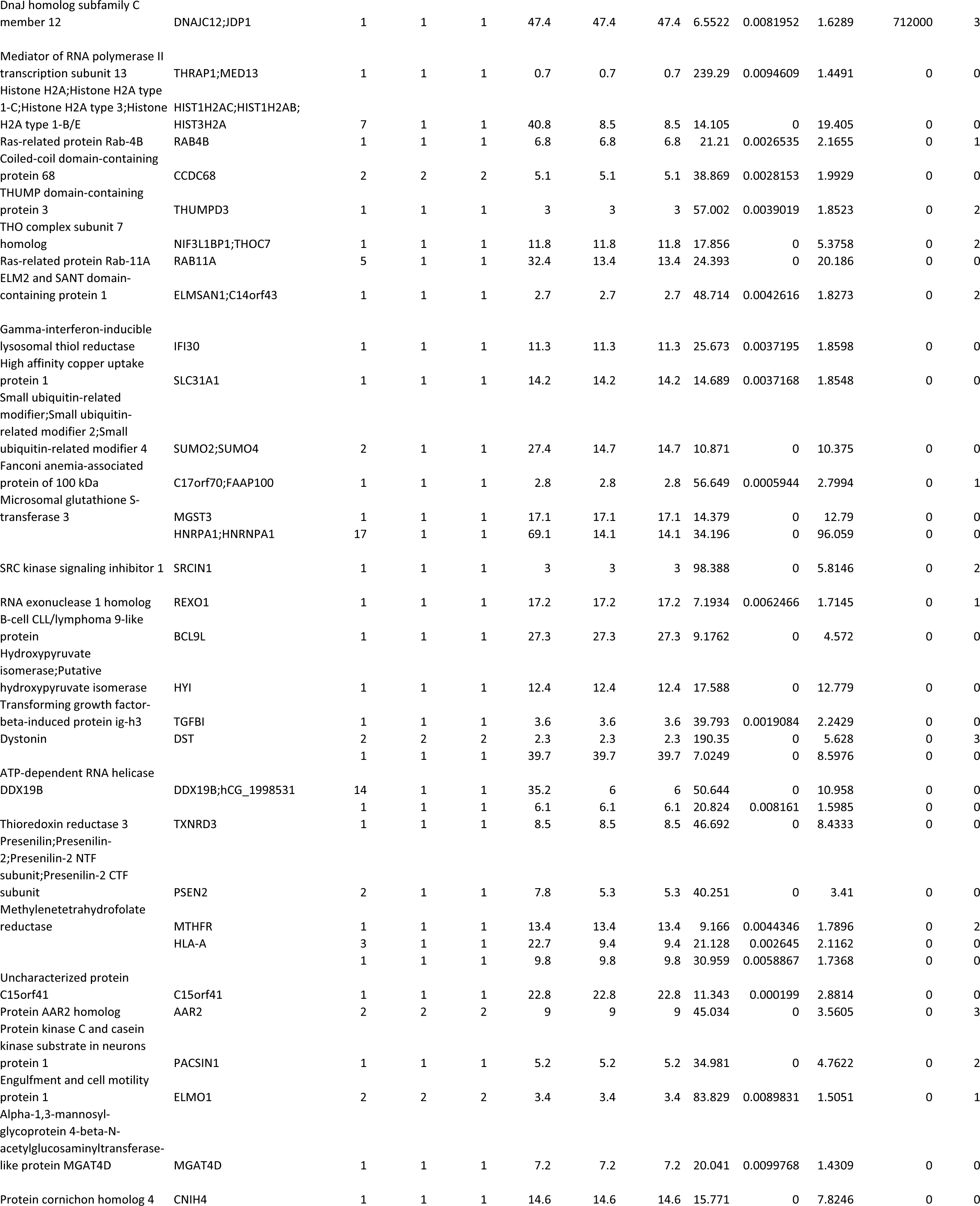
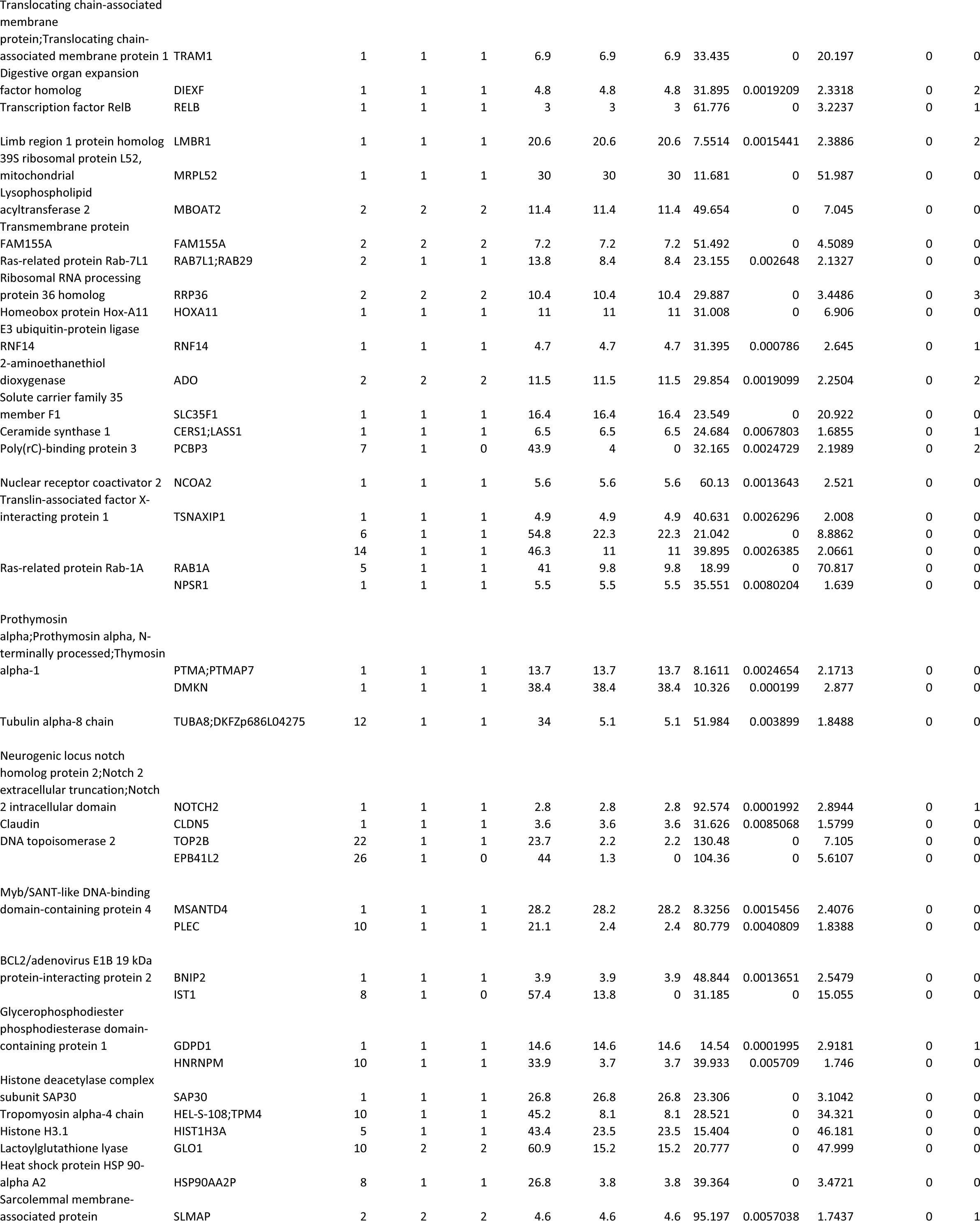
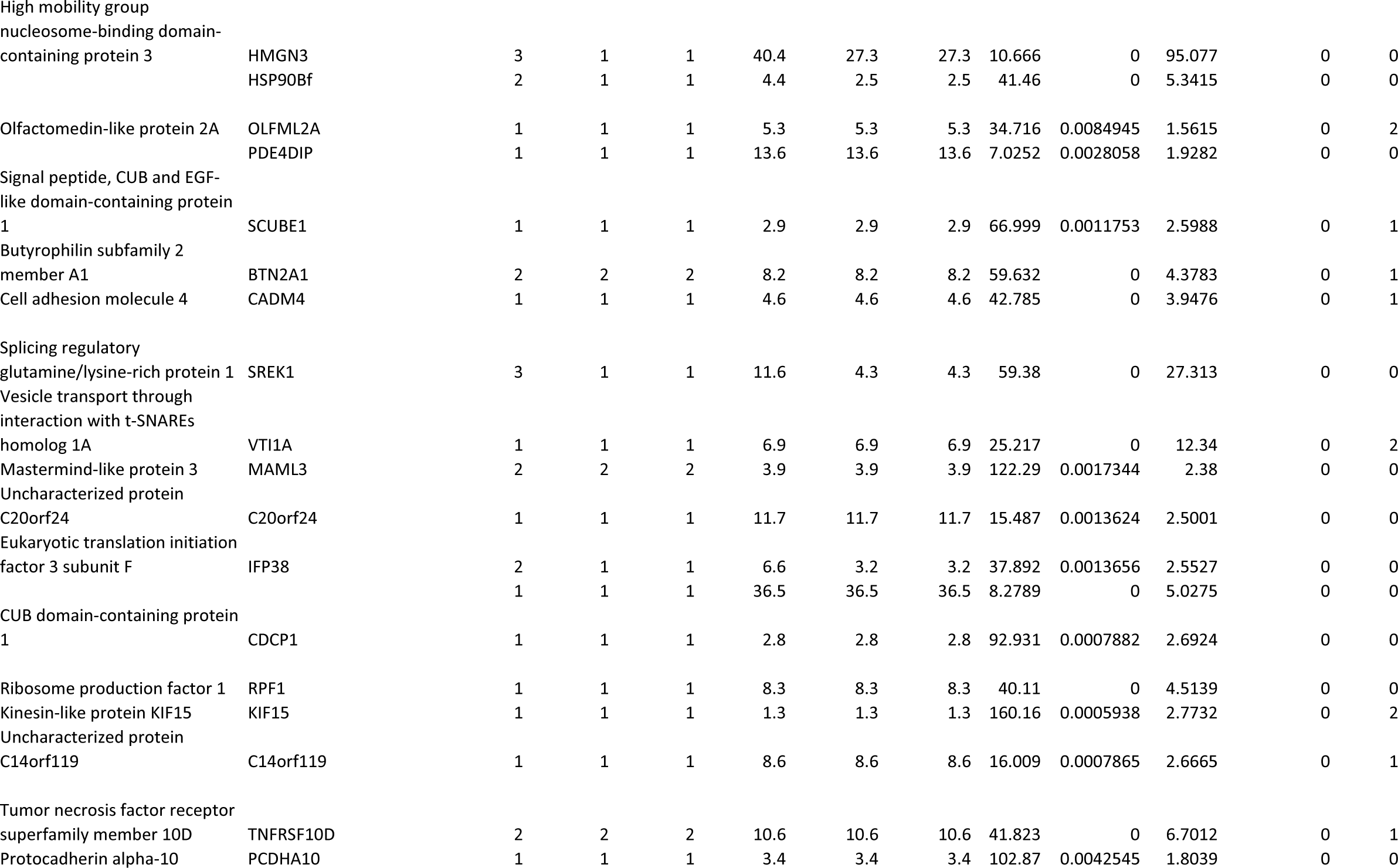

## References

1. T. Yonekawa, A. J. Rauckhorst, S. El-Hattab, M. A. Cuellar, D. Venzke, M. E. Anderson, H. Okuma, A. D. Pewa, E. B. Taylor, K. P. Campbell, Large1 gene transfer in older myd mice with severe muscular dystrophy restores muscle function and greatly improves survival. Sci Adv. 8, eabn0379 (2022).

2. A. S. Walimbe, H. Okuma, S. Joseph, T. Yang, T. Yonekawa, J. M. Hord, D. Venzke, M. E. Anderson, S. Torelli, A. Manzur, M. Devereaux, M. Cuellar, S. Prouty, S. O. Landa, L. Yu, J. Xiao, J. E. Dixon, F. Muntoni, K. P. Campbell, POMK regulates dystroglycan function via LARGE-mediated elongation of matriglycan. Elife. 9, e61388 (2020).

54. S. Joseph, N. J. Schnicker, Z. Xu, T. Yang, J. Hopkins, M. Watkins, S. Chakravarthy, O. Davulcu, M. E. Anderson, D. Venzke, K. P. Campbell, bioRxiv, in press, doi:10.1101/2022.05.12.491222.

4. S. Kunz, N. Sevilla, D. B. McGavern, K. P. Campbell, M. B. A. Oldstone, Molecular analysis of the interaction of LCMV with its cellular receptor α-dystroglycan. J Cell Biology. 155, 301– 310 (2001).

5. M. Kanagawa, F. Saito, S. Kunz, T. Yoshida-Moriguchi, R. Barresi, Y. M. Kobayashi, J. Muschler, J. P. Dumanski, D. E. Michele, M. B. A. Oldstone, K. P. Campbell, Molecular recognition by LARGE is essential for expression of functional dystroglycan. Cell. 117, 953–964 (2004).

6. C. A. Pinkert, Transgenic Animal Technology, 3–12 (2002).

7. J. Jang-Lee, S. J. North, M. S. Smith, D. Goldberg, M. Panico, H. Morris, S. Haslam, A. Dell, Glycomic profiling of cells and tissues by mass spectrometry: fingerprinting and sequencing methodologies. Methods in Enzymology. 415, 59–86 (2006).

8. H. Zhang, F. Zhu, T. Yang, L. Ding, M. Zhou, J. Li, S. M. Haslam, A. Dell, H. Erlandsen, H. Wu, The highly conserved domain of unknown function 1792 has a distinct glycosyltransferase fold. Nature communications. 5, 4339 (2014).

9. A. Shevchenko, H. Tomas, J. Havlis, J. V. Olsen, M. Mann, In-gel digestion for mass spectrometric characterization of proteins and proteomes. Nature Protocols. 1, 2856–2860 (2006).

10. J. R. Wiśniewski, A. Zougman, N. Nagaraj, M. Mann, Universal sample preparation method for proteome analysis. Nature Methods. 6, 359–362 (2009).

11. J. Cox, N. Neuhauser, A. Michalski, R. A. Scheltema, J. V. Olsen, M. Mann, Andromeda: a peptide search engine integrated into the MaxQuant environment. Journal of Proteome Research. 10, 1794–1805 (2011).

12. J. Cox, M. Y. Hein, C. A. Luber, I. Paron, N. Nagaraj, M. Mann, Accurate proteome-wide label-free quantification by delayed normalization and maximal peptide ratio extraction, termed MaxLFQ. Molecular & Cellular Proteomics. 13, 2513–2526 (2014).

13. S. Tyanova, R. Albrechtsen, P. Kronqvist, J. Cox, M. Mann, T. Geiger, Proteomic maps of breast cancer subtypes. Nat Commun. 7, 10259 (2016).

14. S. Tyanova, J. Cox, Perseus: A Bioinformatics Platform for Integrative Analysis of Proteomics Data in Cancer Research. Methods in Molecular Biology. 1711, 133–148 (2018).

15. R. Han, E. P. Rader, J. R. Levy, D. Bansal, K. P. Campbell, Dystrophin deficiency exacerbates skeletal muscle pathology in dysferlin-null mice. Skelet. Muscle. 1, 35 (2011).

16. E. P. Rader, R. Turk, T. Willer, D. Beltrán, K. Inamori, T. A. Peterson, J. Engle, S. Prouty, K. Matsumura, F. Saito, M. E. Anderson, K. P. Campbell, Role of dystroglycan in limiting contraction-induced injury to the sarcomeric cytoskeleton of mature skeletal muscle. Proc. Natl. Acad. Sci. 113, 10992–10997 (2016).

17. H. Okuma, J. M. Hord, I. Chandel, D. Venzke, M. E. Anderson, A. S. Walimbe, S. Joseph, Z Gastel, Y. Hara, F. Saito, K. Matsumura, K. P. Campbell, N-terminal domain on dystroglycan enables LARGE1 to extend matriglycan on α-dystroglycan and prevents muscular dystrophy. Elife 12, e82811 (2023).

## References

1. J. H. Gommers-Ampt, F. V. Leeuwen, A. L. de Beer, J. F. Vliegenthart, M. Dizdaroglu, J. A. Kowalak, P. F. Crain, P. Borst, beta-D-glucosyl-hydroxymethyluracil: a novel modified base present in the DNA of the parasitic protozoan T. brucei. Cell 75, 1129–1136 (1993).

2. C. R. H. Raetz, C. Whitfield, LIPOPOLYSACCHARIDE ENDOTOXINS. Annu. Rev. Biochem. 71, 635–700 (2002).

3. J. L. Daniotti, A. A. Vilcaes, V. T. Demichelis, F. M. Ruggiero, M. Rodriguez-Walker, Glycosylation of glycolipids in cancer: basis for development of novel therapeutic approaches. Frontiers in oncology 3, 306 (2013).

4. H. Nothaft, C. M. Szymanski, Protein glycosylation in bacteria: sweeter than ever. Nat Rev Microbiol 8, 765–778 (2010).

5. C. Reily, T. J. Stewart, M. B. Renfrow, J. Novak, Glycosylation in health and disease. Nat Rev Nephrol 15, 346–366 (2019).

6. C. Xu, D. T. W. Ng, Glycosylation-directed quality control of protein folding. Nature reviews. Molecular cell biology 16, 742–752 (2015).

7. A. Peixoto, M. Relvas-Santos, R. Azevedo, L. L. Santos, J. A. Ferreira, Protein Glycosylation and Tumor Microenvironment Alterations Driving Cancer Hallmarks. Frontiers Oncol 9, 380 (2019).

8. M. R. Kudelka, S. R. Stowell, R. D. Cummings, A. S. Neish, Intestinal epithelial glycosylation in homeostasis and gut microbiota interactions in IBD. Nat Rev Gastroentero 17, 597–617 (2020).

9. C. Boscher, J. W. Dennis, I. R. Nabi, Glycosylation, galectins and cellular signaling. Curr Opin Cell Biol 23, 383–392 (2011).

10. O. Ibraghimov-Beskrovnaya, J. M. Ervasti, C. J. Leveille, C. A. Slaughter, S. W. Sernett, K. P. Campbell, Primary structure of dystrophin-associated glycoproteins linking dystrophin to the extracellular matrix. Nature 355, 696–702 (1992).

11. M. Kanagawa, F. Saito, S. Kunz, T. Yoshida-Moriguchi, R. Barresi, Y. M. Kobayashi, J. Muschler, J. P. Dumanski, D. E. Michele, M. B. A. Oldstone, K. P. Campbell, Molecular recognition by LARGE is essential for expression of functional dystroglycan. Cell 117, 953–964 (2004).

12. J. F. Talts, Z. Andac, W. Göhring, A. Brancaccio, R. Timpl, Binding of the G domains of laminin α1 and α2 chains and perlecan to heparin, sulfatides, αLdystroglycan and several extracellular matrix proteins. Embo J 18, 863–870 (1999).

13. D. C. Briggs, T. Yoshida-Moriguchi, T. Zheng, D. Venzke, M. E. Anderson, A. Strazzulli, M. Moracci, L. Yu, E. Hohenester, K. P. Campbell, Structural basis of laminin binding to the LARGE glycans on dystroglycan. Nat Chem Biol 12, 810–814 (2016).

14. J. Ervasti, K. Campbell, A role for the dystrophin-glycoprotein complex as a transmembrane linker between laminin and actin. J Cell Biology 122, 809–823 (1993).

15. S. H. Gee, F. Montanaro, M. H. Lindenbaum, S. Carbonetto, Dystroglycan-α, a dystrophin-associated glycoprotein, is a functional agrin receptor. Cell 77, 675–686 (1994).

16. J. M. Ervasti, K. P. Campbell, Membrane organization of the dystrophin-glycoprotein complex. Cell 66, 1121–1131 (1991).

17. K. Inamori, T. Yoshida-Moriguchi, Y. Hara, M. E. Anderson, L. Yu, K. P. Campbell, Dystroglycan function requires xylosyl- and glucuronyltransferase activities of LARGE. Science 335, 93–96 (2012).

18. M. M. Goddeeris, B. Wu, D. Venzke, T. Yoshida-Moriguchi, F. Saito, K. Matsumura, S. A. Moore, K. P. Campbell, LARGE glycans on dystroglycan function as a tunable matrix scaffold to prevent dystrophy. Nature 503, 136–140 (2013).

19. T. Willer, K. Inamori, D. Venzke, C. Harvey, G. Morgensen, Y. Hara, D. B. V. de Bernabé, L. Yu, K. M. Wright, K. P. Campbell, The glucuronyltransferase B4GAT1 is required for initiation of LARGE-mediated α-dystroglycan functional glycosylation. Elife 3, e03941 (2014).

20. J. L. Praissman, D. H. Live, S. Wang, A. Ramiah, Z. S. Chinoy, G.-J. Boons, K. W. Moremen, L. Wells, B4GAT1 is the priming enzyme for the LARGE-dependent functional glycosylation of α-dystroglycan. Elife 3, e03943 (2014).

21. H. Manya, Y. Yamaguchi, M. Kanagawa, K. Kobayashi, M. Tajiri, K. Akasaka-Manya, H. Kawakami, M. Mizuno, Y. Wada, T. Toda, T. Endo, The Muscular Dystrophy Gene TMEM5 Encodes a Ribitol β1,4-Xylosyltransferase Required for the Functional Glycosylation of Dystroglycan*. J Biol Chem 291, 24618–24627 (2016).

22. J. L. Praissman, T. Willer, M. O. Sheikh, A. Toi, D. Chitayat, Y.-Y. Lin, H. Lee, S. H. Stalnaker, S. Wang, P. K. Prabhakar, S. F. Nelson, D. L. Stemple, S. A. Moore, K. W. Moremen, K. P. Campbell, L. Wells, The functional O-mannose glycan on α-dystroglycan contains a phospho-ribitol primed for matriglycan addition. Elife 5, e14473 (2016).

23. M. Kanagawa, K. Kobayashi, M. Tajiri, H. Manya, A. Kuga, Y. Yamaguchi, K. Akasaka-Manya, J. Furukawa, M. Mizuno, H. Kawakami, Y. Shinohara, Y. Wada, T. Endo, T. Toda, Identification of a Post-translational Modification with Ribitol-Phosphate and Its Defect in Muscular Dystrophy. Cell reports 14, 2209–2223 (2016).

24. H. Yagi, C.-W. Kuo, T. Obayashi, S. Ninagawa, K.-H. Khoo, K. Kato, Direct Mapping of Additional Modifications on Phosphorylated O-glycans of α-Dystroglycan by Mass Spectrometry Analysis in Conjunction with Knocking Out of Causative Genes for Dystroglycanopathy. Molecular & Cellular Proteomics 15, 3424–3434 (2016).

25. K. Fujimura, H. Sawaki, T. Sakai, T. Hiruma, N. Nakanishi, T. Sato, T. Ohkura, H. Narimatsu, LARGE2 facilitates the maturation of alpha-dystroglycan more effectively than LARGE. Biochemical and biophysical research communications 329, 1162–1171 (2005).

26. M. Brockington, S. Torelli, P. Prandini, C. Boito, N. F. Dolatshad, C. Longman, S. C. Brown, F. Muntoni, Localization and functional analysis of the LARGE family of glycosyltransferases: significance for muscular dystrophy. Hum Mol Genet 14, 657–665 (2005).

27. K. Inamori, T. Willer, Y. Hara, D. Venzke, M. E. Anderson, N. F. Clarke, P. Guicheney, C. G. Bönnemann, S. A. Moore, K. P. Campbell, Endogenous Glucuronyltransferase Activity of LARGE or LARGE2 Required for Functional Modification of α-Dystroglycan in Cells and Tissues*. J Biol Chem 289, 28138–28148 (2014).

28. T. Yoshida-Moriguchi, T. Willer, M. E. Anderson, D. Venzke, T. Whyte, F. Muntoni, H. Lee, S. F. Nelson, L. Yu, K. P. Campbell, SGK196 is a glycosylation-specific O-mannose kinase required for dystroglycan function. Science 341, 896–899 (2013).

29. T. Hiruma, A. Togayachi, K. Okamura, T. Sato, N. Kikuchi, Y.-D. Kwon, A. Nakamura, K. Fujimura, M. Gotoh, K. Tachibana, Y. Ishizuka, T. Noce, H. Nakanishi, H. Narimatsu, A novel human beta1,3-N-acetylgalactosaminyltransferase that synthesizes a unique carbohydrate structure, GalNAcbeta1-3GlcNAc. Journal of Biological Chemistry 279, 14087–14095 (2004).

30. E. Stevens, K. J. Carss, S. Cirak, A. R. Foley, S. Torelli, T. Willer, D. E. Tambunan, S. Yau, L. Brodd, C. A. Sewry, L. Feng, G. Haliloglu, D. Orhan, W. B. Dobyns, G. M. Enns, M. Manning, A. Krause, M. A. Salih, C. A. Walsh, M. Hurles, K. P. Campbell, M. C. Manzini, U. Consortium, D. Stemple, Y.-Y. Lin, F. Muntoni, Mutations in B3GALNT2 cause congenital muscular dystrophy and hypoglycosylation of α-dystroglycan. American journal of human genetics 92, 354–365 (2013).

31. H. Manya, A. Chiba, A. Yoshida, X. Wang, Y. Chiba, Y. Jigami, R. U. Margolis, T. Endo, Demonstration of mammalian protein O-mannosyltransferase activity: Coexpression of POMT1 and POMT2 required for enzymatic activity. Proc National Acad Sci 101, 500–505 (2004).

32. T. Yoshida-Moriguchi, L. Yu, S. H. Stalnaker, S. Davis, S. Kunz, M. Madson, M. B. A. Oldstone, H. Schachter, L. Wells, K. P. Campbell, O-mannosyl phosphorylation of alpha-dystroglycan is required for laminin binding. Science 327, 88–92 (2010).

33. M. Kanagawa, Dystroglycanopathy: From Elucidation of Molecular and Pathological Mechanisms to Development of Treatment Methods. Int J Mol Sci 22, 13162 (2021).

34. T. Willer, H. Lee, M. Lommel, T. Yoshida-Moriguchi, D. B. V. de Bernabe, D. Venzke, S. Cirak, H. Schachter, J. Vajsar, T. Voit, F. Muntoni, A. S. Loder, W. B. Dobyns, T. L. Winder, S. Strahl, K. D. Mathews, S. F. Nelson, S. A. Moore, K. P. Campbell, ISPD loss-of-function mutations disrupt dystroglycan O-mannosylation and cause Walker-Warburg syndrome. Nat Genet 44, 575–580 (2012).

35. Y. Hara, B. Balci-Hayta, T. Yoshida-Moriguchi, M. Kanagawa, D. B. V. de Bernabé, H. Gündeşli, T. Willer, J. S. Satz, R. W. Crawford, S. J. Burden, S. Kunz, M. B. A. Oldstone, A. Accardi, B. Talim, F. Muntoni, H. Topaloglu, P. Dinçer, K. P. Campbell, A dystroglycan mutation associated with limb-girdle muscular dystrophy. New England Journal of Medicine 364, 939–946 (2011).

36. R. Harrison, P. G. Hitchen, M. Panico, H. R. Morris, D. Mekhaiel, R. J. Pleass, A. Dell, J. E. Hewitt, S. M. Haslam, Glycoproteomic characterization of recombinant mouse α-dystroglycan. Glycobiology 22, 662–675 (2012).

37. Y. Hara, M. Kanagawa, S. Kunz, T. Yoshida-Moriguchi, J. S. Satz, Y. M. Kobayashi, Z. Zhu, S. J. Burden, M. B. A. Oldstone, K. P. Campbell, Like-acetylglucosaminyltransferase (LARGE)-dependent modification of dystroglycan at Thr-317/319 is required for laminin binding and arenavirus infection. Proc National Acad Sci 108, 17426–17431 (2011).

38. N. Nakagawa, H. Takematsu, S. Oka, HNK-1 sulfotransferase-dependent sulfation regulating laminin-binding glycans occurs in the post-phosphoryl moiety on α-dystroglycan. Glycobiology 23, 1066–1074 (2013).

39. M. O. Sheikh, D. Venzke, M. E. Anderson, T. Yoshida-Moriguchi, J. N. Glushka, A. V. Nairn, M. Galizzi, K. W. Moremen, K. P. Campbell, L. Wells, HNK-1 sulfotransferase modulates α-dystroglycan glycosylation by 3-O-sulfation of glucuronic acid on matriglycan. Glycobiology 30, 817–829 (2020).

40. M. Riemersma, J. Sandrock, T. J. Boltje, C. Büll, T. Heise, A. Ashikov, G. J. Adema, H. van Bokhoven, D. J. Lefeber, Disease mutations in CMP-sialic acid transporter SLC35A1 result in abnormal α-dystroglycan O-mannosylation, independent from sialic acid. Human molecular genetics 24, 2241–2246 (2015).

41. C. Diesen, A. Saarinen, H. Pihko, C. Rosenlew, B. Cormand, W. B. Dobyns, J. Dieguez, L. Valanne, T. Joensuu, A.-E. Lehesjoki, POMGnT1 mutation and phenotypic spectrum in muscle-eye-brain disease. Journal of Medical Genetics 41, e115–e115 (2004).

42. R. Biancheri, E. Bertini, A. Falace, M. Pedemonte, A. Rossi, A. D’Amico, S. Scapolan, L. Bergamino, S. Petrini, D. Cassandrini, P. Broda, M. Manfredi, F. Zara, F. M. Santorelli, C. Minetti, C. Bruno, POMGnT1 mutations in congenital muscular dystrophy - Genotype-phenotype correlation and expanded clinical spectrum. Archives of Neurology 63, 1491–1495 (2006).

43. B. Ury, S. Potelle, F. Caligiore, M. R. Whorton, G. T. Bommer, The promiscuous binding pocket of SLC35A1 ensures redundant transport of CDP-ribitol to the Golgi. J Biol Chem 296, 100789 (2021).

44. N. Kuwabara, H. Manya, T. Yamada, H. Tateno, M. Kanagawa, K. Kobayashi, K. Akasaka-Manya, Y. Hirose, M. Mizuno, M. Ikeguchi, T. Toda, J. Hirabayashi, T. Senda, T. Endo, R. Kato, Carbohydrate-binding domain of the POMGnT1 stem region modulates O-mannosylation sites of α-dystroglycan. Proceedings of the National Academy of Sciences 113, 9280–9285 (2016).

45. T. Yonekawa, A. J. Rauckhorst, S. El-Hattab, M. A. Cuellar, D. Venzke, M. E. Anderson, H. Okuma, A. D. Pewa, E. B. Taylor, K. P. Campbell, Large1 gene transfer in older myd mice with severe muscular dystrophy restores muscle function and greatly improves survival. Sci Adv 8, eabn0379 (2022).

46. K. M. Gharpure, O. D. Lara, Y. Wen, S. Pradeep, C. LaFargue, C. Ivan, R. Rupaimoole, W. Hu, L. S. Mangala, S. Y. Wu, A. S. Nagaraja, K. Baggerly, A. K. Sood, ADH1B promotes mesothelial clearance and ovarian cancer infiltration. Oncotarget 9, 25115–25126 (2018).

47. N. Pacheco-Fernandez, M. Pakdel, B. Blank, I. Sanchez-Gonzalez, K. Weber, M. L. Tran, T. K.-H. Hecht, R. Gautsch, G. Beck, F. Perez, A. Hausser, S. Linder, J. von Blume, Nucleobindin-1 regulates ECM degradation by promoting intra-Golgi trafficking of MMPs. J. Cell Biol. 219, e201907058 (2020).

48. H. Okuma, J. M. Hord, I. Chandel, D. Venzke, M. E. Anderson, A. S. Walimbe, S. Joseph, Z. Gastel, Y. Hara, F. Saito, K. Matsumura, K. P. Campbell, N-terminal domain on dystroglycan enables LARGE1 to extend matriglycan on α-dystroglycan and prevents muscular dystrophy. Elife 12, e82811 (2023).

49. A. S. Walimbe, H. Okuma, S. Joseph, T. Yang, T. Yonekawa, J. M. Hord, D. Venzke, M. E. Anderson, S. Torelli, A. Manzur, M. Devereaux, M. Cuellar, S. Prouty, S. O. Landa, L. Yu, J. Xiao, J. E. Dixon, F. Muntoni, K. P. Campbell, POMK regulates dystroglycan function via LARGE-mediated elongation of matriglycan. Elife 9, e61388 (2020).

50. D. Vanhoutte, T. G. Schips, J. Q. Kwong, J. Davis, A. Tjondrokoesoemo, M. J. Brody, M. A. Sargent, O. Kanisicak, H. Yi, Q. Q. Gao, J. E. Rabinowitz, T. Volk, E. M. McNally, J. D. Molkentin, Thrombospondin expression in myofibers stabilizes muscle membranes. eLife 5, e17589 (2016).

51. G. Juban, A. Bertrand-Chapel, S. Meyer, A. Kneppers, P. Huchedé, C. Gallerne, R. Benayoun, E. Cohen, A. Lopez-Gonzales, S. B. Larbi, M. Creveaux, L. Vaille, A. Bouvier, M. Théodore, L. Broutier, A. Dutour, M. Cordier-Bussat, J.-Y. Blay, N. Streichenberger, C. Picard, N. Corradini, V. Allamand, P. Castets, R. Mounier, M. Castets, Alterations of the TGFb-sequestration complex member ADAMTSL1 levels are associated with muscular defects and rhabdomyosarcoma aggressiveness. bioRxiv, 2023.03.07.531559 (2023).

