## Supplementary Figures and Legends for "Identification of a short, single site matriglycan that maintains neuromuscular function in the mouse"

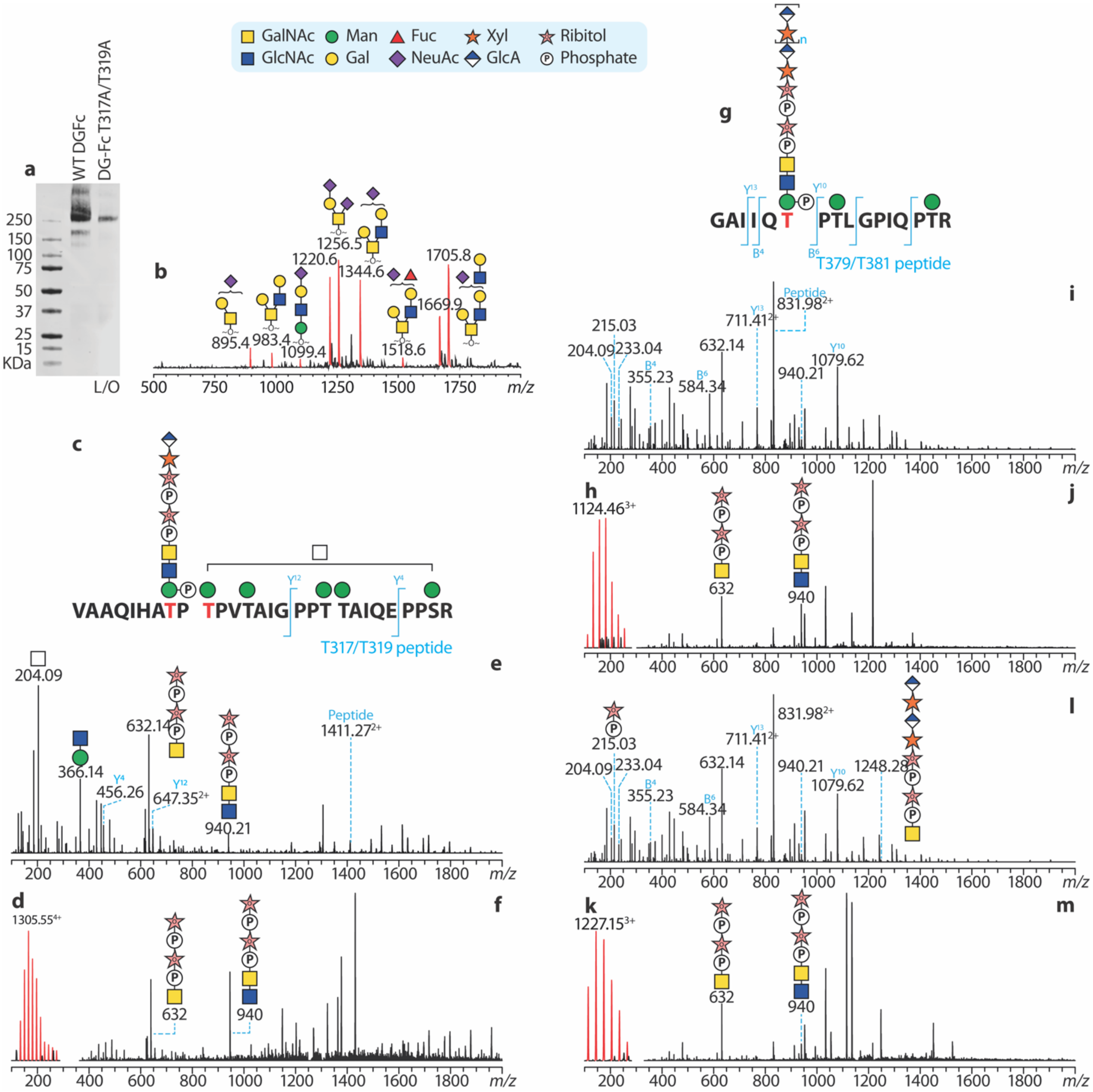


Fig.s1 Biochemical evidence indicating more matriglycosylation sites. (a) Laminin overlay (L/O) of DG-Fc and DG-Fc carrying a 317A/T319A mutation. (b) MALDI-TOF based O-glycomics of full length DAG1749. O-glycans were reductively eliminated and permethylated before MS analysis. MS peaks corresponding to sodiated glycans were colored red and annotated with m/z and glycan structures. (c-f) MS of a T317/T319 glycopeptide carrying the matriglycan precursor (c). A representative MS signal is shown in (d). HCD and CID MSMS are shown in (e) and (f) respectively. (g-m) MS of a T379/T381 glycopeptide (g) carrying the matriglycan precursor (n=0) and matriglycan (n=1). Representative MS signals are shown in (h) and (k). HCD and CID MSMS are shown in (i and j) and (l and m) respectively.

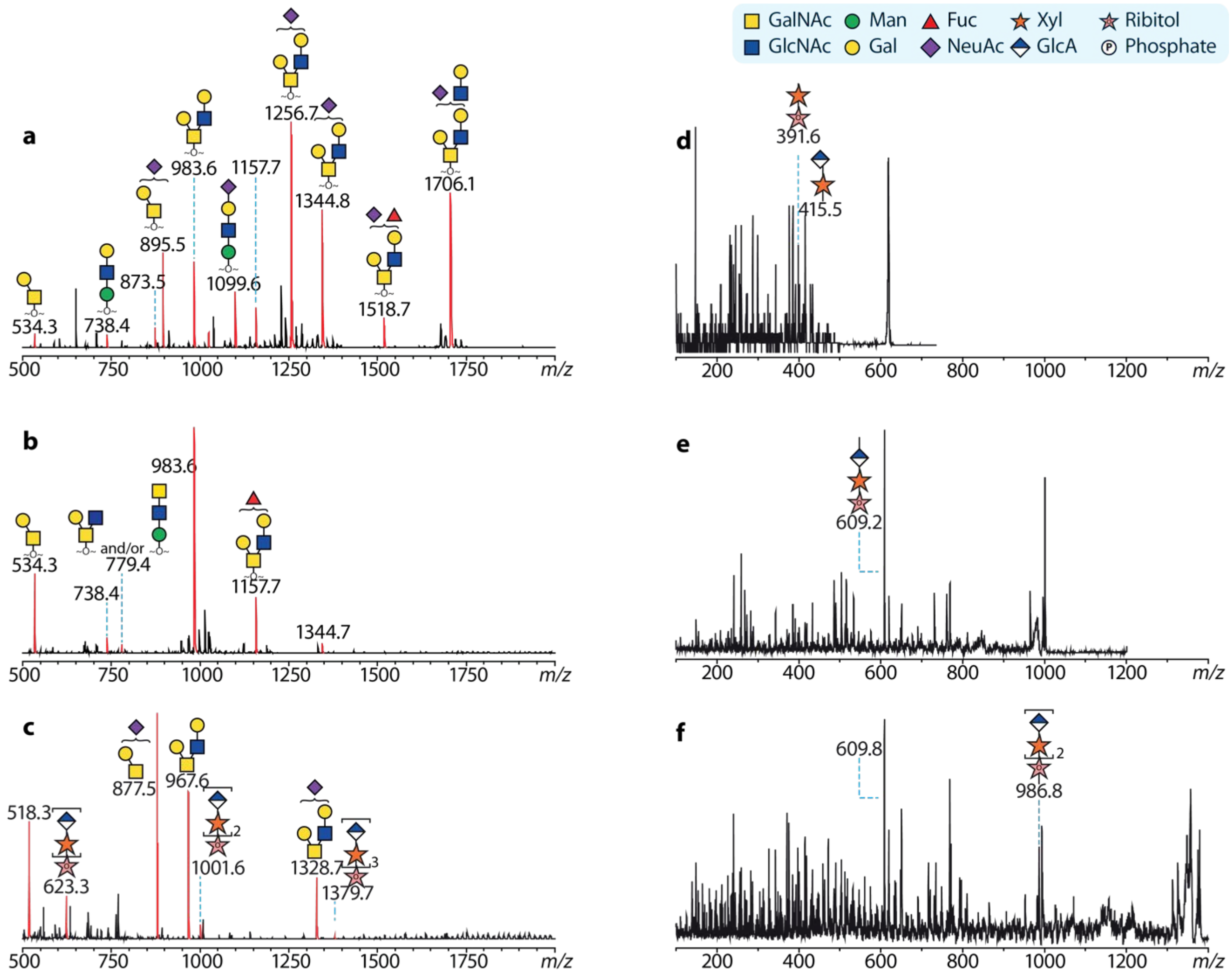


Fig.s2 MALDI-TOF based O-glycomics of full length DG390. MS spectra of O-glycans reductively eliminated from DG390 and de-glycosylated DG390 are shown in (a) and (b), mild HF hydrolysis released from DG390 in (c). MS peaks corresponding to sodiated glycans were colored red and annotated with m/z and glycan structures. MS peaks at *m/z* 632.3, 1001.6 and 1379.7 were further characterized by MALDI-TOF/TOF and spectra are shown in (d-f). The MSMS signals are low due to a lack of amino-sugars in these glycan fragments.

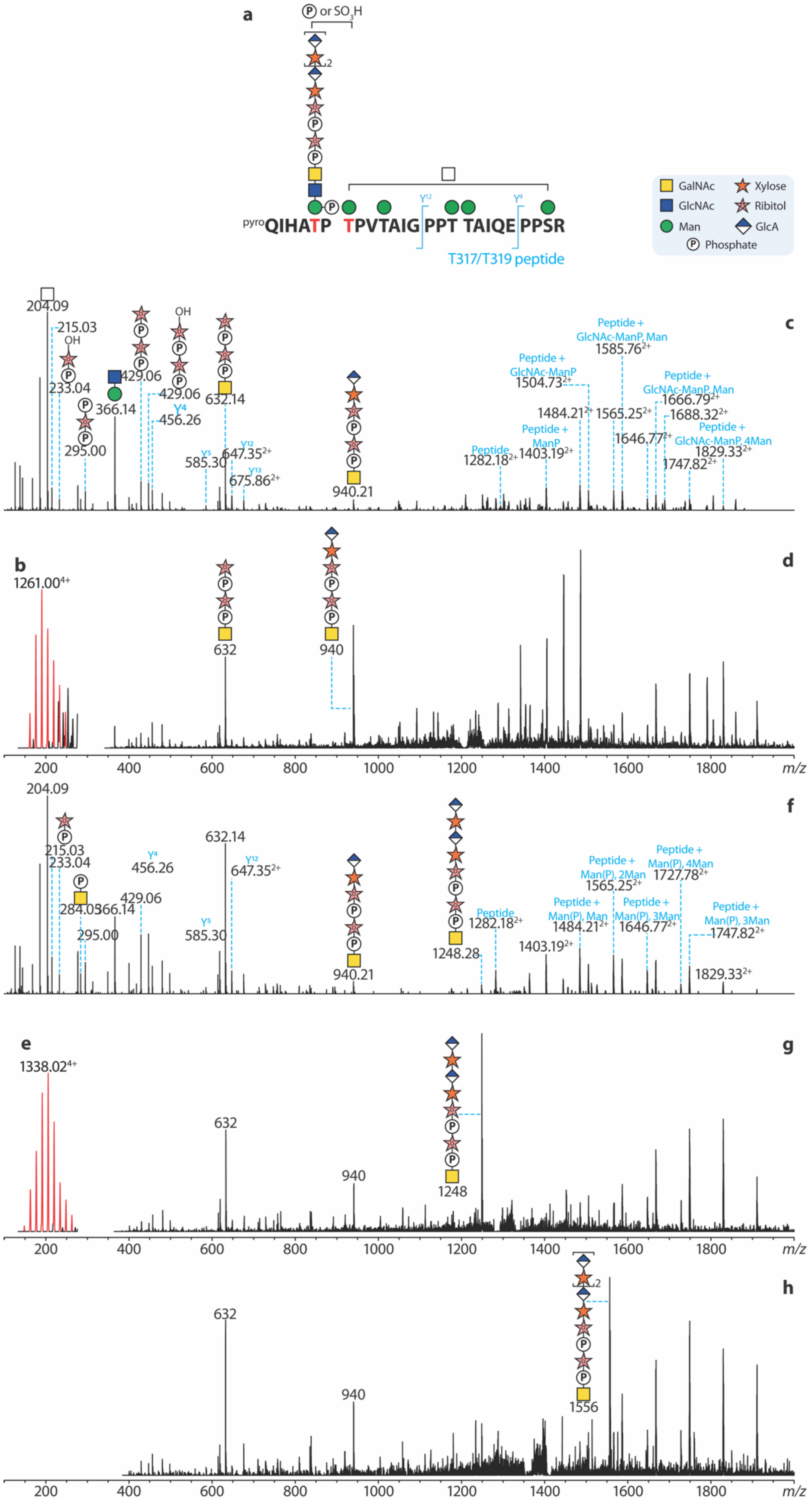
a
P or SO3H
2
pyroQIHATP TPVTAIG PPT TAIQE PPSR
Y12
Y4
T317/T319 peptide
GalNAc
GlcNAc
Man
Xylose
Ribitol
GlcA
Phosphate
b
1261.00 4+
c
204.09
215.03
OH
233.04
295.00
366.14
429.06
429.06
Y4
456.26
632.14
Y5
585.30
Y12
647.35 2+
Y13
675.86 2+
940.21
Peptide
1282.18 2+
Peptide + ManP
1403.19 2+
Peptide + GlcNAc-ManP
1504.73 2+
1484.21 2+
Peptide + GlcNAc-ManP, Man
1585.76 2+
1565.25 2+
Peptide + GlcNAc-ManP, Man
1666.79 2+
1688.32 2+
1646.77 2+
Peptide + GlcNAc-ManP, 4Man
1829.33 2+
1747.82 2+
d
632
940
m/z
e
1338.02 4+
f
204.09
215.03
233.04
284.03
366.14
429.06
295.00
Y4
456.26
632.14
Y12
647.35 2+
Y5
585.30
940.21
1248.28
Peptide
1282.18 2+
1403.19 2+
Peptide + Man(P), Man
1484.21 2+
Peptide + Man(P), 2Man
1565.25 2+
Peptide + Man(P), 3Man
1646.77 2+
Peptide + Man(P), 4Man
1727.78 2+
Peptide + Man(P), 3Man
1747.82 2+
1829.33 2+
g
632
940
1248
h
632
940
1556
200
400
600
800
1000
1200
1400
1600
1800
m/z

Fig.s3 High-resolution MS mapping of T317/T319 matriglycan (a). Representative MS signals corresponding to the glycopeptides carrying the precursor of matriglycan (b) and matriglycan with one repeating unit (e). The two MS signals were further characterized by HCD (c and f) and CID (d and g). (h) shows the CID spectrum of MS signals in **fig.2b**, Note the CID spectra has lower resolution. High resolution CID can be found in raw data. These data supplements MS data in **fig.2** showing the matriglycan being added to the T317/T319 glycopeptides.

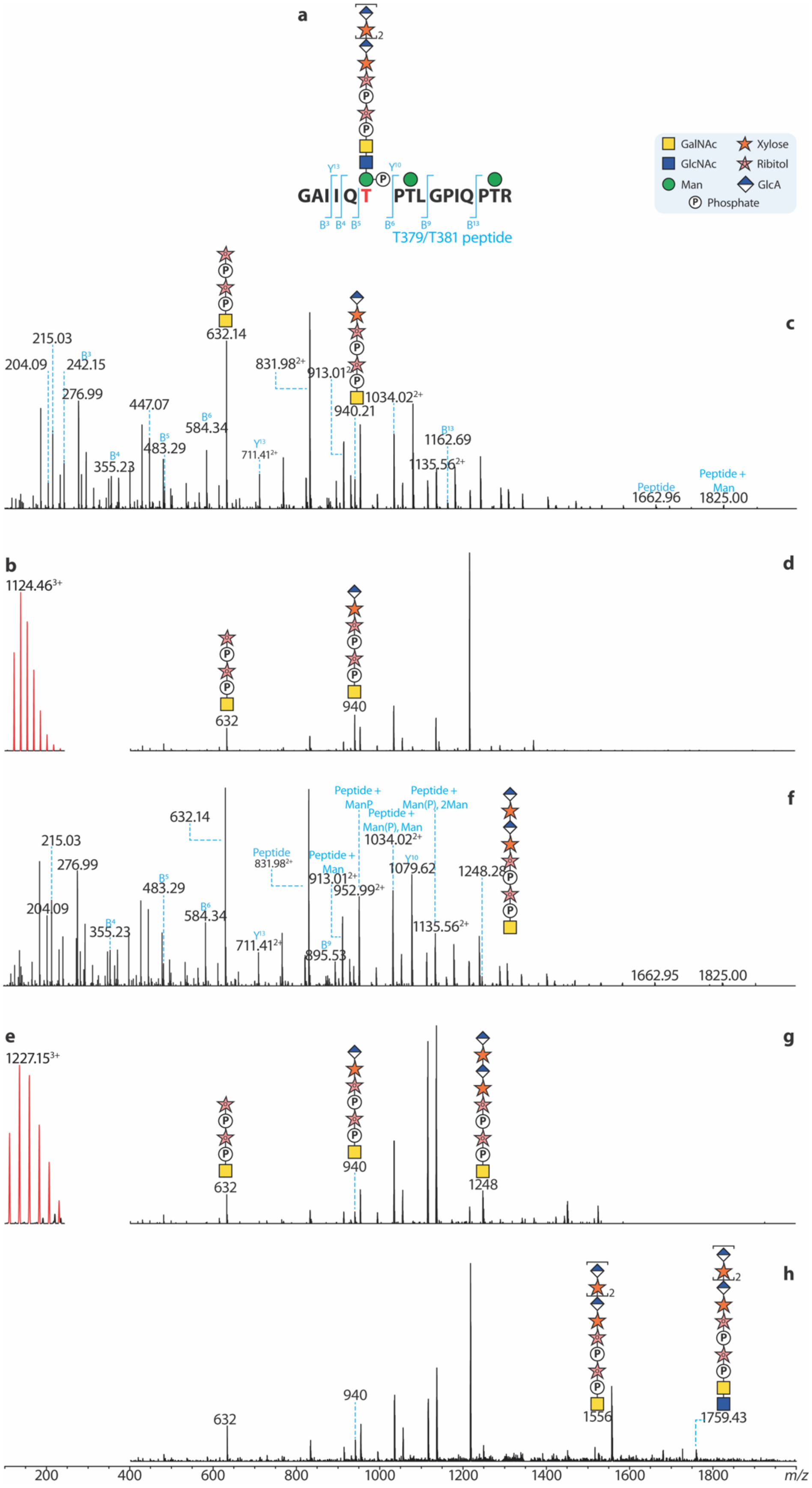

a
b
c
d
e
f
g
h
GalNAc
GlcNAc
Man
Xylose
Ribitol
GlcA
Phosphate
GAI I Q T PTL GPIQ PTR
T379/T381 peptide
1124.46³⁺
1227.15³⁺
632.14
940.21
1034.02²⁺
1162.69
1135.56²⁺
1248.28
1079.62
1662.96
1825.00
Peptide + Man
1759.43
1556
m/z

Fig.s4 High-resolution MS mapping of T379/T381 matriglycan (a). Representative MS signals corresponding to the glycopeptides carrying the precursor of matriglycan (b) and matriglycan with one repeating unit (e). The two MS signals were further characterized by HCD (c and f) and CID (d and g). (h) shows the CID spectrum of MS signals in **fig.2e**, Note the CID spectra has lower resolution. High resolution CID can be found in raw data. These data supplements MS data in **fig.2** showing the matriglycan being added to the T379/T381 glycopeptides.

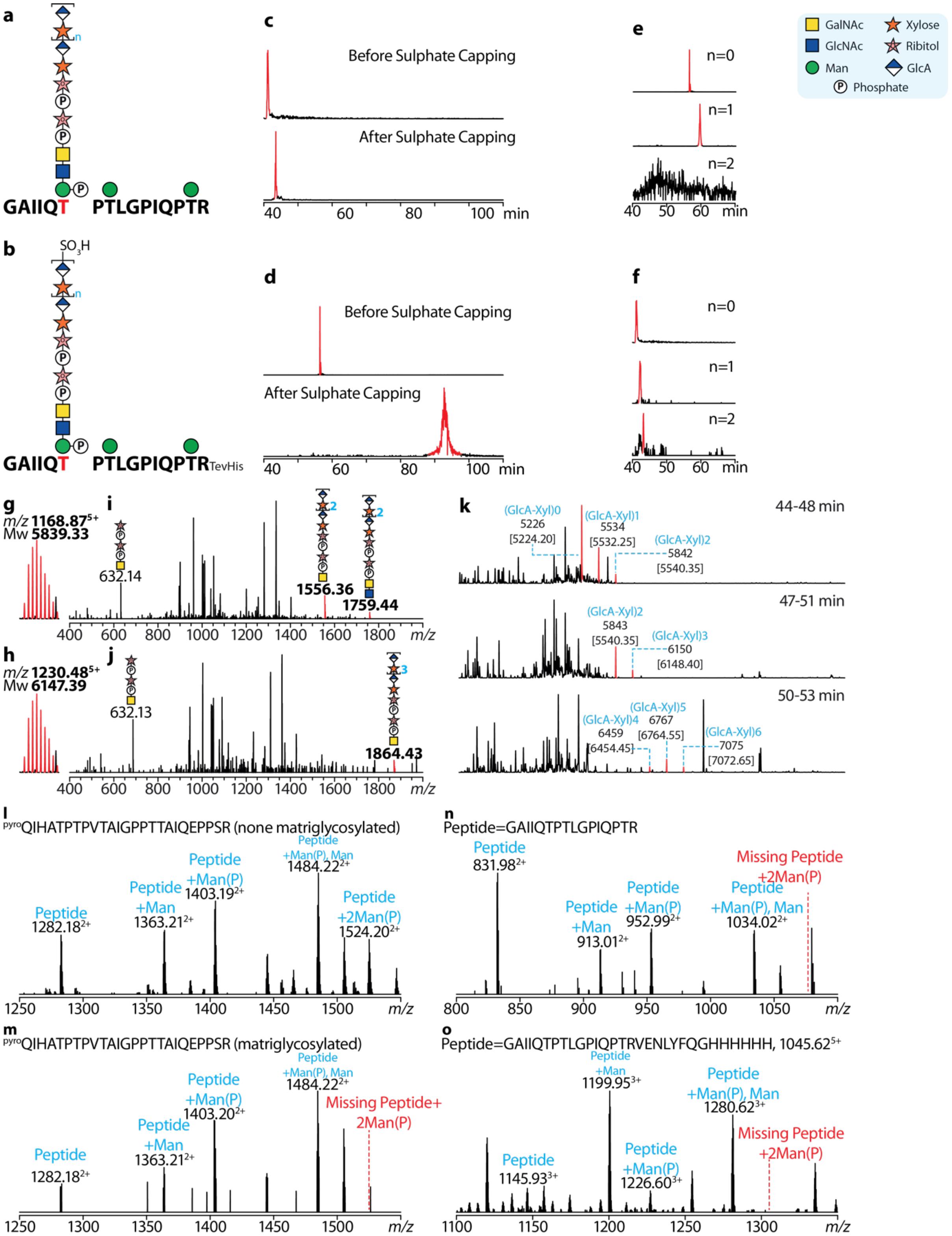

a
GAIIQT PTLGPIQPTR
n
b
SO3H
GAIIQT PTLGPIQPTRTevHis
c
Before Sulphate Capping
After Sulphate Capping
40 60 80 100 min
d
Before Sulphate Capping
After Sulphate Capping
40 60 80 100 min
e
n=0
n=1
n=2
40 50 60 min
f
n=0
n=1
n=2
40 50 60 min
GalNAc
GlcNAc
Man
Xylose
Ribitol
GlcA
Phosphate
g
m/z 1168.87 5+
Mw 5839.33
i
632.14
1556.36
1759.44
400 600 800 1000 1200 1400 1600 1800 m/z
h
m/z 1230.48 5+
Mw 6147.39
j
632.13
1864.43
400 600 800 1000 1200 1400 1600 1800 m/z
k
44-48 min
(GlcA-Xyl)0 5226 [5224.20]
(GlcA-Xyl)1 5534 [5532.25]
(GlcA-Xyl)2 5842 [5540.35]
47-51 min
(GlcA-Xyl)2 5843 [5540.35]
(GlcA-Xyl)3 6150 [6148.40]
50-53 min
(GlcA-Xyl)4 6459 [6454.45]
(GlcA-Xyl)5 6767 [6764.55]
(GlcA-Xyl)6 7075 [7072.65]
l
pyroQIHATPTPVTAIGPPTTAIQEPPSR (none matriglycosylated)
Peptide 1282.18 2+
Peptide +Man 1363.21 2+
Peptide +Man(P) 1403.19 2+
Peptide +Man(P), Man 1484.22 2+
Peptide +2Man(P) 1524.20 2+
1250 1300 1350 1400 1450 1500 m/z
m
pyroQIHATPTPVTAIGPPTTAIQEPPSR (matriglycosylated)
Peptide 1282.18 2+
Peptide +Man 1363.21 2+
Peptide +Man(P) 1403.20 2+
Peptide +Man(P), Man 1484.22 2+
Missing Peptide+ 2Man(P)
1250 1300 1350 1400 1450 1500 m/z
n
Peptide=GAIIQTPTLGPIQPTR
Peptide 831.98 2+
Peptide +Man 913.01 2+
Peptide +Man(P) 952.99 2+
Peptide +Man(P), Man 1034.02 2+
Missing Peptide +2Man(P)
800 850 900 950 1000 1050 m/z
o
Peptide=GAIIQTPTLGPIQPTRVENLYFQGHHHHHH, 1045.62 5+
Peptide 1145.93 3+
Peptide +Man 1199.95 3+
Peptide +Man(P) 1226.60 3+
Peptide +Man(P), Man 1280.62 3+
Missing Peptide +2Man(P)
1100 1150 1200 1250 1300 m/z

Fig.s5 Further MS characterization of polymeric matriglycan on the T379/T381 motif. Matriglycans can be capped by sulphate groups. Diagrams illustrating T379/T381 glycopeptides without (a) and with (b) the sulphate are shown. Matriglycan sulphation significantly delays reverse-phase LC retention times (d) but can be partially compensated by the his-tag (c, both glycopeptides are his-tagged). Extracted ion chromatograms (XICs) of his-tagged T379/T381 glycopeptides with or without sulphate capping show similar retention times (c) but non-his-tagged glycopeptides show very different retention times (d). The his-tag also facilitates ESI-MS characterization of glycopeptides with polymeric matriglycans. XICs of non-his-tagged (e) and his-tagged (f) glycopeptides with matriglycans are shown. It is apparent the longer matriglycan signals were missing when the glycopeptide is not his-tagged.
(g-h) High-resolution CID of glycopeptides with polymeric matriglycans. Nearly intact matriglycan fragments can be readily detected. (k) Deconvoluted MS spectra of the T379/T381 glycopeptides carrying polymeric matriglycans. Theoretical single charged *m/z* are annotated below the observed *m/z* with square brackets.
MSMS fragments of T317/T319 glycopeptides carrying two mannose-phosphate groups can be detected by MS (l), but these fragments were missing in the matriglycosylated T317/T319 glycopeptides (m) and T379/T381 glycopeptides (n and o).

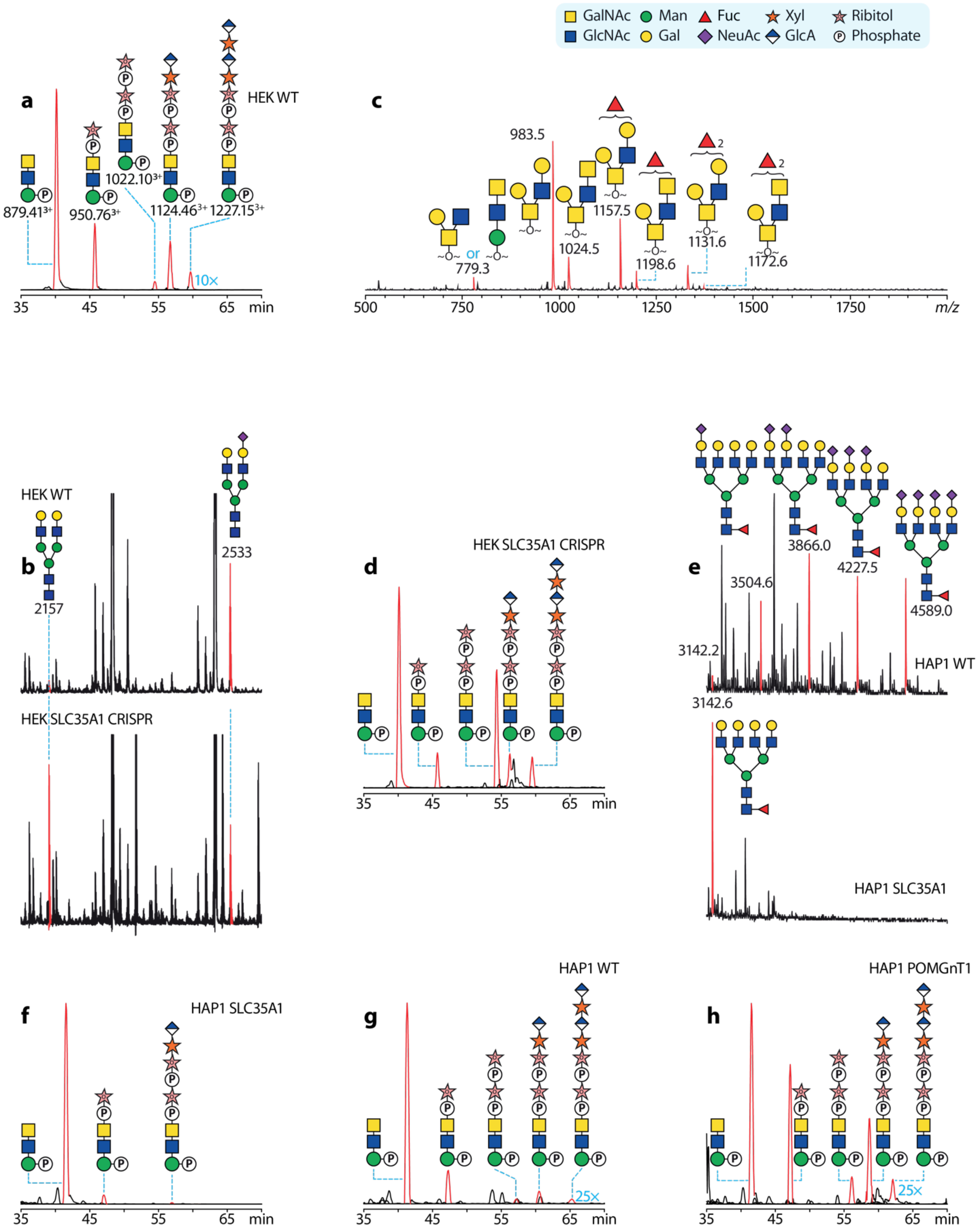

GalNAc
GlcNAc
Man
Gal
Fuc
NeuAc
Xyl
GlcA
Ribitol
Phosphate
a
HEK WT
879.41³⁺
950.76³⁺
1022.10³⁺
1124.46³⁺
1227.15³⁺
10×
35 45 55 65 min
c
983.5
779.3
or
1024.5
1157.5
1198.6
1131.6
1172.6
2
2
500 750 1000 1250 1500 1750 m/z
b
HEK WT
2157
2533
HEK SLC35A1 CRISPR
d
HEK SLC35A1 CRISPR
35 45 55 65 min
e
3142.2
3504.6
3866.0
4227.5
4589.0
HAP1 WT
3142.6
HAP1 SLC35A1
f
HAP1 SLC35A1
35 45 55 65 min
g
HAP1 WT
25×
35 45 55 65 min
h
HAP1 POMGnT1
25×
35 45 55 65 min

Fig.s6 (a) Extracted ion chromatograms (XICs) of MS signals corresponding to glycan structures along matriglycan biosynthetic pathway. The m/z used to generate the XICs are annotated below matriglycan-related structures. Note they are omitted in the following figures. These glycans are carried by the T379/T381 glycopeptides shows in **fig.2d**. (b) Selected region of a MALDI-TOF based N-glycomics data indicating sialylated glycans are not eliminated in our HEK SLC35A1 CRISPR cells. These N-glycans were released by PNGaseF and deutero-permethylated. (c) MALDI-TOF based O-glycomics could not detect sialylated O-glycans on DG390. The O-glycans were reductively eliminated from DG390 and permethylated. Red signals show sodiated O-glycans after derivatization. (d) XIC showing matriglycosylation can be detected on DG390 expressed by our HEK SLC25A1 CRISPR cells. (e) Selected region of a MALDI-TOF based N-glycomics data indicating no sialylation in our HAP SLC35A1 cells. These N-glycans were released by PNGaseF and permethylated. (f) XIC showing ribitol-phosphate containing glycopeptides significantly down-regulated in the SLC35A1 KO (f) but not in WT (g) and POMGnT1 KO HAP1 (h) cells. Note the matriglycosylated XIC signals were amplified to different folds throughout this figure.

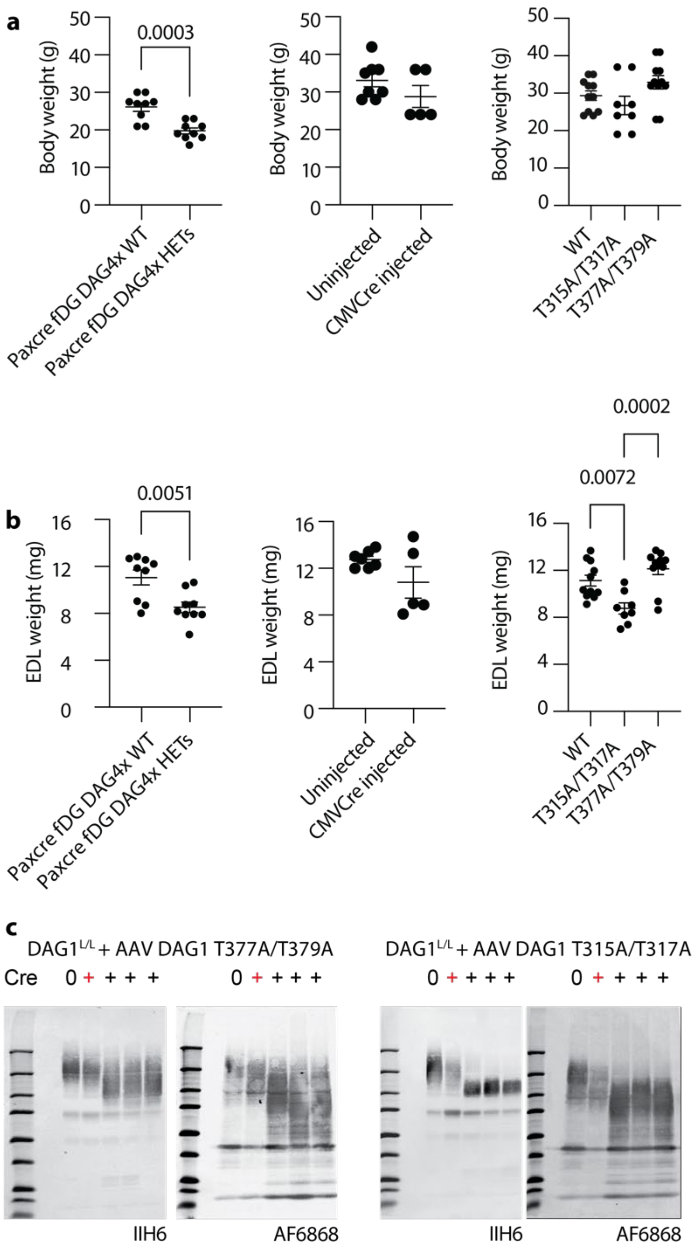


Fig.s7 Body (a) and EDL (b) wight of our transgenic mice. (c) Immunoblot of skeletal from different $DAG1^{L/L}$ mice. Above the blots, "0" means the floxed DAG1 not mutated, "red plus" means floxed DAG1 mutated but not rescued by the viral vector encoding T315A/T317A DAG1 or T377A/T379A DAG1, and the "black plus" means floxed DAG1 mutated and rescued by the correspondence viral vectors. The blot indicates our myf5-cre system does not completely knockout DAG1 in the skeletal muscle, but the DAG1 expressed in the AAV rescued $DAG1^{L/L}$ cre+ mice are mostly introduced by the viral vector.

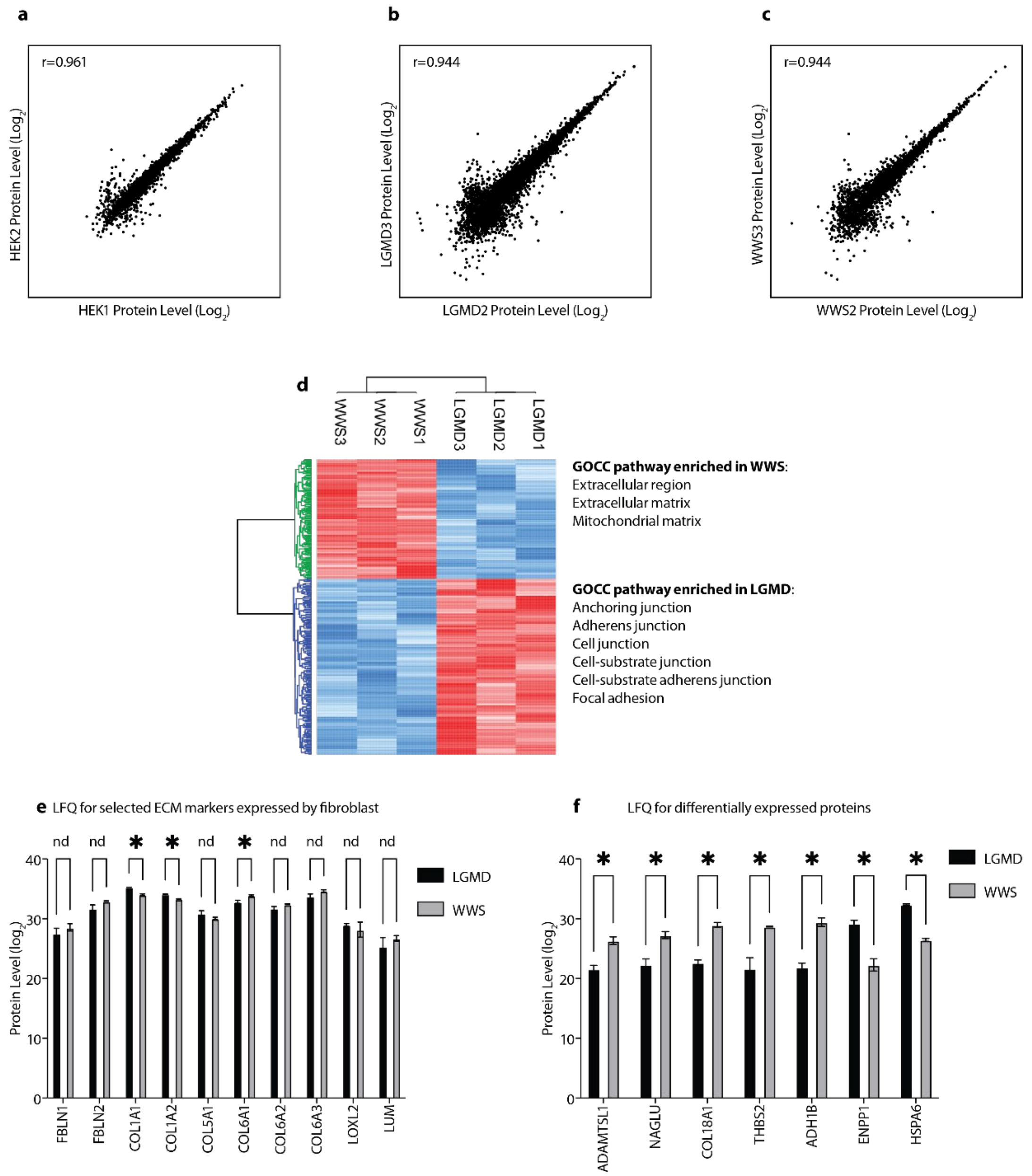


Fig.s8 Pearson correlation for duplicate proteomics of LARGE1 over-expression HEK1 cells and triplicate of LGMD (b) and WWS (c) patient fibroblasts. (d) HCA of differentially expressed proteins in WWS and LGMS patient fibroblasts. The enriched GOCC pathways are annotated at the right of the HCA figure. (e) Most fibroblast expressing ECM markers are not different between the LGMD and WWS fibroblasts. (f) T-test of differentially expressed proteins shown in **fig.7e**.
